# A proximal-to-distal survey of healthy adult human small intestine and colon epithelium by single-cell transcriptomics

**DOI:** 10.1101/2021.10.05.460818

**Authors:** Joseph Burclaff, R. Jarrett Bliton, Keith A Breau, Meryem T Ok, Ismael Gomez-Martinez, Jolene S Ranek, Aadra P Bhatt, Jeremy E Purvis, John T Woosley, Scott T Magness

**Author notes:** These authors contributed equally (Joseph Burclaff, R. Jarrett Bliton). Author Contributions: STM designed and supervised the project JB, RJB, KAB, MTO, IG, STM obtained and analyzed data RJB and KAB performed computational analysis JB, RJB, STM wrote the manuscript JTW analyzed histology slides APB and JSR provided intellectual contributions JSR and JEP reviewed computational analyses All authors reviewed the final manuscript. All data will be available in the NCBI Gene Expression Omnibus: accession number GSE185224 Python Scripts allowing for main steps of our analysis to be performed will be available on GitHub.

## Abstract

**Background and Aims:** Single-cell transcriptomics offer unprecedented resolution of tissue function at the cellular level, yet studies analyzing healthy adult human small intestine and colon are sparse. Here, we present single-cell transcriptomics covering the duodenum, jejunum, ileum, and ascending, transverse, and descending colon from 3 humans.

**Methods:** 12,590 single epithelial cells from three independently processed organ donors were evaluated for organ-specific lineage biomarkers, differentially regulated genes, receptors, and drug targets. Analyses focused on intrinsic cell properties and capacity for response to extrinsic signals along the gut axis across different humans.

**Results:** Cells were assigned to 25 epithelial lineage clusters. Human intestinal stem cells (ISCs) are not specifically marked by many murine ISC markers. Lysozyme expression is not unique to human Paneth cells (PCs), and PCs lack expression of expected niche-factors. BEST4^+^ cells express NPY and show maturational differences between SI and colon. Tuft cells possess a broad ability to interact with the innate and adaptive immune systems through previously unreported receptors. Some classes of mucins, hormones, cell-junction, and nutrient absorption genes show unappreciated regional expression differences across lineages. Differential expression of receptors and drug targets across lineages reveals biological variation and potential for variegated responses.

**Conclusions:** Our study identifies novel lineage marker genes; covers regional differences; shows important differences between mouse and human gut epithelium; and reveals insight into how the epithelium responds to the environment and drugs. This comprehensive cell atlas of the healthy adult human intestinal epithelium resolves likely functional differences across anatomical regions along the gastrointestinal tract and advances our understanding of human intestinal physiology.

## Introduction

Colloquially called the ‘gut’, the small intestine (SI) and colon are distinct organs with overlapping and unique roles in maintaining health. A monolayer of epithelium lines the gut lumen, comprised of stem and differentiated cells that renew the epithelium each week^1^. Broad health conditions develop at the intestinal epithelium, caused by pathological mucosal immunity^2^, dysregulation of carefully orchestrated cell-cell signaling, or disrupted synergy between stem cell-driven self-renewal and production of differentiated lineages. This complexity is little understood at the cellular level.

The generalized function of the gut epithelium is maintaining barrier function, absorbing nutrients, and regulating water. Cellular roles include ion balance, hormone production, mucus production, and signaling through the luminal-epithelial-immune axis. While physiological functions differ across the gut length, how lineages differ along the SI-colon axis is poorly understood. Whether adult gut epithelial lineages adopt regional fates and functions is a central question of human gut physiology and disease.

Single-cell RNA sequencing (scRNAseq) approaches have provided unprecedented transcriptomic resolution of cells and revealed unappreciated cellular heterogeneity. Studies in mouse intestines^3-5^ led to human scRNAseq studies analyzing fetal gut development^6-8^ and adult colonic^9-12^, ileal^13-15^, and duodenal^16^ epithelium. To date, one study compares adult human ileum and regionally-unspecified colon^13^, and one recent report compiles samples from across the gut yet has limited epithelial analysis^17^. Several human gut regions have sparse scRNAseq analysis available, with no studies analyzing regional differences within SI or colon. Addressing these gaps requires technically and logistically challenging approaches.

Here we comprehensively survey adult human gut epithelium using transplant-grade organs. scRNAseq libraries were prepared from epithelial cells from duodenum, jejunum, ileum, and ascending- (AC), transverse- (TC), and descending- (DC) colon from three donors. This robust cellular library avoids intra-donor batch effects and allows for observations between individual patients. Using this dataset, we probe understudied human lineages including Paneth cells (PCs), SI BEST4^+^ cells, and Follicle Associated Epithelium (FAE). We define comprehensive transcriptional signatures for lineages across the entire gut, highlight differences between human and mouse markers, and generate regional atlases of functional gene families across the proximal-to-distal axis. We further probe how lineages might be affected by extrinsic signaling by mapping receptor families and analyzing primary gene targets of approved drugs.

## Methods

### Donor Selection

Human donor intestines were received from HonorBridge (Durham, NC) with acceptance criteria: age _≤_65 years, brain-dead only, HIV(-), hepatitis(-), syphilis(-), tuberculosis(-), COVID-19(-), and no bowel surgery, severe abdominal injury, cancer, or chemotherapy. Pancreas donors were excluded to avoid duodenum loss. UNC IRB determined this study does not constitute human subjects research.

### Tissue processing

Intestines were transported on ice in University of Wisconsin Solution. Tissue dissection began within eight hours of cross-clamping. Fat/connective tissue were trimmed and intestinal regions separated: duodenum (most-proximal 20 cm); jejunum/ileum splitting remaining SI; colon split into thirds for AC/TC/DC. Two 3×3 cm mucosectomies were isolated from the center of each region for dissociation.

Mucosectomies were incubated in 10 mM NAC in dPBS at room temperature for 30 min to remove mucus, then washed in ice-cold Chelating Buffer^18^ + 100 µmol/L Y-27632. Tissues were incubated in Chelating Buffer with 2 mmol/L EDTA and 0.5mmol/L DTT, then shaken to remove crypts. High-yield colon shakes were pooled, with SI shakes pooled to approximate 1:1 villus to crypt tissue by cell mass. Crypts and villi were dissociated to single cells using 4 mg/mL Protease VIII in dPBS + Y-27632 on ice for ∼45min with trituration via a P1000 micropipette every 10 min. Cells was checked under a light microscope then filtered.

### Sample preparation

Single cells were washed with dPBS + Y-27632, resuspended in Advanced DMEM/F12 + 1% Bovine Serum Albumin + Y-27632, then stained with AnnexinV-APC (1:100) and one TotalSeq Anti-Human Hashtag Antibody (1:100) per region to track all six regions with a single library preparation^19^. Cells were washed and resuspended in AdvDMEM + 1% BSA +Y-27632 for sorting on a Sony Cell Sorter SH800Z to enrich for live single epithelial cells. 25,000 cells were collected for each region, then regions were combined pre-sequencing. Library prep was performed with the Chromium Next GEM Single Cell 3’ GEM, Library & Gel Bead Kit v3.1 (PN-100012). Sequencing was performed on an Illumina NextSeq 500.

### Data preparation and Hashtag Calling

Harmony (v0.0.5) was used to integrate the top 40 principal components from each dataset for clustering and visualization^20^. Leiden clustering was initialized with a kNN graph (k=10 neighbors) and a Leiden resolution of 0.92^21^ to resolve most expected physiological lineages. UMAPs were initialized with PAGA of identified Leiden clusters^21,22^, then non-epithelial EPCAM-negative lineages were eliminated. Regional hashtag deconvolution followed published methods: raw hashtag read counts were normalized using centered log ratio transformation followed by k-medoid clustering of hashtag read counts, with k=6 medoids for donor 1 and k=7 medoids for donors 2 and 3^19^. Hashtag noise distributions were determined by removing the highest-expressing cluster, then fitting a negative binomial distribution to the remaining cells. Cells were considered positive for a hashtag with counts above the distribution’s 99^th^ percentile (p<0.01) threshold. Cells positive for multiple hashtags were excluded as likely doublets.

## Results

### Sample processing

We define SI and colon as ‘organs’ and duodenum, jejunum, ileum, AC, TC, and DC as ‘regions’. Intestinal tracts were obtained from three organ donors (Fig. 1A,S1) with no history of cancer or intestinal disease, and healthy mucosa was verified by a pathologist (Fig. S2). Tissue was resected from each donor (Fig. 1A, Fig. S2) then epithelium was dissociated to single cells using cold protease to preserve RNA integrity. Each region’s cells were stained with Cell Hashtag antibody-oligo conjugates^8,19^ to multiplex regions for library preparation and sequencing, avoiding intra-donor batch effects and reducing cost (Fig. S4). FACS excluded dead cells and doublets (Fig. S3) prior to sequencing. After filtering for quality, transcriptional readouts for 12,590 total cells (4,330, 4,465, and 3,795 cells/donor) were obtained (Fig S1,S5).

**Figure 1:**
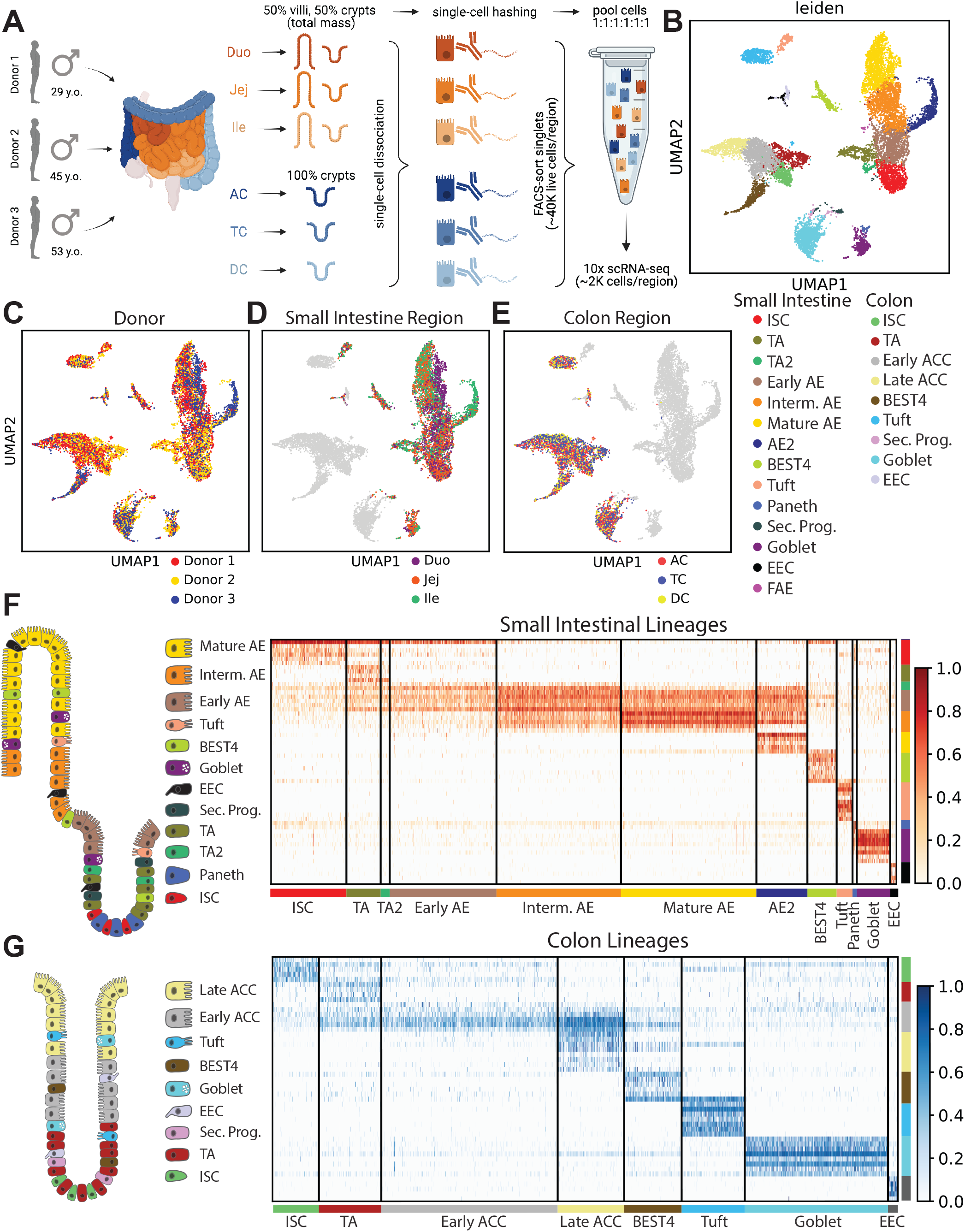
Sample Processing. **A)** Schematic for isolating cells from six intestinal regions for three donors and using hashtag antibodies to sequence regions side-by-side. **B)** UMAP of analyzed cells in 25 lineage clusters. **C-E)** UMAP of all cells by donor (C) or region (D,E). **F-G)** Heatmaps showing unique markers for major lineages in SI (F) and colon (G).

Donor sequencing datasets were individually processed then combined, with principal components integrated with Harmony^20^ to minimize donor-specific differences prior to dimensional reduction and Leiden clustering^21^. Most lineages formed distinct SI/colon-specific clusters, suggesting functional differences by location. No organ-specific clustering occurred for enteroendocrine cells (EECs) and secretory progenitors (Fig. S6). One cluster expressed PC and goblet cell (GC) markers, so sub-clustering resolved these lineages (Fig. S6). Our final dataset identifies all lineages by organ along with rare FAE (Fig. 1B). The integrated dataset shows overlapping cell distributions from each donor and region within all major lineages, demonstrating that post-sequencing hashtag deconvolution preserves transcriptomic features across samples and batches (Fig. 1C-E).

To define significant marker genes across the gut, we calculated differentially expressed genes (DEGs) in each lineage versus all other cells from each organ. We identified DEGs consistently enriched across all three donors and the combined dataset. Through this rigorous statistical evaluation, we developed a unique signature for all major lineages across the human SI and colon epithelium, a previously unavailable resource (Fig. 1F,G, Table S1,S2).

### Proliferative Cells

We found human Intestinal Stem Cells (ISCs) differentially expressed classical markers *LGR5, ASCL2, SLC12A2*, and *RGMB* (Fig. 2A,B)^23-25^. Though enriched, *SMOC2*^26^ was not a DEG in SI ISCs, as PCs express higher levels (see PCs). While *in situ* hybridization showed *OLFM4* marks human SI and colonic ISCs^27^, our data indicate significantly higher levels of *OLFM4* in SI ISCs, with colonic *OLFM4* higher in transit amplifying (TA) cells and early absorptive colonocytes (ACCs) (Fig. 2A,B). This concurs with mouse, where *Olfm4* transcripts marked SI but not colon ISCs^24^. Notably, *RARRES2* was enriched in colon ISCs, with low expression in SI ISCs (Fig. 2B). We found no gut-related literature on this retinoid response gene, providing an intriguing target for future studies.

**Figure 2:**
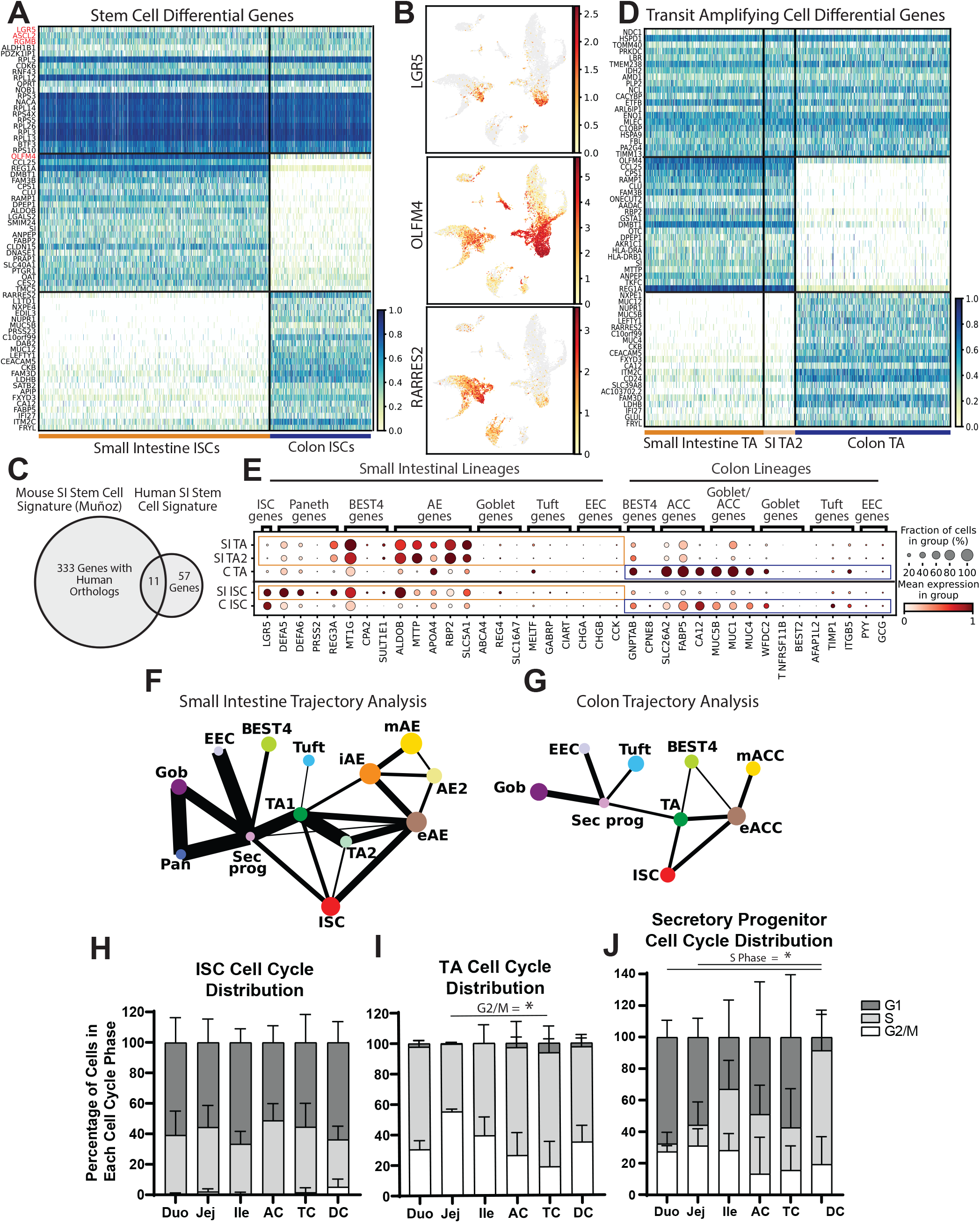
Proliferative crypt populations. **A)** Heatmap of DEGs in ISCs vs. other lineages (top; red: classical markers), SI vs. colon ISCs (middle), colon vs. SI ISCs (bottom). **B)** UMAP of *LGR5, OLFM4*, and *RARRES2* expression. **C)** Venn Diagram showing overlap between murine and human ISC signatures. **D)** Heatmap of DEGs in TA cells vs. other lineages (top), SI vs. colon TA cells (middle), colon vs. SI TA cells (bottom). **E)** Dotplot showing DEGs from SI or colon-specific lineages within organ-delineated ISCs and TA cells. **F**,**G)** Partition-based graph abstraction (PAGA) showing connectivity between major lineages in SI (F) and colon (G) to infer maturation trajectory. Line thickness represents connectivity strength. **H-J)** Regional cell cycle phase distribution in ISCs (H), TA cells (I), and secretory progenitors (J).

We constructed a comprehensive human ISC signature. SI ISCs had 68 DEGs compared to other clusters across SI regions for all three donors, whereas colon ISCs displayed 109 DEGs (Table S1, Fig. S7). To define a human ISC transcriptional signature spanning SI and colon, we identified 46 DEGs enriched in ISCs from both organs (Table S2). This signature includes classical ISC markers along with 30 ribosomal genes. While ribosomal genes are less abundant in murine ISC signatures^26,28,29^, this is consistent with literature describing ribosomal control of transcriptional dynamics in other stem cells^30-32^. To identify ISC DEGs conserved between human and mouse, we compared our 68-gene SI ISC signature with a mouse signature with 344 human homologs^26^. Surprisingly, only 11 genes overlapped between the human and mouse signatures (Fig. 2C), although it is unclear whether this reflects species differences or the higher resolution and stringency of our computational approach. Conserved genes included classical markers: *LGR5, OLFM4, ASCL2, RGMB, SLC12A2*, and *MYC*; genes with known ISC function: *RNF43, ZBTB38, VDR, and CDK6*; and one gene not described in ISC literature: *TRIM24*, a gene involved in p53 degradation. These SI, colon, and full-gut ISC signatures underline key similarities and differences in proximal-distal human ISCs.

TA cells are classically defined by high proliferation and location above ISCs in the intestinal crypt^9,33^.Leiden clustering separated SI TA cells undergoing S/G2 cell-cycle phases (TA) and M-phase (TA2) (File S1,S2). We evaluated DEGs between SI TA, TA2, and colon TA cells to define shared TA markers (Fig. 2D). DEGs were involved in cell cycle as well as in mitochondrial biogenesis and rRNA processing, consistent with increased mitochondrial load and translation levels seen as stem cells differentiate in various systems^30,34-36^.

Several organ-specific markers of differentiated lineages (Fig. S8) were unexpectedly enriched in their respective SI or colon ISC and TA populations (Fig. 2E), hinting that ISCs are already transcriptionally primed for organ-specificity, instead of existing in a naïve, pan-intestinal state. This is consistent with rodent studies showing adult SI ISCs produce daughter cells specific to their originating organ when engrafted into alternative SI or colon sites^37,38^. Studies defining region/organ-specific chromatin or active transcriptomic differences in ISCs were not found; thus, these genes may prove useful for studying differentiation and chromatin dynamics in early fate-determination.

Trajectory analyses computationally investigate lineage transitions, with intestinal analyses primarily using mouse data^39-42^. We used Partition-based Graph Abstraction (PAGA), which estimates connectivity between clusters, to infer connections between proliferative crypt-based lineages and differentiated populations^22^. Absorptive enterocytes (AEs) and ACCs arise nearly exclusively from ISCs and TA cells^15^. The secretory progenitor population arises from ISCs and TA cells and gives rise to PCs, GCs, and EECs (Fig. 2F,G). Interestingly, tuft cells appear to derive from secretory progenitors in colon but not SI, consistent with murine findings^39^. Conversely, SI BEST4^+^ cells apparently arise from secretory progenitors while colon BEST4^+^ cells connect to TA cells, agreeing with absorptive and secretory functions ascribed to this novel lineage^12,16^.

Predicted regional cell cycle phase distributions^43^ were analyzed in ISCs, TA cells, and secretory progenitors (Fig. 2H-J). ISCs showed expected high G1 and S phase representation across regions^13,44^, while highly-proliferative TA cells largely existed in S and G2/M. TC showed a significantly lower proportion of TA cells in G2/M than jejunum, with similar regional differences in rodents^45,46^, but biological implications are unknown. Secretory progenitors had notably different distributions, with S phase proportions increasing proximally-to-distally and higher G1 proportion than TA cells (Fig. S1). Since secretory progenitors differentiate into specialized lineages, elongated G1 may allow for additional reception of differentiation factors, as stem cells are most receptive to such cues during G1^47^.

### Paneth Cells

Murine PCs play important niche-supporting and antimicrobial roles^48^, yet little scRNAseq analysis covers human PCs. Our dataset includes 49 PCs representing all SI regions across three donors, 10-times more than analyzed in recent literature^16^. PCs were defined using *DEFA5, DEFA6, ITLN2*, and *PLA2G2A* (Fig. 3A). Surprisingly, Lysozyme (*LYZ*), an important murine PC marker, was expressed higher in SI BEST4^+^ cells and FAE, with measurable tuft cell expression, making *LYZ* an imprecise human PC marker (Fig. 3B). This is consistent with *LYZ* expression in organoids derived from a fetal human stage too young to form PCs^6^. Since PCs cluster alongside GCs and share *LYZ* expression with BEST4^+^ cells, classical PC markers were plotted to confirm PC identify (Fig. 3C). Importantly, our data indicate the cells described as PCs in a recent scRNAseq publication^13^ are BEST4^+^ cells, with high *LYZ, SPIB, BEST4*, and *CA7*. Similarly, the reported colonic ‘Paneth-Like Cells’ in the study are likely BEST4^+^ cells unrelated to PCs other than *LYZ* expression. The rarity of PCs (<1% of our dataset and others^16^) and divergence of PC and BEST4 markers highlights the precise lineage attribution needed when defining human PCs.

**Figure 3:**
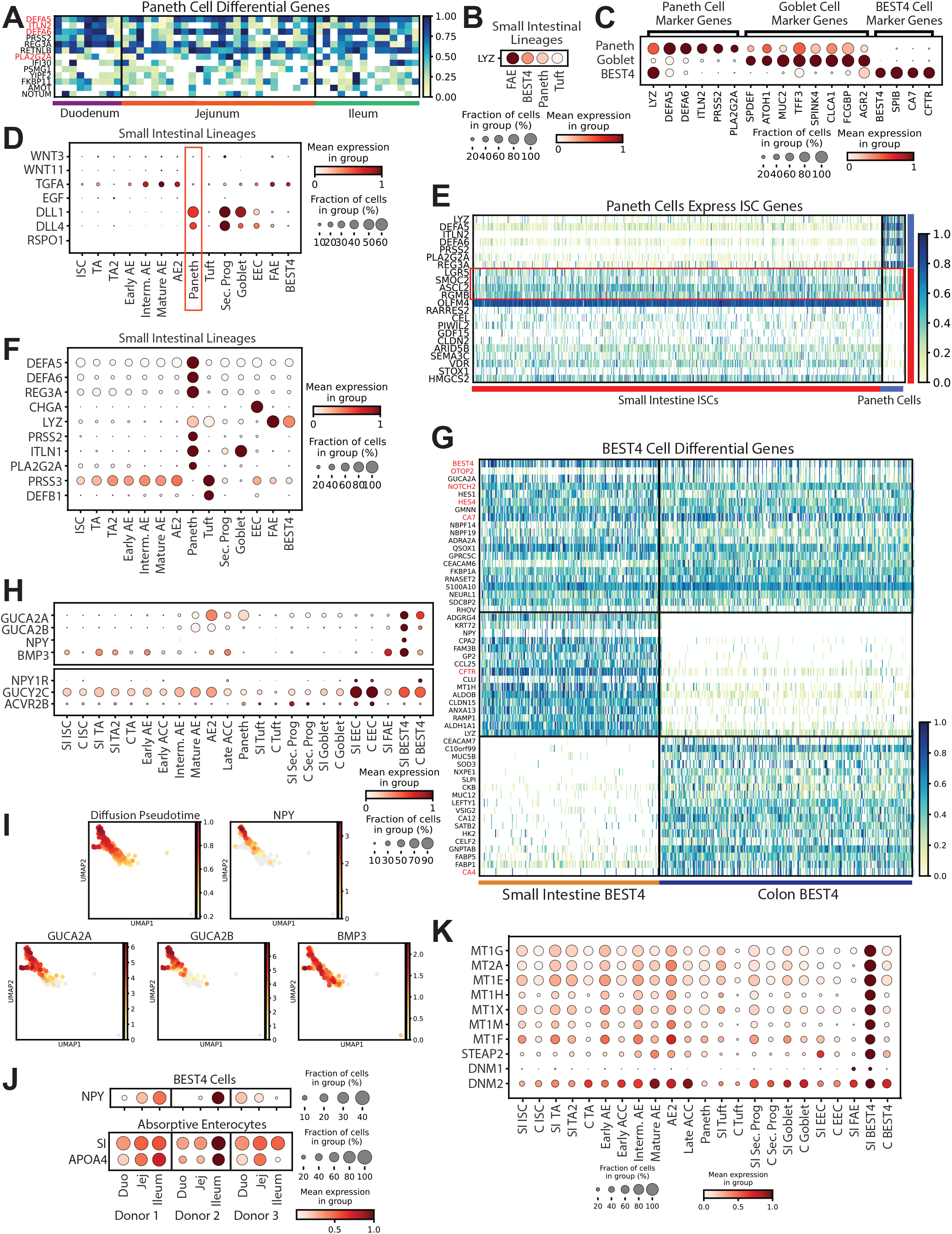
Paneth and BEST4^+^ cells. **A)** Heatmap of DEGs in PCs vs. other lineages (red: classical markers). Dotplot showing Lysozyme across FAE, BEST4, Paneth, and tuft cells. **C)** Dotplot showing PC, goblet, and BEST4 markers. **D)** Dotplot showing growth factors implicated in murine PCs across SI lineages. **E)** Heatmap showing PC and ISC markers across both lineages. **F)** Dotplot showing 10 highest-expressed antimicrobial peptides across SI lineages. **G)** Heatmap of DEGs in BEST4^+^ cells vs other lineages (top; red: classical markers), SI vs. colon BEST4^+^ cells (middle), colon vs. SI BEST4^+^ cells (bottom). **H)** Dotplot showing secreted genes and their receptors across lineages. **I)** UMAPs of BEST4^+^ cells showing predicted diffusion pseudotime and expression of secreted peptides. **J)** Expression of *NPY, SI*, and *APOA1* across regions by donor. **K)** Dotplot showing genes involved in metal-binding and endocytosis across lineages.

We next probed whether human PCs express ISC niche factors. Murine PCs express *Wnt3, Wnt11, Tgfa, Egf, Dll1, Rspo1, and Dll4*^40,48-50^, but one report shows human PCs express no *WNT3/11*^*16*^. Our data confirm this and demonstrate no measurable *EGF* or *RSPO1* and minimal *TGFA*. While human PCs express *DLL1* and *DLL4*, both are higher in secretory progenitors (Fig. 3D). We found no members of any major intestinal growth factor family broadly enriched in PCs (Fig. S9), suggesting human PCs are not major niche-supporting cells. This notion is consistent with non-epithelial sources of WNTs and growth factors in the human niche^6,51-53^, and echoes mouse biology, where PCs are sufficient to support ISCs^48^ yet unnecessary for niche maintenance^48,54-56^.

Unexpectedly, *SMOC2*, a murine ISC marker,^*26*^ was expressed highest in PCs, with other mouse ISC-restricted markers (*LGR5, ASCL2, RGMB)* higher in human PCs than mouse PCs^5^ (Fig. 3E). ISC-PC doublet artifacts could explain this, however lack of ISC markers (e.g., *OLFM4)* does not support this hypothesis. *LGR5, SMOC2*, and *ASCL2* are involved in WNT reception^23,57-59^, suggesting human PCs may receive WNT signals instead of providing WNT signals as in mice^48^. PC-unique expression of *FZD9* supports a WNT-receptive PC role^60^ (Fig. S9), with its expected ligand WNT2^61^ absent in our database yet induced in intestinal inflammation and cancer^62-64^. Mature PCs can dedifferentiate following injury in mice^65-67^, and *Ascl2* is required for dedifferentiation in mouse crypts^68^. Thus, expression of these ISC genes may render human PCs more responsive to dedifferentiation cues; however, this needs functional validation.

A murine subset of colonic GCs termed Paneth-Like Cells or Deep-Crypt Secretory Cells are defined by MUC2, C-KIT, REG4, CD24, EGF, and FZD5^69,70^. We found only two cells positive for both *REG4* and *KIT* across the 1,252 colon GCs and secretory progenitors (Fig. S9D). Colonic tuft cells expressed higher *KIT* and *CD24* but no *REG4*. Based on murine-defined markers, we conclude there is no human equivalent to this population. Since human PCs likely perform little niche-supporting activity, namely not producing *EGF*, human cells with murine PC-like functions may be unnecessary.

Despite striking differences between mouse and human PCs, both supply antimicrobial gene products. Six of the 10 highest-expressed SI antimicrobial peptides are PC-enriched (Fig. 3F). As antimicrobial genes comprise half of human PC DEGs (Table S1), they likely function primarily to protect the ISC-niche from bacterial invasion and regulate microbiota composition^71^.

### Follicle-Associated Epithelium

Rare FAE cells, important for epithelial-immune crosstalk, reside in small puncta throughout the intestines^72^. FAE includes microfold (M)-cells, which transport luminal antigens to immune cells residing within their microfolds^73^. M-cells have almost exclusively been explored in mice^74-76^ or using directed differentiation in vitro^77^, with only one study reporting scRNAseq data from healthy human intestinal M-cells^17^. Our dataset includes a cluster of 19 cells from Donor 2 enriched for M-cell markers^78-80^ and immune cross-talk genes (Fig. S10). Strikingly, several murine M-cell-specific markers were either widely expressed (*MARCKSL1, ANXA5, CXCL16*)^79,81^ or absent (*CCL36, SCG5, TNFRSF9, CCL9, CCL6, PGLYRP1*)^73,79,82^, suggesting species functional differences. With the caveat of including 19 cells from one donor, we defined 145 DEGs (Table S1,S2), finding many FAE-unique genes (Fig. S10). Pathway enrichment analysis implicates these DEGs in immune cell interactions, verifying expected M-cell function (File S3). Differences in chemokines between mouse and human M-cells, with CCL20 the only major shared chemokine, call for enriching for FAE using newly described methods^72^ to probe how human M-cells interact with immune cells.

### BEST4^+^ cells

Recent human single-cell studies describe a novel intestinal lineage, absent in mice, expressing high *BEST4, SPIB, and CA7*^*12*^, with *CFTR* in SI^16,17^ and *OTOP2* in colon. Several papers describe colonic BEST4^+^ cells^11,12^, so we analyze SI BEST4^+^ cells. With BEST4^+^ cell functions largely unknown, DEGs were used to predict physiological roles (Fig. 3G). DEG analysis revealed secreted peptides including *GUCA2A* and *GUCA2B*^*12*^, which can act as pro-hormones effecting a satiety response^83-85^. A previous report showed these genes in SI and colon BEST4^+^ cells^16^, yet we find both expressed higher in SI than colon BEST4^+^ cells (Fig. 3H), indicating a role in satiety signaling in the SI.

We identified two unreported secreted peptides, *NPY* and *BMP3*, specifically in SI BEST4^+^ cells. *NPY* expression was unexpected in intestinal epithelium^86^, and gut *BMP3* is largely studied for antitumor roles^87,88^. Diffusion pseudotime, a computational method to define single-cell differentiation trajectories^89^, indicated that *NPY, GUCA2A*, and *GUCA2B* expression increased with BEST4^+^ cell maturation, while *BMP3* expression appeared independent of maturation (Fig. 3I). Interestingly, receptors for all four genes are enriched in EECs, suggesting cross-talk between these lineages (Fig. 3H).

Since *NPY* is proposed to affect gastrointestinal (GI) motility^90^ and energy homeostasis^91^, we probed if NPY correlated with genes induced following meals. Looking across SI regions for each individual donor, we found a strong positive correlation between SI BEST4^+^ cell *NPY* and AE expression of *SI* (R=0.82) and *APOA4* (R=0.86), which are induced in mice by dietary sugar^92^ and fat^93^ (Fig. 3J). We found further positive correlations with AE genes involved in dietary metabolism (Table S3), and negligible correlation with housekeeping genes *ACTB* (R=-0.22) or *GAPDH* (R=-0.08). Thus, *NPY* expression in SI BEST4^+^ cells may be induced by luminal contents. Further DEGs from SI BEST4^+^ cells included adrenergic and cholinergic neurotransmitter receptors involved in intestinal motility^94^, *ADRA2A* and *CHRM3* (Table S1), supporting that SI BEST4^+^ cells may regulate intestinal motility following meals.

BEST4^+^ cells likely absorb dietary heavy metals. Metallothionein expression has been implicated in colonic BEST4^+^ cells^12^, yet we find seven members of this family, known to bind heavy metals and protect against toxicity^95-97^, specifically enriched in SI BEST4^+^ cells (Fig 3K). Unique expression of *STEAP2*, a metalloreductase for copper and iron^98^, supports a role for SI BEST4^+^ cells in maintaining SI homeostasis for many metal ions^97-99^. SI BEST4^+^ cells express endocytosis effector genes *DNM1* and *DNM2* (Fig. 3K), supporting an absorptive role for these cells. Our data indicate BEST4^+^ cells perform diverse roles within the intestinal epithelium, laying the groundwork for functional studies.

### Tuft Cells

Tuft cells are chemosensory epithelial cells which regulate type-2 immune reactions in the intestinal epithelium through pathogenic metabolite detection and classical taste signal transduction pathways^100-102^. Together, these initiate tuft cell IL-25 release^103-105^. SI and colon tuft cells share many classical markers^16,106^ (Fig. 4A, Table S1,S2). *DCLK1*, a key murine marker^105^, was not observed. Interestingly, *SUCNR1*, a G-protein coupled receptor mediating SI IL-25 release^107^, was negligible in colon, suggesting SI and colon tuft cells differentially detect and respond to luminal succinate-producing pathogens (Fig. 4B,C). Instead, colonic tuft cells likely respond to umami-chemosensory cues, such as microbe-derived free amino acids, as they express heterodimeric umami taste receptor subunits *TAS1R1* and *TAS1R3* (Fig. 4B,C)^108^. We further analyzed downstream taste signal transduction genes, finding SI and colon tuft cells enriched in critical pathway components (*GNB1, GNG13, ITPR2, TRPM5*)^100,101^, with SI-specific *GNAT3*, a G-protein alpha subunit, which likely activates *PDE4D* to decrease intracellular cAMP/cGMP^109^ (Fig. 4B). This suggests human SI tuft cells have varied responses to succinate-producing microbes (e.g., *N. brasiliensis*), whereas colonic tuft cells may respond more broadly to other microbial taxa.

**Figure 4:**
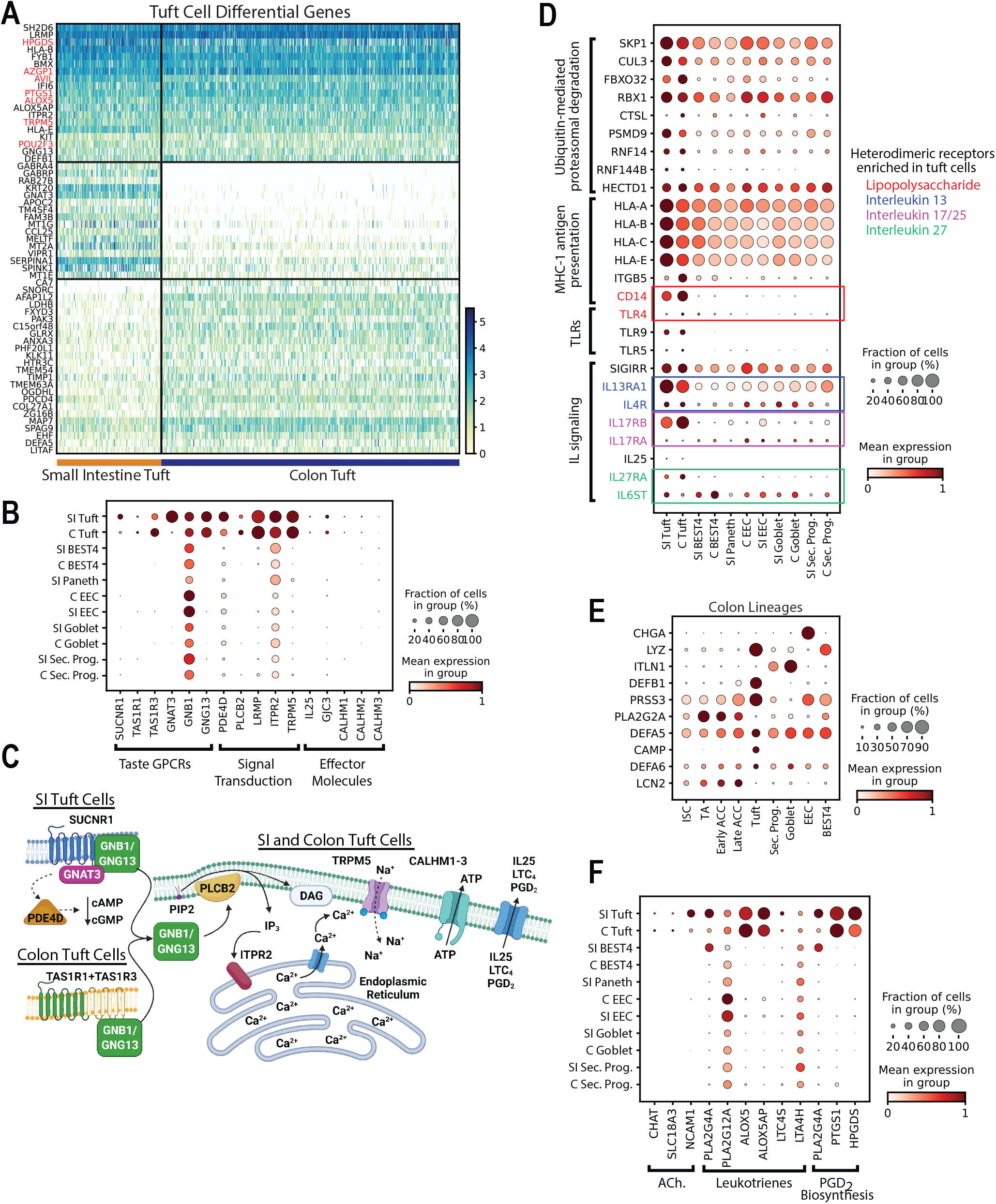
Tuft Cells. **A)** Heatmap of DEGs in tuft cells vs. other lineages (top; red: classical markers), SI vs. colon tuft cells (middle), colon vs. SI tuft cells (bottom). **B)** Dotplot showing tuft cell enrichment of genes specific to taste signal transduction. **C)** Organ-specific signal transduction in SI vs. colon tuft cells. **D)** Dotplot showing tuft cell-enriched genes enabling interactions with innate and adaptive immune system. **E)** Dotplot showing 10 highest-expressed antimicrobial peptides across colon lineages. **F)** Dotplot showing tuft cell-specific genes for producing acetylcholine, eicosanoids, and prostaglandins.

Beyond triggering type-2 immunity, tuft cell DEGs allow broad interaction with the adaptive and innate immune systems. Tuft cells express DEGs involving ubiquitin-mediated proteasome degradation, including SCF complex components (*SKP1, CUL3, FBXO32, RBX1*) which initiate processing of exogenous antigens for presentation^110,111^, as well as genes for the MHC1 antigen presentation complex (Fig. 4D). This suggests tuft cells may interact with the adaptive immune system following luminal stimuli. Human tuft cells also uniquely express previously unappreciated Toll-Like Receptors –*TLR9, TLR5*, and *TLR4* – which bind bacterial and viral DNA, bacterial flagellin, and lipopolysaccharide (LPS), respectively^112-114^ (Fig. 4D). Expression of the LPS coreceptor *CD14* across tuft cells (Fig. 4D) supports a novel role in bacterial-related immune responses^112^.

Tuft cells exhibit possible auto-regulatory mechanisms for these pathogen-response pathways. Tuft cells express heterodimeric IL-25-specific receptor components *IL17RA* and *IL17RB* (Fig. 4D) which may create a positive feedback loop to amplify IL-25 signaling, as in keratinocytes^115^. Tuft cells may also autoregulate their LPS response through *SIGIRR*, which negatively regulates TLR4*-*LPS signaling^116-118^. These implicate tuft cells as dynamic sentinels linking luminal contents to the immune system.

Consistent with a role regulating gut pathogens, tuft cells produce *LYZ, PRSS3, and DEFB1* antimicrobial peptides in the SI (Fig. 3F) and six of the top ten antimicrobial peptides in the colon, without PCs present (Fig. 4E). Finally, human and murine tuft cells both produce neuro-and immunomodulatory compounds. We find genes necessary for acetylcholine synthesis (*CHAT, SLC18A3/VACHT*), communication with neurons (*NCAM1*)^119^, and enzymes involved in eicosanoid production, namely cysteinyl leukotrienes (*ALOX5* and *ALOX5AP*) and Prostaglandin D_2_ (*PTGS1* and *HPGDS*), which broadly regulate inflammation^120^ (Fig. 4F). Altogether, these genetic analyses indicate tuft cells regulate luminal microbes, communicate with the nervous system, and effect systemic immune responses.

### Goblet Cells

GCs produce membrane-bound and secreted mucin glycoproteins that lubricate the gut, act in signaling, support commensal bacteria, and form the protective mucus barrier^103,121-123^. DEGs include classical markers *CLCA1, MUC2*, and *TFF3*, with colon GCs expressing higher mucins (*MUC5B, MUC4, MUC1*) and the antiprotease *WFDC2*^12^ (Fig. 5A). Pathway enrichment analysis of DEGs confirms GCs principally act in mucus secretion, with 15 of 20 top enriched pathways involved in mucus production, including glycosylation, Golgi/ER vesicle transport, and unfolded protein response (File S3)^124-127^. Thus, we mapped regional GC mucin expression (Fig. 5B), finding secreted *MUC2* and transmembrane *MUC13* expressed across all regions and colon-enriched *MUC1, MUC4*, and *MUC5B*. While GCs are the major mucus-secreting intestinal lineage, AE and ACC express transmembrane mucins that form a glycocalyx to protect against pathogenic bacteria^128,129^. Regional mucin expression across these lineages showed high *MUC13, MUC17*, and *MUC3A* in AEs and several enriched in ACCs (Fig. S11), informing studies regarding mucus composition and function across GCs, AEs, and ACCs.

**Figure 5:**
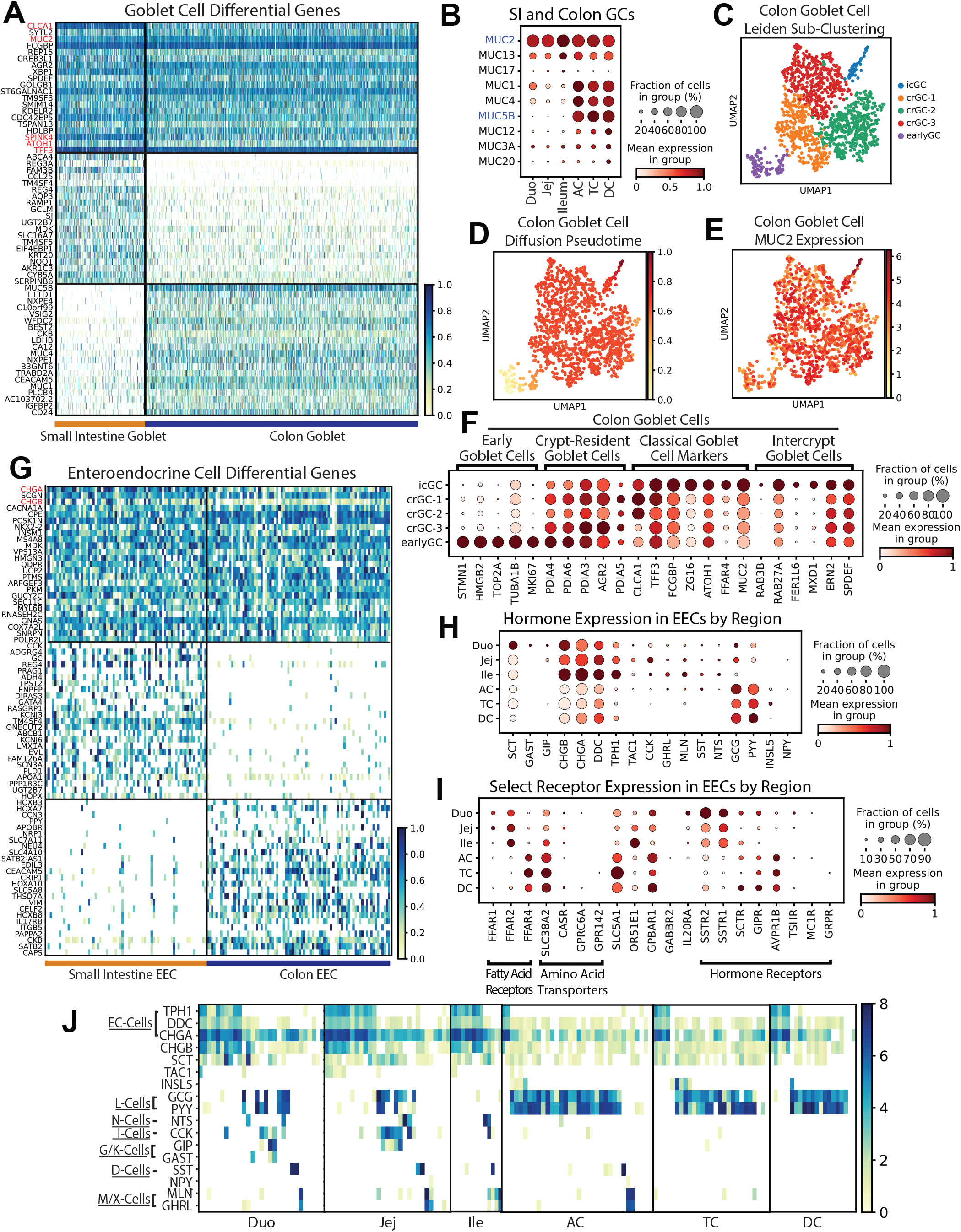
Goblet and Enteroendocrine Cells. **A)** Heatmap of DEGs in GCs vs. other lineages (top; red: classical markers), SI vs. colon GCs (middle), colon vs. SI GCs (bottom). **B)** Dotplot showing 9 highest-expressed mucins across GCs by region (blue: gel-forming mucins). **C)** Leiden sub-clustering of colon GCs. **D)** Diffusion pseudotime of colon GCs. **E)** UMAP of *MUC2* expression in colon GCs. **F)** Dotplot showing mouse-implicated markers in human colon GC subclusters. **G)** Heatmap of DEGs in EECs vs. other lineages (top, red: classical markers), SI vs. colon EECs (middle), colon vs. SI EECs (bottom). **H)** Dotplot of EEC regional hormone gene expression. **I)** Dotplot of EEC expression of select receptors by region. **J)** Heatmap showing hormone expression in each EEC.

GCs are commonly considered fairly homogenous, yet recent work in mouse colon reported transcriptional signatures of early GCs, crypt-resident (crGC), and inter-crypt goblet cells (icGC), with icGCs producing more permeable mucus than crGCs^130^. Human colonic secretory progenitors and GCs subclustered into similar groups marked by genes implicated in mouse GC heterogeneity: early GCs (*STMN1, HMGB2, MKI67*), crGCs (*PDIA3, AGR2,PDIA5*), and icGCs (*MXD1, RAB27A, FER1L6*) (Fig. 5C,F). Some canonical GC markers of mucus secretion (*MUC2, ZG16*) were expressed highest in icGCs, consistent with icGCs constitutively secreting mucus^131^. Diffusion pseudotime confirmed increasing maturity across these sup-populations (Fig. 5D-E). Notably, crGCs expressed higher *MUC5B* while icGCs expressed higher *MUC2, MUC13, MUC1*, and *MUC4* (Fig. S11C), consistent with distinct mucus production in human icGCs shown via lectin staining^130^. Similar sub-clusters were observed in the SI, although mucin differences were less obvious (Fig. S11D-F). This demonstrates human GCs are more heterogeneous than appreciated, necessitating studies to determine functional differences.

### Enteroendocrine Cells

EECs secrete hormones to communicate between the intestine and the body. EEC hormone expression profiles are well characterized at the single-cell level in mice, since EEC reporter models enable enrichment of this rare lineage (<1% of intestinal epithelium)^132,133^. However, transcriptomic differences exist between mouse and human EECs^132,134^. An approach in human organoids with an EEC reporter gene yielded sufficient EECs for detailed scRNAseq analysis, though it is unclear how these cells may differ from primary human EECs^132^. While several human scRNAseq studies show EEC data^11-13,16,17^, our 154 EECs (Fig. 5G) represent the largest dataset of primary human EECs to our knowledge.

First, regional expression of hormones and other signaling machinery in EECs was surveyed. We found *SCT, CHGA, TPH1*, and *DDC* span regions with a SI bias and *GCG* and *PYY* span regions with a colonic bias (Fig. 5H). *GAST* and *GIP* express in proximal SI, while *TAC1, CCK, GHRL, MLN, SST*, and *NTS* express from duodenum through AC/TC, and *INSL5* is colon-specific. Rare *NPY* expression was detected in jejunum and AC. These results expand on an early study using immunohistochemistry in regional biopsies to find CCK, GAST, GIP, NTS, MLN, and SCT segregated to SI regions^135^. Our data confirm the SI bias but show low colonic expression of *CCK, NTS, MLN*, and *SCT*, demonstrating the higher sensitivity in scRNAseq. *NTS* and *CCK* were also absent in a study analyzing region-unspecified colon^13^, showing the importance of analyzing all colon regions. Fatty-acid receptors *FFAR1* and *FFAR2* were enriched in SI EECs and *FFAR4* expression was colon-specific (Fig. 5I). The amino acid transporter *SLC38A2* was well-expressed, with lower expression of other amino acid transporters. EECs also express several hormone receptors, indicating crosstalk amongst EECs. An additional form of gut-brain crosstalk was recently discovered in mice, with EECs found to form synapses with the vagus nerve^136-138^. DEGs from SI and colon EECs (Fig S1,S12A) are consistent with a human equivalent of these mouse EECs, termed neuropods^137^, with 33.7% of genes in the GOCC_Presynapse list expressed highest in EECs (Fig. S12B). Altogether, these patterns support roles for EECs in crosstalk within the gut and between the gut and brain, further illuminating their functional importance.

EECs are classified into subtypes based on hormone expression^139,140^. A regional breakdown of individual EECs was constructed to visualize EEC subtypes across the gut (Fig. 5J). Enterochromaffin cells (*TPH1, DDC, CHGA*) appear in each region with fewer in colon, and ileum unexpectedly lacks L cells (*GCG, PYY*). Multiple EECs express 8-10 hormones, expanding on studies identifying poly-hormonal EECs^141,142^. Despite low numbers, *GAST* and *GIP* largely segregate from duodenal L cells yet overlap in jejunum. We note rare *NPY* expression in *MLN*^*+*^ and *GHRL*^+^ EECs in jejunum and AC. Future studies combining our primary EECs with additional datasets from various regions would improve our understanding of human EECs.

### Absorptive Enterocytes and Colonocytes

AEs and ACCs perform nearly all intestinal absorption^143^. These lineages formed multiple Leiden clusters, with three AE clusters and two ACC clusters consistent with increasing maturity, reflecting other reports^11,16^, and one cluster (AE2) separate from other AEs, largely from Donor 3 ileum (Fig. 1C,6B). A common DEG signature was defined by grouping all AEs and comparing their DEGs with all ACC DEGs. Surprisingly, only five DEGs were shared between organs (Fig. 6A), indicating stark organ differences. Interestingly, the novel AE2 population expressed mature AE markers (Fig. 6B), while uniquely expressing genes for bile acid uptake and processing^144-146^ (Fig. 6C). It is unclear why ileal AEs of Donor 3 clustered separately from other donors. Possible donor-specific factors contributing to this novel cluster include unique demographics (lowest BMI, African American, Type II diabetic) or differences in luminal contents between donors possibly inducing unique gene expression patterns, as described for certain genes^92,93^. These donor-specific differences in transcriptomic signatures highlight the need for data from wide, diverse populations.

**Figure 6:**
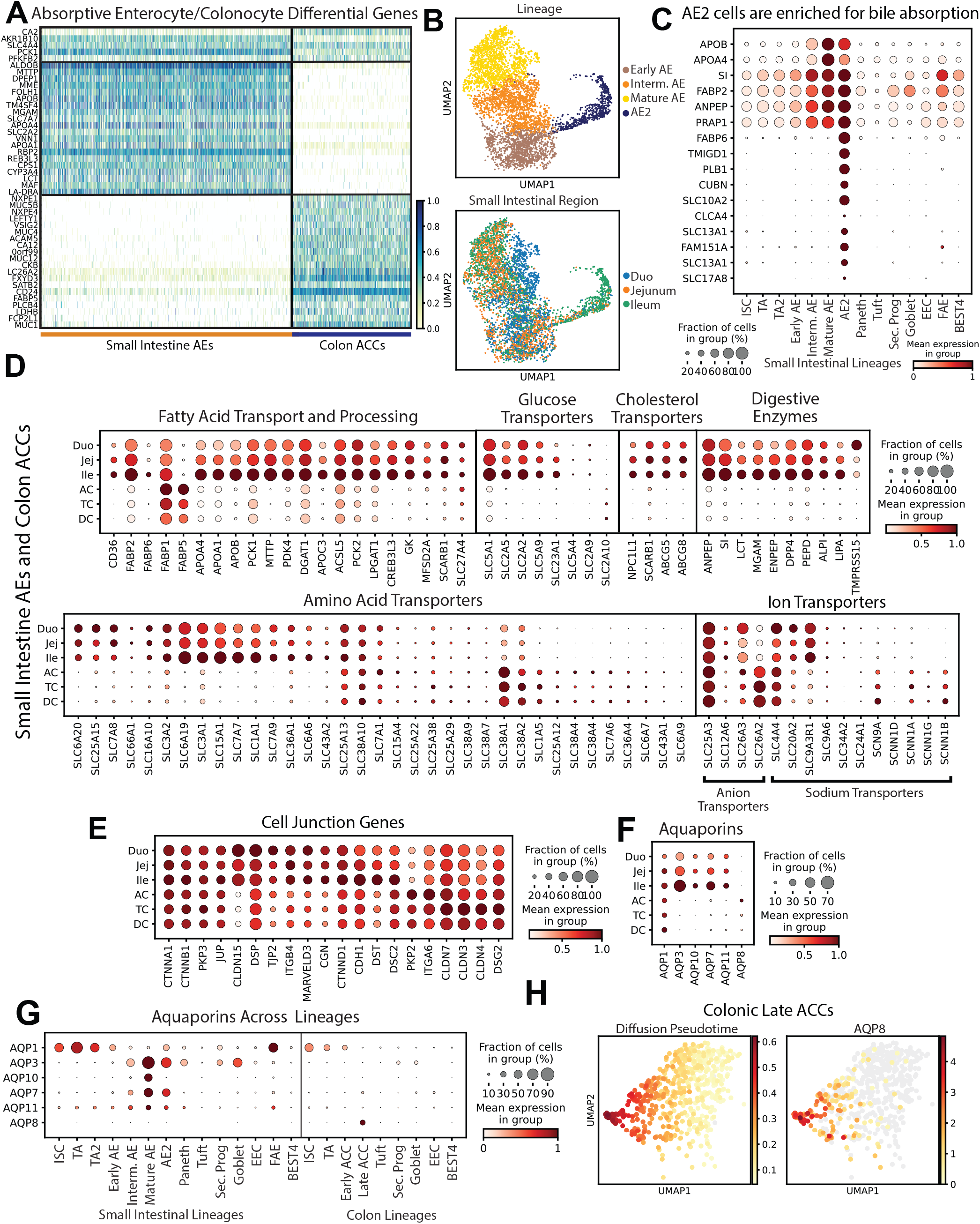
Absorptive cells. **A)** Heatmap of DEGs in absorptive cells vs. other lineages (top) AEs vs. ACCs (middle), ACCs vs. SI AEs (bottom). **B)** UMAPs showing AE2 Leiden cluster (top) and cells by region (bottom). Dotplot of classical Mature AE markers and top 10 DEGs for AE2 cluster. **D)** Dotplots showing regional expression of genes involved in digestion and absorption in all AEs and ACCs. **E)** Dotplots showing 20 highest-expressed cell junction genes in AEs and ACCs by region. **F)** Dotplots showing regional aquaporin expression in AEs and ACCs. **G)** Aquaporin expression across lineages. **H)** UMAPs of late ACCs showing predicted diffusion pseudotime (left) and *AQP8* expression (right).

Macro- and micro-nutrient handling was mapped across all AEs and ACCs (Fig. 6D). Nearly all fatty acid, glucose, and cholesterol transporters enriched in SI, consistent with recent work^13^, but regional data revealed an undescribed trend of increasing expression from duodenum through ileum for most genes (Fig. 6D). Two notable exceptions were *FABP1* and *FABP5*, both with higher colon expression. *FABP1* is expected in the proximal SI^147^ and *FABP5* is not implicated in healthy ACCs to our knowledge, suggesting a possible role for colonic fatty acid uptake. Digestive enzymes exhibited ileal enrichment except for the duodenum-specific serine protease *TMPRSS15*/Enteropeptidase^148^. More regional variability was seen for amino acid transporters, with different genes peaking from duodenum through colon. The neutral amino acid transporter *SLC38A1* expressed in colon, in contrast to a study using colon biopsies^149^. Ion transporters showed the most regional differences, with *SLC25A3* and *SLC4A4* spanning all regions, colon-enriched *SLC26A2*, and SI-enriched *SLC9A3R1*. Finally, *SCNN1* sodium transporter subunits were largely enriched in colon, consistent with its need for regulated water uptake. This regional map of nutrient handling genes expands upon previous organ-level analyses, emphasizing the importance of the ileum in digestion.

Intestinal barrier function, largely conferred by cell-junction proteins, is essential for regulated absorption and physical antimicrobial defense^150^. The 20 highest-expressed cell junction genes were regionally evaluated (Fig. 6E). Several junction genes expressed equally across AEs and ACCs, while others exhibited regional enrichment. Claudins (*CLDN*) are the primary determinants of tight junction barrier function and intestinal epithelial integrity^150,151^. *CLDN1* and *CLDN15* were SI-enriched and *CLDN3, CLDN4*, and *CLDN7* were highest in TC. Notably, no junction genes expressed highest in DC. This is intriguing, as ulcerative colitis often originates in the distal large intestine, raising the possibility that higher junction protein expression in AC and TC might protect against certain inflammatory conditions^152-154^. While our data cannot show pathological implications, the possibility that this differential expression pattern may protect against colitis in AC and TC is intriguing.

Aquaporins (AQPs) are the major transcellular transporters of water and small solutes in the intestine^155^. Our data confirms a previous report showing elevated *AQP3, AQP7*, and *AQP11* in ileum relative to colon and *AQP8* elevated in colon (Fig. 6F)^13^, yet we find *AQP1* widely expressed. Aquaglyceroporins (*AQP3, AQP7, AQP10*) increase from duodenum to ileum, coincident with increasing lipid metabolism genes (Fig. 6D). Viewing *AQP* across lineages (Fig. 6G) shows enriched aquaglyceroporins in mature AEs, defining a likely role for AQP-mediated glycerol transfer in AE triglyceride processing. We note unappreciated specificity of *AQP1* expression in ISCs and TA cells across organs and uniquely restricted *AQP8* expression in late ACCs in the AC. Diffusion pseudotime demonstrates that *AQP8* is expressed in the most mature late ACCs, likely on the surface epithelium (Fig. 6H). Distinct expression of *AQP1* and *AQP8* at the crypt base and surface, respectively, suggests specific physiological roles that should be functionally interrogated.

### Receptors/Drugs

We finish by examining how extrinsic signals may affect the intestinal epithelium. Two approaches were designed to show how associations with receptors, drug targets, and lineage states can be revealed. First, expression plots were compiled of major receptor families across lineages, separated into high-, intermediate-, and low-expressing genes for easier visualization (File S5, Table S4). The five highest-expressing genes from each family were then grouped to visualize expression across lineages (Fig. 7A). Several patterns appear from these 60 receptors: 20 receptors are highest in tuft cells, 11 in EECs, 10 in AE/ACC, 9 in FAE; receptors bias towards villi (12) vs crypts (3); receptors bias toward SI (4) vs colon (0); and many uniquely express in certain lineages (12 in tuft, 3 in EECs, etc.), showing potential ways regions and lineages might be targeted by exogenous signals.

**Figure 7:**
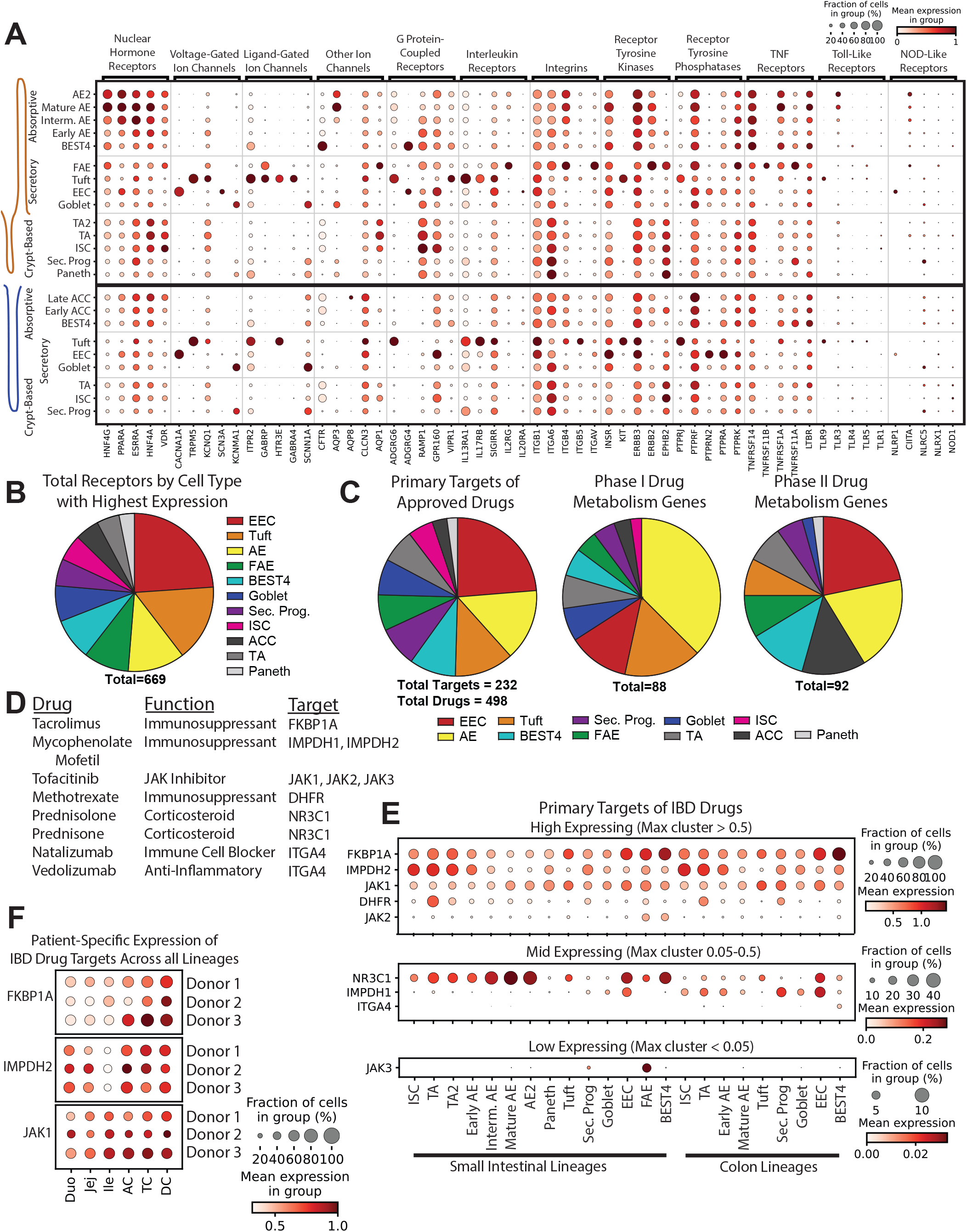
Extrinsic receptors and drug targets. **A)** Dotplot showing five highest-expressing members of major receptor families by lineage. **B)** Pie chart showing receptor genes expressed in the intestinal epithelium by lineage with highest expression. **C)** Pie chart showing primary targets of all approved drugs and Phase I and Phase II drug metabolism genes expressed in the intestinal epithelium by lineage with highest expression. **D-E)** Primary targets of drugs used to treat IBD by expression across lineages. Note scaling changes. **F)** Dotplot showing expression of three targets of IBD drugs across regions split by donor.

To test the novelty of these observations, we probed receptors with unique expression in tuft cells and searched for literature connecting these genes to tuft cells in mice or humans. We found direct connections to intestinal tuft cells for only five of the 12 (*TRPM5, ITPR2, HTR3E, IL13RA1, IL17RB*), with no connection to intestinal tuft cells found for the remaining seven (*GABRA4, ADGRG6/GPR126, SIGIRR, ITGB5, KIT, PTPRJ, TLR9*). These seven newly-defined lineage-specific receptors arose from just the five highest-expressed receptors from each family, and our full dataset includes 669 total receptors (Fig. 7B, File S5). This receptor expression atlas across lineages, organs, regions, and donors provides a powerful foundation to explore how extrinsic signals from local epithelial microenvironments, dietary and microbial influences, and pharmaceuticals affect intestinal epithelial lineages.

We explored how medicines might directly affect the intestinal epithelium. While many drugs have GI-related side effects, few are intended to target the intestinal epithelium itself^156-159^, and side effects such as diarrhea are often unexplained at the cell-lineage level^160^. We searched for primary targets of all FDA-approved drugs and found 498 approved drugs had 232 primary gene targets expressed in our gut epithelial dataset (Fig. 7C, Table S5). As most of these drugs are prescribed for non-GI diseases, primary targets within the intestinal epithelium are likely often overlooked and may contribute to unexpected GI side effects.

While many drugs are metabolized by the liver, oral drugs can be altered by metabolism within the gut epithelium^157-159^. We show expression of genes for Phase I and Phase II drug metabolism proteins by lineage with highest expression in the intestinal epithelium (Fig. 7C) and quantified gene expression by lineage and by region (Table S6). *CES2*, which metabolizes the cancer drug irinotecan into its biologically active form SN-38^161^, is found to be the highest-expressed Phase I metabolism gene in the SI, with AE enrichment. Interestingly, *UGT1A1*, the Phase II gene which inactivates SN-38^162^, has low gut epithelial expression (Table S6). This suggests irinotecan might remain active in the gut, supporting the idea that orally administered irinotecan might be effective against intestinal cancers^163-165^. Our easily-searchable dataset provides expression values for genes important for further studies probing intestinal metabolism of endobiotics, environmental toxicants, and pharmaceuticals.

As an example of a disease-focused approach, we searched for primary targets of drugs prescribed for inflammatory bowel disease (IBD). Most are anti-inflammatory or immunomodulatory, so primary targets are often not expressed in the intestinal epithelium. Our database shows nine primary gene targets of eight IBD drugs with epithelial expression (Fig. 7D). As many of these drugs are intended to affect immune cells, we mapped epithelial expression of their primary target genes to see which lineages might be inadvertently affected (Fig. 7E). We find high *FKBP1A*, a tacrolimus (Prograf^®^) target, in the little-understood BEST4^+^ cells, making this important to follow as their functions become better understood. Mycophenolate mofetil (CellCept^®^) targets *IMPDH2* and *IMPDH1* expressed in proliferative crypt populations and EECs, respectively. The methotrexate target *DHFR* is highest in TA and progenitor cells, while the tofacitinib (Embrel^®^) target *JAK1* has broader expression. These drugs can be orally administered and have side effects including diarrhea, nausea, vomiting, or appetite loss^166-168^. Interrogating this small subset of drugs in our database highlights a spectrum of potential unintended epithelial effects on renewal by ISC or TA cells, EEC hormonal influences on appetite and gut motility, and unknown effects from other lineages.

Personalized precision medicine is an emerging field motivating new technologies^169^. We used our drug-target atlas to evaluate regional variability of tacrolimus, mycophenolate mofetil, and tofacitinib target genes, *FKBP1A, IMPDH2*, and *JAK1*, across individual donors as an approach to inform potential patient-dependent effects (Fig. 7F). We find higher colonic expression of all three targets, suggesting that patients may experience effects of drugs targeting these genes in their lower GI tract. Comparing multiple donors may hint at susceptibility to drug side-effects, with Donors 2 and 3 generally expressing target genes higher than Donor 1.

While our donor number is insufficient for statistically significant conclusions, it provides a framework to generate observations to inform larger studies. We hope our lineage-, regional-, and donor-specific data on primary drug targets will be used across gastroenterology and pharmacology to better understand how drugs may affect the intestinal epithelium.

## Discussion

We use transplant organ tissue to characterize intestinal epithelial cells from duodenum through DC for three adult human donors, allowing us to define comprehensive lineage markers and map regional functionality. Our experimental design has many strengths not found in other studies. DNA-oligo hashtag antibodies allowed for all six regions from each donor to be sequenced together to save cost and avoid intra-donor batch effects, then separated computationally to analyze regional differences. This allowed for comparing gene expression between multiple regions within SI or colon. The regional differences found highlight the importance of regional selection when studying the gut, yet many colonic scRNAseq studies do not specify sample region or mention if pooled samples are from consistent regions. We analyzed cells from donors with no known intestinal diseases or cancer, avoiding effects on non-inflamed adjacent tissues in patients with inflammatory disease^9^.

Our experimental design allowed us to study many specific phenomena. Analyzing cells across six regions allowed us to compile comprehensive transcriptional signatures of genes significantly enriched in each lineage versus other lineages in all six regions and across donors. We map mucins, hormones, transporters, and barrier function genes across all six regions. We analyze PCs, showing drastic differences in growth factor expression from mouse literature and highlighting the insufficiency of *LYZ* for marking human PCs. We also analyze rare FAE cells, showing mouse/human differences and defining DEGs. We use PAGA to infer the differentiation trajectory for each lineage and suggest organ-specific maturation for tuft and BEST4^+^ cells. We propose novel tuft cell interactions with pathogens and the immune system. Finally, we map receptor expression and primary drug targets across lineages and highlight the ease of using this database to find previously undescribed gene expression. We hope our database serves as a resource to understand how drugs affect the intestinal epithelium and as guidance for future precision medicine approaches.

SI and colon from three male donors varying in age, race, and BMI were used in our study. Optimally, future studies will add regional data from diverse donors to build upon this foundation. While multiplexing six regions in one library prep allowed us to make a wide-ranging atlas, this approach results in limited numbers of each lineage analyzed per region. Our trajectory analysis suggests which lineages mature through secretory progenitors, yet future studies including more progenitors may provide more sensitive and high-resolution analyses.

In this study, we provide a comprehensive cell-level transcriptomic view of the SI and colon epithelium with regional resolution across multiple humans. Our analyses independently confirm and advance prior studies, define important differences between mouse and human lineages, and highlight how lineages change along the proximal-distal axis. We include easy-to-search tables for DEGs, receptors, and drug targets that can be interrogated by most investigators and trainees. Overall, our database provides a foundation for understanding individual contributions of diverse epithelial cells across the length of the human intestine and colon to maintaining physiologic function.

## Supporting information

All Supplemental Figures

## Abbreviations Used

SI: Small intestine
scRNAseq: single-cell RNA sequencing
AC: Ascending Colon
TC: Transverse Colon
DC: Descending Colon
PC: Paneth Cell
FAE: Follicle Associated Epithelium
EEC: Enteroendocrine Cell
GC: Goblet Cell
DEG: Differentially Expressed Genes
ISC: Intestinal Stem Cell
TA: Transit Amplifying
ACC: Absorptive Colonocyte
AE: Absorptive Enterocyte
PAGA: Partition-based Graph Abstraction
M-cell: Microfold Cell
GI: Gastrointestinal
crGC: Crypt-Resident Goblet Cell
icGC: Inter-Crypt Goblet Cell

## Supplemental Figures and Files

**Figure S1: Patient characteristics and cell counts. A)** Donor information. **B)** Cells collected per donor region. **C)** Small intestinal lineages collected per donor. **D)** Colonic lineages collected per donor. **E)** Small intestinal lineages per donor region. **F)** Colonic lineages per donor region.

**Figure S2: Tissue histology**. Hemotoxylin and eosin stained tissues from each region for all three donors. All scale bars = 200 µm

**Figure S3: FACS strategy**. FACS strategy for gating out cell fragments, likely doublets, and dead cells. ‘APC-A’ channel is detecting AnnexinV-APC.

**Figure S4: Hashtag deconvolution. A)** Per donor hashtag noise distributions. Blue dotted lines indicate 99^th^ percentile values for noise. Values above this line were called positive for a specific hashtag. **B)** (left) K-medoid clustering for each donor based only on hashtag reads. Cells positive (p<0.01) for mulitple hashtags are removed as likely multiplets. Cells are called as negative if they do not surpass the noise threshold for all hashtags, (right) k-medoid clustering with final hasthag labelling for non-multiplet cells

**Figure S5: Filtering for cell quality**. Total counts, N genes, and mitochondrial gene percentages shown for each donor before and after filtering; (top) pre-filtering and (bottom) post filtering.

**Figure S6: Determining final lineage clusters. A)** Original Leiden clustering for all cells. **B)** Seperating EEC and secretory progenitors by organ. **C-E)** An *ITLN1-*high cluster, all from SI, contains cells expressing PC markers (*DEFA5, DEFA6, ITLN2, LYZ*) along with cells expressing GC marker *MUC2*. (C) cluster defined by *ITLN1* and dotplot showing expression of markes; (D) UMAP expression of PC and GC markers across all cells; (E) UMAP expression of PC and GC markers within the *ITLN1*-high cluster. **F)** Subclustering to define Paneth and goblet cells. **G)** Final lineage clusters used for the rest of the analyses in our study.

**Figure S7: DEG dotplots for each lineage**. Dotplots showing expression of top DEGs (max: 20) for each lineage, as sorted by expression fold-change above the cluster with the next highest expression. DEGs included are genes significantly enriched in both the SI and colon (if applicable).

**Figure S8: Organ-specific lineage markers**. Relating to Fig. 2E, UMAPs showing expression of DEGs from mature lineages found to be higher enriched in SI or colon.

**Figure S9: Additional Paneth cell data. A)** Dotplot showing members of major intestinal growth factor families with expression in PCs across SI lineages. **B)** Dotplot showing expression of all Frizzled family receptors across SI lineages. **C)** Heatmap of GC markers (top), mouse-defined Paneth-Like Cell markers (middle), and PC markers (bottom) plotted across colon GCs, secretroy progenitors, and tuft cells. Cells in each lineage are sorted by increasing *REG4* expression (top row of middle third) to more easily visualize potenital overlap of marker expression.

**Figure S10: Follicle Associated Epithelium. A)** Dotplot showing expression of consrved M-cell markers and other genes known to interact with the immune system across lineages. Bottom third shows genes implicated in mouse M-cells that are not specific to human FAE. **B)** Dotplot showing expression of top 20 FAE DEGs across lineages.

**Figure S11: Additional mucin and goblet cell data. A)** Dotplot showing expression of top nine expressing mucins across GCs and proliferative and absorptive lineages of the SI and colon. **B)** Dotplot showing expression of top 9 expressed mucins in all AE and ACC lineages by intestinal region. **C)** Dotplot showing expression of mucins in colonic intercrypt goblet cells (icGC), crypt-resident goblet cells (crGCs), and early goblet cells. **D)** (Left) Leiden subclustering of SI goblet cells, with subclusters named according to genes with high expression. (Middle) UMAP of SI goblet cells marked by diffusion pseudotime. (Right) UMAP of SI goblet cells marked by *MUC2* expression. **E)** Dotplot showing expression of mucins in SI GC subpopulations. **F)** Dotplot showing expression of mouse-implicated markers in human SI GC subclusters.

**Figure S12: Additional EEC Data. A)** Dotplot showing expression of DEGs found for SI or colon EECs that are present in the GOCC_Presynapse gene list. **B)** Pie chart showing all genes within the GOCC_Presynapse gene list marked by lineage in which they have highest expression (SI and colon lineages are combined when applicable).

**File S1: REACTOME pathway enrichment analysis for TA vs TA2**

**File S2: REACTOME pathway enrichment analysis for TA2 vs TA**

**File S3: REACTOME pathway enrichment analysis for FAE DEGs**

**File S4: REACTOME pathway enrichment analysis for goblet cell DEGs**

**File S5: Receptor Family dotplots by lineage**

**Table S1: Differentially expressed gene signatures for lineages by organ**

**Table S2: Lineage signatures across full gut**

**Table S3: Correlations for BEST4**^**+**^ ***NPY* and AE gene expression**

**Table S4: Receptor family expression by lineage**

**Table S5: Primary targets of approved drugs expressed by lineage**

**Table S6: Phase I and Phase II drug metabolism genes expressed by lineage**

## Acknowledgements

We first thank the organ donors without whom our study would be impossible. We thank HonorBridge (formerly Carolina Donor Services) for assistance in coordinating organ donations. We thank Gabrielle Cannon at the Advanced Analytics core at UNC Chapel Hill and the Histology Research Core at UNC Chapel Hill. This work was supported by NIH R01DK115806, R01DK109559, and P30-DK034987 to STM, F32DK124929 to JRB, and F31-HL156433 and 5T32-GM067553 for JSR. APB is supported by a Career Development Award from the Crohn’s and Colitis Foundation, and the University Cancer Research Fund.

## Notes

All authors declare no conflicts of interest

### Competing Interest Statement

The authors have declared no competing interest.

