## Supplementary material for "A proximal-to-distal survey of healthy adult human small intestine and colon epithelium by single-cell transcriptomics": All Supplemental Figures

**Figure S1: Patient characteristics and cell counts.**

A

| Donor Characteristics | | | | | |
|---|---|---|---|---|---|
| **Donor** | **Sex** | **Age** | **Race** | **BMI** | **Intestinal Status** |
| Donor 1 | Male | 29 | White | 36.4 | Healthy |
| Donor 2 | Male | 45 | White | 52.2 | Healthy |
| Donor 3 | Male | 53 | African American | 25 | Healthy |

B

| Cells collected per donor region | | | | |
|---|---|---|---|---|
| | **Total Cells** | **Donor 1** | **Donor 2** | **Donor 3** |
| **Duodenum** | 2115 | 687 | 423 | 1005 |
| **Jejunum** | 2612 | 869 | 961 | 782 |
| **Ileum** | 2609 | 918 | 1097 | 594 |
| **Asc. Colon** | 1746 | 733 | 557 | 456 |
| **Trans. Colon** | 2171 | 567 | 1042 | 562 |
| **Desc. Colon** | 1337 | 556 | 385 | 396 |

C

| Small intestinal lineages collected per donor | | | | |
|---|---|---|---|---|
| | **Total Cells** | **Donor 1** | **Donor 2** | **Donor 3** |
| **Mature AE** | 1569 | 607 | 372 | 590 |
| **Intermediate AE** | 1443 | 631 | 302 | 510 |
| **EArly AE** | 1237 | 455 | 498 | 284 |
| **ISC** | 873 | 239 | 448 | 186 |
| **AE2** | 591 | 35 | 113 | 443 |
| **TA** | 397 | 111 | 259 | 27 |
| **Goblet** | 388 | 77 | 184 | 127 |
| **BEST4** | 339 | 148 | 73 | 118 |
| **Tuft** | 184 | 81 | 94 | 9 |
| **TA2** | 105 | 29 | 67 | 9 |
| **EEC** | 72 | 19 | 25 | 28 |
| **Sec. Progenitors** | 70 | 34 | 12 | 24 |
| **Paneth** | 49 | 8 | 15 | 26 |
| **FAE** | 19 | 0 | 19 | 0 |

D

| Colonic lineages collected per donor | | | | |
|---|---|---|---|---|
| | **Total Cells** | **Donor 1** | **Donor 2** | **Donor 3** |
| **Early ACC** | 1464 | 783 | 533 | 148 |
| **Goblet** | 1189 | 157 | 435 | 597 |
| **Late ACC** | 552 | 292 | 126 | 134 |
| **Tuft** | 524 | 191 | 230 | 103 |
| **TA** | 518 | 140 | 344 | 34 |
| **BEST4** | 477 | 92 | 81 | 304 |
| **ISC** | 385 | 178 | 171 | 36 |
| **EEC** | 82 | 15 | 22 | 45 |
| **Sec. Progenitors** | 63 | 8 | 42 | 13 |

E

| | | Small Intestinal lineages per donor region | | | | | | | | |
|---|---|---|---|---|---|---|---|---|---|---|
| | | **Donor 1** | | | **Donor 2** | | | **Donor 3** | | |
| **Cell Type** | **Total #** | **Duodenum** | **Jejunum** | **Ileum** | **Duodenum** | **Jejunum** | **Ileum** | **Duodenum** | **Jejunum** | **Ileum** |
| **Mature Abs. Enterocytes** | 1569 | 43 | 192 | 372 | 20 | 33 | 319 | 278 | 311 | 1 |
| **Intermediate Abs. Enterocytes** | 1443 | 262 | 164 | 205 | 15 | 42 | 245 | 368 | 138 | 4 |
| **Early Abs. Enterocytes** | 1237 | 190 | 171 | 94 | 116 | 246 | 136 | 141 | 99 | 44 |
| **Stem Cells** | 873 | 90 | 96 | 53 | 112 | 299 | 37 | 90 | 75 | 21 |
| **Abs. Enterocytes 2** | 591 | 0 | 13 | 22 | 30 | 71 | 12 | 2 | 0 | 441 |
| **Transit Amplifying Cells** | 397 | 40 | 50 | 21 | 29 | 69 | 161 | 17 | 4 | 6 |
| **Goblet Cells** | 388 | 10 | 44 | 23 | 44 | 106 | 34 | 28 | 65 | 34 |
| **BEST4 Cells** | 339 | 18 | 68 | 62 | 12 | 28 | 33 | 49 | 45 | 24 |
| **Tuft Cells** | 184 | 8 | 29 | 44 | 21 | 25 | 48 | 2 | 5 | 2 |
| **Transit Amplifying 2** | 105 | 10 | 17 | 2 | 11 | 15 | 41 | 3 | 5 | 1 |
| **Enteroendocrine Cells** | 72 | 11 | 5 | 3 | 8 | 10 | 7 | 10 | 14 | 4 |
| **Secretory Progenitors** | 70 | 4 | 14 | 16 | 0 | 7 | 5 | 10 | 9 | 5 |
| **Paneth Cells** | 49 | 1 | 6 | 1 | 1 | 8 | 6 | 7 | 12 | 7 |
| **FAE Cells** | 19 | 0 | 0 | 0 | 4 | 2 | 13 | 0 | 0 | 0 |

F

| | | Colon lineages per donor region | | | | | | | | |
|---|---|---|---|---|---|---|---|---|---|---|
| | | **Donor 1** | | | **Donor 2** | | | **Donor 3** | | |
| **Cell Type** | **Total #** | **Asc. Colon** | **Trans. Colon** | **Desc. Colon** | **Asc. Colon** | **Trans. Colon** | **Desc. Colon** | **Asc. Colon** | **Trans. Colon** | **Desc. Colon** |
| **Late Abs. Colonocytes** | 552 | 93 | 99 | 100 | 22 | 82 | 22 | 29 | 76 | 29 |
| **Early Abs. Colonocytes** | 1464 | 314 | 254 | 215 | 128 | 283 | 122 | 33 | 64 | 51 |
| **Stem Cells** | 385 | 68 | 72 | 38 | 63 | 79 | 29 | 10 | 12 | 14 |
| **Transit Amplifying Cells** | 518 | 76 | 29 | 35 | 55 | 233 | 56 | 15 | 12 | 7 |
| **Goblet Cells** | 1189 | 65 | 27 | 65 | 182 | 207 | 46 | 215 | 224 | 158 |
| **Secretory Progenitors** | 63 | 4 | 1 | 3 | 15 | 23 | 4 | 6 | 6 | 1 |
| **BEST4 Cells** | 477 | 26 | 28 | 38 | 21 | 38 | 22 | 100 | 111 | 93 |
| **Tuft Cells** | 524 | 76 | 55 | 60 | 65 | 90 | 75 | 31 | 39 | 33 |
| **Enteroendocrine Cells** | 82 | 11 | 2 | 2 | 6 | 7 | 9 | 17 | 18 | 10 |

**Figure S2: Tissue histology**

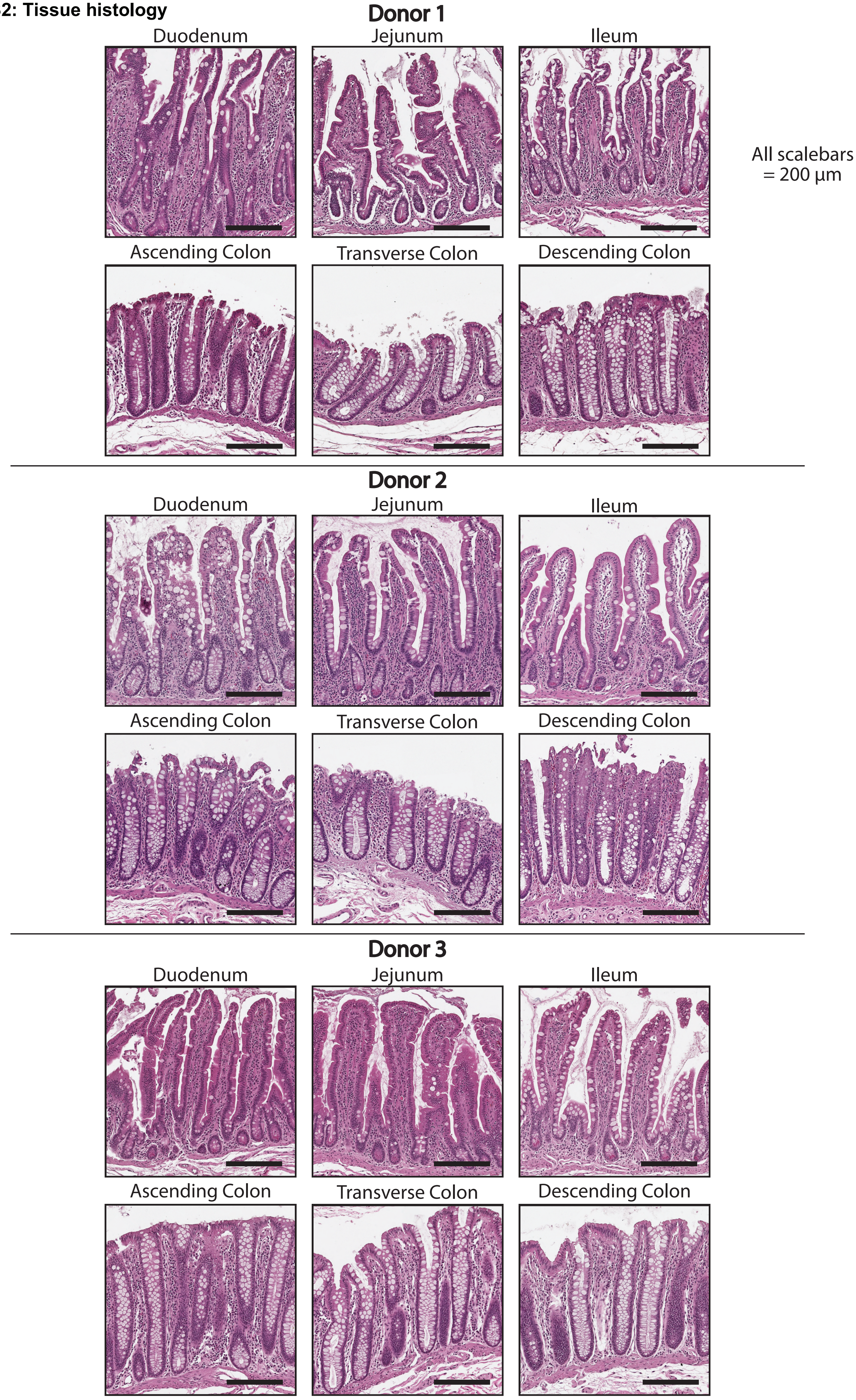

**Figure S3: FACS strategy**

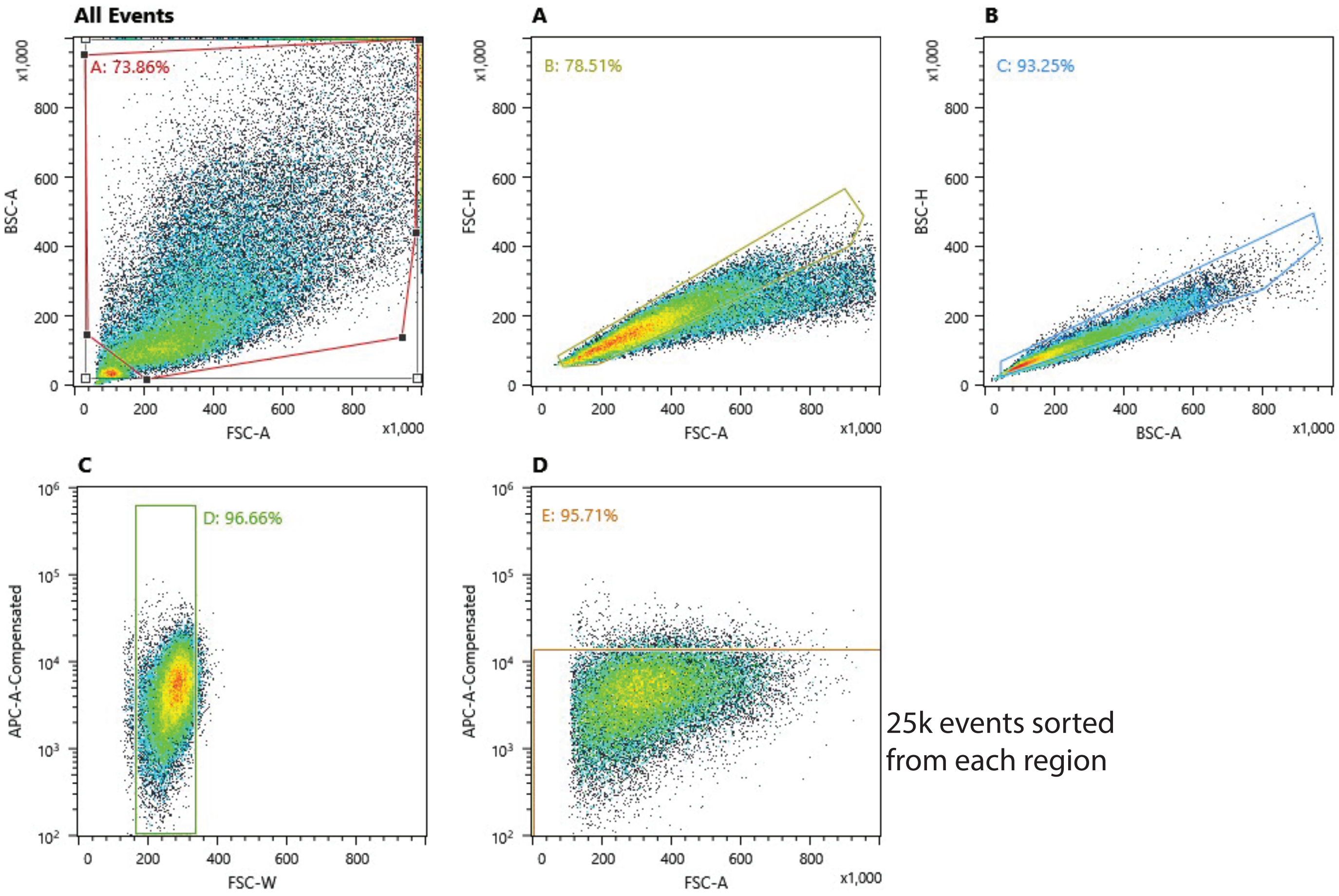

**Figure S4: Hashtag deconvolution**

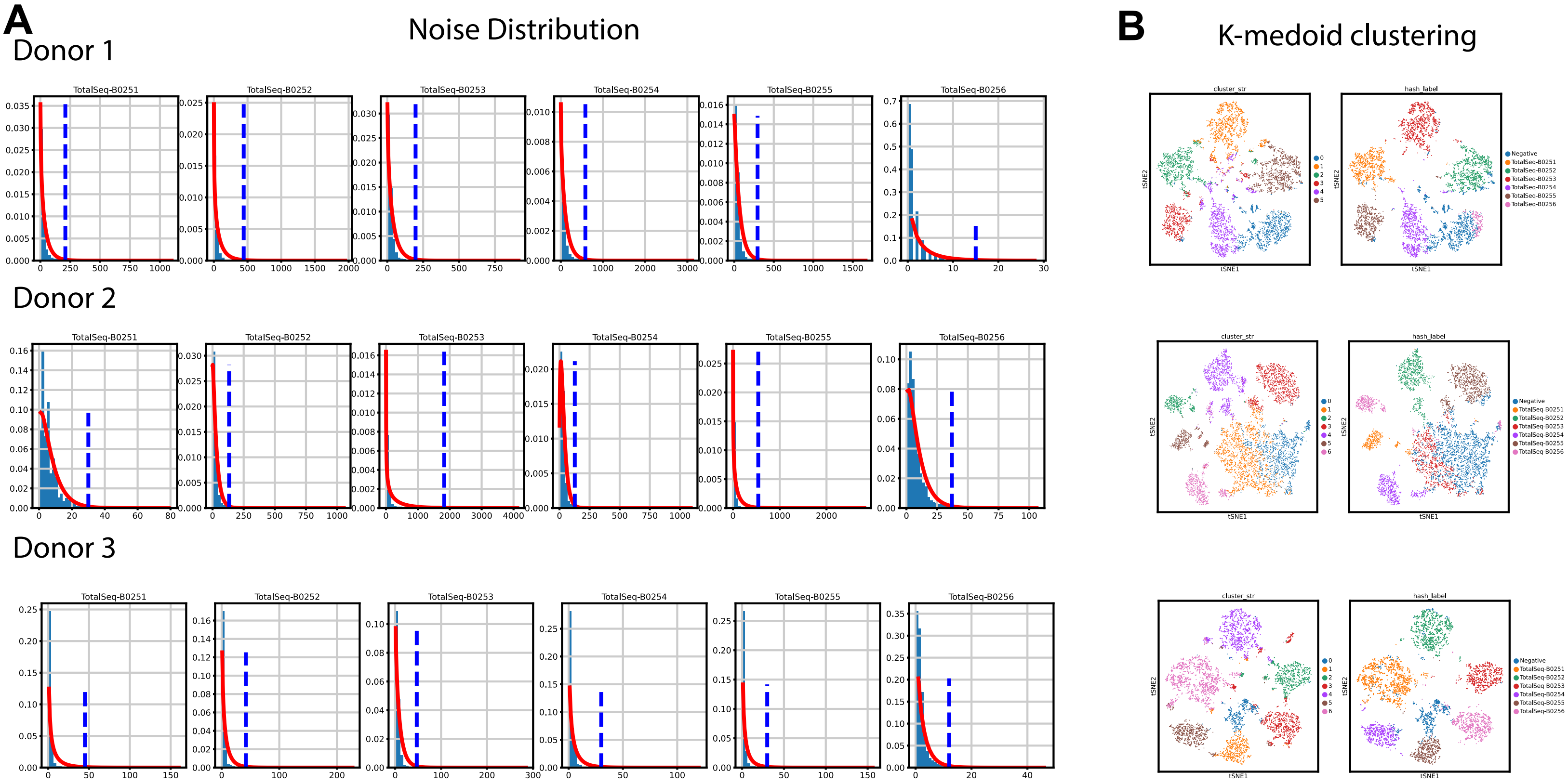

**Figure S5: Filtering for cell quality**

#### Pre-filtering distribution of quality control parameters

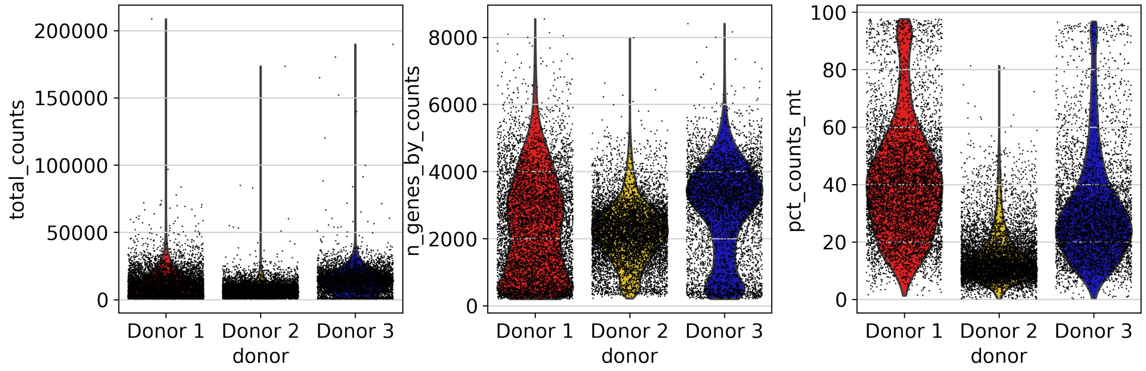


#### Post-filtering distribution of quality control parameters

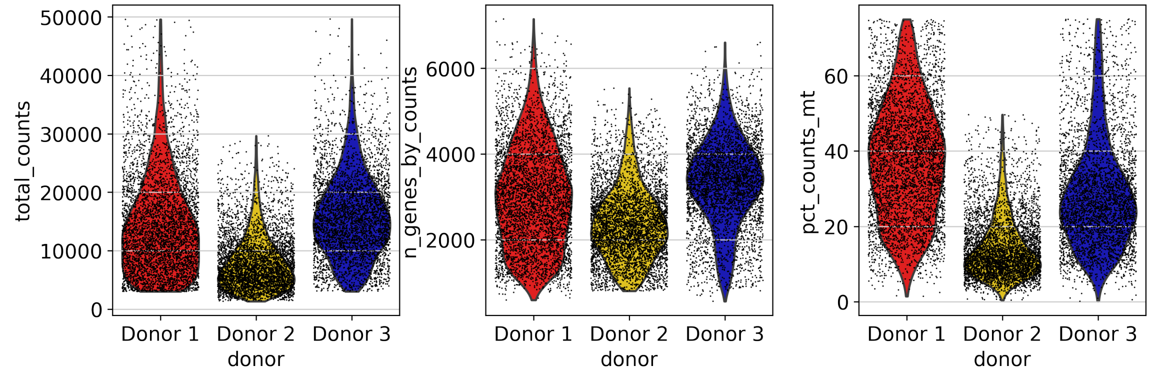

### Figure S6: Determining final lineage clusters.

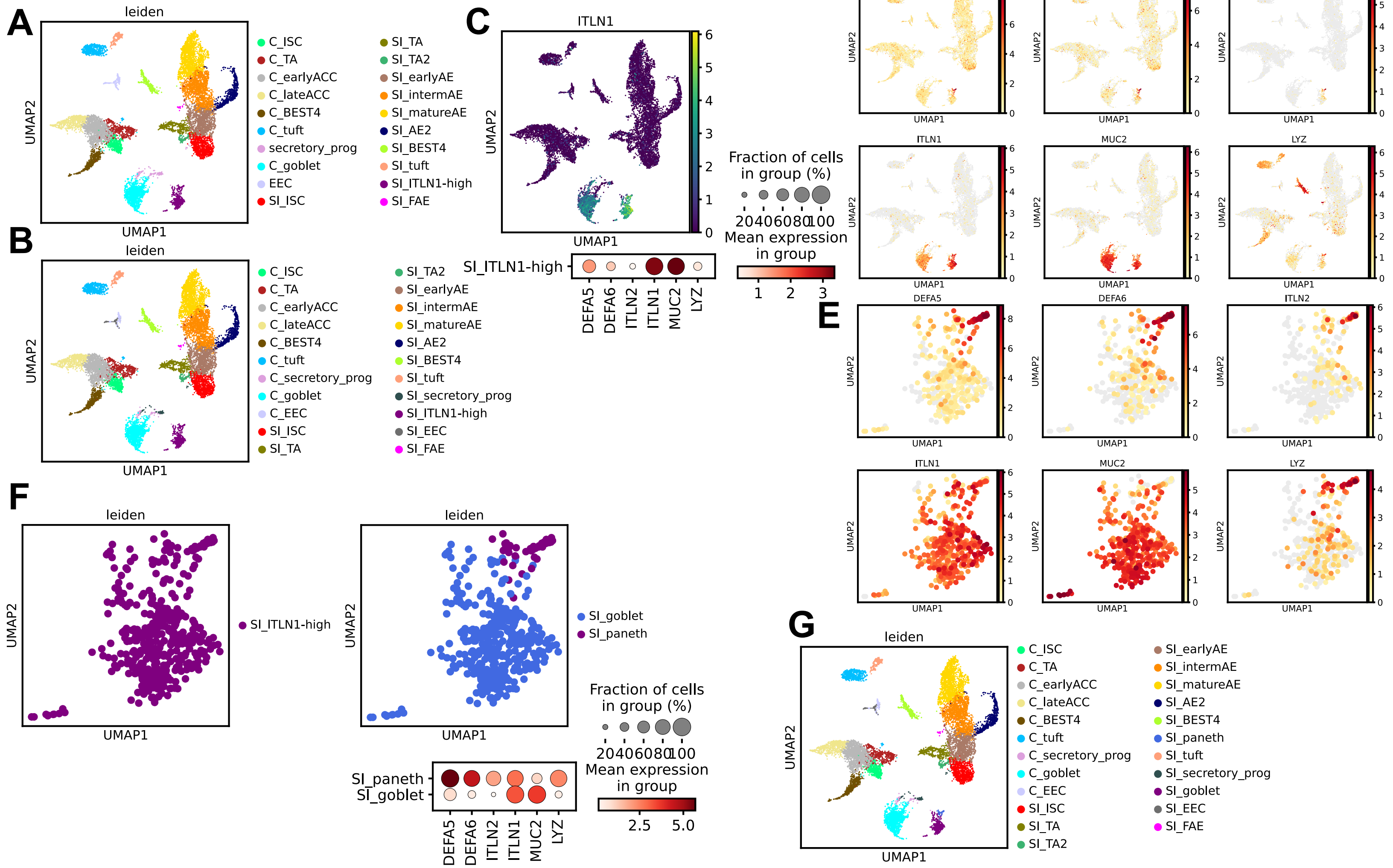

**Figure S7: DEG dotplots for each lineage**

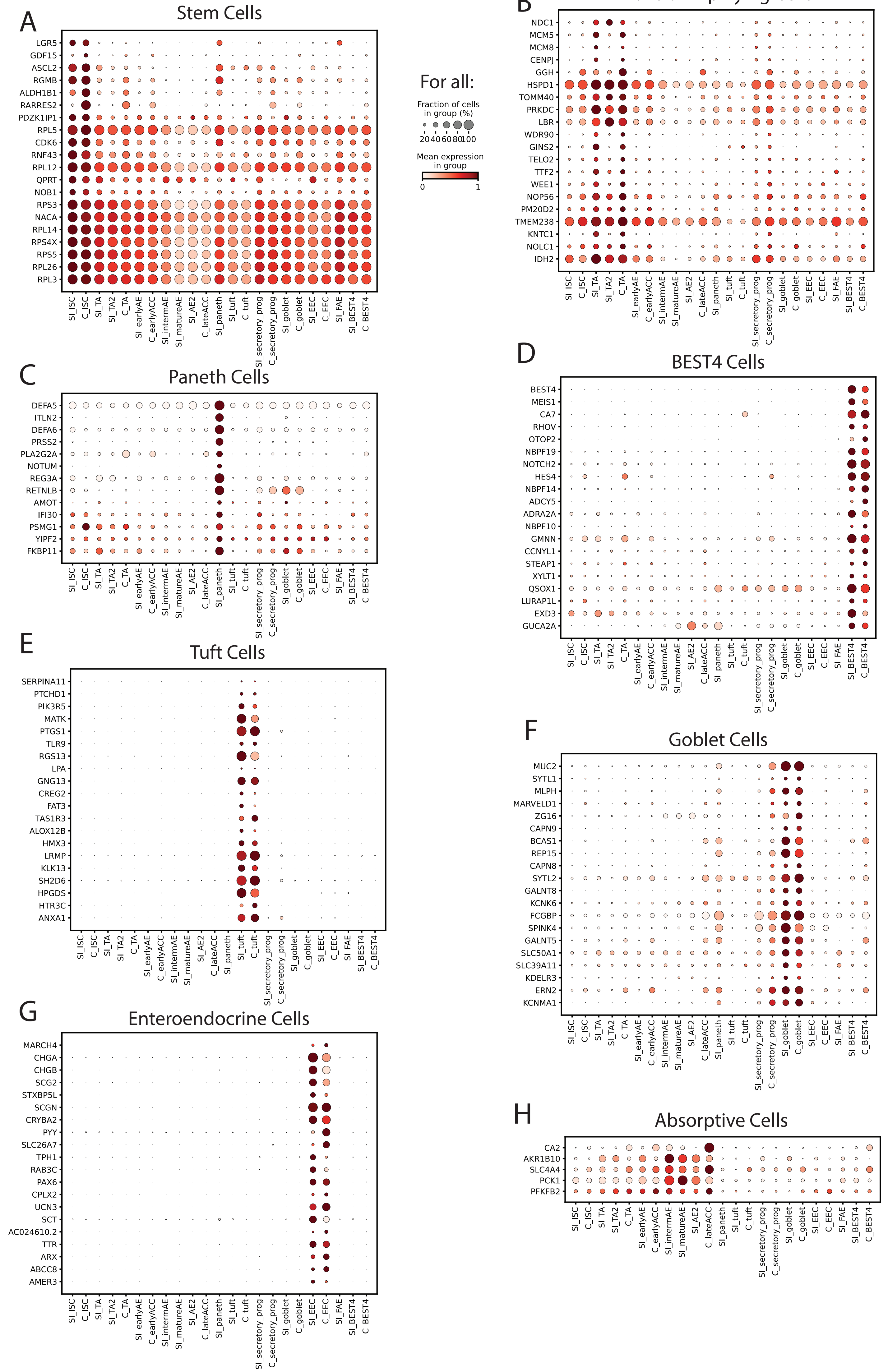

### Figure S8: Organ-specific lineage markers

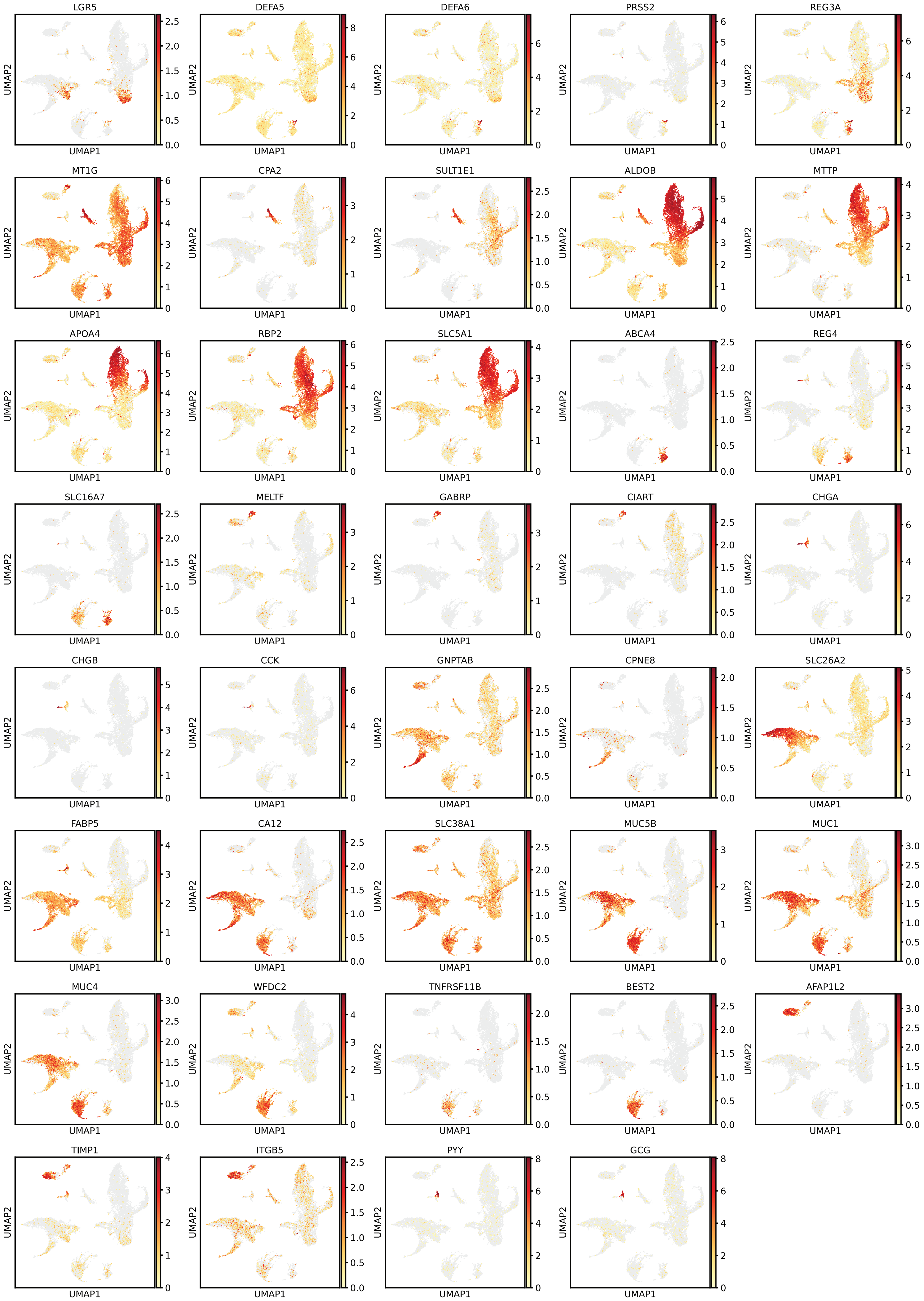

**Figure S9: Additional Paneth cell data**

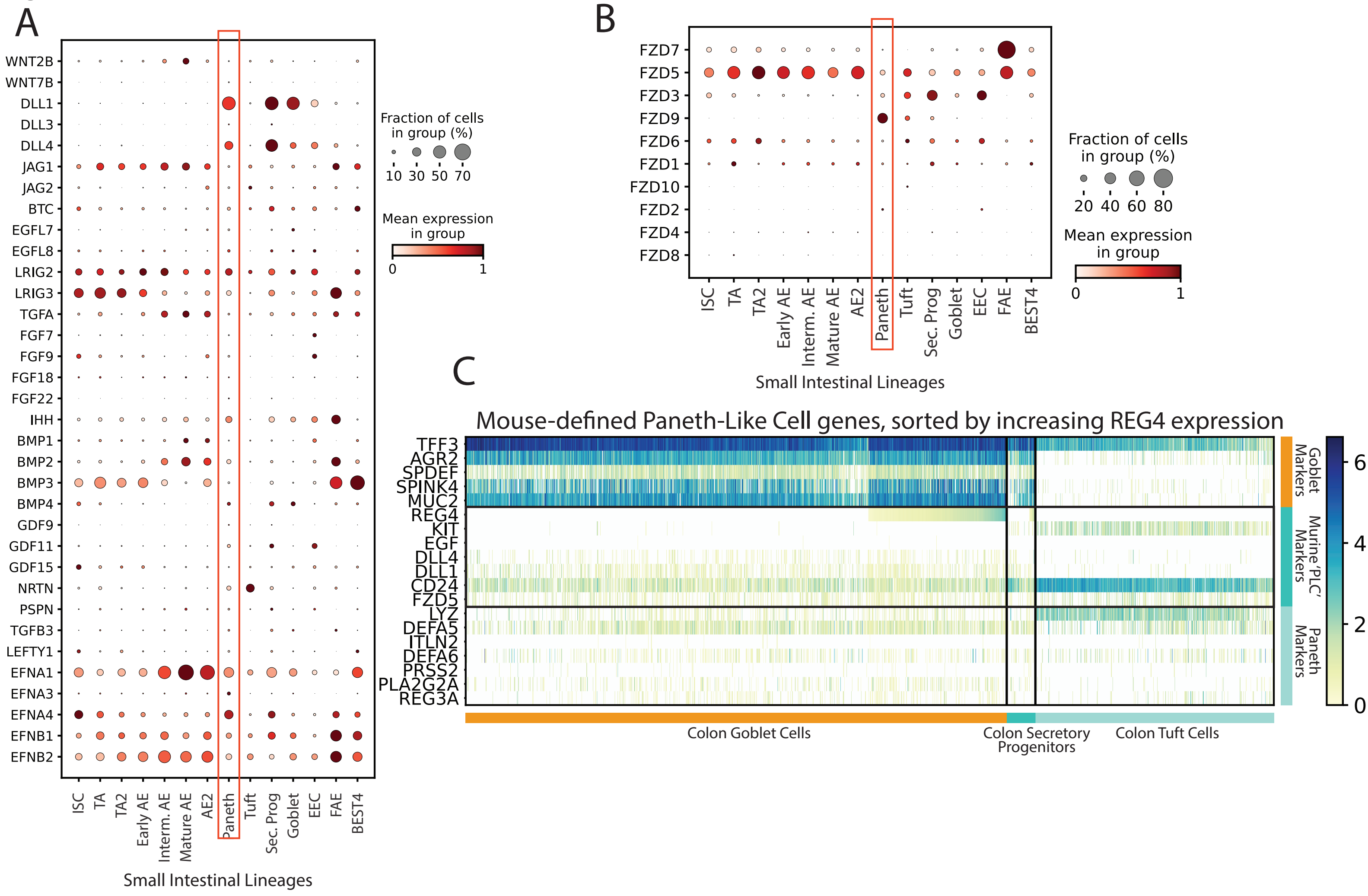

### Figure S10: Follicle Associated Epithelium

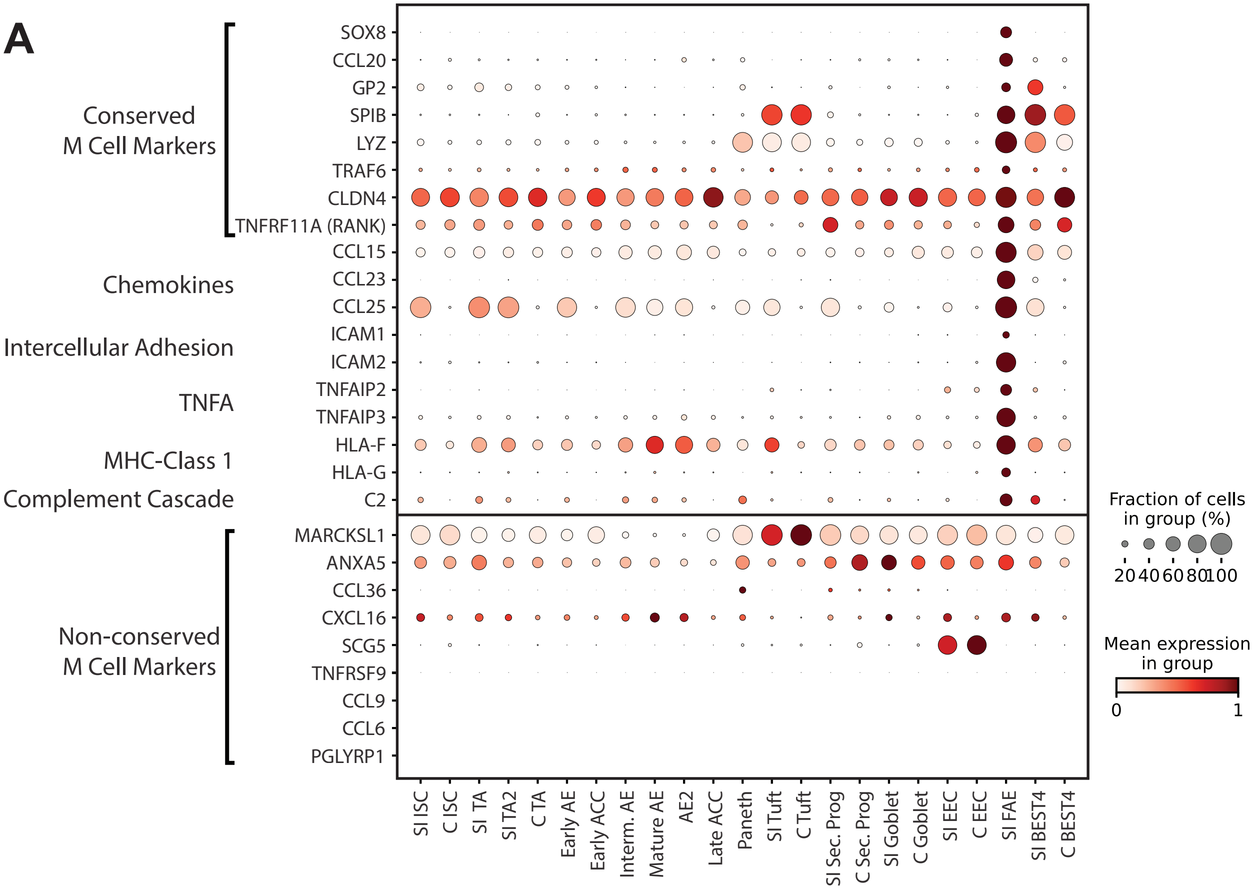


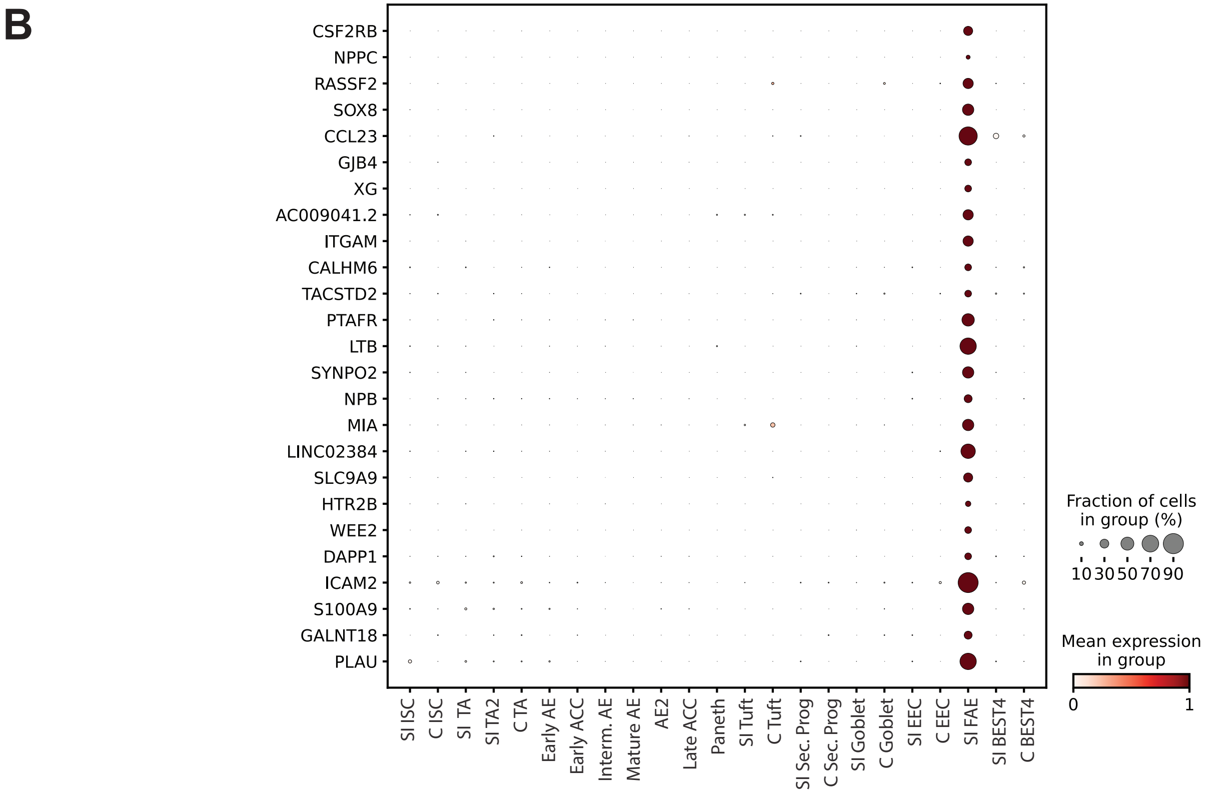

### Figure S11: Additional mucin and goblet cell data

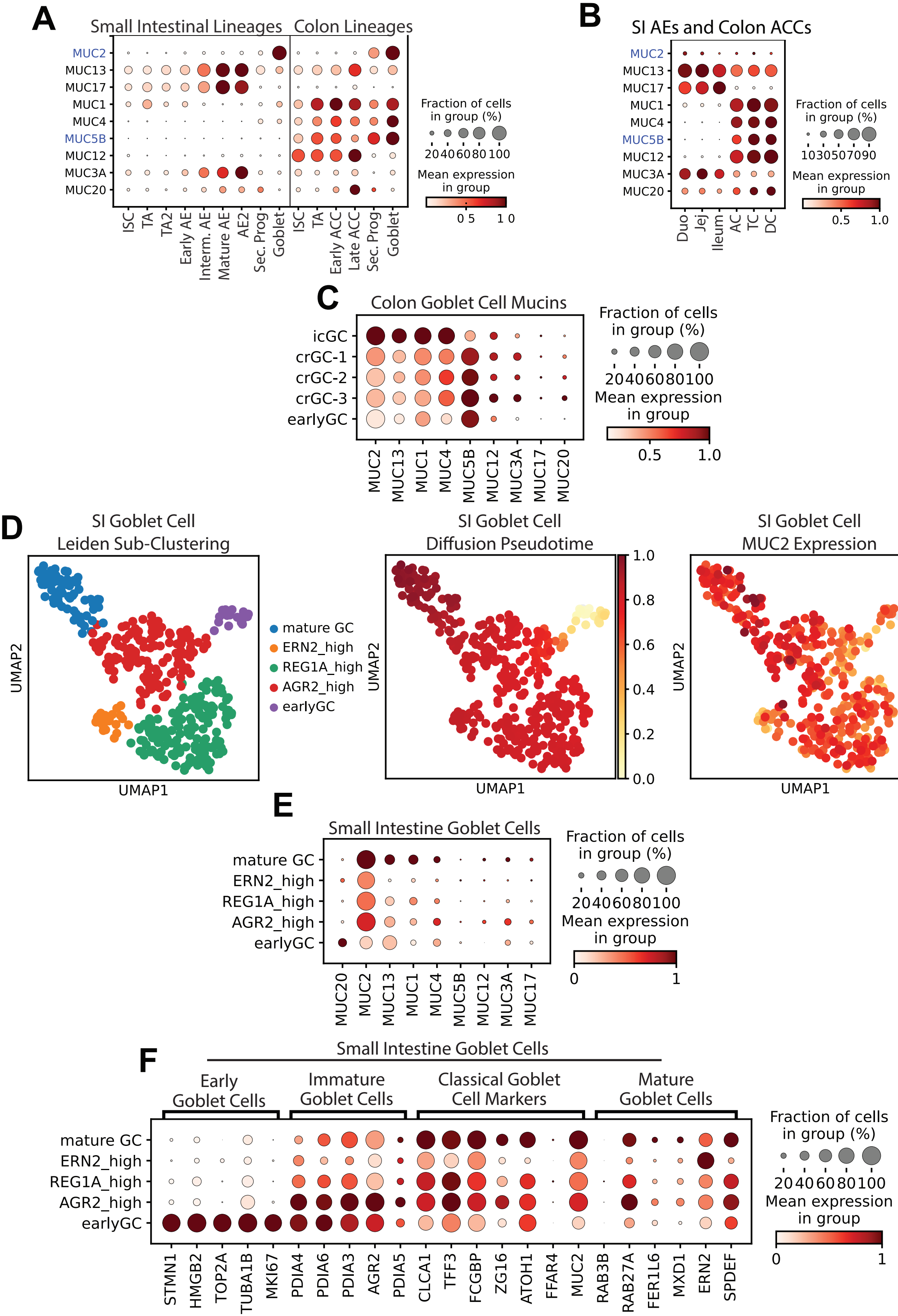

**Figure S12: Additional EEC Data.**

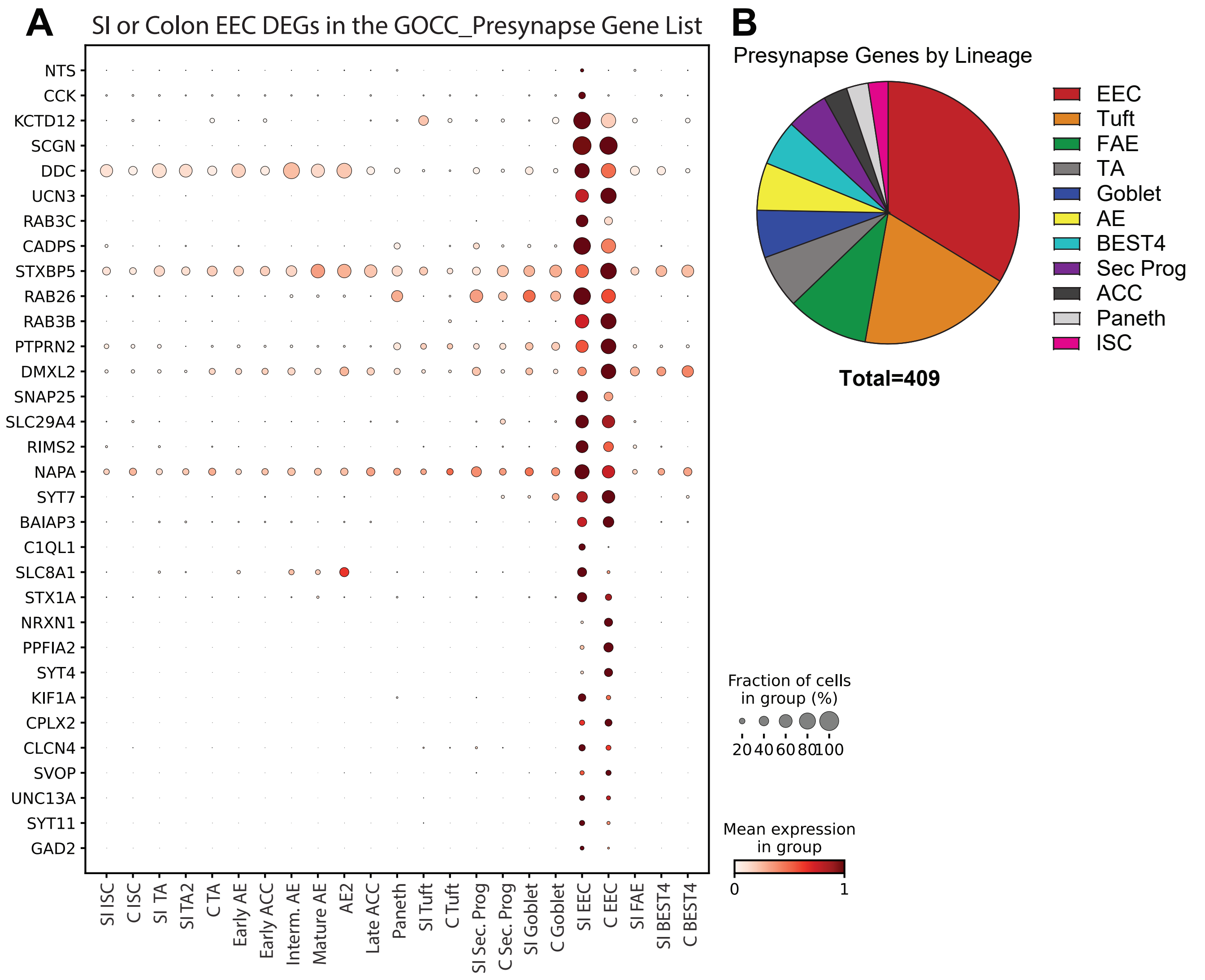
