## Supplemental Methods for "A proximal-to-distal survey of healthy adult human small intestine and colon epithelium by single-cell transcriptomics"

**Reagents used**

| **Reagent** | **Company** | **Catalog Number** |
| --- | --- | --- |
| N-Acetylcholine (NAC) | Sigma-Aldrich | A9165 |
| Dulbecco′s Phosphate Buffered Saline (dPBS) | Gibco | 14190-144 |
| Na2HPO4 | Sigma | S7907 |
| KH2PO4 | Sigma | P5655 |
| NaCl | Sigma | S5886 |
| KCl | Sigma | P5405 |
| Sucrose | Fisher BP | 220-1 |
| d-sorbitol | Fisher BP | 439-500 |
| Y27632 | Selleck Chemical | S6390 |
| Ethylenediaminetetraacetic acid (EDTA) | Corning | 46-034-Cl |
| Dithiothreitol (DTT) | Fisher Scientific BP | 172-5 |
| Protease VIII | Sigma | P5380 |
| Advanced DMEM/F12 | Gibco | 12634-010 |
| Bovine Serum Albumin | Fisher Scientific | BP1600-1 |
| AnnexinV-APC | BioLegend | 640920 |
| TotalSeq Anti-Human Hashtag Antibodies | BioLegend | B0251-B0256 |

**Data processing, filtering, doublet removal, feature selection**

After sequencing, single-cell fastq files were aligned to reference transcriptome GRCh38 with the 10X Cell Ranger pipeline (V4.0.0), and downstream analysis was performed with scanpy (v1.7.2). Annotations for cell cycle phase predictions were added following previously published methods^1^. Quality filtering thresholds for each donor were:

|  | Donor 1 | Donor 2 | Donor 3 |
| --- | --- | --- | --- |
| Minimum genes | >500 | >800 | >500 |
| Percent mitochondrial reads | <75% | <50% | <75% |
| Minimum counts | >3,000 | >1,000 | >3,000 |
| Maximum counts | <50,000 | <30,000 | <50,000 |

Following filtering, read counts were log-transformed and normalized to the median read depth of Donor 2, which had the fewest read counts. Variability due to gene expression count, mitochondrial percentage, and cell-cycle gene expression were regressed out by simple linear regression. Highly variable genes were identified with the Seurat method^2^ (min_dispersion = 0.2, min_mean = 0.0125, max_mean = 6), identifying 2777 genes that were subsequently used for principal component analysis. Genes were scaled to have a mean of zero and unit variance.

**Identifying transcriptionally distinct sub-clusters**

Subclustering was performed to isolate Paneth cells from SI goblet cells, which cluster together in the overall dataset. For Paneth cells, the ‘SI_*ITLN1*-high’ cluster was subset from the main dataset and 40 principal components were recalculated and re-harmonized. Leiden clusters were recalculated on the new principal components based on the same 2777 highly variable genes as the initial dataset with Leiden resolution = 0.15 and k = 15 neighbors.

Further subclustering on SI and colon goblet cells was performed to show goblet cell heterogeneity in the SI and colon separately. For colon goblet cells, leiden clustering settings were as follows: k = 5 nearest neighbors, leiden resolution = 0.3, and 4000 highly variable genes were calculated based on 22 recomputed principal components from the colon goblet cells subset. For SI goblet cells, leiden clustering settings were as follows: k = 10 nearest neighbors, leiden resolution = 0.4, and 2000 highly variable genes were calculated based on 19 recomputed principal components on the SI goblet cell subset.

**Online Databases**

Human homologs for mouse genes were defined using Ensembl version release 104^3^. Pathway enrichment analysis was performed using Reactome^4^, with focus given to pathways with false discovery rate (FDR) <0.05, as calculated by over-representation analysis, and full reports are included as supplemental files. Common drugs prescribed for ulcerative colitis and Crohn’s disease were curated using online literature. Comprehensive receptor family lists and primary target genes for all approved drugs were downloaded from Guide to Pharmacology^5^. Phase I and Phase II Drug metabolism genes were defined via Reactome^4^.

**Trajectory Analysis**

To infer differentiation trajectories, subcustered cell populations were separated into SI and colon as previously described. For each dataset, PAGA (v1.2) was then performed on a k-nearest neighbor graph of 20 neighbors constructed from 40 principle components. The resultant transition connectivity matrix was filtered to remove spurious connections (SI > 0.08, colon > 0.09).

**Differential expression analysis**

To determine genes that consistently mark a lineage, as determined by previously described leiden clustering, in all three donors, the dataset was first separated into small intestine- and colon-specific data. For each organ, depth-normalized expression of each gene was used fit to a negative binomial general linear model with the diffxpy package (v0.7.4). A Wald test was used to iterate through all cell lineages, testing a null model in which only donor-specific batch effects were included,

$$x_{i}= \beta_{0}+\beta_{1}Donor$$

against an alternative model where a cell’s inclusion in the current test lineage was included as a binary independent variable, correcting for multiple testing using the Benjamini-Hochberg procedure.

$$x_{i}= \beta_{0}+\beta_{1}Donor+\beta_{2}Lineage$$

Each test was then repeated independently on each donor including at least 10 cells of the lineage, excluding donor as a covariate. A gene was determined to be a marker gene for a particular lineage if it met these thresholds:

1. Maximum expression in the lineage of interest
2. q value < 0.05 in the combined dataset and all three donors individually
3. minimum log2-fold-change (compared to the next highest expressing lineage) > 0.25
4. Mean in-lineage normalized expression > 0.2
