## Supplementary material for "A proximal-to-distal survey of healthy adult human small intestine and colon epithelium by single-cell transcriptomics": File S1

### Pathway Analysis Report

This report contains the pathway analysis results for the submitted sample ''. Analysis was performed against Reactome version 77 on 25/08/2021. The web link to these results is:

https://reactome.org/PathwayBrowser/#/ANALYSIS=MjAyMTA4MjUxMzE1MzRfMTEyOA%3D%3D

Please keep in mind that analysis results are temporarily stored on our server. The storage period depends on usage of the service but is at least 7 days. As a result, please note that this URL is only valid for a limited time period and it might have expired.

### Table of Contents


1. Introduction
2. Properties
3. Genome-wide overview
4. Most significant pathways
5. Pathways details
6. Identifiers found
7. Identifiers not found

### 1. Introduction

Reactome is a curated database of pathways and reactions in human biology. Reactions can be considered as pathway 'steps'. Reactome defines a 'reaction' as any event in biology that changes the state of a biological molecule. Binding, activation, translocation, degradation and classical biochemical events involving a catalyst are all reactions. Information in the database is authored by expert biologists, entered and maintained by Reactome's team of curators and editorial staff. Reactome content frequently cross-references other resources e.g. NCBI, Ensembl, UniProt, KEGG (Gene and Compound), ChEBI, PubMed and GO. Orthologous reactions inferred from annotation for Homo sapiens are available for 17 non-human species including mouse, rat, chicken, puffer fish, worm, fly, yeast, rice, and Arabidopsis. Pathways are represented by simple diagrams following an SBGN-like format.

Reactome's annotated data describe reactions possible if all annotated proteins and small molecules were present and active simultaneously in a cell. By overlaying an experimental dataset on these annotations, a user can perform a pathway over-representation analysis. By overlaying quantitative expression data or time series, a user can visualize the extent of change in affected pathways and its progression. A binomial test is used to calculate the probability shown for each result, and the p-values are corrected for the multiple testing (Benjamini–Hochberg procedure) that arises from evaluating the submitted list of identifiers against every pathway.

To learn more about our Pathway Analysis, please have a look at our relevant publications:

Fabregat A, Sidiropoulos K, Garapati P, Gillespie M, Hausmann K, Haw R, … D'Eustachio P (2016). The reactome pathway knowledgebase. Nucleic Acids Research, 44(D1), D481–D487. https://doi.org/10.1093/nar/gkv1351.

Fabregat A, Sidiropoulos K, Viteri G, Forner O, Marin-Garcia P, Arnau V, … Hermjakob H (2017). Reactome pathway analysis: a high-performance in-memory approach. BMC Bioinformatics, 18.

### 2. Properties

- This is an **overrepresentation** analysis: A statistical (hypergeometric distribution) test that determines whether certain Reactome pathways are over-represented (enriched) in the submitted data. It answers the question 'Does my list contain more proteins for pathway X than would be expected by chance?' This test produces a probability score, which is corrected for false discovery rate using the Benjamani-Hochberg method.
- 86 out of 122 identifiers in the sample were found in Reactome, where 727 pathways were hit by at least one of them.
- All non-human identifiers have been converted to their human equivalent.
- This report is filtered to show only results for species 'Homo sapiens' and resource 'all resources'.
- The unique ID for this analysis (token) is MjAyMTA4MjUxMzE1MzRfMTEyOA%3D%3D. This ID is valid for at least 7 days in Reactome's server. Use it to access Reactome services with your data.

#### 3. Genome-wide overview

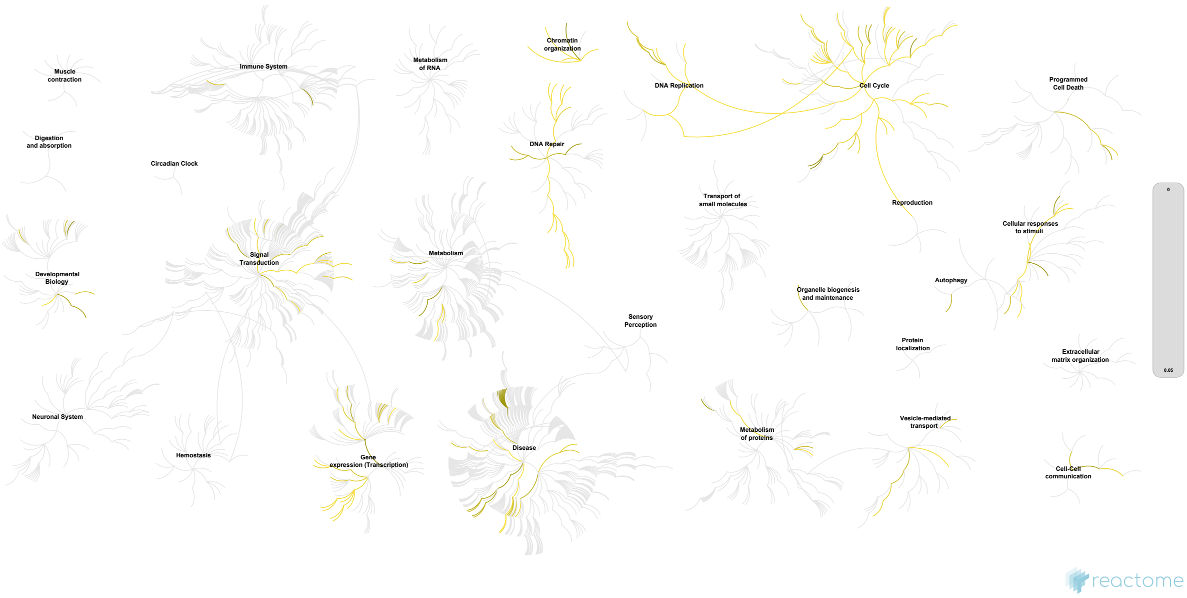


This figure shows a genome-wide overview of the results of your pathway analysis. Reactome pathways are arranged in a hierarchy. The center of each of the circular "bursts" is the root of one top-level pathway, for example "DNA Repair". Each step away from the center represents the next level lower in the pathway hierarchy. The color code denotes over-representation of that pathway in your input dataset. Light grey signifies pathways which are not significantly over-represented.

### 4. Most significant pathways

The following table shows the 25 most relevant pathways sorted by p-value.

| Pathway name | Entities | | | | Reactions | |
|---|---|---|---|---|---|---|
| | found | ratio | p-value | FDR* | found | ratio |
| Cell Cycle, Mitotic | 44 / 596 | 0.041 | 1.11e-16 | 2.09e-14 | 181 / 350 | 0.026 |
| Cell Cycle | 51 / 734 | 0.05 | 1.11e-16 | 2.09e-14 | 228 / 449 | 0.033 |
| G1/S Transition | 21 / 150 | 0.01 | 1.11e-16 | 2.09e-14 | 30 / 61 | 0.005 |
| Mitotic G1 phase and G1/S transition | 22 / 173 | 0.012 | 1.11e-16 | 2.09e-14 | 42 / 99 | 0.007 |
| G1/S-Specific Transcription | 12 / 43 | 0.003 | 8.47e-14 | 1.27e-11 | 11 / 28 | 0.002 |
| G2/M Checkpoints | 15 / 154 | 0.011 | 1.75e-10 | 2.19e-08 | 8 / 24 | 0.002 |
| Unwinding of DNA | 6 / 12 | 8.25e-04 | 5.16e-09 | 4.85e-07 | 3 / 4 | 2.96e-04 |
| DNA strand elongation | 8 / 38 | 0.003 | 1.12e-08 | 8.05e-07 | 13 / 15 | 0.001 |
| Cell Cycle Checkpoints | 17 / 280 | 0.019 | 1.18e-08 | 8.05e-07 | 12 / 56 | 0.004 |
| Meiosis | 10 / 92 | 0.006 | 7.78e-08 | 4.51e-06 | 11 / 15 | 0.001 |
| Activation of the pre-replicative complex | 7 / 36 | 0.002 | 1.61e-07 | 8.03e-06 | 8 / 9 | 6.66e-04 |
| Meiotic recombination | 8 / 58 | 0.004 | 2.75e-07 | 1.21e-05 | 5 / 9 | 6.66e-04 |
| DNA Replication | 11 / 142 | 0.01 | 4.89e-07 | 2.01e-05 | 31 / 47 | 0.003 |
| M Phase | 18 / 416 | 0.029 | 6.41e-07 | 2.50e-05 | 50 / 91 | 0.007 |
| S Phase | 12 / 180 | 0.012 | 7.01e-07 | 2.59e-05 | 32 / 54 | 0.004 |
| Reproduction | 10 / 123 | 0.008 | 1.07e-06 | 3.74e-05 | 11 / 24 | 0.002 |
| RMTs methylate histone arginines | 7 / 53 | 0.004 | 2.07e-06 | 6.62e-05 | 18 / 22 | 0.002 |
| Synthesis of DNA | 10 / 133 | 0.009 | 2.13e-06 | 6.62e-05 | 18 / 26 | 0.002 |
| Diseases of DNA repair | 6 / 37 | 0.003 | 3.56e-06 | 1.07e-04 | 5 / 29 | 0.002 |
| Activation of ATR in response to replication stress | 6 / 39 | 0.003 | 4.79e-06 | 1.39e-04 | 2 / 9 | 6.66e-04 |
| Polo-like kinase mediated events | 5 / 23 | 0.002 | 5.86e-06 | 1.58e-04 | 5 / 15 | 0.001 |
| Diseases of DNA Double-Strand Break Repair | 5 / 24 | 0.002 | 7.18e-06 | 1.72e-04 | 4 / 4 | 2.96e-04 |
| Defective HDR through Homologous Recombination (HRR) due to PALB2 loss of function | 5 / 24 | 0.002 | 7.18e-06 | 1.72e-04 | 2 / 2 | 1.48e-04 |
| Defective HDR through Homologous Recombination Repair (HRR) due to PALB2 loss of BRCA2/RAD51/RAD51C binding function | 5 / 24 | 0.002 | 7.18e-06 | 1.72e-04 | 1 / 1 | 7.40e-05 |
| Defective HDR through Homologous Recombination Repair (HRR) due to PALB2 loss of BRCA1 binding function | 5 / 24 | 0.002 | 7.18e-06 | 1.72e-04 | 1 / 1 | 7.40e-05 |

* False Discovery Rate

### 5. Pathways details

For every pathway of the most significant pathways, we present its diagram, as well as a short summary, its bibliography and the list of inputs found in it.

#### 1. Cell Cycle, Mitotic (R-HSA-69278)

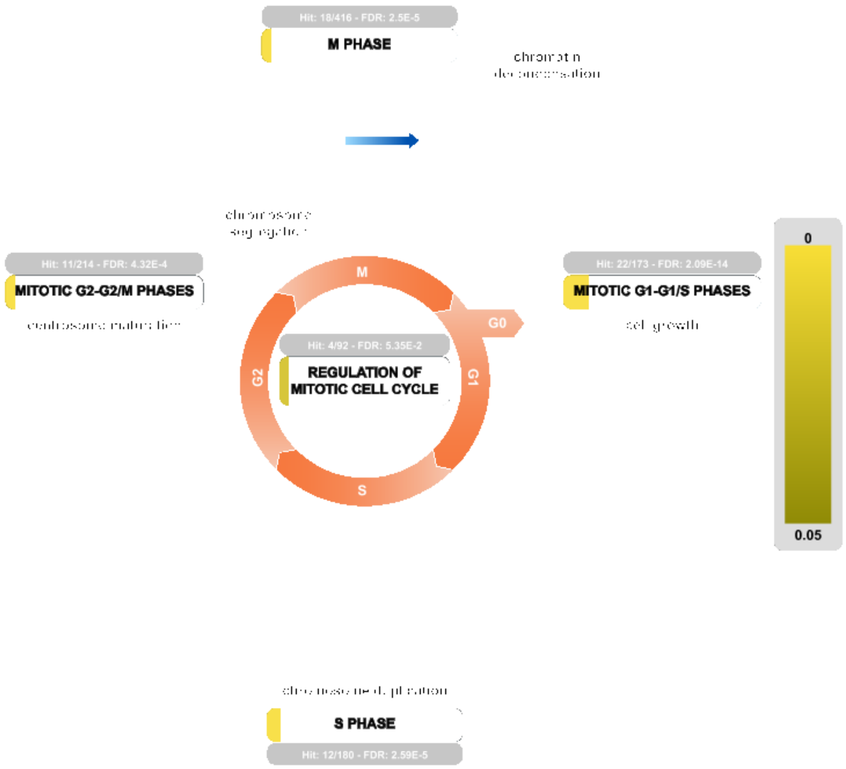


The events of replication of the genome and the subsequent segregation of chromosomes into daughter cells make up the cell cycle. DNA replication is carried out during a discrete temporal period known as the S (synthesis)-phase, and chromosome segregation occurs during a massive reorganization of cellular architecture at mitosis. Two gap-phases separate these cell cycle events: G1 between mitosis and S-phase, and G2 between S-phase and mitosis. Cells can exit the cell cycle for a period and enter a quiescent state known as G0, or terminally differentiate into cells that will not divide again, but undergo morphological development to carry out the wide variety of specialized functions of individual tissues.

A family of protein serine/threonine kinases known as the cyclin-dependent kinases (CDKs) controls progression through the cell cycle. As the name suggests, the kinase activity of the catalytic subunits is dependent on binding to cyclin partners, and control of cyclin abundance is one of several mechanisms by which CDK activity is regulated throughout the cell cycle.

A complex network of regulatory processes determines whether a quiescent cell (in G0 or early G1) will leave this state and initiate the processes to replicate its chromosomal DNA and divide. This regulation, during the **Mitotic G1-G1/S phases** of the cell cycle, centers on transcriptional regulation by the DREAM complex, with major roles for D and E type cyclin proteins.

Chromosomal DNA synthesis occurs in the **S phase**, or the synthesis phase, of the cell cycle. The cell duplicates its hereditary material, and two copies of each chromosome are formed. A key aspect of the **regulation of DNA** replication is the assembly and modification of a pre-replication complex assembled on ORC proteins.

**Mitotic G2-G2/M phases** encompass the interval between the completion of DNA synthesis and the beginning of mitosis. During G2, the cytoplasmic content of the cell increases. At G2/M transition, duplicated centrosomes mature and separate and CDK1:cyclin B complexes become active, setting the stage for spindle assembly and chromosome condensation at the start of mitotic **M phase**. Mitosis, or M phase, results in the generation of two daughter cells each with a complete diploid set of chromosomes. Events of the **M/G1 transition**, progression out of mitosis and division of the cell into two daughters (cytokinesis) are regulated by the Anaphase Promoting Complex.

The Anaphase Promoting Complex or Cyclosome (APC/C) plays additional roles in **regulation of the mitotic cell cycle**, insuring the appropriate length of the G1 phase. The APC/C itself is regulated by phosphorylation and interactions with checkpoint proteins.

#### References

#### Edit history

| Date | Action | Author |
|---|---|---|
| 2005-01-01 | Authored | Walworth N, Bosco G, O'Donnell M |
| 2005-01-01 | Created | Walworth N, Bosco G, O'Donnell M |
| 2010-01-19 | Revised | Matthews L |
| 2011-06-15 | Reviewed | Grana X |
| 2011-08-25 | Reviewed | MacPherson D |
| 2011-08-27 | Revised | Orlic-Milacic M |
| 2013-11-25 | Edited | Matthews L, Gopinathrao G |
| 2018-07-10 | Reviewed | Manfredi JJ |
| 2021-05-22 | Modified | Shorser S |

#### Entities found in this pathway (33)

| Input | UniProt Id | Input | UniProt Id | Input | UniProt Id |
|---|---|---|---|---|---|
| CCNB2 | O95067 | CDK1 | P06493, P24941 | CDK4 | P11802 |
| CDKN2C | P42773 | CENPF | P49454 | CENPU | Q71F23 |
| CHMP2A | O43633 | DHFR | P00374 | FBXO5 | Q9UKT4 |
| GINS2 | Q9Y248 | GINS3 | Q9BRX5 | H2AFJ | Q9BTM1 |
| HIST1H2AC | Q93077 | HIST1H3F | P68431 | HIST1H4C | P62805 |
| HSP90AA1 | P07900 | HSP90AB1 | P08238 | LBR | Q14739 |
| LMNA | P02545-1, P02545-2 | MCM10 | Q7L590 | MCM3 | P25205 |
| MCM4 | P33991 | MCM5 | P33992 | MCM7 | P33993 |
| ORC6 | Q9Y5N6 | PCNA | P12004 | PKMYT1 | Q99640 |
| POLA2 | Q14181 | PTTG1 | O95997 | RAD21 | O60216 |
| RRM2 | P31350 | TUBB | P04350, P07437 | TYMS | P04818 |

| Input | Ensembl Id | Input | Ensembl Id | Input | Ensembl Id |
|---|---|---|---|---|---|
| CCNB2 | ENSG00000157456 | CDK1 | ENSG00000170312 | CENPF | ENSG00000117724 |
| DHFR | ENSG00000228716 | FBXO5 | ENSG00000112029 | PCNA | ENSG00000132646 |
| RRM2 | ENSG00000171848 | TYMS | ENSG00000176890 | | |

#### 2. Cell Cycle (R-HSA-1640170)

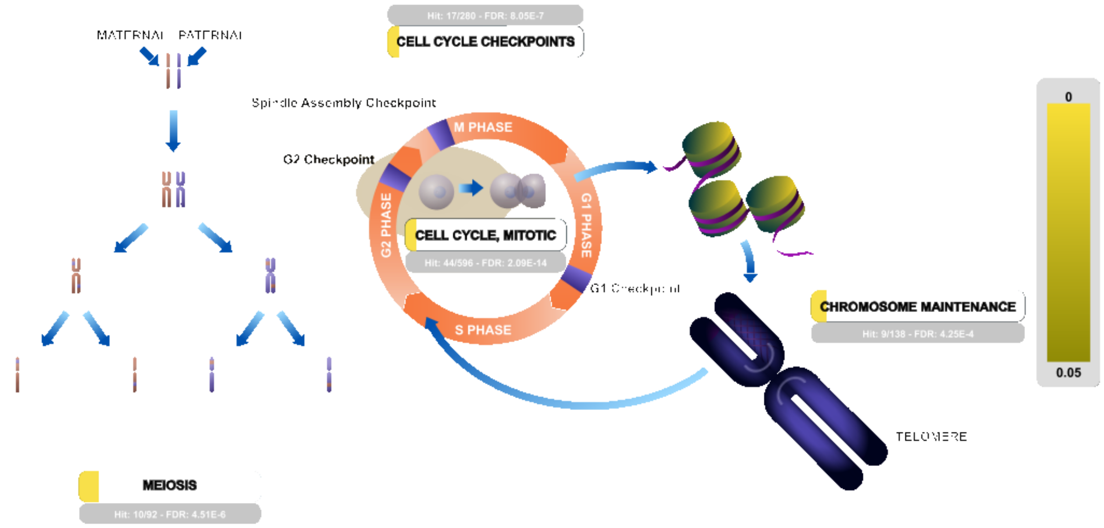


The replication of the genome and the subsequent segregation of chromosomes into daughter cells are controlled by a series of events collectively known as the **cell cycle**. DNA replication is carried out during a discrete temporal period known as the S (synthesis)-phase, and chromosome segregation occurs during a massive reorganization to cellular architecture at mitosis. Two gap-phases separate these major cell cycle events: G1 between mitosis and S-phase, and G2 between S-phase and mitosis. In the development of the human body, cells can exit the cell cycle for a period and enter a quiescent state known as G0, or terminally differentiate into cells that will not divide again, but undergo morphological development to carry out the wide variety of specialized functions of individual tissues.

A family of protein serine/threonine kinases known as the cyclin-dependent kinases (CDKs) controls progression through the cell cycle. As the name suggests, the activity of the catalytic subunit is dependent on binding to a cyclin partner. The human genome encodes several cyclins and several CDKs, with their names largely derived from the order in which they were identified. The oscillation of cyclin abundance is one important mechanism by which these enzymes phosphorylate key substrates to promote events at the relevant time and place. Additional post-translational modifications and interactions with regulatory proteins ensure that CDK activity is precisely regulated, frequently confined to a narrow window of activity.

In addition, genome integrity in the cell cycle is maintained by the action of a number of signal transduction pathways, known as **cell cycle checkpoints**, which monitor the accuracy and completeness of DNA replication during S phase and the orderly chromosomal condensation, pairing and partition into daughter cells during mitosis.

Replication of telomeric DNA at the ends of human chromosomes and packaging of their centromeres into chromatin are two aspects of **chromosome maintenance** that are integral parts of the cell cycle.

**Meiosis** is the specialized form of cell division that generates haploid gametes from diploid germ cells, associated with recombination (exchange of genetic material between chromosomal homologs).

#### References

#### Edit history

| Date | Action | Author |
|---|---|---|
| 2011-10-10 | Edited | Matthews L |
| 2011-10-10 | Created | Matthews L |
| 2021-05-22 | Modified | Shorser S |

#### Entities found in this pathway (40)

| Input | UniProt Id | Input | UniProt Id | Input | UniProt Id |
|---|---|---|---|---|---|
| BLM | P54132 | BRCA1 | P38398 | BRCA2 | P51587 |
| BRIP1 | Q9BX63 | CCNB2 | O95067 | CDK1 | P06493, P24941 |
| CDK4 | P11802 | CDKN2C | P42773 | CENPF | P49454 |
| CENPU | Q71F23 | CENPX | A8MT69 | CHMP2A | O43633 |
| DHFR | P00374 | EXO1 | Q9UQ84 | FBXO5 | Q9UKT4 |
| GINS2 | Q9Y248 | GINS3 | Q9BRX5 | H2AFJ | Q9BTM1 |
| HIST1H2AC | Q93077 | HIST1H3F | P68431 | HIST1H4C | P62805 |
| HSP90AA1 | P07900 | HSP90AB1 | P08238 | LBR | Q14739 |
| LMNA | P02545-1, P02545-2 | MCM10 | Q7L590 | MCM3 | P25205 |
| MCM4 | P33991 | MCM5 | P33992 | MCM7 | P33993 |
| ORC6 | Q9Y5N6 | PCNA | P12004 | PKMYT1 | Q99640 |
| POLA2 | Q14181 | PTTG1 | O95997 | RAD21 | O60216 |
| RRM2 | P31350 | TUBB | P04350, P07437 | TYMS | P04818 |
| YWHAZ | P63104 | | | | |

| Input | Ensembl Id | Input | Ensembl Id | Input | Ensembl Id |
|---|---|---|---|---|---|
| CCNB2 | ENSG00000157456 | CDK1 | ENSG00000170312 | CENPF | ENSG00000117724 |
| DHFR | ENSG00000228716 | FBXO5 | ENSG00000112029 | PCNA | ENSG00000132646 |
| RRM2 | ENSG00000171848 | TYMS | ENSG00000176890 | | |

#### 3. G1/S Transition (R-HSA-69206)

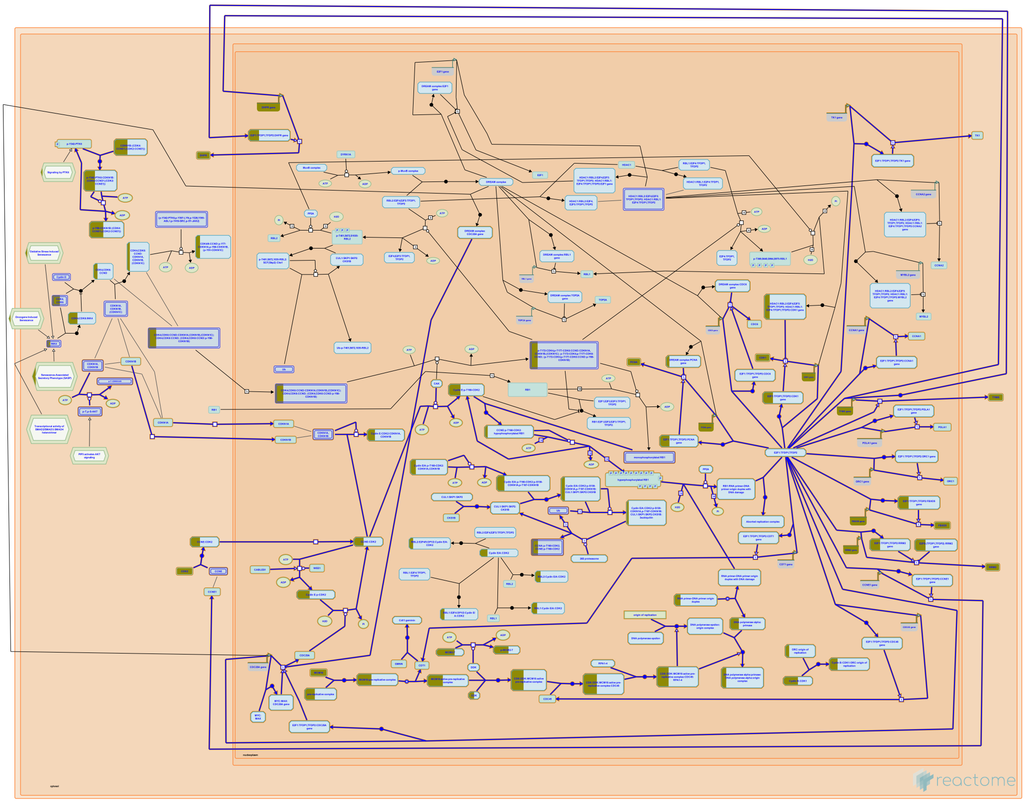

Cyclin E - Cdk2 complexes control the transition from G1 into S-phase. In this case, the binding of p21Cip1/Waf1 or p27kip1 is inhibitory. Important substrates for Cyclin E - Cdk2 complexes include proteins involved in the initiation of DNA replication. The two Cyclin E proteins are subjected to ubiquitin-dependent proteolysis, under the control of an E3 ubiquitin ligase known as the SCF. Cyclin A - Cdk2 complexes, which are also regulated by p21Cip1/Waf1 and p27kip1, are likely to be important for continued DNA synthesis, and progression into G2. An additional level of control of Cdk2 is reversible phosphorylation of Threonine-14 (T14) and Tyrosine-15 (Y15), catalyzed by the Wee1 and Myt1 kinases, and dephosphorylation by the three Cdc25 phosphatases, Cdc25A, B and C.

##### References

##### Edit history

| Date | Action | Author |
|---|---|---|
| 2003-06-05 | Created | Walworth N, O'Donnell M |
| 2021-05-22 | Modified | Shorser S |

##### Entities found in this pathway (14)

| Input | UniProt Id | Input | UniProt Id | Input | UniProt Id |
|---|---|---|---|---|---|
| CDK1 | P06493, P24941 | CDK4 | P11802 | DHFR | P00374 |
| FBXO5 | Q9UKT4 | MCM10 | Q7L590 | MCM3 | P25205 |
| MCM4 | P33991 | MCM5 | P33992 | MCM7 | P33993 |
| ORC6 | Q9Y5N6 | PCNA | P12004 | POLA2 | Q14181 |
| RRM2 | P31350 | TYMS | P04818 | | |

| Input | Ensembl Id | Input | Ensembl Id | Input | Ensembl Id |
|---|---|---|---|---|---|
| CDK1 | ENSG00000170312 | DHFR | ENSG00000228716 | FBXO5 | ENSG00000112029 |
| PCNA | ENSG00000132646 | RRM2 | ENSG00000171848 | TYMS | ENSG00000176890 |

#### 4. Mitotic G1 phase and G1/S transition (R-HSA-453279)

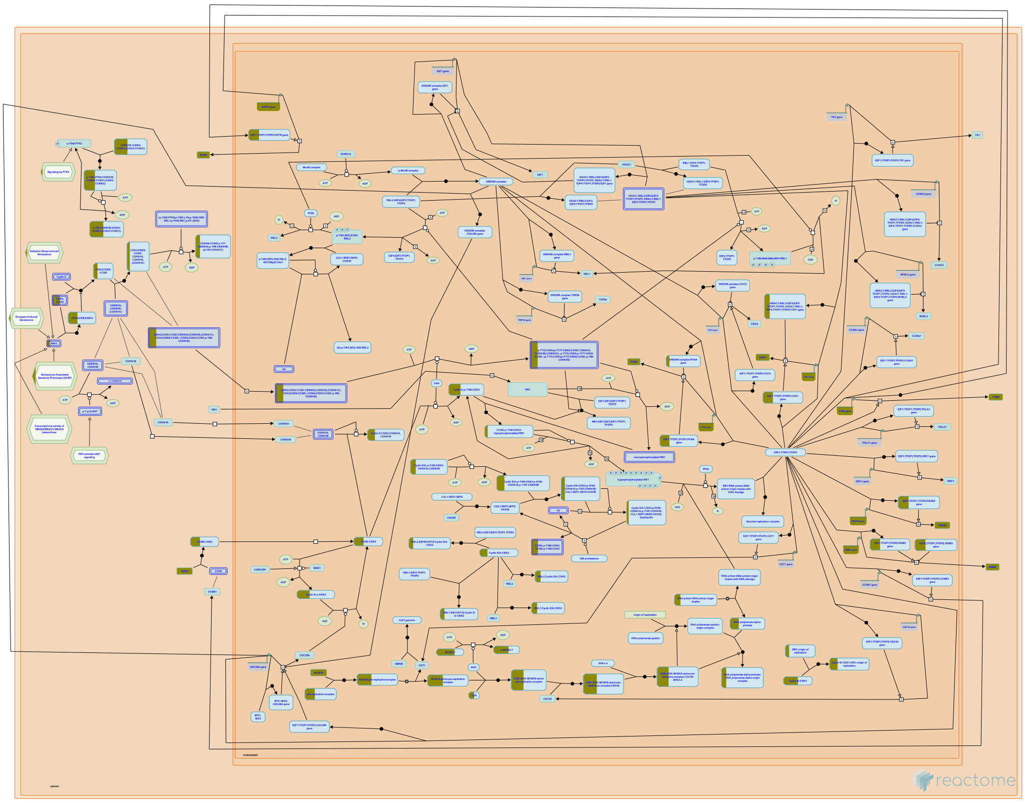

Mitotic G1-G1/S phase involves G1 phase of the mitotic interphase and G1/S transition, when a cell commits to DNA replication and divison genetic and cellular material to two daughter cells.

During early G1, cells can enter a quiescent G0 state. In quiescent cells, the evolutionarily conserved DREAM complex, consisting of the pocket protein family member p130 (RBL2), bound to E2F4 or E2F5, and the MuvB complex, represses transcription of cell cycle genes (reviewed by Sadasivam and DeCaprio 2013).

During early G1 phase in actively cycling cells, transcription of cell cycle genes is repressed by another pocket protein family member, p107 (RBL1), which forms a complex with E2F4 (Ferreira et al. 1998, Cobrinik 2005). RB1 tumor suppressor, the product of the retinoblastoma susceptibility gene, is the third member of the pocket protein family. RB1 binds to E2F transcription factors E2F1, E2F2 and E2F3 and inhibits their transcriptional activity, resulting in prevention of G1/S transition (Chellappan et al. 1991, Bagchi et al. 1991, Chittenden et al. 1991, Lees et al. 1993, Hiebert 1993, Wu et al. 2001). Once RB1 is phosphorylated on serine residue S795 by Cyclin D:CDK4/6 complexes, it can no longer associate with and inhibit E2F1-3. Thus, CDK4/6-mediated phosphorylation of RB1 leads to transcriptional activation of E2F1-3 target genes needed for the S phase of the cell cycle (Connell-Crowley et al. 1997). CDK2, in complex with cyclin E, contributes to RB1 inactivation and also activates proteins needed for the initiation of DNA replication (Zhang 2007). Expression of D type cyclins is regulated by extracellular mitogens (Cheng et al. 1998, Depoortere et al. 1998). Catalytic activities of CDK4/6 and CDK2 are controlled by CDK inhibitors of the INK4 family (Serrano et al. 1993, Hannon and Beach 1994, Guan et al. 1994, Guan et al. 1996, Parry et al. 1995) and the Cip/Kip family, respectively.

#### Edit history

| Date | Action | Author |
|---|---|---|
| 2010-01-19 | Edited | Matthews L |
| 2010-01-20 | Authored | Matthews L |
| 2010-01-20 | Created | Matthews L |
| 2011-06-15 | Reviewed | Grana X |
| 2011-08-25 | Reviewed | MacPherson D |
| 2011-08-26 | Revised | Orlic-Milacic M |
| 2011-08-26 | Authored | Orlic-Milacic M |
| 2017-02-08 | Edited | Orlic-Milacic M |
| 2018-07-10 | Reviewed | Manfredi JJ |
| 2021-05-22 | Modified | Shorser S |

#### Entities found in this pathway (15)

| Input | UniProt Id | Input | UniProt Id | Input | UniProt Id |
|---|---|---|---|---|---|
| CDK1 | P06493, P24941 | CDK4 | P11802 | CDKN2C | P42773 |
| DHFR | P00374 | FBXO5 | Q9UKT4 | MCM10 | Q7L590 |
| MCM3 | P25205 | MCM4 | P33991 | MCM5 | P33992 |
| MCM7 | P33993 | ORC6 | Q9Y5N6 | PCNA | P12004 |
| POLA2 | Q14181 | RRM2 | P31350 | TYMS | P04818 |

| Input | Ensembl Id | Input | Ensembl Id | Input | Ensembl Id |
|---|---|---|---|---|---|
| CDK1 | ENSG00000170312 | DHFR | ENSG00000228716 | FBXO5 | ENSG00000112029 |
| PCNA | ENSG00000132646 | RRM2 | ENSG00000171848 | TYMS | ENSG00000176890 |

#### 5. G1/S-Specific Transcription (R-HSA-69205)

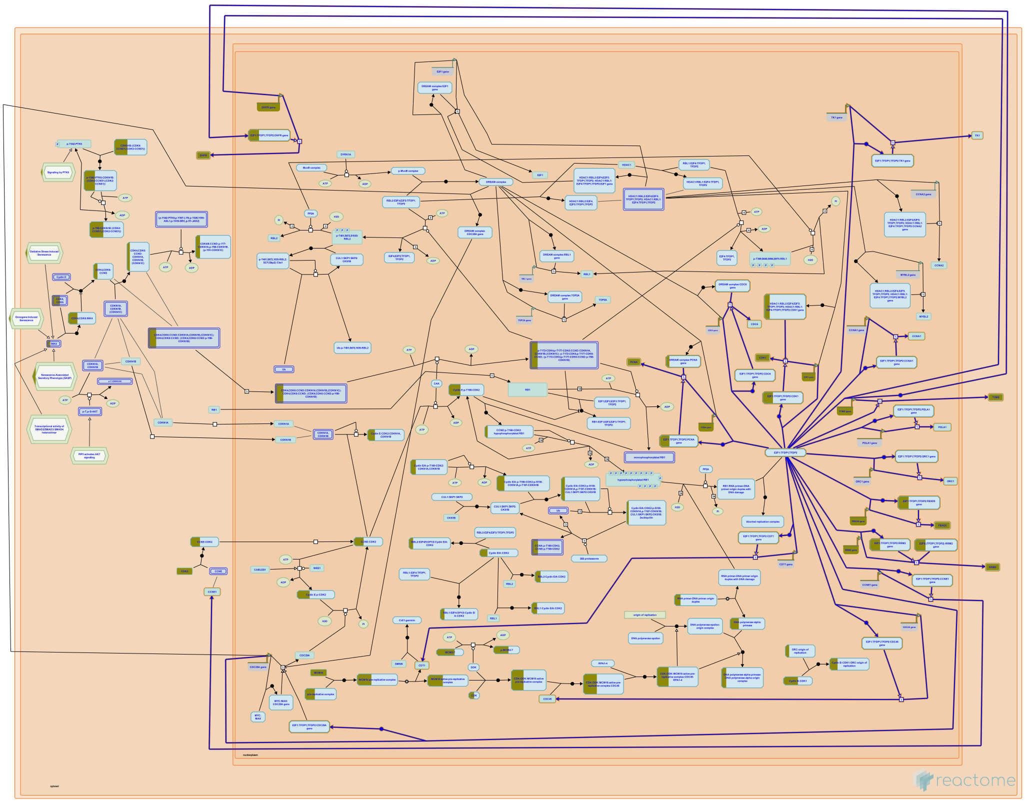

**Cellular compartments:** nucleoplasm.

The E2F family of transcription factors regulate the transition from the G1 to the S phase in the cell cycle. E2F activity is regulated by members of the retinoblastoma protein (pRb) family, resulting in the tight control of the expression of E2F-responsive genes. Phosphorylation of pRb by cyclin D:CDK complexes releases pRb from E2F, inducing E2F-targeted genes such as cyclin E.

E2F1 binds to E2F binding sites on the genome activating the synthesis of the target proteins. For annotation purposes, the reactions regulated by E2F1 are grouped under this pathway and information about the target genes alone are displayed for annotation purposes.

Cellular targets for activation by E2F1 include thymidylate synthase (TYMS) (DeGregori et al. 1995), Rir2 (RRM2) (DeGregori et al. 1995, Giangrande et al. 2004), Dihydrofolate reductase (DHFR) (DeGregori et al. 1995, Wells et al. 1997, Darbinian et al. 1999), Cdc2 (CDK1) (Furukawa et al. 1994, DeGregori et al. 1995, Zhu et al. 2004), Cyclin A1 (CCNA1) (DeGregori et al. 1995, Liu et al. 1998), CDC6 (DeGregori et al. 1995, Yan et al. 1998; Ohtani et al. 1998), CDT1 (Yoshida and Inoue 2004), CDC45 (Arata et al. 2000), Cyclin E (CCNE1) (Ohtani et al. 1995), Emi1 (FBXO5) (Hsu et al. 2002), and ORC1 (Ohtani et al. 1996, Ohtani et al. 1998). The activation of TK1 (Dnk1) (Dou et al. 1994, DeGregori et al. 1995, Giangrande et al. 2004) and CDC25A (DeGregori et al. 1995, Vigo et al. 1999) by E2F1 is conserved in Drosophila (Duronio and O'Farrell 1994, Reis and Edgar 2004).

RRM2 protein is involved in dNTP level regulation and activation of this enzyme results in higher levels of dNTPs in anticipation of S phase. E2F activation of RRM2 has been shown also in Drosophila by Duronio and O'Farrell (1994). E2F1 activation of CDC45 is shown in mouse cells by using human E2F1 construct (Arata et al. 2000). Cyclin E is also transcriptionally regulated by E2F1. Cyclin E protein plays important role in the transition of G1 in S phase by associating with CDK2 (Ohtani et al. 1996). E2F1-mediated activation of PCNA has been demonstrated in Drosophila (Duronio and O'Farrell 1994) and in some human cells by using recombinant adenovirus constructs (DeGregori et al. 1995). E2F1-mediated activation of the DNA polymerase alpha subunit p180 (POLA1) has been demonstrated in some human cells. It has also been demonstrated in Drosophila by Ohtani and Nevins (1994). It has been observed in Drosophila that E2F1 induced expression of Orc1 stimulates ORC1 6 complex formation and binding to the origin of replication (Asano and Wharton 1999). ORC1 6 recruit CDC6 and CDT1 that are required to recruit the MCM2 7 replication helicases. E2F1 regulation incorporates a feedback mechanism wherein Geminin (GMNN) can inhibit MCM2 7 recruitment of ORC1 6 complex by interacting with CDC6/CDT1. The activation of CDC25A and TK1 (Dnk1) by E2F1 has been inferred from similar events in Drosophila (Duronio RJ and O'Farrell 1994; Reis and Edgar 2004). E2F1 activates string (CDC25) that in turn activates the complex of Cyclin B and CDK1. A similar phenomenon has been observed in mouse NIH 3T3 cells and in Rat1 cells.


#### Edit history

| Date | Action | Author |
|---|---|---|
| 2003-06-05 | Created | Walworth N, O'Donnell M |
| 2018-12-21 | Modified | D'Eustachio P |

#### Entities found in this pathway (6)

| Input | UniProt Id | Input | UniProt Id | Input | UniProt Id |
|---|---|---|---|---|---|
| CDK1 | P06493 | DHFR | P00374 | FBXO5 | Q9UKT4 |
| PCNA | P12004 | RRM2 | P31350 | TYMS | P04818 |

| Input | Ensembl Id | Input | Ensembl Id | Input | Ensembl Id |
|---|---|---|---|---|---|
| CDK1 | ENSG00000170312 | DHFR | ENSG00000228716 | FBXO5 | ENSG00000112029 |

| Input | Ensembl Id | Input | Ensembl Id | Input | Ensembl Id |
|---|---|---|---|---|---|
| PCNA | ENSG00000132646 | RRM2 | ENSG00000171848 | TYMS | ENSG00000176890 |

#### 6. G2/M Checkpoints (R-HSA-69481)

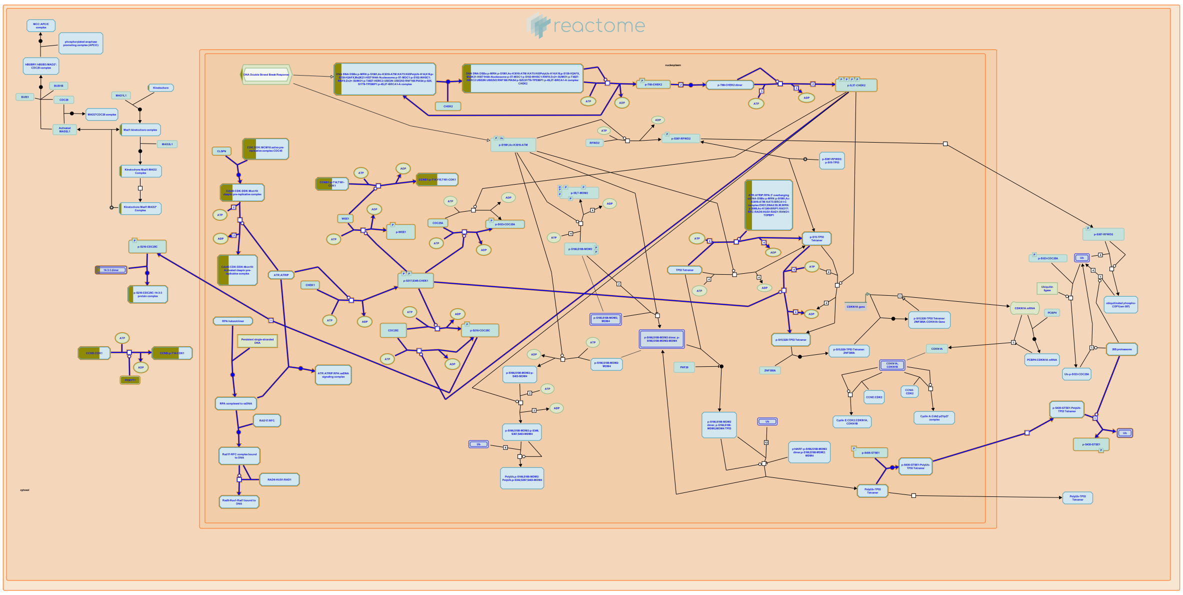

G2/M checkpoints include the checks for damaged DNA, unreplicated DNA, and checks that ensure that the genome is replicated once and only once per cell cycle. If cells pass these checkpoints, they follow normal transition to the M phase. However, if any of these checkpoints fail, mitotic entry is prevented by specific G2/M checkpoint events.

The G2/M checkpoints can fail due to the presence of unreplicated DNA or damaged DNA. In such instances, the cyclin-dependent kinase, Cdc2(Cdk1), is maintained in its inactive, phosphorylated state, and mitotic entry is prevented. Events that ensure that origins of DNA replication fire once and only once per cell cycle are also an example of a G2/M checkpoint.

In the event of high levels of DNA damage, the cells may also be directed to undergo apoptosis (not covered).

##### References

##### Edit history

| Date | Action | Author |
|---|---|---|
| 2003-06-05 | Created | Walworth N, O'Donnell M |
| 2021-05-22 | Modified | Shorser S |

##### Entities found in this pathway (15)

| Input | UniProt Id | Input | UniProt Id | Input | UniProt Id |
|---|---|---|---|---|---|
| BLM | P54132 | BRCA1 | P38398 | BRIP1 | Q9BX63 |
| CCNB2 | O95067 | CDK1 | P06493 | EXO1 | Q9UQ84 |
| HIST1H4C | P62805 | MCM10 | Q7L590 | MCM3 | P25205 |
| MCM4 | P33991 | MCM5 | P33992 | MCM7 | P33993 |
| ORC6 | Q9Y5N6 | PKMYT1 | Q99640 | YWHAZ | P63104 |

#### 7. Unwinding of DNA (R-HSA-176974)

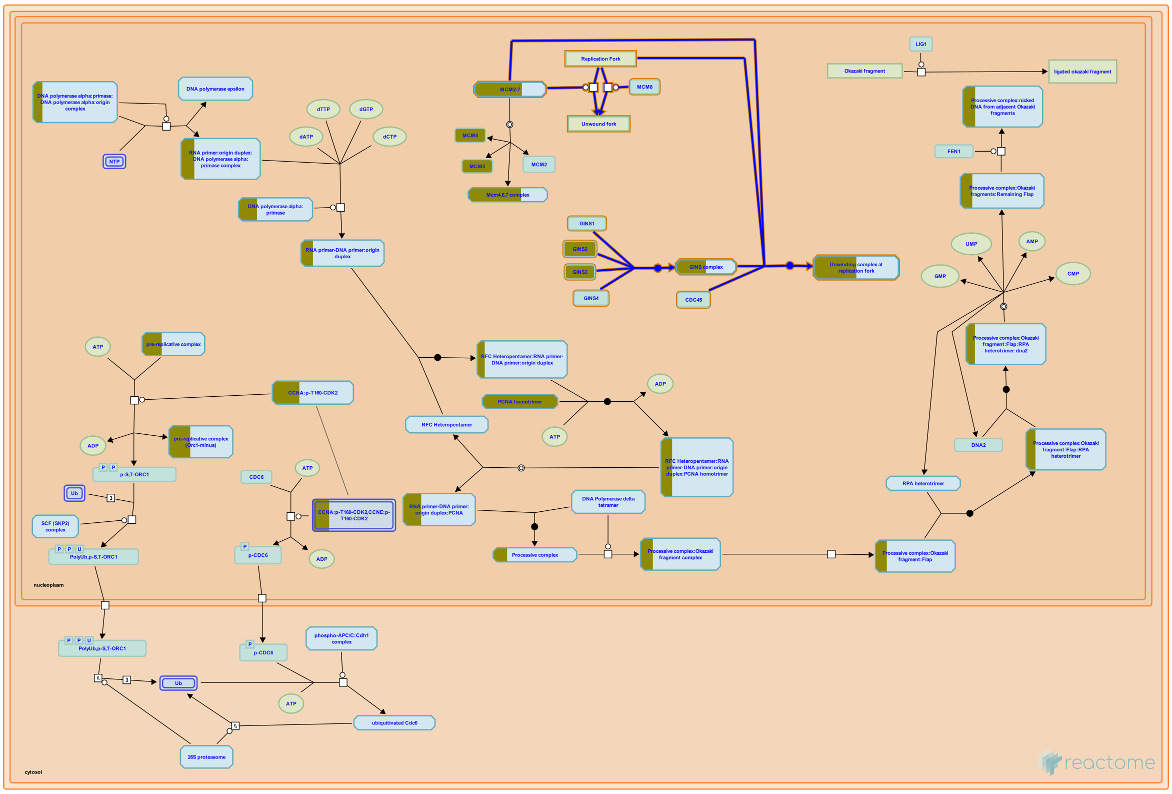


**Cellular compartments:** nucleoplasm.

DNA Replication is regulated accurately and precisely by various protein complexes. Many members of the MCM protein family are assembled into the pre-Replication Complexes (pre-RC) at the end of M phase of the cell cycle. DNA helicase activity of some of the MCM family proteins are important for the unwinding of DNA and initiation of replication processes. This section contains four events which have been proved in different eukaryotic experimental systems to involve various proteins for this essential step during DNA Replication.

##### Edit history

| Date | Action | Author |
|---|---|---|
| 2006-03-17 | Edited | Gopinathrao G |
| 2006-03-17 | Authored | Tye BK |
| 2006-03-18 | Created | Gopinathrao G |
| 2021-05-22 | Modified | Shorser S |

##### Entities found in this pathway (6)

| Input | UniProt Id | Input | UniProt Id | Input | UniProt Id |
|---|---|---|---|---|---|
| GINS2 | Q9Y248 | GINS3 | Q9BRX5 | MCM3 | P25205 |
| MCM4 | P33991 | MCM5 | P33992 | MCM7 | P33993 |

#### 8. DNA strand elongation (R-HSA-69190)

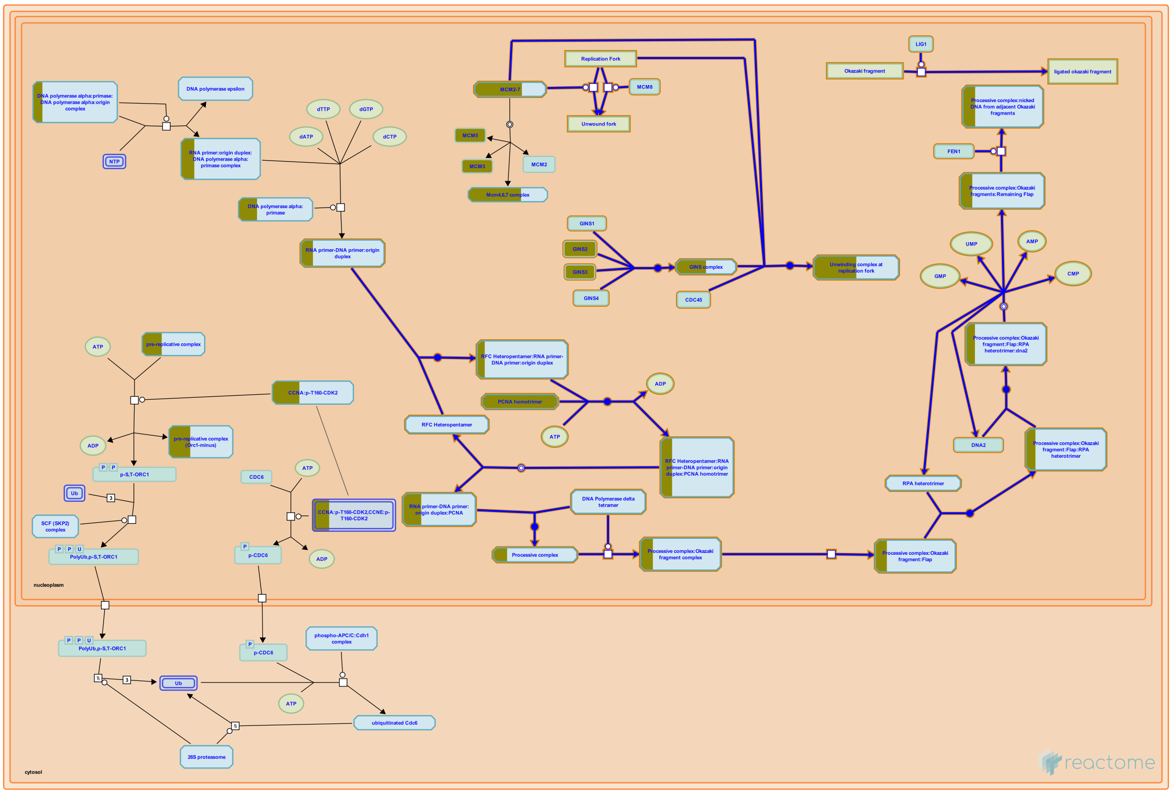

**Cellular compartments:** nucleoplasm.

Accurate and efficient genome duplication requires coordinated processes to replicate two template strands at eucaryotic replication forks. Knowledge of the fundamental reactions involved in replication fork progression is derived largely from biochemical studies of the replication of simian virus and from yeast genetic studies. Since duplex DNA forms an anti-parallel structure, and DNA polymerases are unidirectional, one of the new strands is synthesized continuously in the direction of fork movement. This strand is designated as the leading strand. The other strand grows in the direction away from fork movement, and is called the lagging strand. Several specific interactions among the various proteins involved in DNA replication underlie the mechanism of DNA synthesis, on both the leading and lagging strands, at a DNA replication fork. These interactions allow the replication enzymes to cooperate in the replication process (Hurwitz et al 1990; Brush et al 1996; Ayyagari et al 1995; Budd & Campbell 1997; Bambara et al 1997).

#### Edit history

| Date | Action | Author |
|---|---|---|
| 2003-06-05 | Authored | Tom S, Bambara RA |
| 2003-06-05 | Created | Tom S, Bambara RA |
| 2021-05-22 | Modified | Shorser S |

#### Entities found in this pathway (8)

| Input | UniProt Id | Input | UniProt Id | Input | UniProt Id |
|---|---|---|---|---|---|
| GINS2 | Q9Y248 | GINS3 | Q9BRX5 | MCM3 | P25205 |
| MCM4 | P33991 | MCM5 | P33992 | MCM7 | P33993 |
| PCNA | P12004 | POLA2 | Q14181 | | |

#### 9. Cell Cycle Checkpoints (R-HSA-69620)

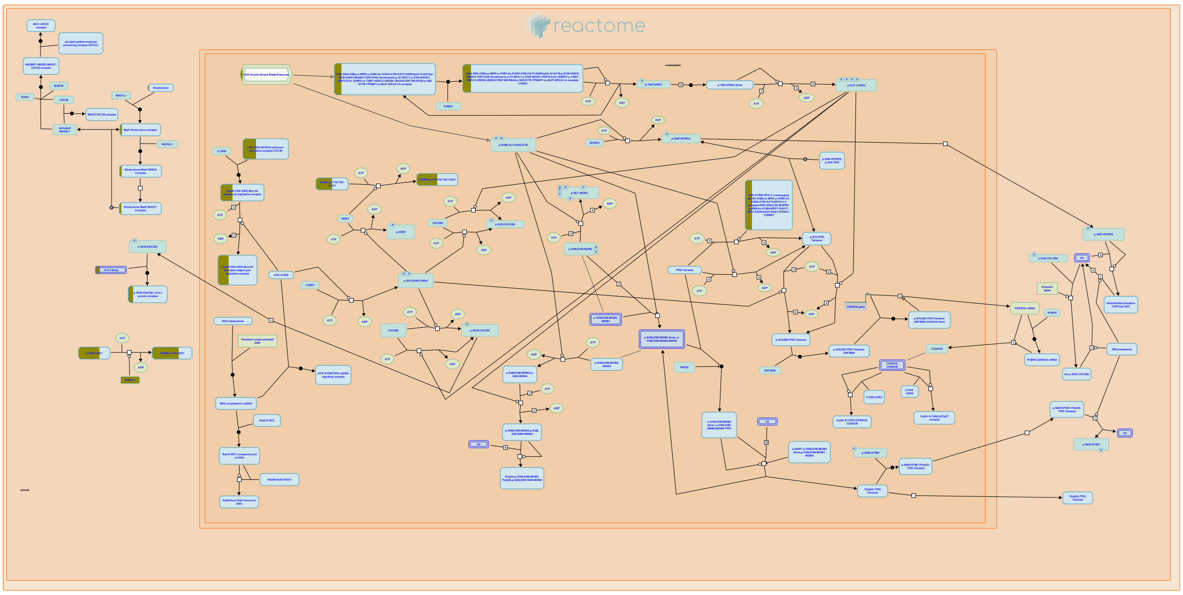

A hallmark of the human cell cycle in normal somatic cells is its precision. This remarkable fidelity is achieved by a number of signal transduction pathways, known as checkpoints, which monitor cell cycle progression ensuring an interdependency of S-phase and mitosis, the integrity of the genome and the fidelity of chromosome segregation.

Checkpoints are layers of control that act to delay CDK activation when defects in the division program occur. As the CDKs functioning at different points in the cell cycle are regulated by different means, the various checkpoints differ in the biochemical mechanisms by which they elicit their effect. However, all checkpoints share a common hierarchy of a sensor, signal transducers, and effectors that interact with the CDKs.

The stability of the genome in somatic cells contrasts to the almost universal genomic instability of tumor cells. There are a number of documented genetic lesions in checkpoint genes, or in cell cycle genes themselves, which result either directly in cancer or in a predisposition to certain cancer types. Indeed, restraint over cell cycle progression and failure to monitor genome integrity are likely prerequisites for the molecular evolution required for the development of a tumor. Perhaps most notable amongst these is the p53 tumor suppressor gene, which is mutated in >50% of human tumors. Thus, the importance of the checkpoint pathways to human biology is clear.

##### References

##### Edit history

| Date | Action | Author |
|---|---|---|
| 2005-01-01 | Authored | Walworth N, Hoffmann I, Yen TJ, O'Donnell M, Khanna KK |
| 2005-01-01 | Created | Walworth N, Hoffmann I, Yen TJ, O'Donnell M, Khanna KK |
| 2013-11-25 | Edited | Matthews L |
| 2021-05-18 | Reviewed | Sanchez Y, Knudsen E, Hardwick KG |
| 2021-05-22 | Modified | Shorser S |

##### Entities found in this pathway (17)

| Input | UniProt Id | Input | UniProt Id | Input | UniProt Id |
|---|---|---|---|---|---|
| BLM | P54132 | BRCA1 | P38398 | BRIP1 | Q9BX63 |
| CCNB2 | O95067 | CDK1 | P06493 | CENPF | P49454 |
| CENPU | Q71F23 | EXO1 | Q9UQ84 | HIST1H4C | P62805 |
| MCM10 | Q7L590 | MCM3 | P25205 | MCM4 | P33991 |
| MCM5 | P33992 | MCM7 | P33993 | ORC6 | Q9Y5N6 |
| PKMYT1 | Q99640 | YWHAZ | P63104 | | |

#### 10. Meiosis (R-HSA-1500620)

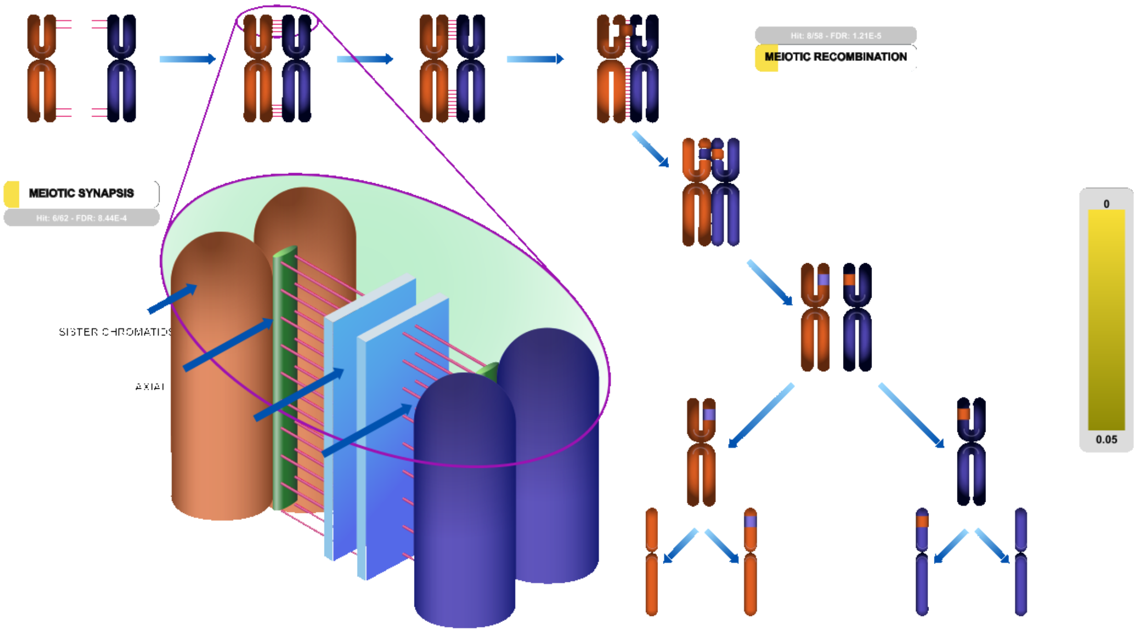


**Cellular compartments:** nuclear envelope, nucleoplasm.

During meiosis the replicated chromosomes of a single diploid cell are segregated into 4 haploid daughter cells by two successive divisions, meiosis I and meiosis II. In meiosis I, the distinguishing event of meiosis, pairs (bivalents) of homologous chromosomes in the form of sister chromatids are paired by **synapsis** along their regions of homologous DNA (Yang and Wang 2009), and then segregated, resulting in haploid daughters containing sister chromatids paired at their centromeres (Cohen et al. 2006, Handel and Schimenti 2010). The sister chromatids are then separated and segregated during meiosis II.

**Recombination** between chromosomal homologues but not between sister chromatids occurs during prophase of meiosis I (Inagaki et al. 2010). Though hundreds of recombination events are initiated, most are resolved without crossovers and only tens proceed to become crossovers. In mammals recombination events are required between homologues for normal pairing, synapsis, and segregation.

##### Edit history

| Date | Action | Author |
|---|---|---|
| 2011-02-05 | Reviewed | Cohen PE, Schimenti JC, Holloway JK |
| 2011-02-25 | Reviewed | Lyndaker A, Bolcun-Filas E, Strong E |
| 2011-08-19 | Edited | May B |
| 2011-08-19 | Authored | May B |
| 2011-08-19 | Created | May B |
| 2018-11-14 | Modified | Matthews L |

#### Entities found in this pathway (10)

| Input | UniProt Id | Input | UniProt Id | Input | UniProt Id |
|---|---|---|---|---|---|
| BLM | P54132 | BRCA1 | P38398 | BRCA2 | P51587 |
| CDK4 | P11802 | H2AFJ | Q9BTM1 | HIST1H2AC | Q93077 |
| HIST1H3F | P68431 | HIST1H4C | P62805 | LMNA | P02545-2 |
| RAD21 | O60216 | | | | |

#### 11. Activation of the pre-replicative complex (R-HSA-68962)

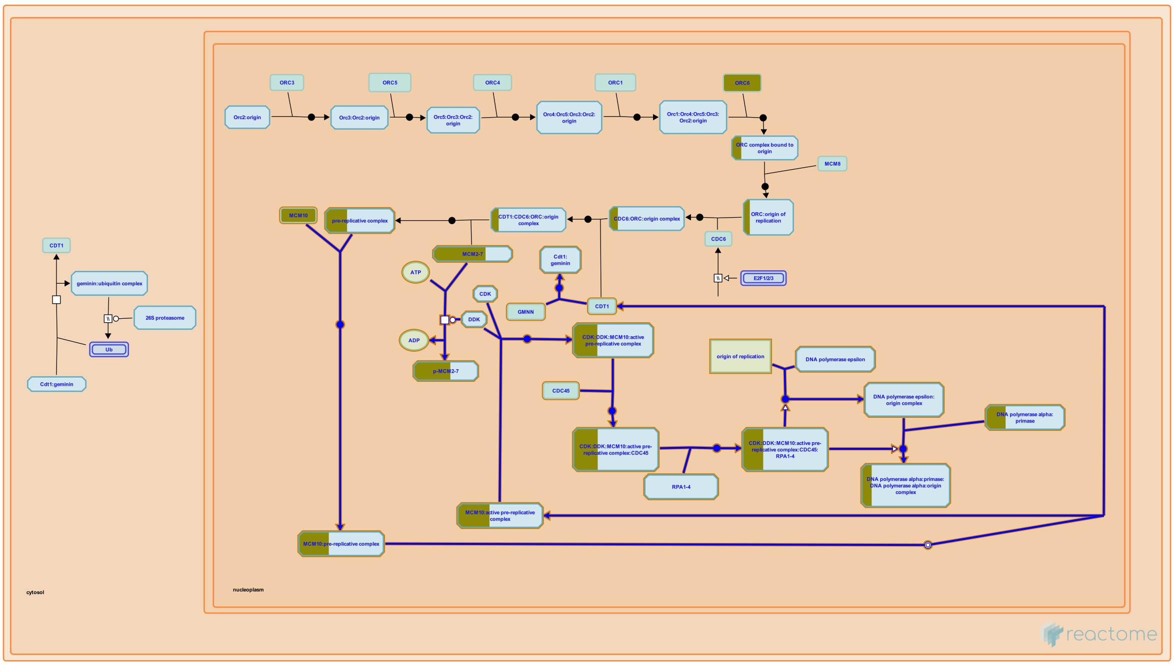


**Cellular compartments:** nucleoplasm.

In S. cerevisiae, two ORC subunits, Orc1 and Orc5, both bind ATP, and Orc1 in addition has ATPase activity. Both ATP binding and ATP hydrolysis appear to be essential functions in vivo. ATP binding by Orc1 is unaffected by the association of ORC with origin DNA (ARS) sequences, but ATP hydrolysis is ARS-dependent, being suppressed by associated double-stranded DNA and stimulated by associated single-stranded DNA. These data are consistent with the hypothesis that ORC functions as an ATPase switch, hydrolyzing bound ATP and changing state as DNA unwinds at the origin immediately before replication. It is attractive to speculate that ORC likewise functions as a switch as human pre-replicative complexes are activated, but human Orc proteins are not well enough characterized to allow the model to be critically tested. mRNAs encoding human orthologs of all six Orc proteins have been cloned, and ATP-binding amino acid sequence motifs have been identified in Orc1, Orc4, and Orc5. Interactions among proteins expressed from the cloned genes have been characterized, but the ATP-binding and hydrolyzing properties of these proteins and complexes of them have not been determined.

#### Edit history

| Date | Action | Author |
|---|---|---|
| 2003-06-05 | Created | Davey MJ, O'Donnell M |
| 2021-05-22 | Modified | Shorser S |

#### Entities found in this pathway (7)

| Input | UniProt Id | Input | UniProt Id | Input | UniProt Id |
|---|---|---|---|---|---|
| MCM10 | Q7L590 | MCM3 | P25205 | MCM4 | P33991 |
| MCM5 | P33992 | MCM7 | P33993 | ORC6 | Q9Y5N6 |
| POLA2 | Q14181 | | | | |

#### 12. Meiotic recombination (R-HSA-912446)

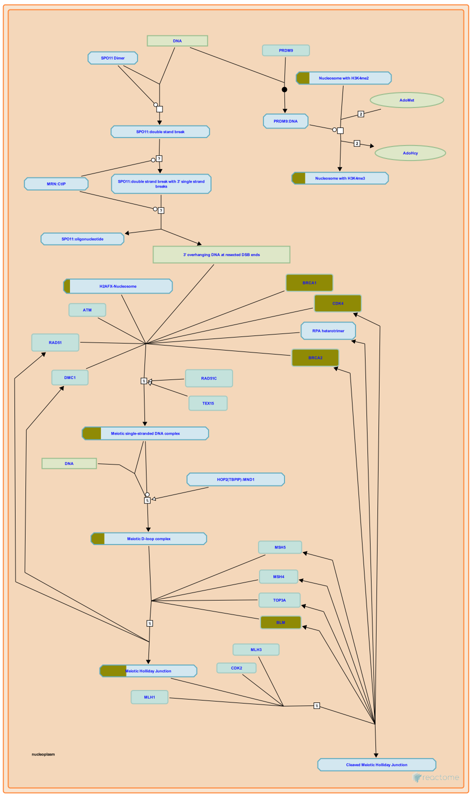

**Cellular compartments:** nucleoplasm.

Meiotic recombination exchanges segments of duplex DNA between chromosomal homologs, generating genetic diversity (reviewed in Handel and Schimenti 2010, Inagaki et al. 2010, Cohen et al. 2006). There are two forms of recombination: non-crossover (NCO) and crossover (CO). In mammals, the former is required for correct pairing and synapsis of homologous chromosomes, while CO intermediates called chiasmata are required for correct segregation of bivalents.

Meiotic recombination is initiated by double-strand breaks created by SPO11, which remains covalently attached to the 5' ends after cleavage. SPO11 is removed by cleavage of single DNA strands adjacent to the covalent linkage. The resulting 5' ends are further resected to produce protruding 3' ends. The single-stranded 3' ends are bound by RAD51 and DMC1, homologs of RecA that catalyze a search for homology between the bound single strand and duplex DNA of the chromosomal homolog. RAD51 and DMC1 then catalyze the invasion of the single strand into the homologous duplex and the formation of a D-loop heteroduplex. Approximately 90% of heteroduplexes are resolved without crossovers (NCO), probably by synthesis-dependent strand annealing.

The invasive strand is extended along the homolog and ligated back to its original duplex, creating a double Holliday junction. The mismatch repair proteins MSH4, MSH5 participate in this process, possibly by stabilizing the duplexes. The mismatch repair proteins MLH1 and MLH3 are then recruited to the double Holliday structure and an unidentified resolvase (Mus81? Gen1?) cleaves the junctions to yield a crossover.

Crossovers are not randomly distributed: The histone methyltransferase PRDM9 recruits the recombination machinery to genetically determined hotspots in the genome and each incipient crossover somehow inhibits formation of crossovers nearby, a phenomenon called crossover interference. Each chromosome bivalent, including the X-Y body in males, has at least one crossover and this is required for meiosis to proceed correctly.


##### Edit history

| Date | Action | Author |
|---|---|---|
| 2010-07-03 | Edited | May B |
| 2010-07-03 | Authored | May B |
| 2010-07-09 | Created | May B |
| 2011-02-05 | Reviewed | Cohen PE, Schimenti JC, Holloway JK |
| 2011-02-25 | Reviewed | Lyndaker A, Bolcun-Filas E, Strong E |
| 2017-09-29 | Modified | May B |

##### Entities found in this pathway (8)

| Input | UniProt Id | Input | UniProt Id | Input | UniProt Id |
|---|---|---|---|---|---|
| BLM | P54132 | BRCA1 | P38398 | BRCA2 | P51587 |
| CDK4 | P11802 | H2AFJ | Q9BTM1 | HIST1H2AC | Q93077 |
| HIST1H3F | P68431 | HIST1H4C | P62805 | | |

#### 13. DNA Replication (R-HSA-69306)

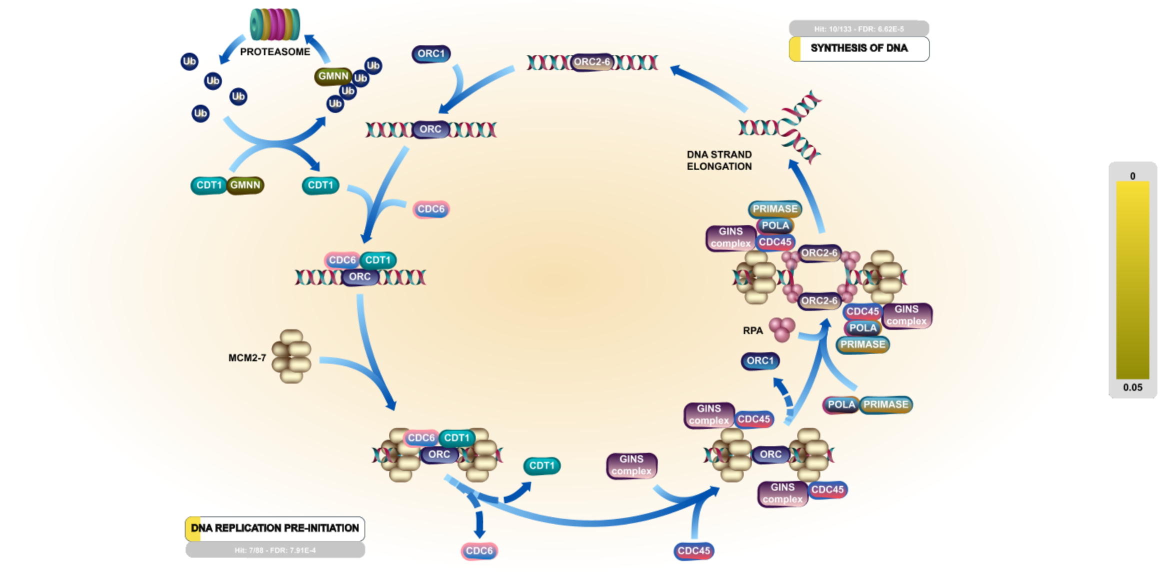


**Cellular compartments:** nucleoplasm, cytosol.

Studies in the past decade have suggested that the basic mechanism of DNA replication initiation is conserved in all kingdoms of life. Initiation in unicellular eukaryotes, in particular Saccharomyces cerevisiae (budding yeast), is well understood, and has served as a model for studies of DNA replication initiation in multicellular eukaryotes, including humans. In general terms, the first step of initiation is the binding of the replication initiator to the origin of replication. The replicative helicase is then assembled onto the origin, usually by a helicase assembly factor. Either shortly before or shortly after helicase assembly, some local unwinding of the origin of replication occurs in a region rich in adenine and thymine bases (often termed a DNA unwinding element, DUE). The unwound region provides the substrate for primer synthesis and initiation of DNA replication. The best-defined eukaryotic origins are those of S. cerevisiae, which have well-conserved sequence elements for initiator binding, DNA unwinding and binding of accessory proteins. In multicellular eukaryotes, unlike S. cerevisiae, these loci appear not to be defined by the presence of a DNA sequence motif. Indeed, choice of replication origins in a multicellular eukaryote may vary with developmental stage and tissue type. In cell-free models of metazoan DNA replication, such as the one provided by Xenopus egg extracts, there are only limited DNA sequence specificity requirements for replication initiation (Kelly & Brown 2000; Bell & Dutta 2002; Marahrens & Stillman 1992; Cimbora & Groudine 2001; Mahbubani et al 1992, Hyrien & Mechali 1993).

#### Edit history

| Date | Action | Author |
|---|---|---|
| 2003-01-06 | Authored | Catlett M, Davey MJ, Tye BK, O'Donnell M, Forsburg SL et al. |
| 2003-01-06 | Created | Catlett M, Davey MJ, Tye BK, O'Donnell M, Forsburg SL et al. |
| 2005-09-07 | Revised | Tye BK, Borowiec JA, Mendez J, Aladjem M |
| 2021-05-18 | Edited | Joshi-Tope G, Nickerson E, D'Eustachio P |
| 2021-05-18 | Reviewed | Mendez J, Aladjem M |
| 2021-05-22 | Modified | Shorser S |

#### Entities found in this pathway (11)

| Input | UniProt Id | Input | UniProt Id | Input | UniProt Id |
|---|---|---|---|---|---|
| CDK1 | P24941 | GINS2 | Q9Y248 | GINS3 | Q9BRX5 |
| MCM10 | Q7L590 | MCM3 | P25205 | MCM4 | P33991 |
| MCM5 | P33992 | MCM7 | P33993 | ORC6 | Q9Y5N6 |
| PCNA | P12004 | POLA2 | Q14181 | | |

#### 14. M Phase (R-HSA-68886)

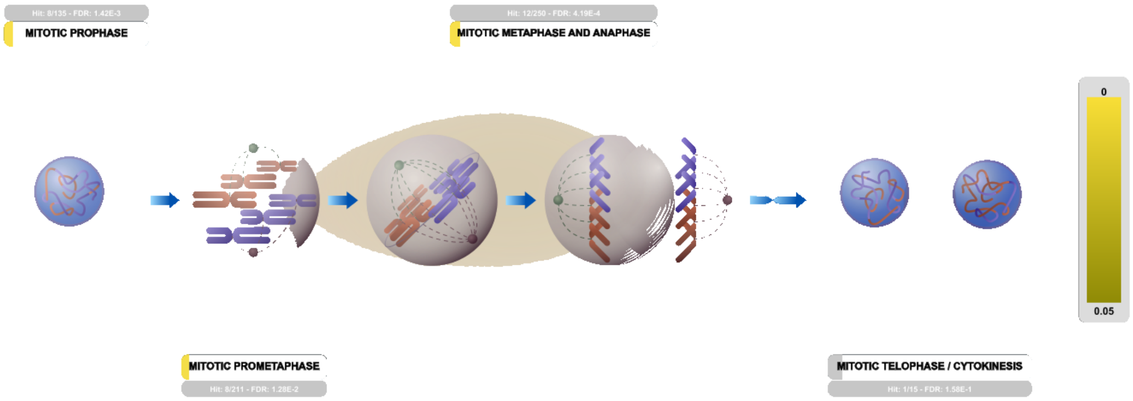


Mitosis, or the M phase, involves nuclear division and cytokinesis, where two identical daughter cells are produced. Mitosis involves prophase, prometaphase, metaphase, anaphase, and telophase. Finally, cytokinesis leads to cell division. The phase between two M phases is called the interphase; it encompasses the G1, S, and G2 phases of the cell cycle.

##### References

##### Edit history

| Date | Action | Author |
|---|---|---|
| 2018-07-10 | Reviewed | Manfredi JJ |
| 2021-05-22 | Modified | Shorser S |

##### Entities found in this pathway (16)

| Input | UniProt Id | Input | UniProt Id | Input | UniProt Id |
|---|---|---|---|---|---|
| CCNB2 | O95067 | CDK1 | P06493 | CENPF | P49454 |
| CENPU | Q71F23 | CHMP2A | O43633 | FBXO5 | Q9UKT4 |
| H2AFJ | Q9BTM1 | HIST1H2AC | Q93077 | HIST1H3F | P68431 |
| HIST1H4C | P62805 | HSP90AA1 | P07900 | LBR | Q14739 |
| LMNA | P02545-1, P02545-2 | PTTG1 | O95997 | RAD21 | O60216 |
| TUBB | P04350, P07437 | | | | |

#### 15. S Phase (R-HSA-69242)

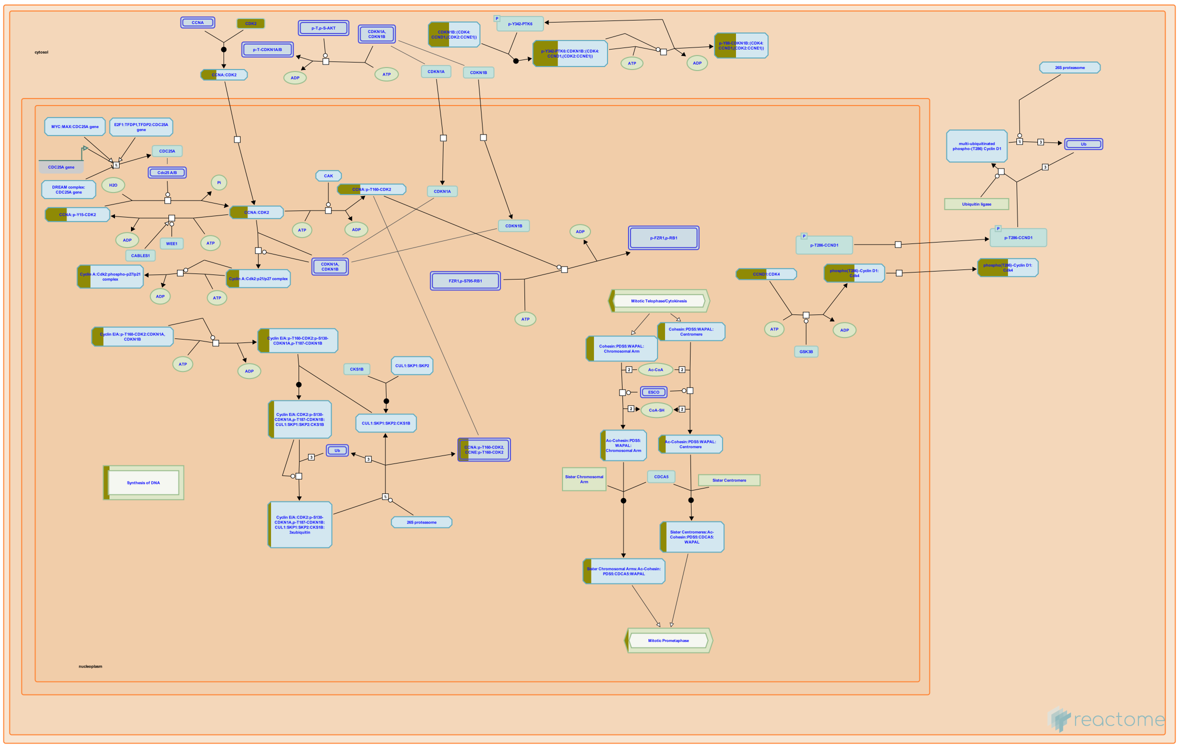

DNA synthesis occurs in the S phase, or the synthesis phase, of the cell cycle. The cell duplicates its hereditary material, and two copies of the chromosome are formed. As DNA replication continues, the E type cyclins shared by the G1 and S phases, are destroyed and the levels of the mitotic cyclins rise.

##### References

##### Edit history

| Date | Action | Author |
|---|---|---|
| 2018-07-10 | Reviewed | Manfredi JJ |
| 2021-05-22 | Modified | Shorser S |

##### Entities found in this pathway (12)

| Input | UniProt Id | Input | UniProt Id | Input | UniProt Id |
|---|---|---|---|---|---|
| CDK1 | P24941 | CDK4 | P11802 | GINS2 | Q9Y248 |
| GINS3 | Q9BRX5 | MCM3 | P25205 | MCM4 | P33991 |
| MCM5 | P33992 | MCM7 | P33993 | ORC6 | Q9Y5N6 |
| PCNA | P12004 | POLA2 | Q14181 | RAD21 | O60216 |

#### 16. Reproduction (R-HSA-1474165)

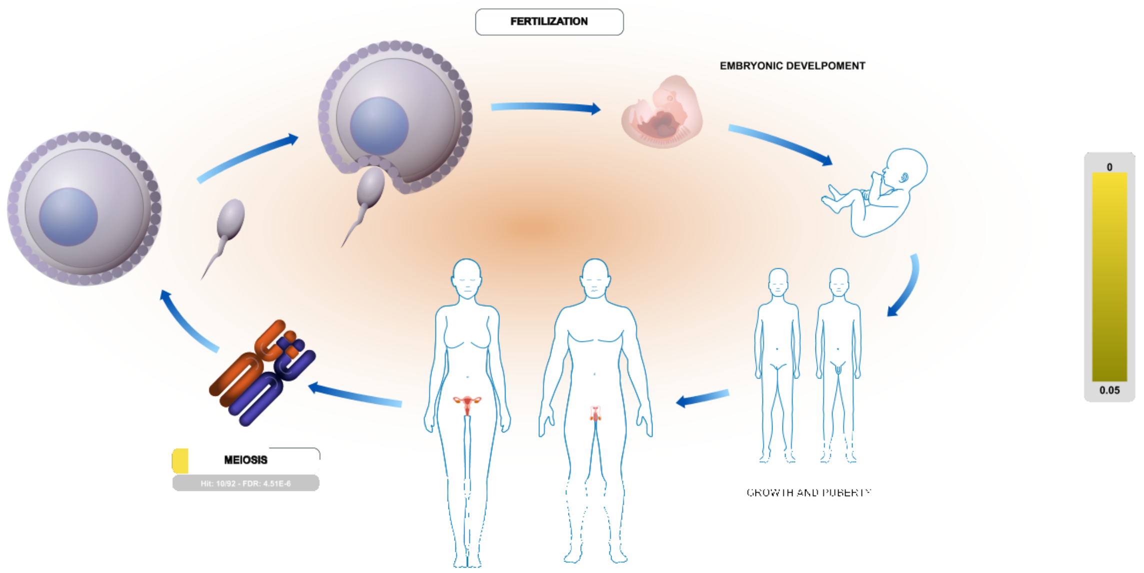


Human reproduction mixes the genomes of two individuals creating a new organism. The offspring individuals produced by sexual reproduction differ from their parents and from their siblings. Reproduction includes the reproductive system, sperm and egg production (haploid cells), fertilization, and the early stages embryo development.

##### Edit history

| Date | Action | Author |
|---|---|---|
| 2011-08-04 | Created | Gillespie ME |
| 2013-02-13 | Authored | Gillespie ME |
| 2013-05-21 | Reviewed | Lishko PV |
| 2013-05-23 | Edited | Gillespie ME |
| 2021-05-22 | Modified | Shorser S |

##### Entities found in this pathway (10)

| Input | UniProt Id | Input | UniProt Id | Input | UniProt Id |
|---|---|---|---|---|---|
| BLM | P54132 | BRCA1 | P38398 | BRCA2 | P51587 |
| CDK4 | P11802 | H2AFJ | Q9BTM1 | HIST1H2AC | Q93077 |
| HIST1H3F | P68431 | HIST1H4C | P62805 | LMNA | P02545-2 |
| RAD21 | O60216 | | | | |

#### 17. RMTs methylate histone arginines (R-HSA-3214858)

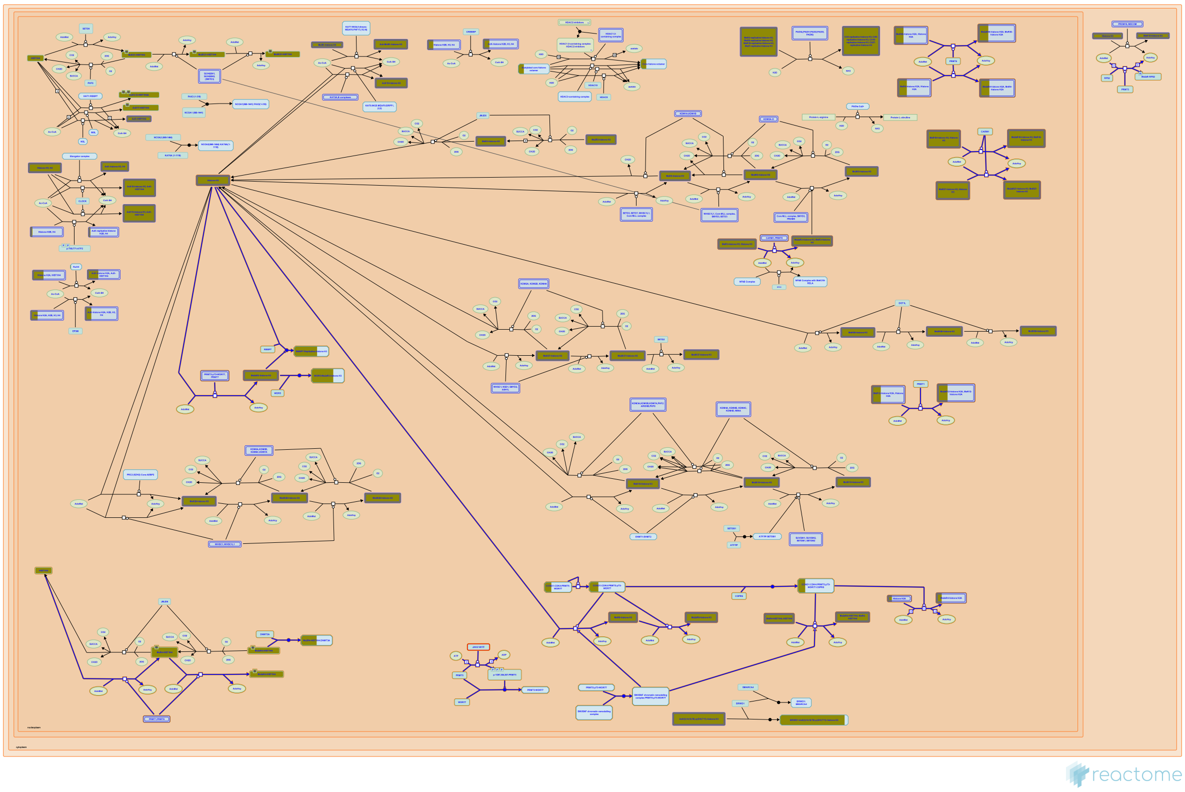

Arginine methylation is a common post-translational modification; around 2% of arginine residues are methylated in rat liver nuclei (Boffa et al. 1977). Arginine can be methylated in 3 different ways: monomethylarginine (MMA); NG,NG-asymmetric dimethylarginine (ADMA) and NG,N'G-symmetric dimethylarginine (SDMA). The formation of MMA, ADMA and SDMA in mammalian cells is carried out by members of a family of nine protein arginine methyltransferases (PRMTs) (Bedford & Clarke 2009).

Type I, II and III PRMTs generate MMA on one of the two terminal guanidino nitrogen atoms. Subsequent generation of asymmetric dimethylarginine (ADMA) is catalysed by the type I enzymes PRMT1, PRMT2, PRMT3, co-activator-associated arginine methyltransferase 1 (CARM1), PRMT6 and PRMT8. Production of symmetric dimethylarginine (SDMA) is catalysed by the type II enzymes PRMT5 and PRMT7. On certain substrates, PRMT7 also functions as a type III enzyme, generating MMA only. PRMT9 activity has not been characterized. No known enzyme is capable of both ADMA and SDMA modifications. Arginine methylation is regarded as highly stable; no arginine demethylases are known (Yang & Bedford 2013).

Most PRMTs methylate glycine- and arginine-rich (GAR) motifs in their substrates (Boffa et al. 1977). CARM1 methylates a proline-, glycine- and methionine-rich (PGM) motif (Cheng et al. 2007). PRMT5 can dimethylate arginine residues in GAR and PGM motifs (Cheng et al. 2007, Branscombe et al. 2001).

PRMTs are widely expressed and are constitutively active as purified recombinant proteins. However, PRMT activity can be regulated through PTMs, association with regulatory proteins, sub-cellular compartmentalization and factors that affect enzyme-substrate interactions. The target sites of PRMTs are influenced by the presence of other PTMs on their substrates. The best characterized examples of this are for histones. Histone H3 lysine-19 acetylation (H3K18ac) primes the histone tail for asymmetric dimethylation at arginine-18 (H3R17me2a) by CARM1 (An et al. 2003, Daujat et al. 2002, Yue et al. 2007). H3 lysine-10 acetylation (H3K9ac) blocks arginine-9 symmetric dimethylation (H3R8me2s) by PRMT5 (Pal et al. 2004). H4R3me2a catalyzed by PRMT1 favours subsequent acetylation of the histone H4 tail (Huang et al. 2005). At the same time histone H4 lysine-5 acetylation (H4K5ac) makes the H4R3 motif a better substrate for PRMT5 compared with PRMT1, thereby moving the balance from an activating ADMA mark to a suppressive SDMA mark at the H4R3 motif (Feng et al. 2011). Finally methylation of Histone H3 on arginine-3 (H3R2me2a) by PRMT6 blocks methylation of H3 lysine-5 by the MLL complex (H3K4me3), and vice versa, methylation of H3K4me3 prevents H3R2me2a methylation (Guccione et al. 2007, Kirmizis et al. 2007, Hyllus et al. 2007).

N.B. The coordinates of post-translational modifications represented and described here follow UniProt standard practice whereby coordinates refer to the translated protein before any further processing. Histone literature typically refers to coordinates of the protein after the initiating methionine has been removed. Therefore the coordinates of post-translated residues in the Reactome database and described here are frequently +1 when compared with the literature.


#### Edit history

| Date | Action | Author |
|---|---|---|
| 2013-03-12 | Authored | Jupe S |
| 2013-03-12 | Created | Jupe S |
| 2013-03-15 | Edited | Jupe S |
| 2014-05-09 | Reviewed | Guccione E |
| 2014-07-23 | Reviewed | Fischle W |
| 2021-05-22 | Modified | Shorser S |

#### Entities found in this pathway (7)

| Input | UniProt Id | Input | UniProt Id | Input | UniProt Id |
|---|---|---|---|---|---|
| CDK4 | P11802 | H2AFJ | Q9BTM1 | HIST1H2AC | Q93077 |
| HIST1H2AH | Q96KK5 | HIST1H2AL | P0C0S8 | HIST1H3F | P68431 |
| HIST1H4C | P62805 | | | | |

#### 18. Synthesis of DNA (R-HSA-69239)

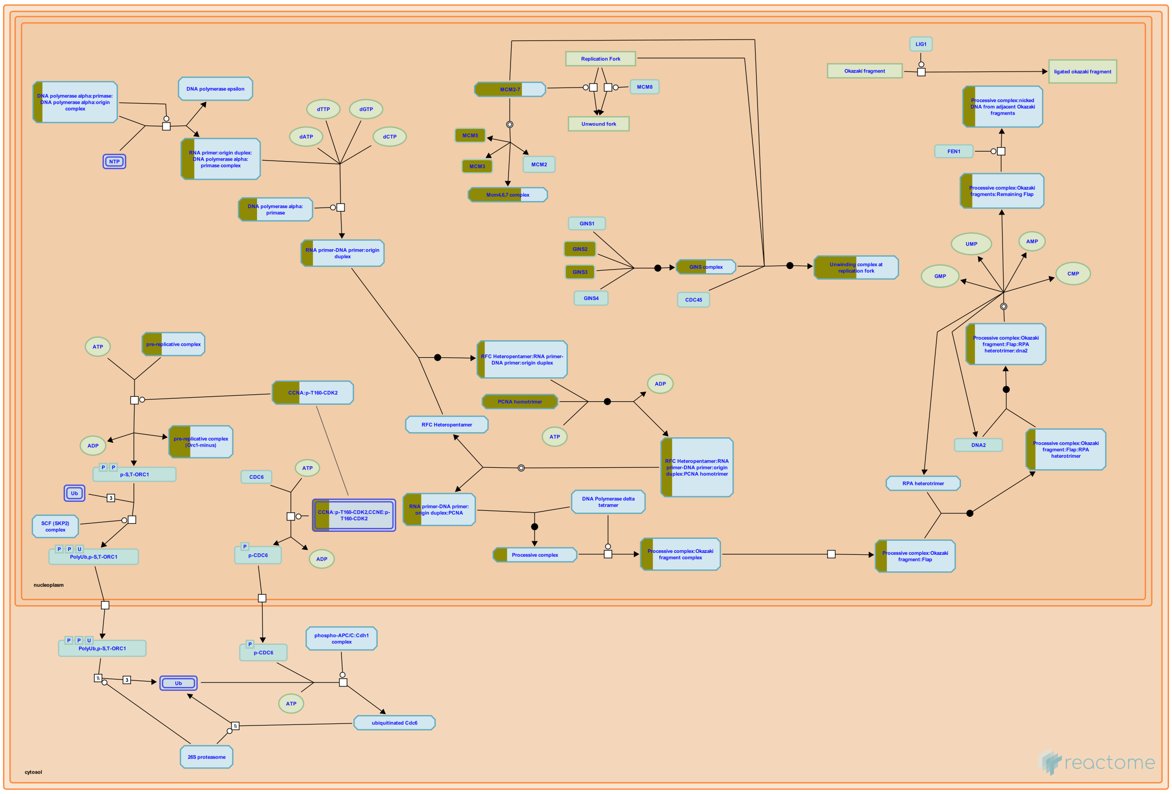

**Cellular compartments:** nucleoplasm, cytosol.

The actual synthesis of DNA occurs in the S phase of the cell cycle. This includes the initiation of DNA replication, when the first nucleotide of the new strand is laid down during the synthesis of the primer. The DNA replication preinitiation events begin in late M or early G1 phase.

##### References

##### Edit history

| Date | Action | Author |
|---|---|---|
| 2021-05-22 | Modified | Shorser S |

##### Entities found in this pathway (10)

| Input | UniProt Id | Input | UniProt Id | Input | UniProt Id |
|---|---|---|---|---|---|
| CDK1 | P24941 | GINS2 | Q9Y248 | GINS3 | Q9BRX5 |
| MCM3 | P25205 | MCM4 | P33991 | MCM5 | P33992 |
| MCM7 | P33993 | ORC6 | Q9Y5N6 | PCNA | P12004 |
| POLA2 | Q14181 | | | | |

#### 19. Diseases of DNA repair (R-HSA-9675135)

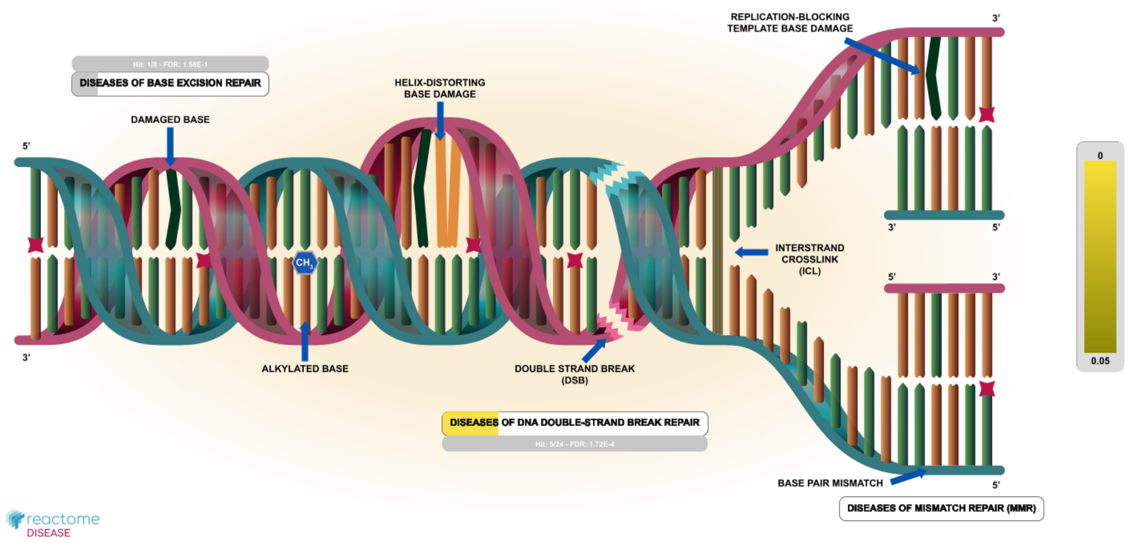


**Diseases:** genetic disease.

Germline and somatic defects in genes that encode proteins that participate in DNA repair give rise to genetic instability that can lead to malignant transformation or trigger cellular senescence or apoptosis. Germline defects in DNA repair genes are an underlying cause of familial cancer syndromes and premature ageing syndromes. Somatic defects in DNA repair genes are frequently found in tumors. For review, please refer to Tiwari and Wilson 2019.

We have so far annotated diseases of mismatch repair, diseases of base excision repair and diseases of DNA double-strand break repair.

Defects in mammalian DNA mismatch repair (MMR) genes (MLH1, PMS2, MSH2, and MSH6) result in microsatellite instability (MSI) and reduced fidelity during replication and repair steps. Defective variants of MMR genes are associated with sporadic cancers with hypermutation phenotypes as well as hereditary cancer syndromes such as Lynch syndrome (hereditary non-polyposis colorectal cancer) and constitutional mismatch repair deficiency syndrome (CMMRD). MSI is an important predictor of sensitivity to cancer immunotherapy as the high mutational burden renders MSI tumors immunogenic and sensitive to programmed cell death-1 (PD-1) immune checkpoint inhibitors (Mandal et al. 2019). For review, please refer to Pena-Diaz and Rasmussen 2016, Sijmons and Hofstra 2016, Tabori et al. 2017, Baretti and Le 2018.

Germline mutations, single nucleotide polymorphisms (SNPs) and somatic mutations in several genes involved in base excision repair (BER), a DNA repair pathway where a damaged DNA base is excised and replaced with a correct base, are involved in the development of cancer and several oxidative stress-related diseases. For review, please refer to Fu et al. 2012, Fletcher and Houlston 2010, Brenerman et al. 2014, Patrono et al. 2014, and D'Errico et al. 2017.

Germline mutations in genes involved in repair of DNA double-strand breaks (DSBs) are the underlying cause of several cancer predisposition syndromes, some of which also encompass developmental disorders associated with immune dysfunction, radiosensitivity and neurodegeneration. Somatic mutations in genes involved in DSB repair also occur in sporadic cancers. For review, please refer to McKinnon and Caldecott 2007, Keijzers et al. 2017, and Jachimowicz et al. 2019.


#### Edit history

| Date | Action | Author |
|---|---|---|
| 2020-01-31 | Created | Orlic-Milacic M |
| 2020-02-21 | Authored | Orlic-Milacic M |
| 2020-02-24 | Edited | Orlic-Milacic M |
| 2020-02-24 | Reviewed | D'Eustachio P |
| 2020-11-11 | Reviewed | D'Eustachio P |
| 2020-11-12 | Modified | Orlic-Milacic M |
| 2020-11-12 | Edited | Orlic-Milacic M |

#### Entities found in this pathway (6)

| Input | UniProt Id | Input | UniProt Id | Input | UniProt Id |
|---|---|---|---|---|---|
| BLM | P54132 | BRCA1 | P38398 | BRCA2 | P51587 |
| BRIP1 | Q9BX63 | EXO1 | Q9UQ84 | NEIL3 | Q8TAT5 |

#### 20. Activation of ATR in response to replication stress (R-HSA-176187)

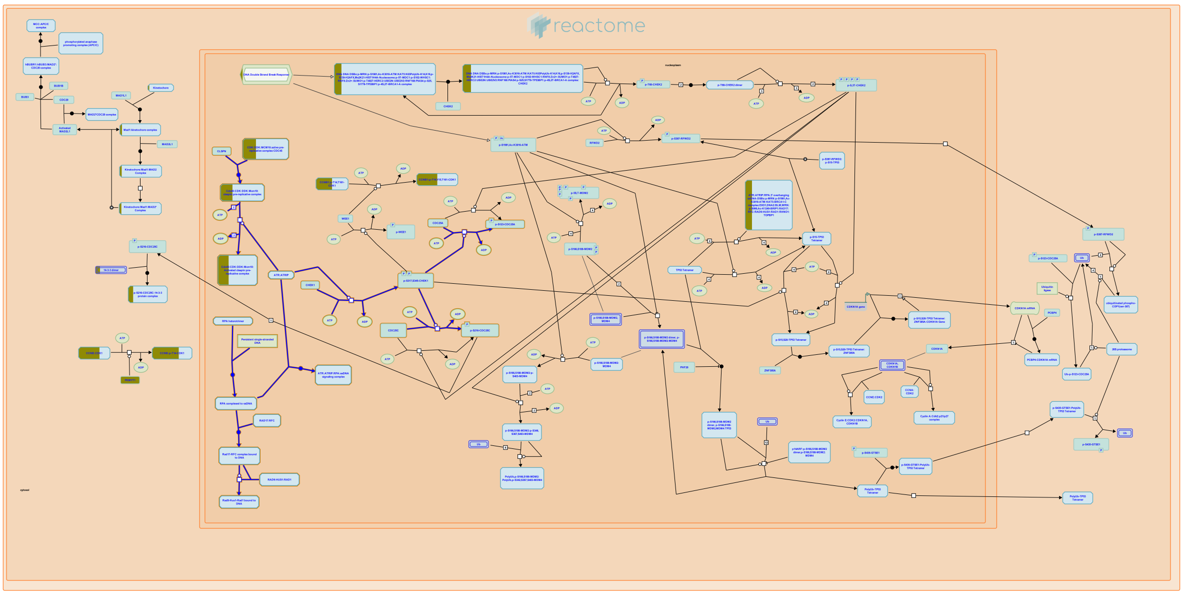

**Cellular compartments:** nucleoplasm.

Genotoxic stress caused by DNA damage or stalled replication forks can lead to genomic instability. To guard against such instability, genotoxically-stressed cells activate checkpoint factors that halt or slow cell cycle progression. Among the pathways affected are DNA replication by reduction of replication origin firing, and mitosis by inhibiting activation of cyclin-dependent kinases (Cdks). A key factor involved in the response to stalled replication forks is the ATM- and rad3-related (ATR) kinase, a member of the phosphoinositide-3-kinase-related kinase (PIKK) family. Rather than responding to particular lesions in DNA, ATR and its binding partner ATRIP (ATR-interacting protein) sense replication fork stalling indirectly by associating with persistent ssDNA bound by RPA. These structures would be formed, for example, by dissociation of the replicative helicase from the leading or lagging strand DNA polymerase when the polymerase encounters a DNA lesion that blocks DNA synthesis. Along with phosphorylating the downstream transducer kinase Chk1 and the tumor suppressor p53, activated ATR modifies numerous factors that regulate cell cycle progression or the repair of DNA damage. The persistent ssDNA also stimulates recruitment of the RFC-like Rad17-Rfc2-5 alternative clamp-loading complex, which subsequently loads the Rad9-Hus1-Rad1 complex onto the DNA. The latter '9-1-1' complex serves to facilitate Chk1 binding to the stalled replication fork, where Chk1 is phosphorylated by ATR and thereby activated. Upon activation, Chk1 can phosphorylate additional substrates including the Cdc25 family of phosphatases (Cdc25A, Cdc25B, and Cdc25C). These enzymes catalyze the removal of inhibitory phosphate residues from cyclin-dependent kinases (Cdks), allowing their activation. In particular, Cdc25A primarily functions at the G1/S transition to dephosphorylate Cdk2 at Thr 14 and Tyr 15, thus positively regulating the Cdk2-cyclin E complex for S-phase entry. Cdc25A also has mitotic functions. Phosphorylation of Cdc25A at Ser125 by Chk1 leads to Cdc25A ubiquitination and degradation, thus inhibiting DNA replication origin firing. In contrast, Cdc25B and Cdc25C regulate the onset of mitosis through dephosphorylation and activation of Cdk1-cyclin B complexes. In response to replication stress, Chk1 phosphorylates Cdc25B and Cdc25C leading to Cdc25B/C complex formation with 14-3-3 proteins. As these complexes are sequestered in the cytoplasm, they are unable to activate the nuclear Cdk1-cyclin B complex for mitotic entry.

These events are outlined in the figure. Persistent single-stranded DNA associated with RPA binds claspin (A) and ATR:ATRIP (B), leading to claspin phosphorylation (C). In parallel, the same single-stranded DNA:RPA complex binds RAD17:RFC (D), enabling the loading of RAD9:HUS1:RAD1 (9-1-1) complex onto the DNA (E). The resulting complex of proteins can then repeatedly bind (F) and phosphorylate (G) CHK1, activating multiple copies of CHK1.


#### Edit history

| Date | Action | Author |
|---|---|---|
| 2006-02-25 | Edited | D'Eustachio P |
| 2006-02-25 | Authored | Borowiec JA |
| 2006-03-03 | Created | D'Eustachio P |
| 2021-05-22 | Modified | Shorser S |

#### Entities found in this pathway (6)

| Input | UniProt Id | Input | UniProt Id | Input | UniProt Id |
|---|---|---|---|---|---|
| MCM10 | Q7L590 | MCM3 | P25205 | MCM4 | P33991 |
| MCM5 | P33992 | MCM7 | P33993 | ORC6 | Q9Y5N6 |

#### 21. Polo-like kinase mediated events (R-HSA-156711)

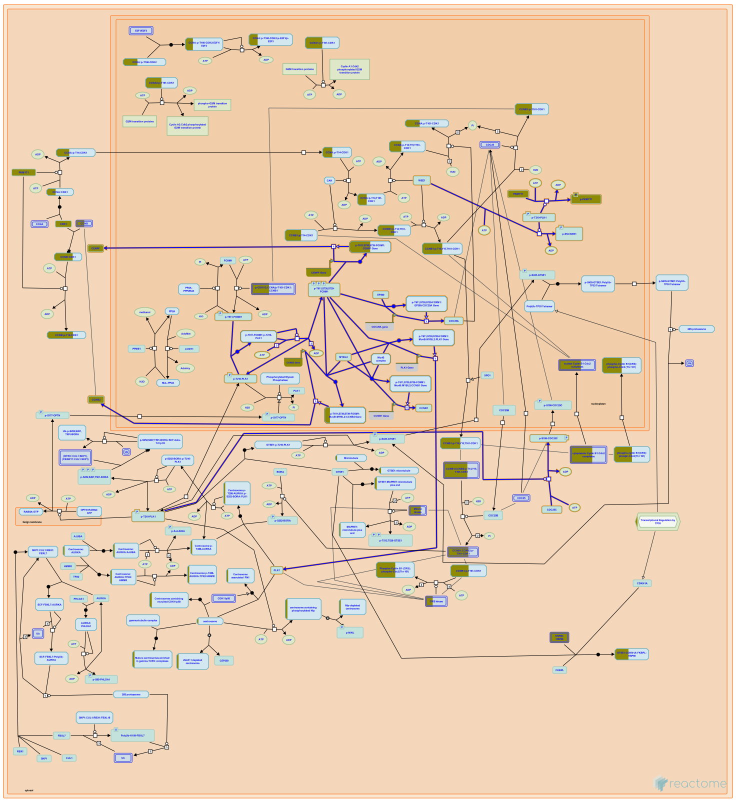

**Cellular compartments:** nucleoplasm.

At mitotic entry, Plk1 phosphorylates and activates Cdc25C phosphatase, whereas it phosphorylates and down-regulates Wee1A (Watanabe et al. 2004). Plk1 also phosphorylates and inhibits Myt1 activity (Sagata 2005). Cyclin B1-bound Cdc2, which is the target of Cdc25C, Wee1A, and Myt1, functions in a feedback loop and phosphorylates the latter components (Cdc25C, Wee1A, Myt1). The Cdc2- dependent phosphorylation provides docking sites for the polo-box domain of Plk1, thus promoting the Plk1-dependent regulation of these components and, as a result, activation of Cdc2-Cyclin B1.

PLK1 phosphorylates and activates the transcription factor FOXM1 which stimulates the expression of a number of genes needed for G2/M transition, including PLK1, thereby creating a positive feedback loop (Laoukili et al. 2005, Fu et al. 2008, Sadasivam et al. 2012, Chen et al. 2013).

#### Edit history

| Date | Action | Author |
|---|---|---|
| 2004-12-09 | Authored | Lee KS |
| 2004-12-09 | Created | Gillespie ME |
| 2013-08-21 | Reviewed | Bruinsma W |
| 2021-05-18 | Edited | Gillespie ME |
| 2021-05-22 | Modified | Shorser S |

#### Entities found in this pathway (3)

| Input | UniProt Id | Input | UniProt Id | Input | UniProt Id |
|---|---|---|---|---|---|
| CCNB2 | O95067 | CENPF | P49454 | PKMYT1 | Q99640 |

| Input | Ensembl Id | Input | Ensembl Id |
|---|---|---|---|
| CCNB2 | ENSG00000157456 | CENPF | ENSG00000117724 |

#### 22. Diseases of DNA Double-Strand Break Repair (R-HSA-9675136)

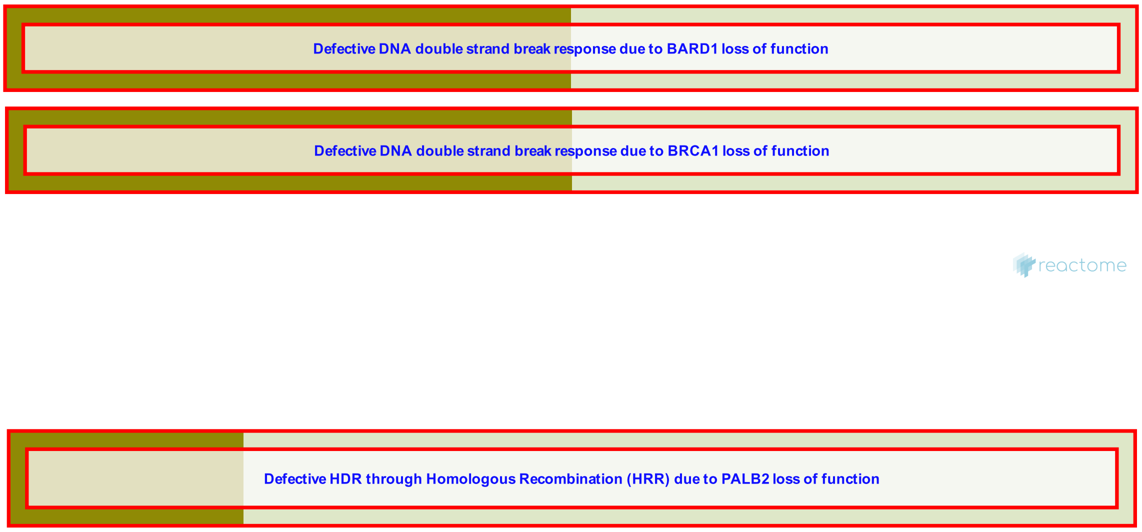


**Diseases:** cancer.

Diseases of DNA double-strand break repair (DSBR) are caused by mutations in genes involved in repair of double strand breaks (DSBs), one of the most cytotoxic types of DNA damage. Unrepaired DSBs can lead to cell death, cellular senescence, or malignant transformation.

Germline mutations in DSBR genes are responsible for several developmental disorders associated with increased predisposition to cancer:

Ataxia telangiectasia, characterized by cerebellar neurodegeneration, hematologic malignancies and immunodeficiency, is usually caused by germline mutations in the ATM gene;

Nijmegen breakage syndrome 1, characterized by microcephaly, short stature and recurrent infections, is caused by germline mutations in the NBN (NBS1) gene;

Seckel syndrome, characterized by short stature, skeletal deformities and microcephaly, is caused by germline mutations in the ATR or RBBP8 (CtIP) genes.

Heterozygous germline mutations in BRCA1, BRCA2 or PALB2 cause the hereditary breast and ovarian cancer syndrome (HBOC), while homozygous germline mutations in BRCA2 and PALB2 cause Fanconi anemia, a developmental disorder characterized by short stature, microcephaly, skeletal defects, bone marrow failure, and predisposition to cancer.

Somatic mutations in DSBR genes are also frequently found in sporadic cancers.

We have so far annotated defects in DSB response caused by loss-of-function mutations in BRCA1 and its heterodimerization partner BARD1, which prevent the formation of the BRCA1:BARD1 complex.

For review, please refer to McKinnon and Caldecott 2007, Keijzers et al. 2017, and Jachimowicz et al. 2019.

#### Edit history

| Date | Action | Author |
|---|---|---|
| 2020-01-31 | Created | Orlic-Milacic M |
| 2020-11-11 | Reviewed | D'Eustachio P |
| 2020-11-11 | Authored | Orlic-Milacic M |
| 2020-11-12 | Modified | Orlic-Milacic M |
| 2020-11-12 | Edited | Orlic-Milacic M |

#### Entities found in this pathway (5)

| Input | UniProt Id | Input | UniProt Id | Input | UniProt Id |
|---|---|---|---|---|---|
| BLM | P54132 | BRCA1 | P38398 | BRCA2 | P51587 |
| BRIP1 | Q9BX63 | EXO1 | Q9UQ84 | | |

#### 23. Defective HDR through Homologous Recombination (HRR) due to PALB2 loss of function (R-HSA-9701193)

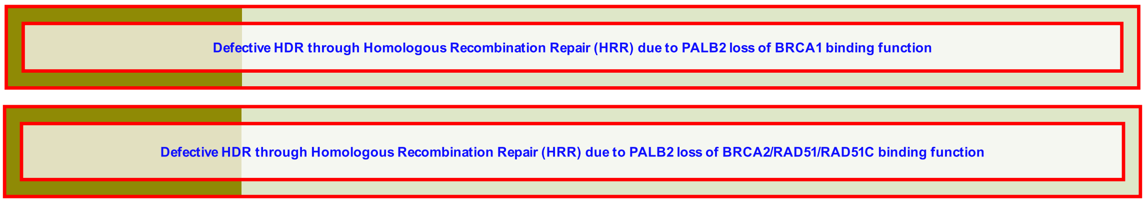


**Cellular compartments:** nucleoplasm.

**Diseases:** cancer.

Biallelic loss-of-function mutations in *PALB2* results in Fanconi anemia subtype N (FA-N), which is phenotypically very similar to Fanconi anemia subtype D1, caused by biallelic loss-of-function of *BRCA2* (Reid et al. 2007). FA-D1 and FA-N are characterized by developmental abnormalities, bone marrow failure and childhood cancer susceptibility, especially childhood solid tumors, such as Wilms tumor and medulloblastoma. Monoallelic *PALB2* loss-of-function is an underlying cause of hereditary breast cancer in particular, but inactivating *PALB2* mutations are also to a lesser extent found in some other cancer types (Erkko et al. 2007, Erkko et al. 2008, Antoniou et al. 2014, Yang et al. 2020). Germline *PALB2* mutations are somewhat less frequent than those occurring in *BRCA1* and *BRCA2*, but cause a comparably high risk of developing breast cancer.

PALB2 interacts with both BRCA1 and BRCA2, and serves as a bridge that connects BRCA2 with BRCA1 at sites of DNA double-strand break repair (DSBR). PALB2 loss-of-function mutations can affect its interaction with BRCA1 when they affect the N-terminal coiled-coil domain that is necessary for BRCA1 binding (Sy et al. 2009, Foo et al. 2017). PALB2 missense mutants that do not bind to BRCA1 can still be recruited to DSBR sites, probably through interaction with other proteins involved in DSBR, but they are unable to restore efficient gene conversion in PALB2-deficient cells and they render cells hypersensitive to the DNA damaging agent mitomycin C (Sy et al. 2009).

Mutations affecting the C-terminal WD40 domain of PALB2 impair its ability to interact with BRCA2, RAD51 and/or RAD51C (Erkko et al. 2007, Park et al. 2014, Simhadri et al. 2019). Mutations affecting the C-terminal domain of PALB2 are more frequent than mutations that affect the N-terminus and have been observed, as germline mutations, in familial breast cancer and in Fanconi anemia, but somatic mutations also occur in sporadic cancers. Cells that express PALB2 mutants defective in BRCA2, RAD51 and/or RAD51C binding show reduced ability to perform DSBR via homologous recombination repair, form fewer RAD51 foci at DSBR sites, and are sensitive to DNA crosslinking agents such as mitomycin C (Erkko et al. 2007, Parker et al. 2014).

For review, please refer to Tischkowitz and Xia 2010, Pauty et al. 2014, Park et al. 2014, Nepomuceno et al. 2017, Ducy et al. 2019, and Wu et al. 2020.

#### Edit history

| Date | Action | Author |
|---|---|---|
| 2020-09-24 | Created | Orlic-Milacic M |
| 2020-10-09 | Authored | Orlic-Milacic M |
| 2021-02-25 | Reviewed | Pospiech H, Winqvist R |
| 2021-03-25 | Edited | Orlic-Milacic M |
| 2021-05-19 | Modified | Shorser S |

#### Entities found in this pathway (5)

| Input | UniProt Id | Input | UniProt Id | Input | UniProt Id |
|---|---|---|---|---|---|
| BLM | P54132 | BRCA1 | P38398 | BRCA2 | P51587 |
| BRIP1 | Q9BX63 | EXO1 | Q9UQ84 | | |

#### 24. Defective HDR through Homologous Recombination Repair (HRR) due to PALB2 loss of BRCA2/RAD51/RAD51C binding function (R-HSA-9704646)

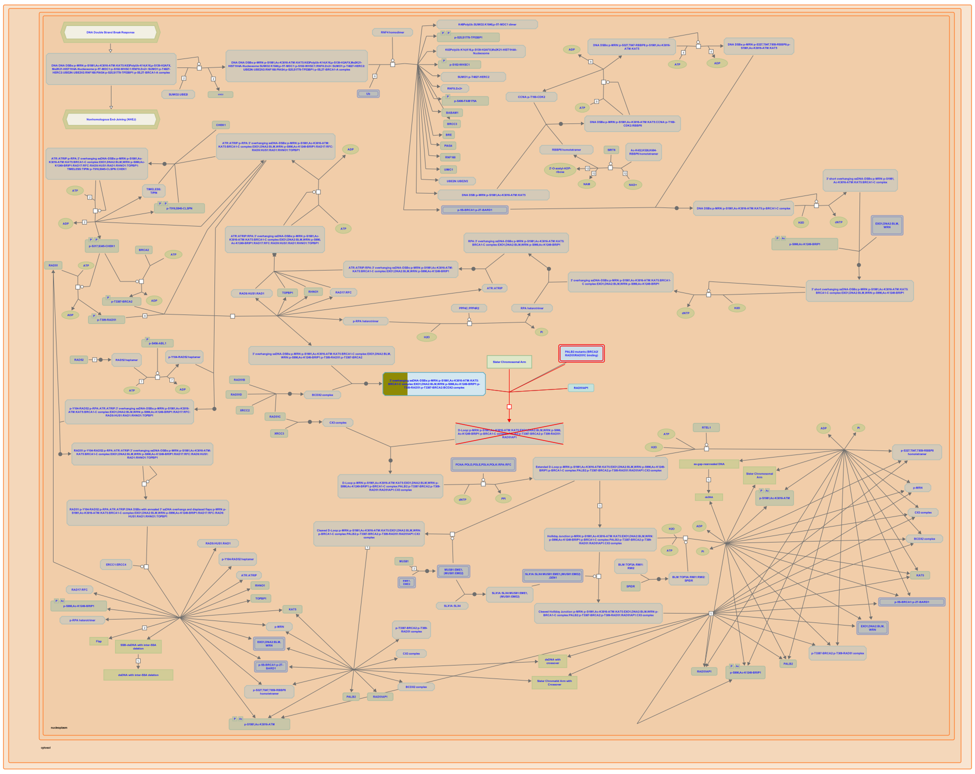

**Cellular compartments:** nucleoplasm.

**Diseases:** cancer.

Mutations affecting the C-terminal WD40 domain of PALB2 impair its ability to interact with BRCA2, RAD51 and/or RAD51C (Erkko et al. 2007, Park et al. 2014). Mutations affecting the C-terminal domain of PALB2 are more frequent than mutations that affect the N-terminus and have been observed, as germline mutations, in familial breast cancer and in Fanconi anemia, but somatic mutations also occur in sporadic cancers. Cells that express PALB2 mutants defective in BRCA2, RAD51 and/or RAD51C binding show reduced ability to perform DSBR via homologous recombination repair, form fewer RAD51 foci at DSBR sites, and are sensitive to DNA crosslinking agents such as mitomycin C (Erkko et al. 2007, Park et al. 2014).

##### Edit history

| Date | Action | Author |
| --- | --- | --- |
| 2020-10-09 | Authored | Orlic-Milacic M |
| 2020-10-09 | Created | Orlic-Milacic M |

| Date | Action | Author |
|---|---|---|
| 2021-02-25 | Reviewed | Pospiech H, Winqvist R |
| 2021-03-25 | Modified | Orlic-Milacic M |
| 2021-03-25 | Edited | Orlic-Milacic M |

#### Entities found in this pathway (5)

| Input | UniProt Id | Input | UniProt Id | Input | UniProt Id |
|---|---|---|---|---|---|
| BLM | P54132 | BRCA1 | P38398 | BRCA2 | P51587 |
| BRIP1 | Q9BX63 | EXO1 | Q9UQ84 | | |

#### 25. Defective HDR through Homologous Recombination Repair (HRR) due to PALB2 loss of BRCA1 binding function (R-HSA-9704331)

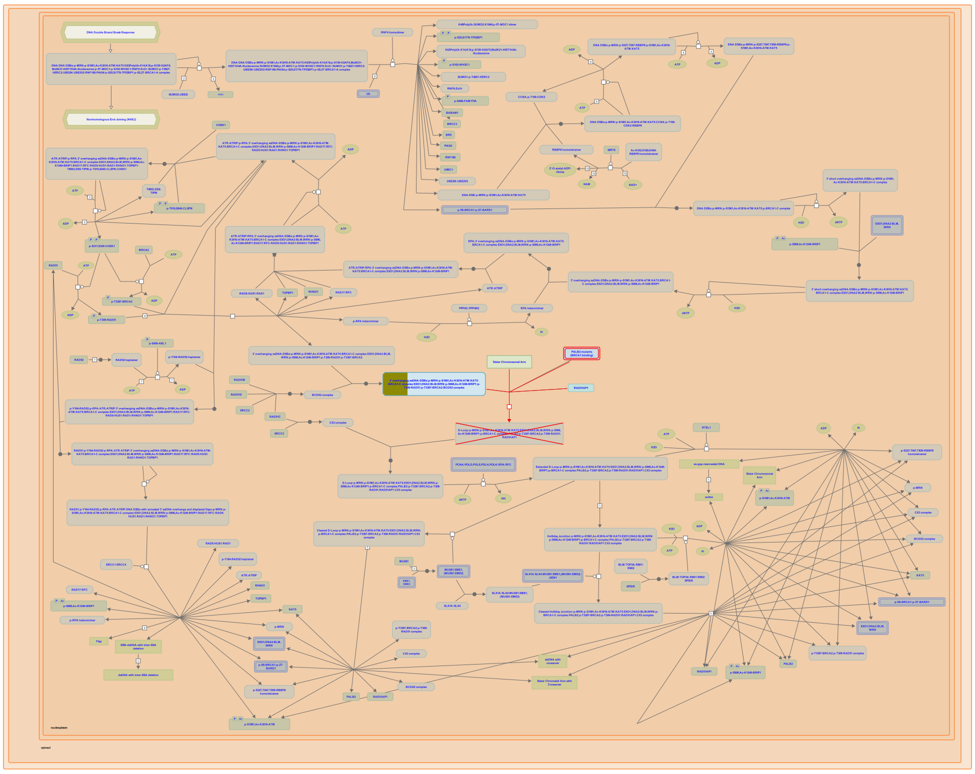

**Cellular compartments:** nucleoplasm.

**Diseases:** cancer.

Mutations in the N-terminal coiled-coil domain of PALB2, involved in self-interaction and BRCA1 binding, impair the interaction of PALB2 with BRCA1 (Sy et al. 2009, Foo et al. 2017, Boonen et al. 2020). PALB2 missense mutants that do not bind to BRCA1 can still be recruited to DNA double-strand break repair (DSBR) sites, probably through interaction with other proteins involved in DS-BR, but they are unable to restore efficient gene conversion in PALB2-deficient cells and they render cells hypersensitive to the DNA damaging agent mitomycin C (Sy et al. 2009).


##### Edit history

| Date | Action | Author |
|---|---|---|
| 2020-10-07 | Created | Orlic-Milacic M |

| Date | Action | Author |
|---|---|---|
| 2020-10-09 | Authored | Orlic-Milacic M |
| 2021-02-25 | Reviewed | Pospiech H, Winqvist R |
| 2021-03-25 | Modified | Orlic-Milacic M |
| 2021-03-25 | Edited | Orlic-Milacic M |

#### Entities found in this pathway (5)

| Input | UniProt Id | Input | UniProt Id | Input | UniProt Id |
|---|---|---|---|---|---|
| BLM | P54132 | BRCA1 | P38398 | BRCA2 | P51587 |
| BRIP1 | Q9BX63 | EXO1 | Q9UQ84 | | |

### 6. Identifiers found

Below is a list of the input identifiers that have been found or mapped to an equivalent element in Reactome, classified by resource.

#### Entities (86)

| Input | UniProt Id | Input | UniProt Id | Input | UniProt Id |
|---|---|---|---|---|---|
| ACTB | P60709 | ACTG1 | P60709 | ANXA2 | P07355 |
| ATAD2 | Q6PL18 | BLM | P54132 | BRCA1 | P38398 |
| BRCA2 | P51587 | BRIP1 | Q9BX63 | CALM3 | P0DP23 |
| CALR | P27797 | CCDC14 | O60662 | CCNB2 | O95067 |
| CDK1 | P06493, P24941 | CDK4 | P11802 | CDKN2C | P42773 |
| CENPF | P49454 | CENPU | Q71F23 | CENPX | A8MT69 |
| CHMP2A | O43633 | CTNNB1 | P35222 | CTSS | P25774 |
| DHFR | P00374 | DNMT1 | P26358 | DSP | P15924 |
| DTL | Q9NZJ0 | DUT | Q6SW70 | E2F8 | A0AVK6 |
| EPCAM | P16422 | ERN1 | O75460 | EXO1 | Q9UQ84 |
| FABP1 | P07148 | FABP6 | P51161 | FBXO5 | Q9UKT4 |
| GINS2 | Q9Y248 | GINS3 | Q9BRX5 | H2AFJ | Q9BTM1 |
| HIST1H2AC | Q93077 | HIST1H2AH | Q96KK5 | HIST1H2AL | P0C0S8 |
| HIST1H3F | P68431 | HIST1H4C | P62805 | HMGCS1 | Q01581 |
| HNRNPM | P52272 | HSP90AA1 | P07900 | HSP90AB1 | P08238 |
| HSP90B1 | P14625 | KRT18 | P05783 | KRT19 | P08727 |
| KRT20 | P35900 | KRT8 | P05787 | KTN1 | Q86UP2 |
| LBR | Q14739 | LDHA | P00338 | LGALS3 | P17931 |
| LGR4 | Q9BXB1 | LMNA | P02545-1, P02545-2 | MCM10 | Q7L590 |
| MCM3 | P25205 | MCM4 | P33991 | MCM5 | P33992 |
| MCM7 | P33993 | MVD | P53602 | MYDGF | Q969H8 |
| NEIL3 | Q8TAT5 | OAT | P04181 | ORC6 | Q9Y5N6 |
| PCLAF | Q15004 | PCNA | P12004 | PKMYT1 | Q99640 |
| POLA2 | Q14181 | PSIP1 | O75475 | PTTG1 | O95997 |
| RAD21 | O60216 | RPL36AL | P83881, Q969Q0 | RRM1 | P23921 |
| RRM2 | P31350 | S100A11 | P31949 | SEC61G | P60059 |
| SLC25A44 | Q96H78 | TFRC | P02786 | TMSB4X | P62328 |
| TOP1 | P11387 | TUBB | P04350, P07437 | TYMS | P04818 |
| UGP2 | Q16851 | YWHAZ | P63104 | | |

| Input | Ensembl Id | Input | Ensembl Id | Input | Ensembl Id |
|---|---|---|---|---|---|
| ACTB | ENST00000331789 | ANXA2 | ENSG00000182718 | BRCA1 | ENSG00000012048 |
| CALR | ENSG00000179218 | CCNB2 | ENSG00000157456 | CDK1 | ENSG00000170312 |
| CENPF | ENSG00000117724 | DHFR | ENSG00000228716 | FABP1 | ENSG00000163586 |
| FABP6 | ENSG00000170231 | FBXO5 | ENSG00000112029 | HMGCS1 | ENSG00000112972 |
| HSP90AA1 | ENSG00000080824 | HSP90B1 | ENSG00000166598 | LGALS3 | ENSG00000131981 |
| LMNA | ENSG00000160789 | MVD | ENSG00000167508 | MYDGF | ENSG00000074842 |
| PCNA | ENSG00000132646 | RRM2 | ENSG00000171848 | TFRC | ENST00000360110 |
| TYMS | ENSG00000176890 | | | | |

### 7. Identifiers not found

These 36 identifiers were not found neither mapped to any entity in Reactome.

| | | | | | | | |
|---|---|---|---|---|---|---|---|
| AC010997.5 | AL353708.3 | ATP5IF1 | CARHSP1 | CCDC15 | CDCA7 | CIB1 | DANT2 |
| FAM111B | FIGNL1 | HELLS | HEXIM1 | HINT3 | ILF3-DT | INAVA | KIAA1324 |
| KLF6 | KMT2E-AS1 | LINC02453 | LPP | LRRC23 | MALAT1 | MIR194-2HG | MTDH |
| NAP1L4 | NASP | NUP62CL | OSTC | PTMS | RASSF9 | S100A14 | S100A6 |
| SCAND1 | TCF19 | WDR76 | ZGRF1 | | | | |
