## Supplementary material for "A proximal-to-distal survey of healthy adult human small intestine and colon epithelium by single-cell transcriptomics": File S3

reactome

### Pathway Analysis Report

This report contains the pathway analysis results for the submitted sample ''. Analysis was performed against Reactome version 77 on 24/09/2021. The web link to these results is:

https://reactome.org/PathwayBrowser/#/ANALYSIS=MjAyMTA5MjQxMzE0MzdfMzA3Ng%3D%3D

Please keep in mind that analysis results are temporarily stored on our server. The storage period depends on usage of the service but is at least 7 days. As a result, please note that this URL is only valid for a limited time period and it might have expired.

### Table of Contents


1. Introduction
2. Properties
3. Genome-wide overview
4. Most significant pathways
5. Pathways details
6. Identifiers found
7. Identifiers not found

### 1. Introduction

Reactome is a curated database of pathways and reactions in human biology. Reactions can be considered as pathway 'steps'. Reactome defines a 'reaction' as any event in biology that changes the state of a biological molecule. Binding, activation, translocation, degradation and classical biochemical events involving a catalyst are all reactions. Information in the database is authored by expert biologists, entered and maintained by Reactome’s team of curators and editorial staff. Reactome content frequently cross-references other resources e.g. NCBI, Ensembl, UniProt, KEGG (Gene and Compound), ChEBI, PubMed and GO. Orthologous reactions inferred from annotation for Homo sapiens are available for 17 non-human species including mouse, rat, chicken, puffer fish, worm, fly, yeast, rice, and Arabidopsis. Pathways are represented by simple diagrams following an SBGN-like format.

Reactome's annotated data describe reactions possible if all annotated proteins and small molecules were present and active simultaneously in a cell. By overlaying an experimental dataset on these annotations, a user can perform a pathway over-representation analysis. By overlaying quantitative expression data or time series, a user can visualize the extent of change in affected pathways and its progression. A binomial test is used to calculate the probability shown for each result, and the p-values are corrected for the multiple testing (Benjamini–Hochberg procedure) that arises from evaluating the submitted list of identifiers against every pathway.

To learn more about our Pathway Analysis, please have a look at our relevant publications:

Fabregat A, Sidiropoulos K, Garapati P, Gillespie M, Hausmann K, Haw R, … D’Eustachio P (2016). The reactome pathway knowledgebase. Nucleic Acids Research, 44(D1), D481–D487. https://doi.org/10.1093/nar/gkv1351.

Fabregat A, Sidiropoulos K, Viteri G, Forner O, Marin-Garcia P, Arnau V, … Hermjakob H (2017). Reactome pathway analysis: a high-performance in-memory approach. BMC Bioinformatics, 18.

### 2. Properties

- This is an **overrepresentation** analysis: A statistical (hypergeometric distribution) test that determines whether certain Reactome pathways are over-represented (enriched) in the submitted data. It answers the question ‘Does my list contain more proteins for pathway X than would be expected by chance?’ This test produces a probability score, which is corrected for false discovery rate using the Benjamani-Hochberg method.
- 102 out of 145 identifiers in the sample were found in Reactome, where 541 pathways were hit by at least one of them.
- All non-human identifiers have been converted to their human equivalent.
- This report is filtered to show only results for species 'Homo sapiens' and resource 'all resources'.
- The unique ID for this analysis (token) is MjAyMTA5MjQxMzE0MzdfMzA3Ng%3D%3D. This ID is valid for at least 7 days in Reactome’s server. Use it to access Reactome services with your data.

### 3. Genome-wide overview

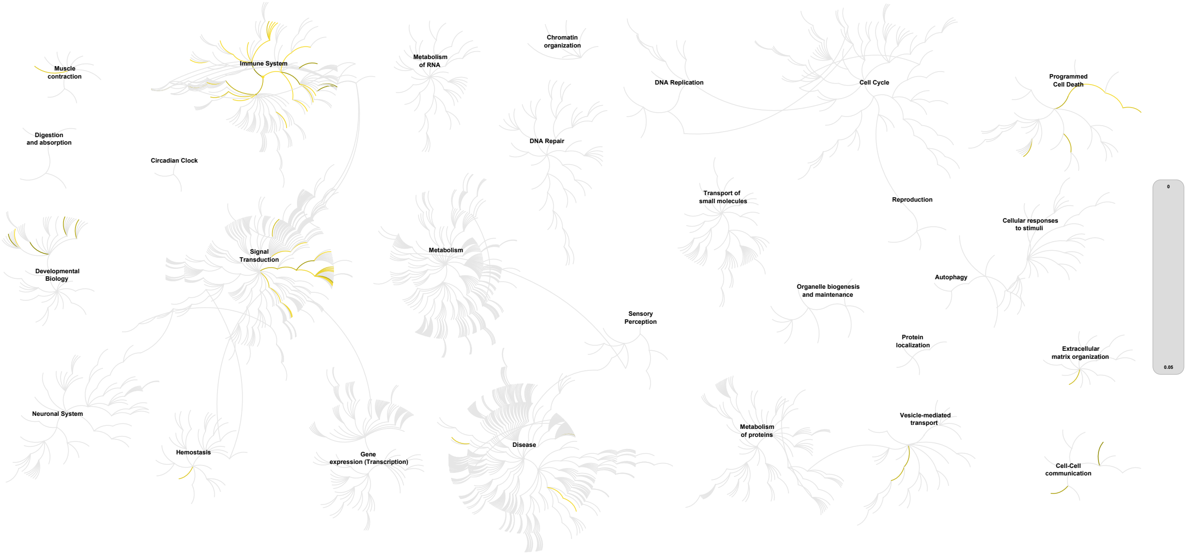


This figure shows a genome-wide overview of the results of your pathway analysis. Reactome pathways are arranged in a hierarchy. The center of each of the circular "bursts" is the root of one top-level pathway, for example "DNA Repair". Each step away from the center represents the next level lower in the pathway hierarchy. The color code denotes over-representation of that pathway in your input dataset. Light grey signifies pathways which are not significantly over-represented.

### 4. Most significant pathways

The following table shows the 25 most relevant pathways sorted by p-value.

| Pathway name | Entities | | | | Reactions | |
|---|---|---|---|---|---|---|
| | found | ratio | p-value | FDR* | found | ratio |
| Generation of second messenger molecules | 7 / 59 | 0.004 | 7.30e-06 | 0.003 | 6 / 17 | 0.001 |
| Cytokine Signaling in Immune system | 30 / 1,092 | 0.075 | 1.05e-05 | 0.003 | 137 / 708 | 0.052 |
| Interferon gamma signaling | 12 / 250 | 0.017 | 4.45e-05 | 0.008 | 4 / 16 | 0.001 |
| TNF receptor superfamily (TNFSF) members mediating non-canonical NF-kB pathway | 4 / 17 | 0.001 | 5.37e-05 | 0.008 | 9 / 12 | 8.88e-04 |
| Translocation of ZAP-70 to Immunological synapse | 5 / 42 | 0.003 | 1.51e-04 | 0.014 | 4 / 4 | 2.96e-04 |
| Immune System | 51 / 2,681 | 0.184 | 1.76e-04 | 0.014 | 252 / 1,623 | 0.12 |
| Phosphorylation of CD3 and TCR zeta chains | 5 / 45 | 0.003 | 2.07e-04 | 0.014 | 5 / 7 | 5.18e-04 |
| PD-1 signaling | 5 / 45 | 0.003 | 2.07e-04 | 0.014 | 1 / 5 | 3.70e-04 |
| Interferon Signaling | 14 / 394 | 0.027 | 2.46e-04 | 0.014 | 7 / 69 | 0.005 |
| TNFR2 non-canonical NF-kB pathway | 7 / 104 | 0.007 | 2.49e-04 | 0.014 | 22 / 43 | 0.003 |
| Cross-presentation of particulate exogenous antigens (phagosomes) | 3 / 13 | 8.94e-04 | 5.14e-04 | 0.026 | 3 / 3 | 2.22e-04 |
| Activation of RAC1 | 3 / 15 | 0.001 | 7.76e-04 | 0.036 | 4 / 4 | 2.96e-04 |
| Interleukin-4 and Interleukin-13 signaling | 9 / 211 | 0.015 | 9.28e-04 | 0.039 | 18 / 47 | 0.003 |
| Costimulation by the CD28 family | 6 / 97 | 0.007 | 0.001 | 0.039 | 2 / 35 | 0.003 |
| RHO GTPases Activate NADPH Oxidases | 4 / 38 | 0.003 | 0.001 | 0.039 | 13 / 14 | 0.001 |
| Regulation of necroptotic cell death | 4 / 38 | 0.003 | 0.001 | 0.039 | 3 / 18 | 0.001 |
| RHOV GTPase cycle | 4 / 39 | 0.003 | 0.001 | 0.04 | 1 / 2 | 1.48e-04 |
| Regulation of TNFR1 signaling | 4 / 41 | 0.003 | 0.001 | 0.044 | 7 / 17 | 0.001 |
| Neutrophil degranulation | 14 / 480 | 0.033 | 0.002 | 0.044 | 9 / 10 | 7.40e-04 |
| WNT5:FZD7-mediated leishmania damping | 3 / 20 | 0.001 | 0.002 | 0.044 | 7 / 9 | 6.66e-04 |
| Killing mechanisms | 3 / 20 | 0.001 | 0.002 | 0.044 | 7 / 9 | 6.66e-04 |
| TCR signaling | 7 / 147 | 0.01 | 0.002 | 0.044 | 16 / 52 | 0.004 |
| RHOU GTPase cycle | 4 / 44 | 0.003 | 0.002 | 0.044 | 1 / 4 | 2.96e-04 |
| MHC class II antigen presentation | 7 / 148 | 0.01 | 0.002 | 0.044 | 24 / 26 | 0.002 |
| RIPK1-mediated regulated necrosis | 4 / 46 | 0.003 | 0.002 | 0.049 | 4 / 34 | 0.003 |

* False Discovery Rate

### 5. Pathways details

For every pathway of the most significant pathways, we present its diagram, as well as a short summary, its bibliography and the list of inputs found in it.

#### 1. Generation of second messenger molecules (R-HSA-202433)

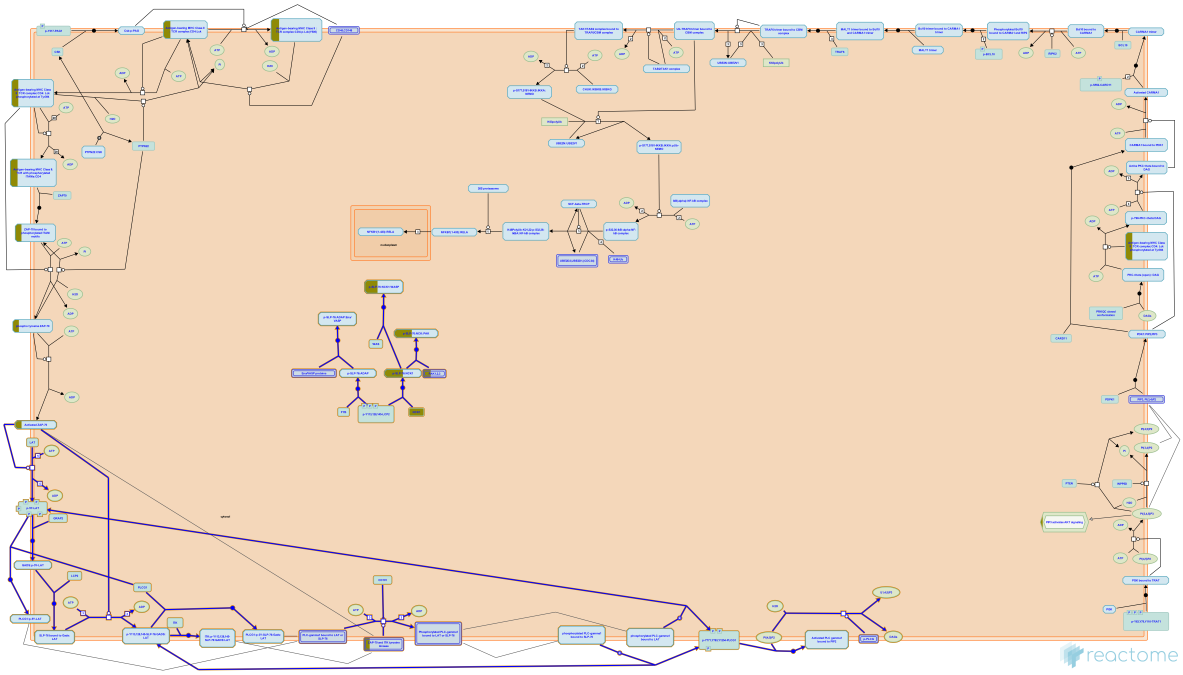

**Cellular compartments:** plasma membrane.

In addition to serving as a scaffold via auto-phosphorylation, ZAP70 also phosphorylates a restricted set of substrates following TCR stimulation - including LAT (step 13) and LCP2. These substrates have been recognized to play pivotal role in TCR signaling by releasing second messengers. When phosphorylated, LAT and SLP-76 act as adaptor proteins which serve as nucleation points for the construction of a higher order signalosome: PLC-gamma1 (step 14) and GRAP2 (step 15) bind to the LAT on the phosphorylated tyrosine residues. LCP2 is then moved to the signalosome by interacting with the SH3 domains of GRAP2 using their proline rich sequences (step 16). Once LCP2 binds to GRAP2, three LCP2 acidic domain N-term tyrosine residues are phosphorylated by ZAP70 (step 17). These phospho-tyrosine residues act as binding sites to the SH2 domains of ITK (steps 18) and PLC-gamma1 (step 19).
PLC-gamma1 is activated by dual phosphorylation on the tyrosine residues at positions 771, 783 and 1254 by ITK (step 20) and ZAP70 (step 21). Phosphorylated PLC-gamma1 subsequently detaches from LAT and LCP2 and translocates to the plasma membrane by binding to phosphatidylinositol-4,5-bisphosphate (PIP2) via its PH domain (step 22). PLC-gamma1 goes on to hydrolyse PIP2 to second messengers DAG and IP3 (step 23). These second messengers are involved in PKC and NF-kB activation and calcium mobilization.

#### Edit history

| Date | Action | Author |
|---|---|---|
| 2007-10-29 | Created | Garapati P V |
| 2008-01-24 | Authored | Rudd C.E., Garapati P V, de Bono B |
| 2008-02-26 | Reviewed | Trowsdale J |
| 2021-05-22 | Modified | Shorser S |

#### Entities found in this pathway (6)

| Input | UniProt Id | Input | UniProt Id | Input | UniProt Id |
|---|---|---|---|---|---|
| HLA-DPA1 | P20036 | HLA-DPB1 | P04440 | HLA-DQB1 | P01920 |
| HLA-DRA | P01903, P01909 | NCK1 | P16333 | PAK1 | Q13153 |

#### 2. Cytokine Signaling in Immune system (R-HSA-1280215)

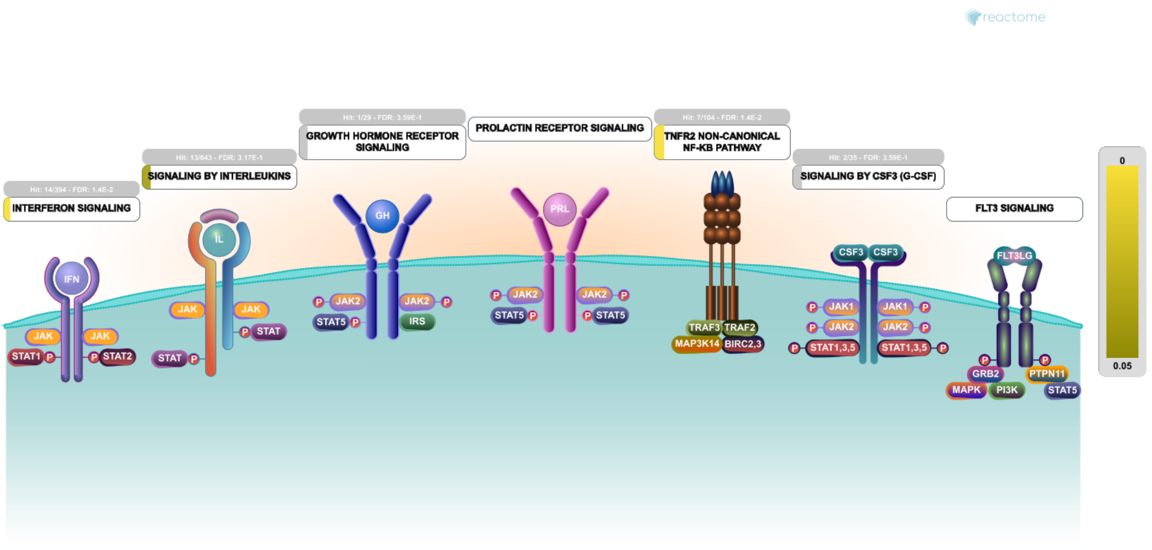


Cytokines are small proteins that regulate and mediate immunity, inflammation, and hematopoiesis. They are secreted in response to immune stimuli, and usually act briefly, locally, at very low concentrations. Cytokines bind to specific membrane receptors, which then signal the cell via second messengers, to regulate cellular activity.

IMMPORT:Bioinformatics for the future of immunology. Retrieved from https://www.immport.org/immportWeb/queryref/geneListSummary.do

COPE. Retrieved from http://www.copewithcytokines.org/cope.cgi

Santamaria P (2003). Cytokines and chemokines in autoimmune disease: an overview. Adv Exp Med Biol, 520, 1-7.


##### Edit history

| Date | Action | Author |
|---|---|---|
| 2011-05-12 | Created | Garapati P V |
| 2011-05-22 | Edited | Ray KP, Jupe S, Garapati P V |
| 2011-05-22 | Authored | Ray KP, Jupe S, Garapati P V |
| 2011-05-29 | Reviewed | Abdul-Sater AA, Schindler C, Pinteaux E |
| 2021-05-22 | Modified | Shorser S |

##### Entities found in this pathway (17)

| Input | UniProt Id | Input | UniProt Id | Input | UniProt Id |
|---|---|---|---|---|---|
| BIRC3 | Q13489, Q13490 | CSF2RB | P24394, P32927 | EIF4G2 | P78344, Q04637 |
| HLA-DPA1 | P20036 | HLA-DPB1 | P04440 | HLA-DQB1 | P01920 |
| HLA-DRA | P01903, P01909 | IL23A | Q9NPF7 | IL2RG | P31785 |

| Input | UniProt Id |
|---|---|
| ITGAM | P11215, P20702 |
| PTAFR | P25105 |
| TNFRSF11A | Q9Y6Q6 |

| Input | UniProt Id |
|---|---|
| LTB | Q06643 |
| RELB | Q01201 |
| TNFRSF11B | O00300 |

| Input | UniProt Id |
|---|---|
| NFKB2 | Q00653 |
| SOCS1 | O15524 |

| Input | Ensembl Id |
|---|---|
| HLA-DPA1 | ENSG00000231389 |
| HLA-DRA | ENSG00000204287 |
| PTAFR | ENSG00000169403 |

| Input | Ensembl Id |
|---|---|
| HLA-DPB1 | ENSG00000223865 |
| IL23A | ENST00000619177 |
| SOCS1 | ENSG00000185338 |

| Input | Ensembl Id |
|---|---|
| HLA-DQB1 | ENSG00000179344 |
| ITGAM | ENSG00000169896 |

#### 3. Interferon gamma signaling (R-HSA-877300)

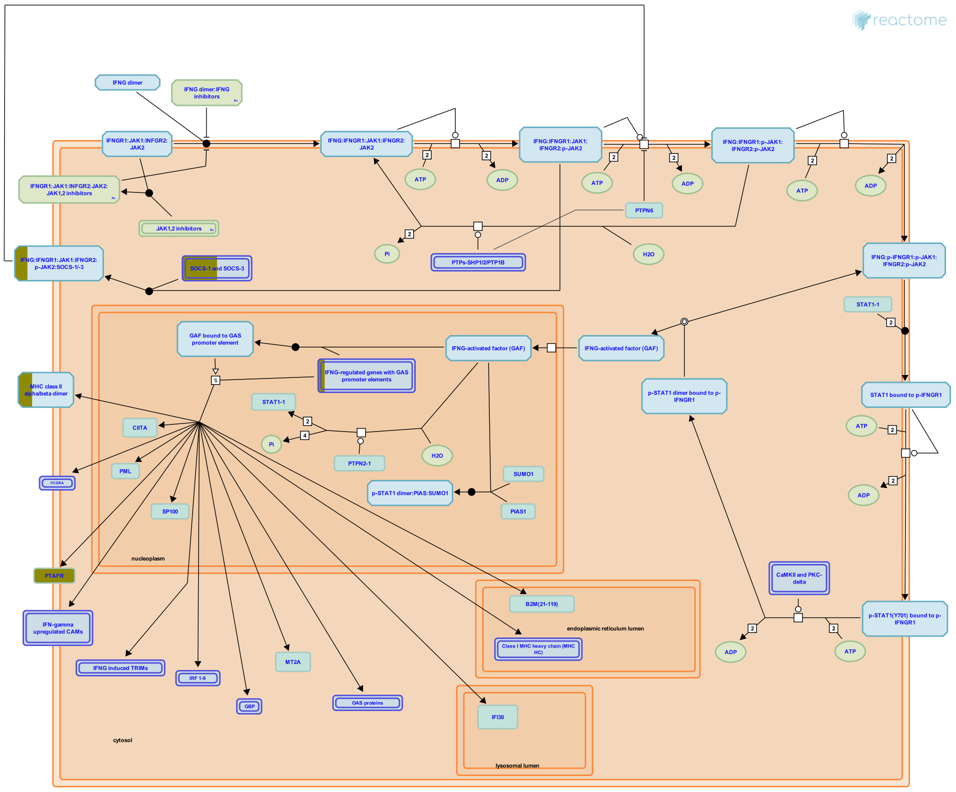


Interferon-gamma (IFN-gamma) belongs to the type II interferon family and is secreted by activated immune cells-primarily T and NK cells, but also B-cells and APC. INFG exerts its effect on cells by interacting with the specific IFN-gamma receptor (IFNGR). IFNGR consists of two chains, namely IFNGR1 (also known as the IFNGR alpha chain) and IFNGR2 (also known as the IFNGR beta chain). IFNGR1 is the ligand binding receptor and is required but not sufficient for signal transduction, whereas IFNGR2 do not bind IFNG independently but mainly plays a role in IFNG signaling and is generally the limiting factor in IFNG responsiveness. Both IFNGR chains lack intrinsic kinase/phosphatase activity and thus rely on other signaling proteins like Janus-activated kinase 1 (JAK1), JAK2 and Signal transducer and activator of transcription 1 (STAT-1) for signal transduction. IFNGR complex in its resting state is a preformed tetramer and upon IFNG association undergoes a conformational change. This conformational change induces the phosphorylation and activation of JAK1, JAK2, and STAT1 which in turn induces genes containing the gamma-interferon activation sequence (GAS) in the promoter.

#### Edit history

| Date | Action | Author |
|---|---|---|
| 2010-06-08 | Edited | Garapati P V |
| 2010-06-08 | Authored | Garapati P V |
| 2010-06-11 | Created | Garapati P V |
| 2010-08-17 | Reviewed | Abdul-Sater AA, Schindler C |
| 2021-05-31 | Modified | Shorser S |

#### Entities found in this pathway (6)

| Input | UniProt Id | Input | UniProt Id | Input | UniProt Id |
|---|---|---|---|---|---|
| HLA-DPA1 | P20036 | HLA-DPB1 | P04440 | HLA-DQB1 | P01920 |
| HLA-DRA | P01903, P01909 | PTAFR | P25105 | SOCS1 | O15524 |

| Input | Ensembl Id | Input | Ensembl Id | Input | Ensembl Id |
|---|---|---|---|---|---|
| HLA-DPA1 | ENSG00000231389 | HLA-DPB1 | ENSG00000223865 | HLA-DQB1 | ENSG00000179344 |
| HLA-DRA | ENSG00000204287 | PTAFR | ENSG00000169403 | | |

#### 4. TNF receptor superfamily (TNFSF) members mediating non-canonical NF-kB pathway (R-HSA-5676594)

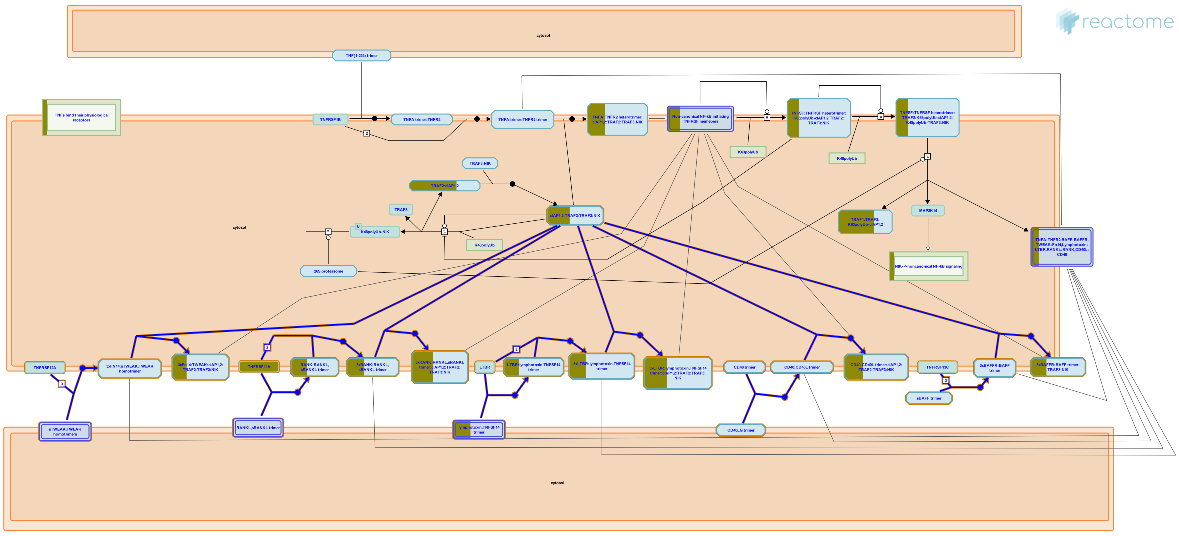

**Cellular compartments:** plasma membrane, cytosol, extracellular region.

Activation of NF-kB is fundamental to signal transduction by members of the TNFRSF. Expression of NF-kB target genes is essential for mounting innate immune responses to infectious microorganisms but is also important for the proper development and cellular compartmentalization of secondary lymphoid organs necessary to orchestrate an adaptive immune response.

NF-kB transcription factor family is activated by two distinct pathways: the canonical pathway involving NF-kB1 and the non-canonical pathway involving NF-kB2. Unlike NF-kB1 signalling, which can be activated by a wide variety of receptors, the NF-kB2 pathway is typically activated by a subset of receptor and ligand pairs belonging to the tumor necrosis factor receptor (TNF) super family (TNFRSF) members. These members include TNFR2 (Rauert et al. 2010), B cell activating factor of the TNF family receptor (BAFFR also known as TNFRSF13C) (Kayagaki et al. 2002, CD40 (also known as TNFRSF5) (Coope et al. 2002, lymphotoxin beta-receptor (LTBR also known as TNFRSF3) (Dejardin et al. 2002), receptor activator for nuclear factor kB (RANK also known as TNFRSF11A) (Novack et al. 2003), CD27 and Fibroblast growth factor-inducible immediate-early response protein 14 (FN14 also known as TNFRSF12A) etc. These receptors each mediate specific biological roles of the non-canonical NF-kB. These non-canonical NF-kB-stimulating receptors have one thing in common and is the presence of a TRAF-binding motif, which recruits different TNF receptor-associated factor (TRAF) members, particularly TRAF2 and TRAF3, to the receptor complex during ligand ligation (Grech et al. 2004, Bishop & Xie 2007). Receptor recruitment of these TRAF members leads to their degradation which is a critical step leading to the activation of NIK and induction of p100 processing (Sun 2011, 2012).

#### Edit history

| Date | Action | Author |
|---|---|---|
| 2015-01-26 | Edited | Garapati P V |
| 2015-01-26 | Authored | Garapati P V |
| 2015-02-20 | Created | Garapati P V |
| 2015-05-12 | Reviewed | Virgen-Slane R, Ware CF, Rajput A |
| 2021-05-21 | Modified | Shorser S |

#### Entities found in this pathway (3)

| Input | UniProt Id | Input | UniProt Id | Input | UniProt Id |
|---|---|---|---|---|---|
| BIRC3 | Q13489, Q13490 | LTB | Q06643 | TNFRSF11A | Q9Y6Q6 |

#### 5. Translocation of ZAP-70 to Immunological synapse (R-HSA-202430)

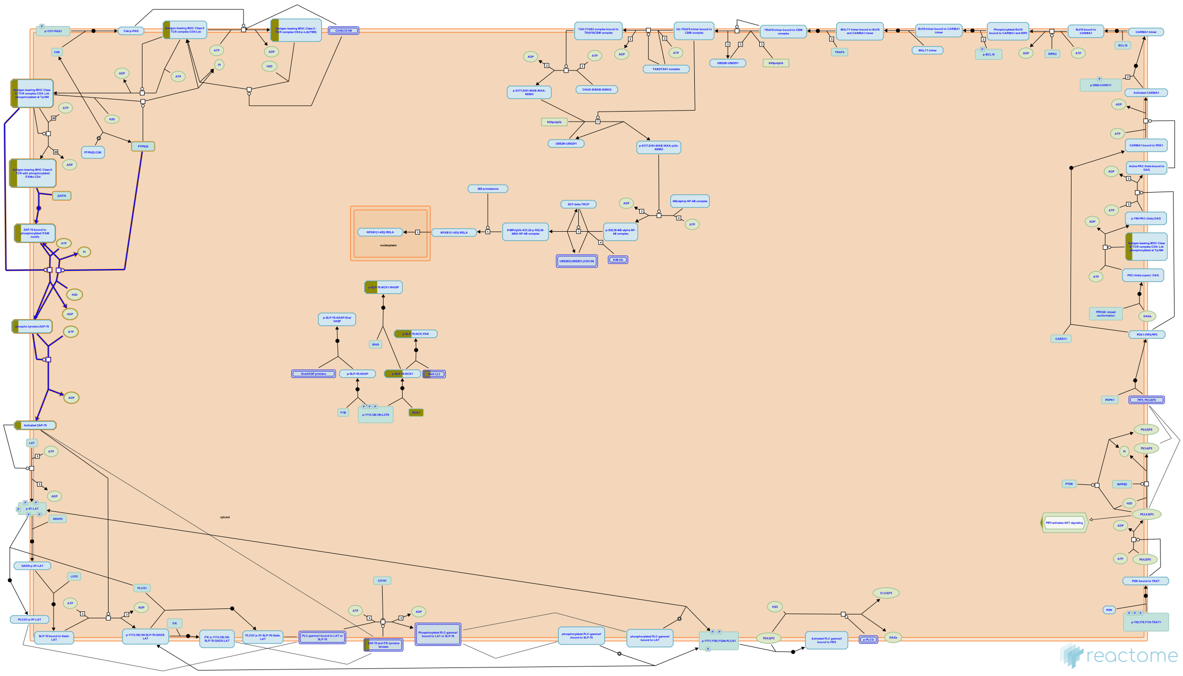

**Cellular compartments:** plasma membrane.

The dual phosphorylated ITAMs recruit SYK kinase ZAP70 via their tandem SH2 domains (step 8). ZAP70 subsequently undergoes phosphorylation on multiple tyrosine residues for further activation. ZAP70 includes both positive and negative regulatory sites. Tyrosine 493 is a conserved regulatory site found within the activation loop of the kinase domain. This site has shown to be a positive regulatory site required for ZAP70 kinase activity and is phosphorylated by LCK (step 9). This phosphorylation contributes to the active conformation of the catalytic domain. Later ZAP70 undergoes trans-autophosphorylation at Y315 and Y319 (step 10). These sites appear to be positive regulatory sites. ZAP70 achieves its full activation after the trans-autophosphorylation. Activated ZAP70 along with LCK phosphorylates the multiple tyrosine residues in the adaptor protein LAT (step 11). PTPN22 can dephosphorylate and inhibit ZAP70 activity to downregulate TCR signaling (step 12).


##### Edit history

| Date | Action | Author |
|---|---|---|
| 2007-10-29 | Created | Garapati P V |
| 2008-01-24 | Authored | Rudd C.E., Garapati P V, de Bono B |
| 2008-02-26 | Reviewed | Trowsdale J |
| 2016-05-10 | Revised | Stanford S, Bottini N |
| 2021-05-21 | Modified | Shorser S |

##### Entities found in this pathway (4)

| Input | UniProt Id | Input | UniProt Id |
|---|---|---|---|
| HLA-DPA1 | P20036 | HLA-DPB1 | P04440 |
| HLA-DQB1 | P01920 | HLA-DRA | P01903, P01909 |

#### 6. Immune System (R-HSA-168256)

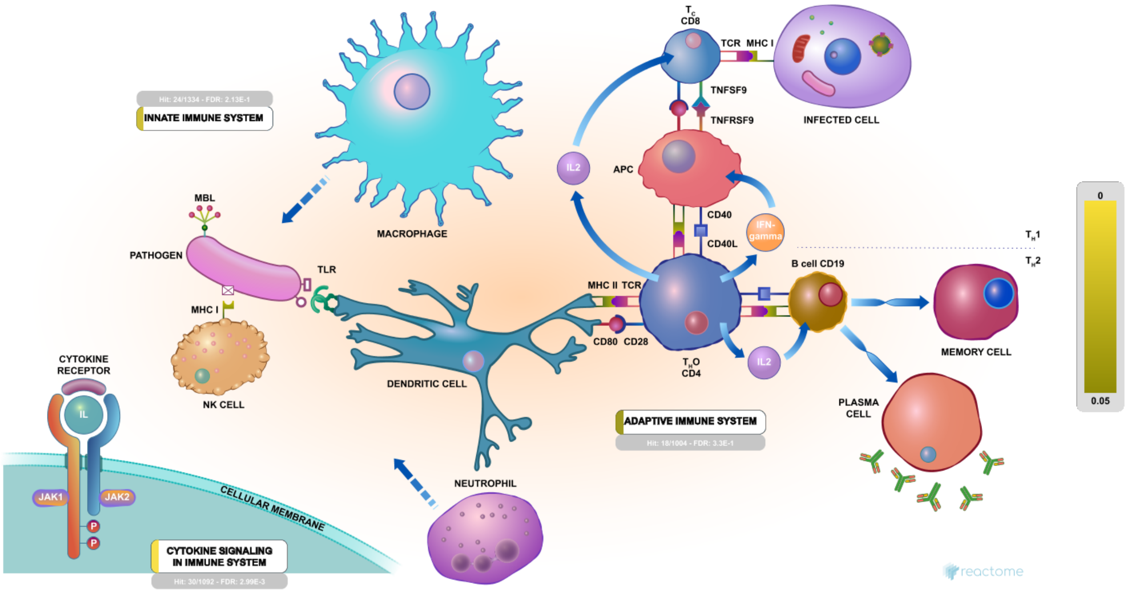


Humans are exposed to millions of potential pathogens daily, through contact, ingestion, and inhalation. Our ability to avoid infection depends on the adaptive immune system and during the first critical hours and days of exposure to a new pathogen, our innate immune system.

##### References

##### Edit history

| Date | Action | Author |
|---|---|---|
| 2005-11-12 | Created | Gillespie ME |
| 2006-03-30 | Authored | Luo F, Ouwehand WH, Gillespie ME, de Bono B |
| 2006-04-19 | Reviewed | Zwaginga JJ, D'Eustachio P, Gay NJ, Gale M Jr |
| 2021-05-22 | Modified | Shorser S |

##### Entities found in this pathway (38)

| Input | UniProt Id | Input | UniProt Id | Input | UniProt Id |
|---|---|---|---|---|---|
| ADAM8 | P78325 | BIRC3 | Q13489, Q13490 | BTN2A2 | Q8WVV5 |
| CD74 | P04233 | COTL1 | Q14019 | CSF2RB | P24394, P32927 |
| CTSZ | Q9UBR2 | CYBA | P13498 | DAPP1 | Q9UN19 |
| DBNL | Q9UJU6 | EIF4G2 | P78344, Q04637 | HLA-DMA | P28067 |
| HLA-DPA1 | P20036 | HLA-DPB1 | P04440 | HLA-DQB1 | P01920 |
| HLA-DRA | P01903, P01909 | ICAM2 | P13598 | IL23A | Q9NPF7 |
| IL2RG | P31785 | ITGAM | P11215, P20702 | ITGAV | P06756 |
| LTB | Q06643 | LYZ | P61626 | NCF1 | P14598 |
| NCK1 | P16333 | NFKB2 | Q00653 | PAK1 | Q13153 |
| PIK3AP1 | Q6ZUJ8 | PLAU | P00749 | PTAFR | P25105 |
| RELB | Q01201 | S100A11 | P31949 | S100A9 | P06702 |
| SERPINB1 | P30740 | SOCS1 | O15524 | TNFAIP3 | P21580 |
| TNFRSF11A | Q9Y6Q6 | TNFRSF11B | O00300 | | |

| Input | Ensembl Id | Input | Ensembl Id | Input | Ensembl Id |
|---|---|---|---|---|---|
| HLA-DPA1 | ENSG00000231389 | HLA-DPB1 | ENSG00000223865 | HLA-DQB1 | ENSG00000179344 |
| HLA-DRA | ENSG00000204287 | IL23A | ENST00000619177 | ITGAM | ENSG00000169896 |
| PTAFR | ENSG00000169403 | SOCS1 | ENSG00000185338 | | |

#### 7. Phosphorylation of CD3 and TCR zeta chains (R-HSA-202427)

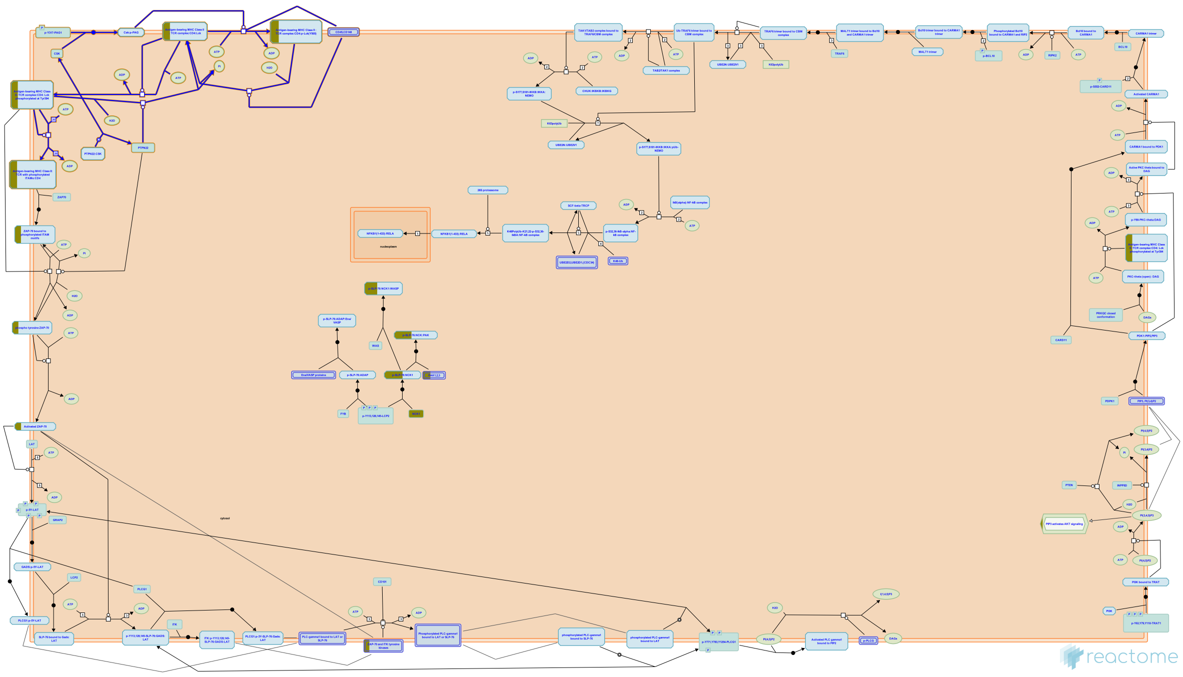

**Cellular compartments:** plasma membrane.

Prior to T cell receptor (TCR) stimulation, CD4/CD8 associated LCK remains seperated from the TCR and is maintained in an inactive state by the action of CSK. PAG bound CSK phosphorylates the negative regulatory tyrosine of LCK and inactivates the LCK kinase domain (step 1). CSK also inhibits PTPN22 by sequestering it via binding (step 2).
Upon TCR stimulation, CSK dissociates from PAG1 (step 3) and PTPN22 (step4) and is unable to inhibit LCK. Furthermore, LCK becomes activated via PTPRC-mediated dephosphorylation of negative regulatory tyrosine residues (step 5). CD4/CD8 binds MHCII receptor in APC and the associated LCK co-localizes with the TCR.
LCK is further activated by trans-autophosphorylation on the tyrosine residue on its activation loop (step 6). Active LCK further phosphorylates the tyrosine residues on CD3 chains. The signal-transducing CD3 delta/epsilon/gamma and TCR zeta chains contain a critical signaling motif known as the immunoreceptor tyrosine-based activation motif (ITAM). The two critical tyrosines of each ITAM motif are phosphorylated by LCK (step 7).


##### Edit history

| Date | Action | Author |
|---|---|---|
| 2007-10-29 | Created | Garapati P V |
| 2008-01-24 | Authored | Rudd C.E., Garapati P V, de Bono B |
| 2008-02-26 | Reviewed | Trowsdale J |
| 2016-05-10 | Revised | Stanford S, Bottini N |
| 2021-05-21 | Modified | Shorser S |

#### Entities found in this pathway (4)

| Input | UniProt Id | Input | UniProt Id |
|---|---|---|---|
| HLA-DPA1 | P20036 | HLA-DPB1 | P04440 |
| HLA-DQB1 | P01920 | HLA-DRA | P01903, P01909 |

#### 8. PD-1 signaling (R-HSA-389948)

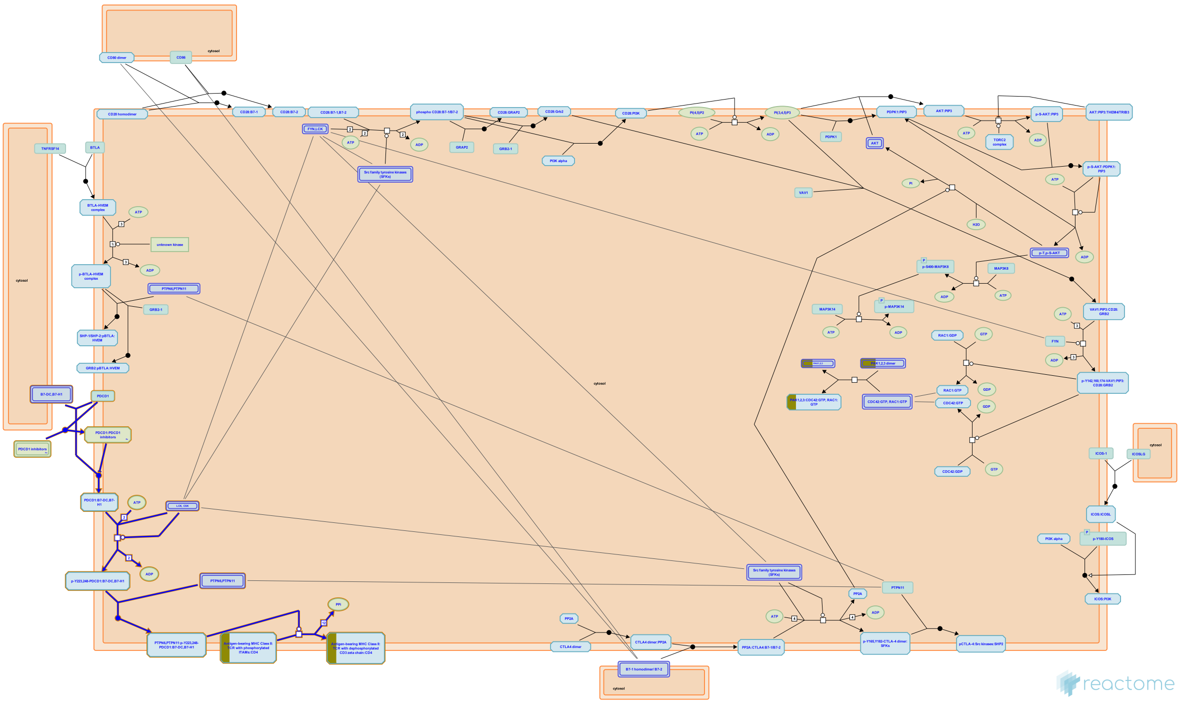

**Cellular compartments:** plasma membrane.

The Programmed cell death protein 1 (PD-1) is one of the negative regulators of TCR signaling. PD-1 may exert its effects on cell differentiation and survival directly by inhibiting early activation events that are positively regulated by CD28 or indirectly through IL-2. PD-1 ligation inhibits the induction of the cell survival factor Bcl-xL and the expression of transcription factors associated with effector cell function, including GATA-3, Tbet, and Eomes. PD-1 exerts its inhibitory effects by bringing phosphatases SHP-1 and SHP-2 into the immune synapse, leading to dephosphorylation of CD3-zeta chain, PI3K and AKT.


##### Edit history

| Date | Action | Author |
|---|---|---|
| 2008-12-16 | Edited | Garapati P V |
| 2008-12-16 | Authored | Garapati P V |
| 2009-01-21 | Created | Garapati P V |
| 2009-06-01 | Reviewed | Bluestone JA, Esensten J |
| 2021-05-21 | Modified | Shorser S |

##### Entities found in this pathway (4)

| Input | UniProt Id | Input | UniProt Id |
| --- | --- | --- | --- |
| HLA-DPA1 | P20036 | HLA-DPB1 | P04440 |
| HLA-DQB1 | P01920 | HLA-DRA | P01903, P01909 |

#### 9. Interferon Signaling (R-HSA-913531)

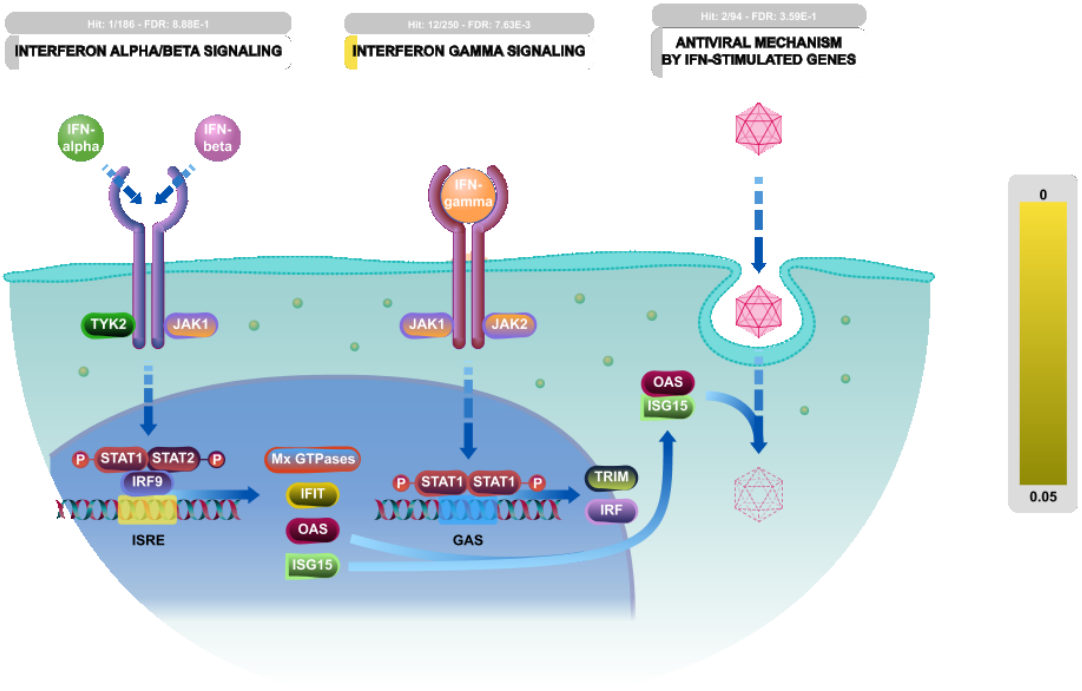


Interferons (IFNs) are cytokines that play a central role in initiating immune responses, especially antiviral and antitumor effects. There are three types of IFNs:Type I (IFN-alpha, -beta and others, such as omega, epsilon, and kappa), Type II (IFN-gamma) and Type III (IFN-lamda). In this module we are mainly focusing on type I IFNs alpha and beta and type II IFN-gamma. Both type I and type II IFNs exert their actions through cognate receptor complexes, IFNAR and IFNGR respectively, present on cell surface membranes. Type I IFNs are broadly expressed heterodimeric receptors composed of the IFNAR1 and IFNAR2 subunits, while the type II IFN receptor consists of IFNGR1 and IFNGR2. Type III interferon lambda has three members: lamda1 (IL-29), lambda2 (IL-28A), and lambda3 (IL-28B) respectively. IFN-lambda signaling is initiated through unique heterodimeric receptor composed of IFN-LR1/IF-28Ralpha and IL10R2 chains.

Type I IFNs typically recruit JAK1 and TYK2 proteins to transduce their signals to STAT1 and 2; in combination with IRF9 (IFN-regulatory factor 9), these proteins form the heterotrimeric complex ISGF3. In nucleus ISGF3 binds to IFN-stimulated response elements (ISRE) to promote gene induction.

Type II IFNs in turn rely upon the activation of JAKs 1 and 2 and STAT1. Once activated, STAT1 dimerizes to form the transcriptional regulator GAF (IFNG activated factor) and this binds to the IFNG activated sequence (GAS) elements and initiate the transcription of IFNG-responsive genes.

Like type I IFNs, IFN-lambda recruits TYK2 and JAK1 kinases and then promote the phosphorylation of STAT1/2, and induce the ISRE3 complex formation.

#### Edit history

| Date | Action | Author |
|---|---|---|
| 2010-07-07 | Edited | Garapati P V |
| 2010-07-07 | Authored | Garapati P V |
| 2010-07-16 | Created | Garapati P V |
| 2010-08-17 | Reviewed | Abdul-Sater AA, Schindler C |
| 2021-05-22 | Modified | Shorser S |

#### Entities found in this pathway (7)

| Input | UniProt Id | Input | UniProt Id | Input | UniProt Id |
|---|---|---|---|---|---|
| EIF4G2 | P78344, Q04637 | HLA-DPA1 | P20036 | HLA-DPB1 | P04440 |
| HLA-DQB1 | P01920 | HLA-DRA | P01903, P01909 | PTAFR | P25105 |
| SOCS1 | O15524 | | | | |

| Input | Ensembl Id | Input | Ensembl Id | Input | Ensembl Id |
|---|---|---|---|---|---|
| HLA-DPA1 | ENSG00000231389 | HLA-DPB1 | ENSG00000223865 | HLA-DQB1 | ENSG00000179344 |
| HLA-DRA | ENSG00000204287 | PTAFR | ENSG00000169403 | | |

#### 10. TNFR2 non-canonical NF-kB pathway (R-HSA-5668541)

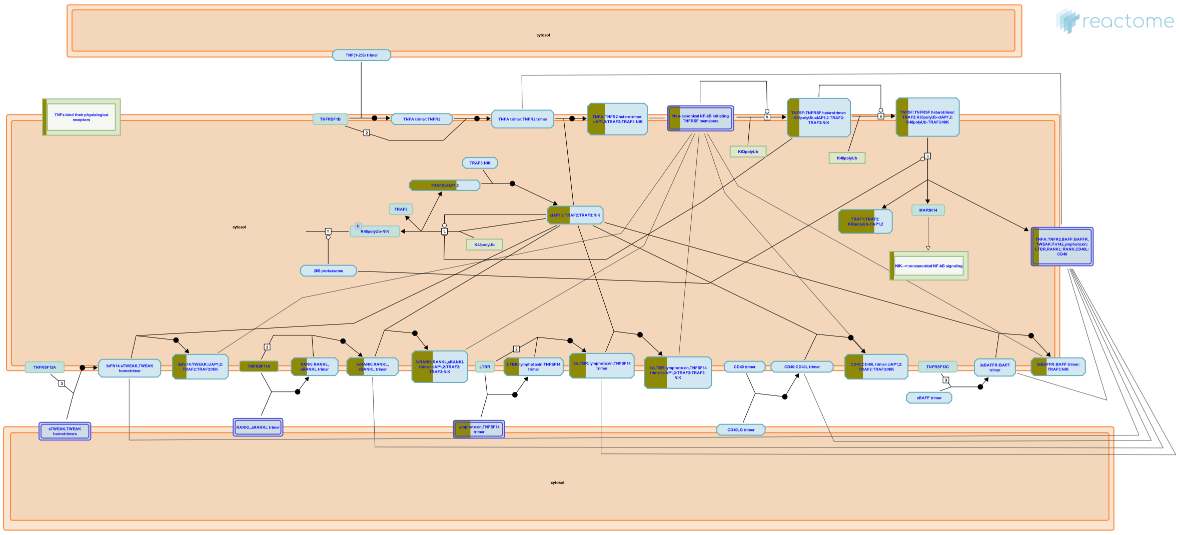

**Cellular compartments:** plasma membrane, nucleoplasm.

Tumor necrosis factor-alpha (TNFA) exerts a wide range of biological effects through TNF receptor 1 (TNFR1) and TNF receptor 2 (TNFR2). Under normal physiological conditions TNFR2 exhibits more restricted expression, being found on certain subpopulation of immune cells and few other cell types (Grell et al. 1995 ). TNFR1 mediated signalling pathways have been very well characterized but, TNFR2 has been much less well studied. TNFR1 upon activation by TNFA activates apoptosis through two pathways, involving the adaptor proteins TNFR1-associated death domain (TRADD) and fas-associated death domain (FADD). In contrast, TNFR2 signalling especially in highly activated T cells, induces cell survival pathways that can result in cell proliferation by activating transcription factor NF-kB (nuclear factor-kB) via the alternative non-canonical route. TNFR2 signalling seems to play an important role, in particular for the function of regulatory T cells. It offers protective roles in several disorders, including autoimmune diseases, heart diseases, demyelinating and neurodegenerative disorders and infectious diseases (Faustman & Davis 2010).

Activation of the non-canonical pathway by TNFR2 is mediated through a signalling complex that includes TNF receptor-associated factor (TRAF2 and TRAF3), cellular inhibitor of apoptosis (cIAP1 and cIAP2), and NF-kB-inducing kinase (NIK). In this complex TRAF3 functions as a bridging factor between the cIAP1/2:TRAF2 complex and NIK. In resting cells cIAP1/2 in the signalling complex mediates K48-linked polyubiquitination of NIK and subsequent proteasomal degradation making NIK levels invisible. Upon TNFR2 stimulation, TRAF2 is recruited to the intracellular TRAF binding motif and this also indirectly recruits TRAF1 and cIAP1/2, as well as TRAF3 and NIK which are already bound to TRAF2 in unstimulated cells. TRAF2 mediates K63-linked ubiquitination of cIAP1/2 and this in turn mediates cIAP dependent K48-linked ubiquitination of TRAF3 leading to the proteasome-dependent degradation of the latter. As TRAF3 is degraded, NIK can no longer interact with TRAF1/2:cIAP complex. As a result NIK concentration in the cytosol increases and NIK gets stabilised and activated. Activated NIK phosphorylates IKKalpha, which in turn phosphorylates p100 (NFkB2) subunit. Phosphorylated p100 is also ubiquitinated by the SCF-beta-TRCP ubiquitin ligase complex and is subsequently processed by the proteaseome to p52, which is a transcriptionally competent NF-kB subunit in conjunction with RelB (Petrus et al. 2011, Sun 2011, Vallabhapurapu & Karin 2009).

#### Edit history

| Date | Action | Author |
|---|---|---|
| 2015-01-26 | Edited | Garapati P V |
| 2015-01-26 | Authored | Garapati P V |
| 2015-01-26 | Created | Garapati P V |
| 2015-05-12 | Reviewed | Virgen-Slane R, Ware CF, Rajput A |
| 2021-05-22 | Modified | Shorser S |

#### Entities found in this pathway (6)

| Input | UniProt Id | Input | UniProt Id | Input | UniProt Id |
|---|---|---|---|---|---|
| BIRC3 | Q13489, Q13490 | LTB | Q06643 | NFKB2 | Q00653 |
| RELB | Q01201 | TNFRSF11A | Q9Y6Q6 | TNFRSF11B | O00300 |

#### 11. Cross-presentation of particulate exogenous antigens (phagosomes) (R-HSA-1236973)

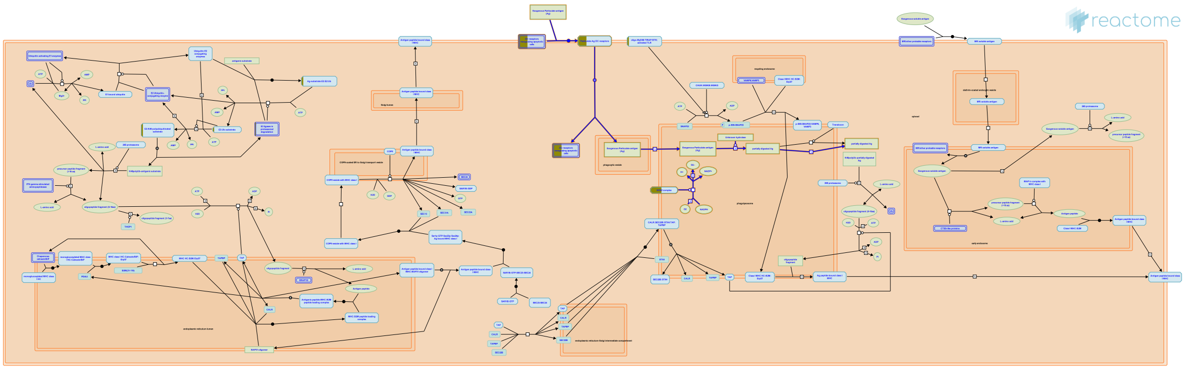

Dendritic cells (DCs) take up and process exogenous particulate or cell-associated antigens such as microbes or tumor cells for MHC-I cross-presentation. Particulate antigens have been reported to be more efficiently cross-presented than soluble antigens by DCs (Khor et al. 2008). Particulate antigens are internalized by phagosomes. There are two established models that explain the mechanism by which exogenous particulate antigens are presented through MHC I; the cytosolic pathway where internalized antigens are somehow translocated from phagosomes into cytosol for proteasomal degradation and the vacuolar pathway (Lin et al. 2008, Amigorena et al. 2010).

##### References

##### Edit history

| Date | Action | Author |
|---|---|---|
| 2011-03-28 | Edited | Garapati P V |
| 2011-03-28 | Authored | Garapati P V |
| 2011-03-28 | Created | Garapati P V |
| 2011-05-13 | Reviewed | Desjardins M, English L |
| 2021-05-22 | Modified | Shorser S |

##### Entities found in this pathway (3)

| Input | UniProt Id | Input | UniProt Id | Input | UniProt Id |
|---|---|---|---|---|---|
| CYBA | P13498 | ITGAV | P06756 | NCF1 | P14598 |

#### 12. Activation of RAC1 (R-HSA-428540)

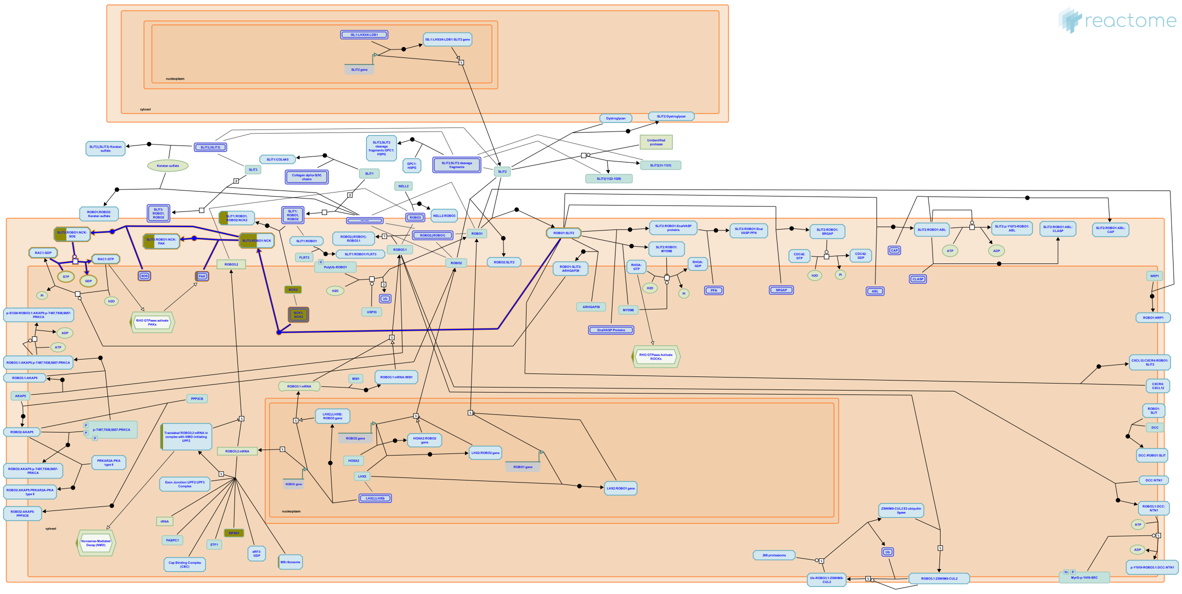

A low level of RAC1 activity is essential to maintain axon outgrowth. ROBO activation recruits SOS, a dual specificity GEF, to the plasma membrane via Dock homolog NCK (NCK1 or NCK2) to activate RAC1 during midline repulsion.

##### References

##### Edit history

| Date | Action | Author |
|---|---|---|
| 2008-09-05 | Edited | Garapati P V |
| 2008-09-05 | Authored | Garapati P V |
| 2009-07-06 | Created | Garapati P V |
| 2009-08-18 | Reviewed | Kidd T |
| 2017-06-26 | Edited | Orlic-Milacic M |
| 2017-07-31 | Reviewed | Jaworski A |
| 2020-11-11 | Modified | D'Eustachio P |

##### Entities found in this pathway (2)

| Input | UniProt Id | Input | UniProt Id |
|---|---|---|---|
| NCK1 | O43639, P16333 | PAK1 | Q13153 |

#### 13. Interleukin-4 and Interleukin-13 signaling (R-HSA-6785807)

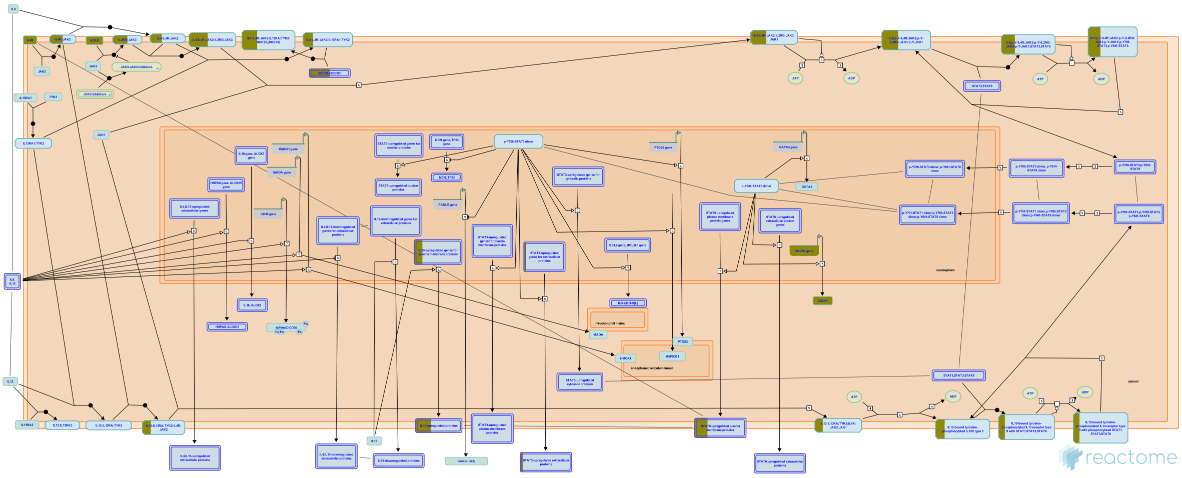

Interleukin-4 (IL4) is a principal regulatory cytokine during the immune response, crucially important in allergy and asthma (Nelms et al. 1999). When resting T cells are antigen-activated and expand in response to Interleukin-2 (IL2), they can differentiate as Type 1 (Th1) or Type 2 (Th2) T helper cells. The outcome is influenced by IL4. Th2 cells secrete IL4, which both stimulates Th2 in an autocrine fashion and acts as a potent B cell growth factor to promote humoral immunity (Nelms et al. 1999).

Interleukin-13 (IL13) is an immunoregulatory cytokine secreted predominantly by activated Th2 cells. It is a key mediator in the pathogenesis of allergic inflammation. IL13 shares many functional properties with IL4, stemming from the fact that they share a common receptor subunit. IL13 receptors are expressed on human B cells, basophils, eosinophils, mast cells, endothelial cells, fibroblasts, monocytes, macrophages, respiratory epithelial cells, and smooth muscle cells, but unlike IL4, not T cells. Thus IL13 does not appear to be important in the initial differentiation of CD4 T cells into Th2 cells, rather it is important in the effector phase of allergic inflammation (Hershey et al. 2003).

IL4 and IL13 induce “alternative activation” of macrophages, inducing an anti-inflammatory phenotype by signaling through IL4R alpha in a STAT6 dependent manner. This signaling plays an important role in the Th2 response, mediating anti-parasitic effects and aiding wound healing (Gordon & Martinez 2010, Loke et al. 2002)

There are two types of IL4 receptor complex (Andrews et al. 2006). Type I IL4R (IL4R1) is predominantly expressed on the surface of hematopoietic cells and consists of IL4R and IL2RG, the common gamma chain. Type II IL4R (IL4R2) is predominantly expressed on the surface of nonhematopoietic cells, it consists of IL4R and IL13RA1 and is also the type II receptor for IL13. (Obiri et al. 1995, Aman et al. 1996, Hilton et al. 1996, Miloux et al. 1997, Zhang et al. 1997). The second receptor for IL13 consists of IL4R and Interleukin-13 receptor alpha 2 (IL13RA2), sometimes called Interleukin-13 binding protein (IL13BP). It has a high affinity receptor for IL13 (Kd = 250 pmol/L) but is not sufficient to render cells responsive to IL13, even in the presence of IL4R (Donaldson et al. 1998). It is reported to exist in soluble form (Zhang et al. 1997) and when overexpressed reduces JAK-STAT signaling (Kawakami et al. 2001). It's function may be to prevent IL13 signalling via the functional IL4R:IL13RA1 receptor. IL13RA2 is overexpressed and enhances cell invasion in some human cancers (Joshi & Puri 2012).

The first step in the formation of IL4R1 (IL4:IL4R:IL2RB) is the binding of IL4 with IL4R (Hoffman et al. 1995, Shen et al. 1996, Hage et al. 1999). This is also the first step in formation of IL4R2 (IL4:IL4R:IL13RA1). After the initial binding of IL4 and IL4R, IL2RB binds (LaPorte et al. 2008), to form IL4R1. Alternatively, IL13RA1 binds, forming IL4R2. In contrast, the type II IL13 complex (IL13R2) forms with IL13 first binding to IL13RA1 followed by recruitment of IL4R (Wang et al. 2009).

Crystal structures of the IL4:IL4R:IL2RG, IL4:IL4R:IL13RA1 and IL13:IL4R:IL13RA1 complexes have been determined (LaPorte et al. 2008). Consistent with these structures, in monocytes IL4R is tyrosine phosphorylated in response to both IL4 and IL13 (Roy et al. 2002, Gordon & Martinez 2010) while IL13RA1 phosphorylation is induced only by IL13 (Roy et al. 2002, LaPorte et al. 2008) and IL2RG phosphorylation is induced only by IL4 (Roy et al. 2002).

Both IL4 receptor complexes signal through Jak/STAT cascades. IL4R is constitutively-associated with JAK2 (Roy et al. 2002) and associates with JAK1 following binding of IL4 (Yin et al. 1994) or IL13 (Roy et al. 2002). IL2RG constitutively associates with JAK3 (Boussiotis et al. 1994, Russell et al. 1994). IL13RA1 constitutively associates with TYK2 (Umeshita-Suyama et al. 2000, Roy et al. 2002, LaPorte et al. 2008, Bhattacharjee et al. 2013).

IL4 binding to IL4R1 leads to phosphorylation of JAK1 (but not JAK2) and STAT6 activation (Takeda et al. 1994, Ratthe et al. 2007, Bhattacharjee et al. 2013).

IL13 binding increases activating tyrosine-99 phosphorylation of IL13RA1 but not that of IL2RG. IL4 binding to IL2RG leads to its tyrosine phosphorylation (Roy et al. 2002). IL13 binding to IL4R2 leads to TYK2 and JAK2 (but not JAK1) phosphorylation (Roy & Cathcart 1998, Roy et al. 2002).

Phosphorylated TYK2 binds and phosphorylates STAT6 and possibly STAT1 (Bhattacharjee et al. 2013).

A second mechanism of signal transduction activated by IL4 and IL13 leads to the insulin receptor substrate (IRS) family (Kelly-Welch et al. 2003). IL4R1 associates with insulin receptor substrate 2 and activates the PI3K/Akt and Ras/MEK/Erk pathways involved in cell proliferation, survival and translational control. IL4R2 does not associate with insulin receptor substrate 2 and consequently the PI3K/Akt and Ras/MEK/Erk pathways are not activated (Busch-Dienstfertig & González-Rodríguez 2013).


#### Edit history

| Date | Action | Author |
|---|---|---|
| 2015-07-01 | Authored | Jupe S |
| 2015-07-01 | Created | Jupe S |
| 2016-09-02 | Edited | Jupe S |
| 2016-09-02 | Reviewed | Leibovich SJ |

| Date | Action | Author |
|---|---|---|
| 2021-05-31 | Modified | Shorser S |

#### Entities found in this pathway (5)

| Input | UniProt Id | Input | UniProt Id | Input | UniProt Id |
|---|---|---|---|---|---|
| CSF2RB | P24394 | IL23A | Q9NPF7 | IL2RG | P31785 |
| ITGAM | P11215, P20702 | SOCS1 | O15524 | | |

| Input | Ensembl Id | Input | Ensembl Id | Input | Ensembl Id |
|---|---|---|---|---|---|
| IL23A | ENST00000619177 | ITGAM | ENSG00000169896 | SOCS1 | ENSG00000185338 |

#### 14. Costimulation by the CD28 family (R-HSA-388841)

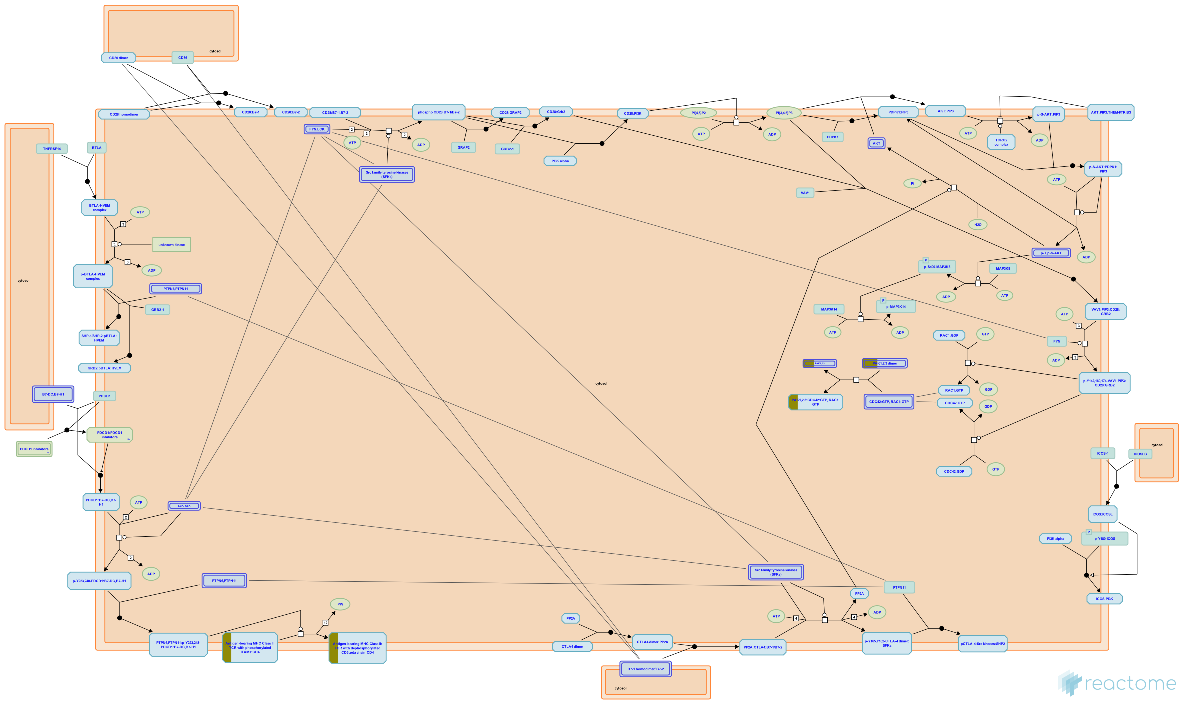

**Cellular compartments:** plasma membrane.

Optimal activation of T-lymphocytes requires at least two signals. A primary one is delivered by the T-cell receptor (TCR) complex after antigen recognition and additional costimulatory signals are delivered by the engagement of costimulatory receptors such as CD28. The best-characterized costimulatory pathways are mediated by a set of cosignaling molecules belonging to the CD28 superfamily, including CD28, CTLA4, ICOS, PD1 and BTLA receptors. These proteins deliver both positive and negative second signals to T-cells by interacting with B7 family ligands expressed on antigen presenting cells. Different subsets of T-cells have very different requirements for costimulation. CD28 family mediated costimulation is not required for all T-cell responses in vivo, and alternative costimulatory pathways also exist. Different receptors of the CD28 family and their ligands have different regulation of expression. CD28 is constitutively expressed on naive T cells whereas CTLA4 expression is dependent on CD28/B7 engagement and the other receptor members ICOS, PD1 and BTLA are induced after initial T-cell stimulation.

The positive signals induced by CD28 and ICOS molecules are counterbalanced by other members of the CD28 family, including cytotoxic T-lymphocyte associated antigen (CTLA)4, programmed cell death (PD)1, and B and T lymphocyte attenuator (BTLA), which dampen immune responses. The balance of stimulatory and inhibitory signals is crucial to maximize protective immune responses while maintaining immunological tolerance and preventing autoimmunity.

The costimulatory receptors CD28, CTLA4, ICOS and PD1 are composed of single extracellular IgV-like domains, whereas BTLA has one IgC-like domain. Receptors CTLA4, CD28 and ICOS are covalent homodimers, due to an interchain disulphide linkage. The costimulatory ligands B71, B72, B7H2, B7H1 and B7DC, have a membrane proximal IgC-like domain and a membrane distal IgV-like domain that is responsible for receptor binding and dimerization. CD28 and CTLA4 have no known intrinsic enzymatic activity. Instead, engagement by their physiologic ligands B71 and B72 leads to the physical recruitment and activation of downstream T-cell effector molecules.


#### Edit history

| Date | Action | Author |
|---|---|---|
| 2008-12-16 | Edited | Garapati P V |
| 2008-12-16 | Authored | Garapati P V |
| 2008-12-16 | Created | Garapati P V |
| 2009-06-01 | Reviewed | Bluestone JA, Esensten J |
| 2021-05-22 | Modified | Shorser S |

#### Entities found in this pathway (5)

| Input | UniProt Id | Input | UniProt Id | Input | UniProt Id |
|---|---|---|---|---|---|
| HLA-DPA1 | P20036 | HLA-DPB1 | P04440 | HLA-DQB1 | P01920 |
| HLA-DRA | P01903, P01909 | PAK1 | Q13153 | | |

#### 15. RHO GTPases Activate NADPH Oxidases (R-HSA-5668599)

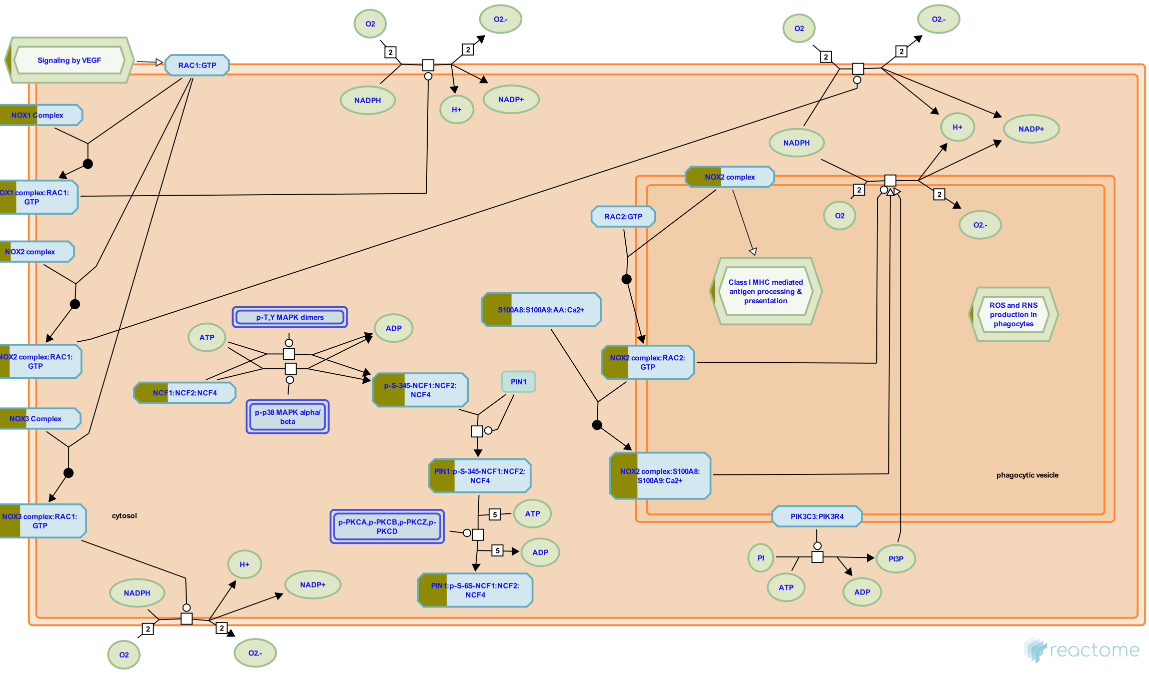


**Cellular compartments:** cytosol, plasma membrane, phagocytic vesicle membrane, phagolysosome.

NADPH oxidases (NOX) are membrane-associated enzymatic complexes that use NADPH as an electon donor to reduce oxygen and produce superoxide (O2-) that serves as a secondary messenger (Brown and Griendling 2009).

NOX2 complex consists of CYBB (NOX2), CYBA (p22phox), NCF1 (p47phox), NCF2 (p67phox) and NCF4 (p40ohox). RAC1:GTP binds NOX2 complex in response to VEGF signaling by directly interracting with CYBB and NCF2, leading to enhancement of VEGF-signaling through VEGF receptor VEGFR2, which plays a role in angiogenesis (Ushio-Fukai et al. 2002, Bedard and Krause 2007). RAC2:GTP can also activate the NOX2 complex by binding to CYBB and NCF2, leading to production of superoxide in phagosomes of neutrophils which is necessary fo the microbicidal activity of neutrophils (Knaus et al. 1991, Roberts et al. 1999, Kim and Dinauer 2001, Jyoti et al. 2014).

NOX1 complex (composed of NOX1, NOXA1, NOXO1 and CYBA) and NOX3 complex (composed of NOX3, CYBA, NCF1 amd NCF2 or NOXA1) can also be activated by binding to RAC1:GTP to produce superoxide (Cheng et al. 2006, Miyano et al. 2006, Ueyama et al. 2006).

#### Edit history

| Date | Action | Author |
|---|---|---|
| 2014-10-24 | Authored | Orlic-Milacic M |
| 2014-12-26 | Authored | Rivero Crespo F |
| 2015-01-26 | Created | Orlic-Milacic M |
| 2015-02-02 | Edited | Orlic-Milacic M |
| 2018-11-07 | Edited | Shamovsky V |
| 2018-11-08 | Revised | Shamovsky V |
| 2021-05-22 | Modified | Shorser S |

#### Entities found in this pathway (4)

| Input | UniProt Id | Input | UniProt Id |
|---|---|---|---|
| CYBA | P13498 | NCF1 | P14598 |
| NOXO1 | Q8NFA2 | S100A9 | P06702 |

#### 16. Regulation of necroptotic cell death (R-HSA-5675482)

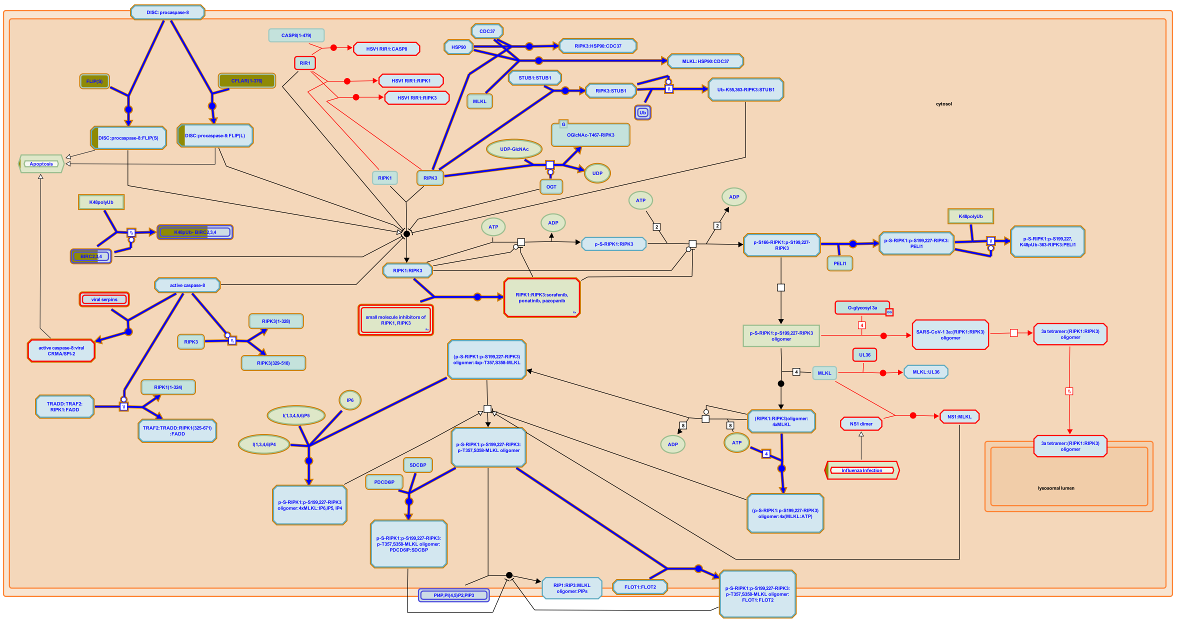

A regulated balance between cell survival and cell death is essential for normal development and homeostasis of multicellular organisms. Defects in control of this balance may contribute to autoimmune disease, neurodegeneration, ischemia/reperfusion injury, non alcoholic steatohepatitis (NASH) and cancer.


##### Edit history

| Date | Action | Author |
|---|---|---|
| 2015-02-10 | Authored | Shamovsky V |
| 2015-02-15 | Edited | Shamovsky V |
| 2015-02-15 | Reviewed | Chan FK |

| Date | Action | Author |
|---|---|---|
| 2015-02-16 | Created | Shamovsky V |
| 2020-08-28 | Reviewed | Murphy JM |
| 2021-05-31 | Modified | Shorser S |

#### Entities found in this pathway (2)

| Input | UniProt Id | Input | UniProt Id |
|---|---|---|---|
| BIRC3 | Q13489, Q13490 | CFLAR | O15519-1, O15519-2 |

#### 17. RHOV GTPase cycle (R-HSA-9013424)

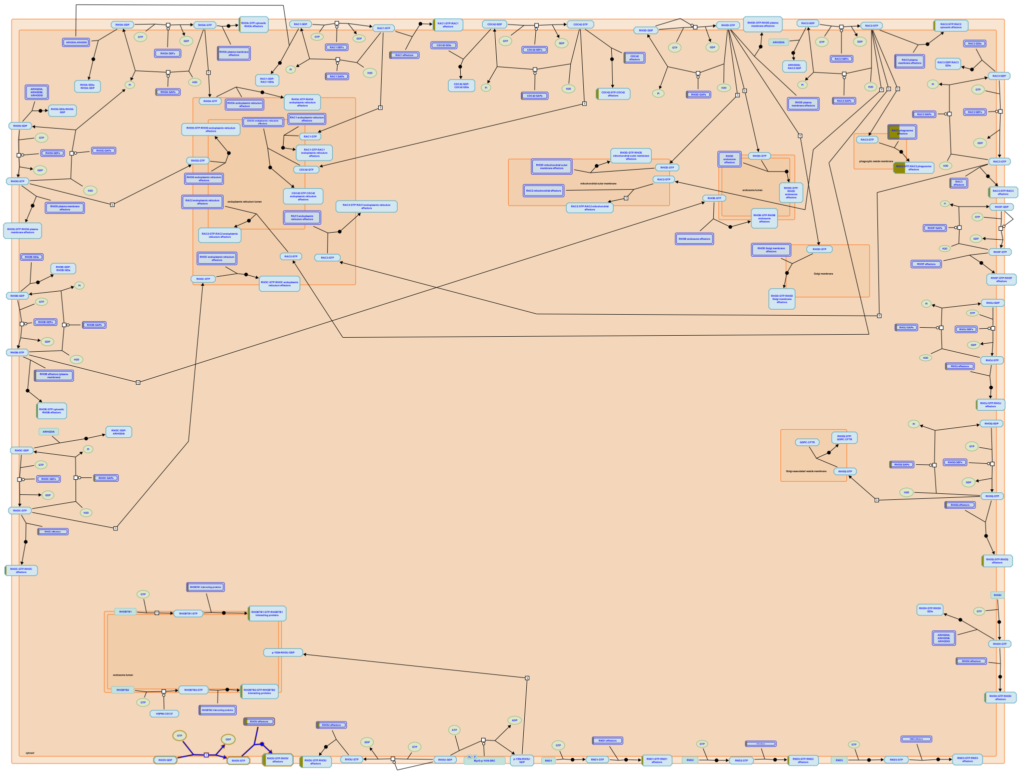

RHOV (also known as Chp) is an atypical RHO GTPase that is thought to be constitutively active due to its high intrinsic guanine nucleotide exchange activity. No guanine nucleotide exchange factors (GEFs) nor GTPase activator proteins (GAPs) that act on RHOV have been identified. RHOV is expressed at very low levels. The expression of RHOV is detected during embryonic development in fish (Tay et al. 2010), frog (Guémar et al. 2007) and chicken (Notarnicola et al. 2008). RHOV is involved in neural crest formation, where its expression is induced downstream of WNT signaling. RHOV is thought to regulate cell adhesion, as its zebrafish orthologue is required for proper localization of E-cadherin and beta-catenin at adherens junctions. RHOV activates JNK and induces apoptosis in rat pheochromocytoma cell line PC12 (Shepelev et al 2011) and in macrophages (Song et al. 2015).

RHOV gene overexpression is a molecular marker of human lung adenocarcinoma (Shepelev and Korobko 2013, Shukla et al. 2017, Ma et al. 2020, Zhang et al. 2020), where RHOV is likely to act as an oncogene (Chen et al. 2021).

For review, please refer to Faure and Fort 2015, and Hodge and Ridley 2020.

#### Edit history

| Date | Action | Author |
|---|---|---|
| 2017-07-25 | Created | Orlic-Milacic M |
| 2020-07-14 | Authored | Orlic-Milacic M |
| 2021-02-05 | Reviewed | Fort P |
| 2021-02-25 | Edited | Orlic-Milacic M |
| 2021-03-30 | Reviewed | Shepelev MV |
| 2021-04-15 | Edited | Orlic-Milacic M |
| 2021-05-22 | Modified | Shorser S |

#### Entities found in this pathway (3)

| Input | UniProt Id | Input | UniProt Id | Input | UniProt Id |
|---|---|---|---|---|---|
| NCK1 | O43639, P16333 | PAK1 | Q13153 | TPM4 | P67936 |

#### 18. Regulation of TNFR1 signaling (R-HSA-5357905)

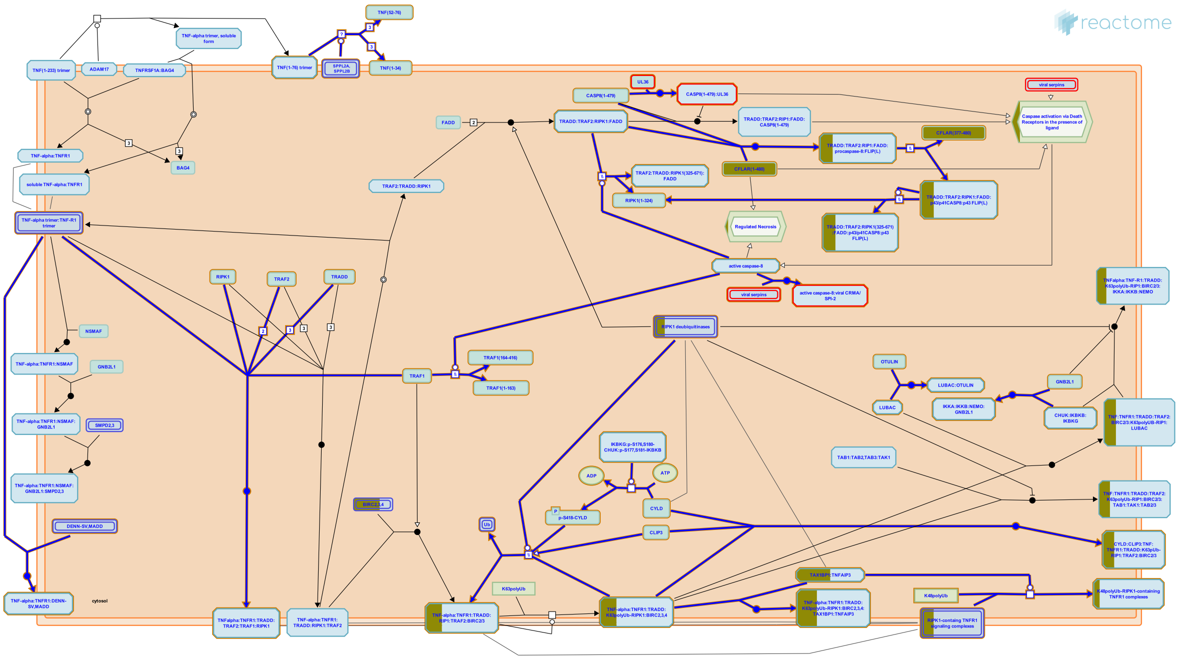

Tumor necrosis factor-alpha (TNFalpha) is an inflammatory cytokine, that activates either cell survival (e.g.,inflammation, proliferation) or cell death upon association with TNF receptor 1 (TNFR1). Stimuli and the cellular context dictate cell fate decisions between survival and death which rely on tightly regulated mechanisms with checkpoints on many levels. TNFR1-mediated NFkappaB activation leads to the pro-survival transcriptional program that is both anti-apoptotic and highly proinflammatory. The constitutive NFkappaB or AP1 activation may lead to excessive inflammation which has been associated with a variety of aggressive tumor types (Jackson-Bernitsas DG et al. 2007; Zhang JY et al. 2007). Thus, the tight regulation of TNFalpha:TNFR1 signaling is required to ensure the appropriate cell response to stimuli.


##### Edit history

| Date | Action | Author |
|---|---|---|
| 2014-03-26 | Created | Shamovsky V |
| 2015-02-15 | Edited | Shamovsky V |
| 2015-03-12 | Reviewed | Gillespie ME |
| 2015-05-12 | Authored | Shamovsky V |
| 2015-08-25 | Reviewed | Wajant H |
| 2021-05-31 | Modified | Shorser S |

#### Entities found in this pathway (3)

| Input | UniProt Id | Input | UniProt Id | Input | UniProt Id |
|---|---|---|---|---|---|
| BIRC3 | Q13489, Q13490 | CFLAR | O15519-1 | TNFAIP3 | P21580 |

#### 19. Neutrophil degranulation (R-HSA-6798695)

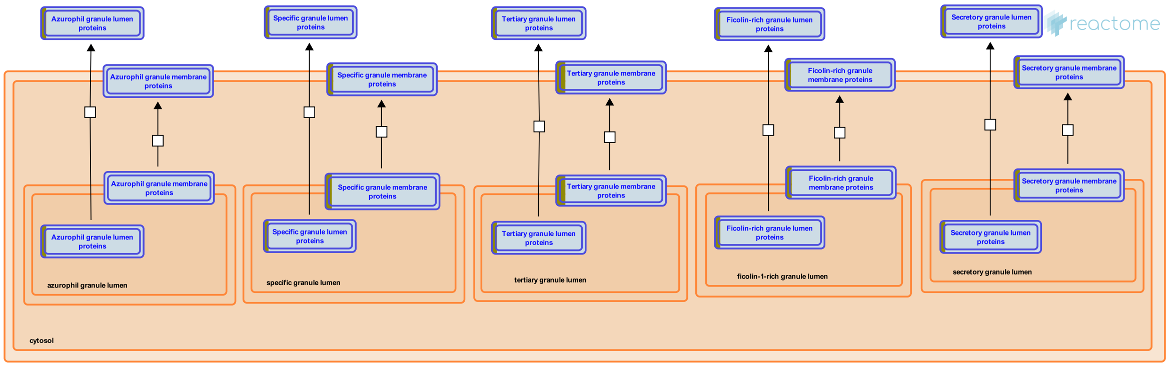


Neutrophils are the most abundant leukocytes (white blood cells), indispensable in defending the body against invading microorganisms. In response to infection, neutrophils leave the circulation and migrate towards the inflammatory focus. They contain several subsets of granules that are mobilized to fuse with the cell membrane or phagosomal membrane, resulting in the exocytosis or exposure of membrane proteins. Traditionally, neutrophil granule constituents are described as antimicrobial or proteolytic, but granules also introduce membrane proteins to the cell surface, changing how the neutrophil responds to its environment (Borregaard et al. 2007). Primed neutrophils actively secrete cytokines and other inflammatory mediators and can present antigens via MHC II, stimulating T-cells (Wright et al. 2010).

Granules form during neutrophil differentiation. Granule subtypes can be distinguished by their content but overlap in structure and composition. The differences are believed to be a consequence of changing protein expression and differential timing of granule formation during the terminal processes of neutrophil differentiation, rather than sorting (Le Cabec et al. 1996).

The classical granule subsets are Azurophil or primary granules (AG), secondary granules (SG) and gelatinase granules (GG). Neutrophils also contain exocytosable storage cell organelles, storage vesicles (SV), formed by endocytosis they contain many cell-surface markers and extracellular, plasma proteins (Borregaard et al. 1992). Ficolin-1-rich granules (FG) are like GGs highly exocytosable but gelatinase-poor (Rorvig et al. 2009).

#### Edit history

| Date | Action | Author |
|---|---|---|
| 2015-09-21 | Authored | Jupe S |
| 2015-09-21 | Created | Jupe S |
| 2016-06-13 | Edited | Jupe S |
| 2016-06-13 | Reviewed | Heegaard N |
| 2021-05-22 | Modified | Shorser S |

#### Entities found in this pathway (13)

| Input | UniProt Id | Input | UniProt Id | Input | UniProt Id |
|---|---|---|---|---|---|
| ADAM8 | P78325 | COTL1 | Q14019 | CTSZ | Q9UBR2 |
| CYBA | P13498 | DBNL | Q9UJU6 | ITGAM | P11215, P20702 |
| ITGAV | P06756 | LYZ | P61626 | PLAU | P00749 |
| PTAFR | P25105 | S100A11 | P31949 | S100A9 | P06702 |
| SERPINB1 | P30740 | | | | |

#### 20. WNT5:FZD7-mediated leishmania damping (R-HSA-9673324)

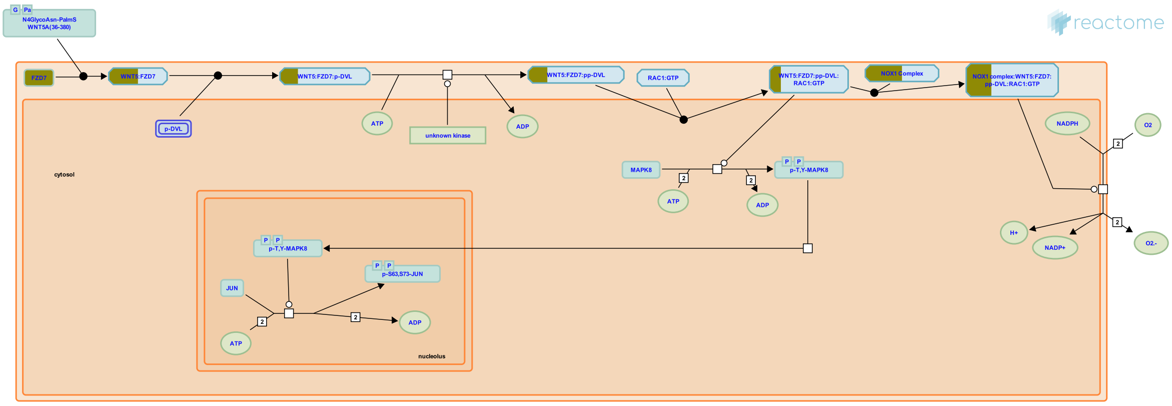


**Diseases:** cutaneous leishmaniasis.

Wnt-5a (WNT5) is known for being a highly specific regulated gene in response to microbial infection (Blumenthal et al. 2006, Pereira et al. 2008 & Ljungberg et al. 2019) including leishmaniasis (Chakraborty et al. 2017), where it seems to be involve in mechanisms that dampen the parasite load within main host macrophages (Chakraborty et al. 2017). In addition, WNT5 is a highly responsive gene in human macrophages present in chronic diseases such as rheumatoid arthritis (Sen et al. 2000), cancer (Pukrop et al. 2006), atherosclerosis (Christman et al. 2008) and obesity (Ouchi et al. 2010 & Ljungberg et al. 2019).

Frizzled-7 (FZD7) acts as a receptor of WNT5 which, upon binding, is implicated in the initiation of the non-canonical WNT pathway that ends up in the re-accommodation of the cytoskeleton to allow a process called planar cell polarity (PCP) (Ljungberg et al. 2019). The activation of the WNT5:FZD7 non-canonical signalling cascade that drives PCP is being studied for its involvement in inflammatory responses (Shao et al. 2016). Treatment of RAW264.7 macrophages with recombinant Wnt5a induced NADPH oxidase-mediated ROS production, which has been suggested to contribute to the macrophage control of L. donovani. Consequently, detailed understanding of how WNT signaling network defines host responses to infection could be important to identify potential targets (Ljungberg et al. 2019).

#### Edit history

| Date | Action | Author |
|---|---|---|
| 2020-01-07 | Created | Murillo JI |
| 2020-01-09 | Edited | Jassal B, Murillo JI |
| 2020-01-09 | Authored | Jassal B, Murillo JI |
| 2020-02-04 | Reviewed | Gregory DJ |
| 2020-02-06 | Modified | Murillo JI |

#### Entities found in this pathway (3)

| Input | UniProt Id | Input | UniProt Id | Input | UniProt Id |
|---|---|---|---|---|---|
| CYBA | P13498 | FZD7 | O75084 | NOXO1 | Q8NFA2 |

#### 21. Killing mechanisms (R-HSA-9664420)

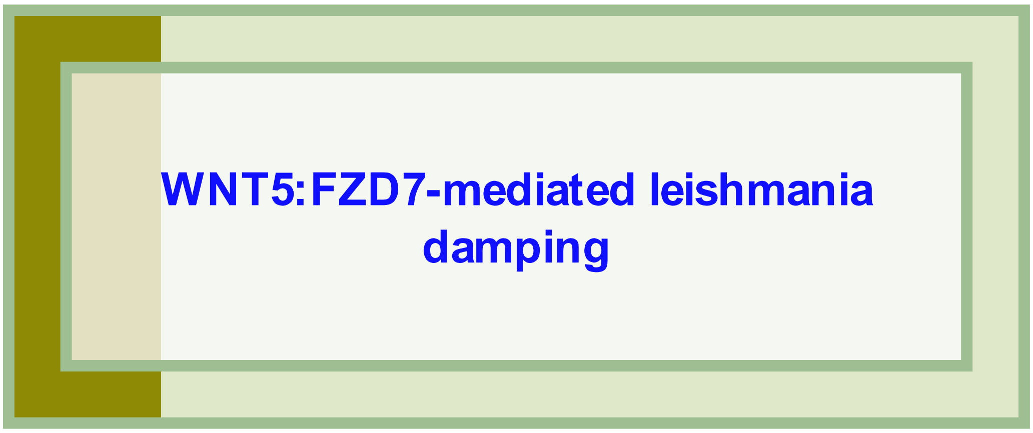


**Diseases:** cutaneous leishmaniasis.

The long-lasting Leishmania infection is established within macrophages in which the most effective killing response is the production of reactive oxygen species (ROS) and reactive nitrogen species (RNS). (Rossi and Fasel 2018). Additionally, autophagy has been described as an innate immune mechanism for eliminating intracellular pathogens, although its role in restricting Leishmania replication is unclear (Veras et al. 2019)

##### Edit history

| Date | Action | Author |
|---|---|---|
| 2019-10-22 | Created | Murillo JI |
| 2020-01-07 | Authored | Jassal B |
| 2020-01-29 | Authored | Murillo JI |
| 2020-02-04 | Reviewed | Gregory DJ |
| 2020-02-05 | Edited | Jassal B, Murillo JI |
| 2020-02-06 | Modified | Murillo JI |

#### Entities found in this pathway (3)

| Input | UniProt Id | Input | UniProt Id | Input | UniProt Id |
|---|---|---|---|---|---|
| CYBA | P13498 | FZD7 | O75084 | NOXO1 | Q8NFA2 |

#### 22. TCR signaling (R-HSA-202403)

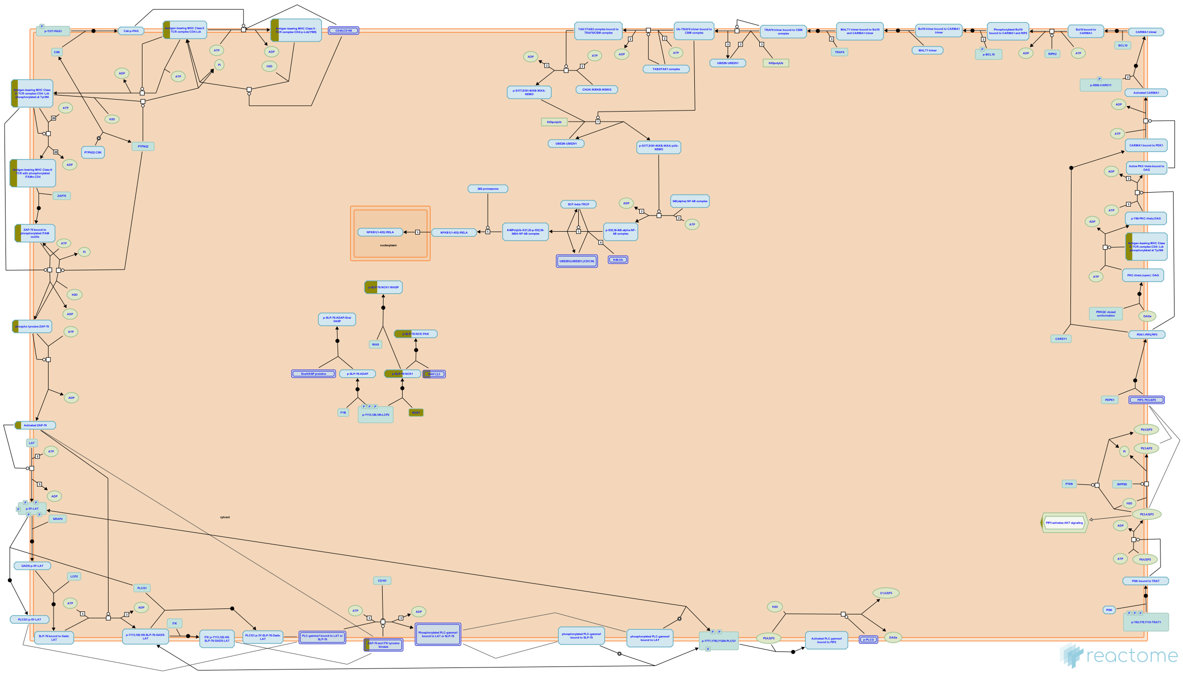

The TCR is a multisubunit complex that consists of clonotypic alpha/beta chains noncovalently associated with the invariant CD3 delta/epsilon/gamma and TCR zeta chains. T cell activation by antigen presenting cells (APCs) results in the activation of protein tyrosine kinases (PTKs) that associate with CD3 and TCR zeta subunits and the co-receptor CD4. Members of the Src kinases (Lck), Syk kinases (ZAP-70), Tec (Itk) and Csk families of nonreceptor PTKs play a crucial role in T cell activation. Activation of PTKs following TCR engagement results in the recruitment and tyrosine phosphorylation of enzymes such as phospholipase C gamma1 and Vav as well as critical adaptor proteins such as LAT, SLP-76 and Gads. These proximal activation leads to reorganization of the cytoskeleton as well as transcription activation of multiple genes leading to T lymphocyte proliferation, differentiation and/or effector function.


##### Edit history

| Date | Action | Author |
|---|---|---|
| 2007-10-29 | Created | Garapati P V |
| 2008-01-24 | Authored | Rudd C.E., Garapati P V, de Bono B |
| 2008-02-26 | Reviewed | Trowsdale J |
| 2021-05-22 | Modified | Shorser S |

##### Entities found in this pathway (6)

| Input | UniProt Id | Input | UniProt Id | Input | UniProt Id |
|---|---|---|---|---|---|
| HLA-DPA1 | P20036 | HLA-DPB1 | P04440 | HLA-DQB1 | P01920 |
| HLA-DRA | P01903, P01909 | NCK1 | P16333 | PAK1 | Q13153 |

#### 23. RHOU GTPase cycle (R-HSA-9013420)

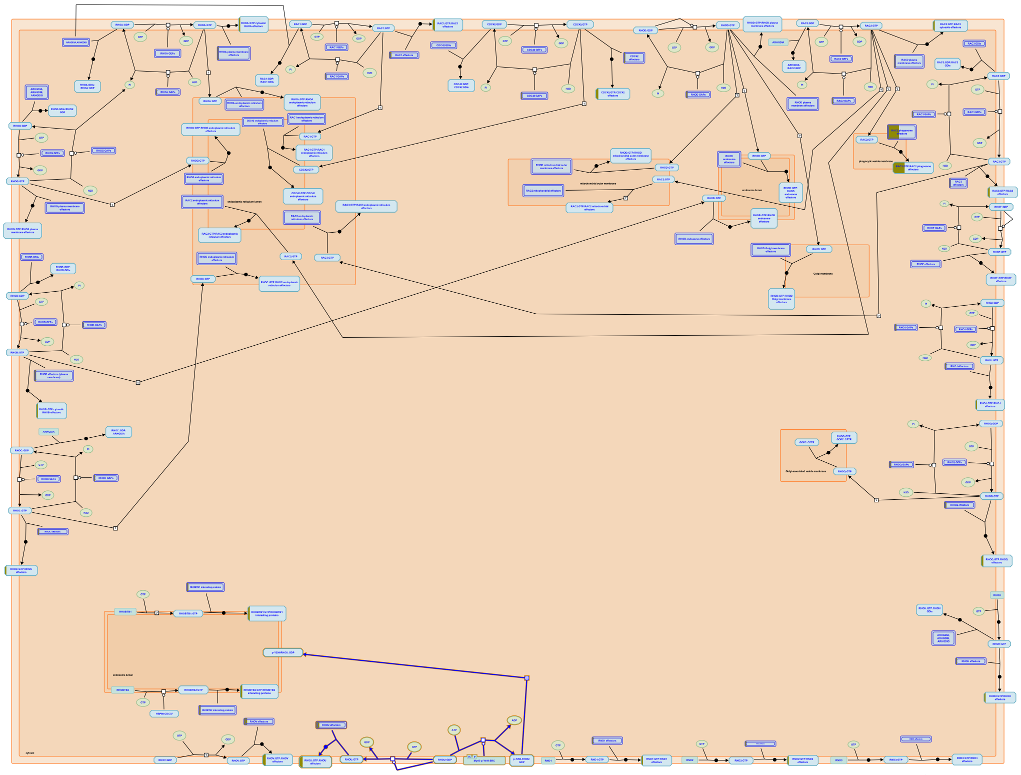

RHO GTPase RHOU (Wrch-1) possesses a high intrinsic guanine nucleotide exchange activity and is constitutively present in the active GTP-bound state in the absence of guanine nucleotide exchange factors (GEFs) (Shutes et al. 2004, Saras et al. 2004). RHOU does not possess a GTPase activity (Saras et al. 2004). RHOU has been reported to interact with some GTPase activator proteins (GAPs) (Bagci et al. 2020), which may serve as effectors that enable cross-talk with other RHO GTPases. RHOU was shown to regulate cytoskeletal dynamics, cell migration and adhesion. RHOU is expressed during embryonic development and regulates cardiac (Dickover et al. 2014) and intestinal (Slaymi et al. 2019) development. RHOU activates JNK and AKT signaling during cell migration (Chuang et al. 2007).

For review, please refer to Faure and Fort 2015, and Hodge and Ridley 2020.

#### Edit history

| Date | Action | Author |
|---|---|---|
| 2017-07-25 | Created | Orlic-Milacic M |
| 2020-07-14 | Authored | Orlic-Milacic M |
| 2021-02-05 | Reviewed | Fort P |
| 2021-02-25 | Edited | Orlic-Milacic M |
| 2021-03-30 | Reviewed | Shepelev MV |
| 2021-04-12 | Revised | Orlic-Milacic M |
| 2021-04-15 | Edited | Orlic-Milacic M |
| 2021-05-22 | Modified | Shorser S |

#### Entities found in this pathway (3)

| Input | UniProt Id | Input | UniProt Id | Input | UniProt Id |
|---|---|---|---|---|---|
| MYO6 | Q9UM54 | NCK1 | O43639, P16333 | PAK1 | Q13153 |

### 24. MHC class II antigen presentation (R-HSA-2132295)

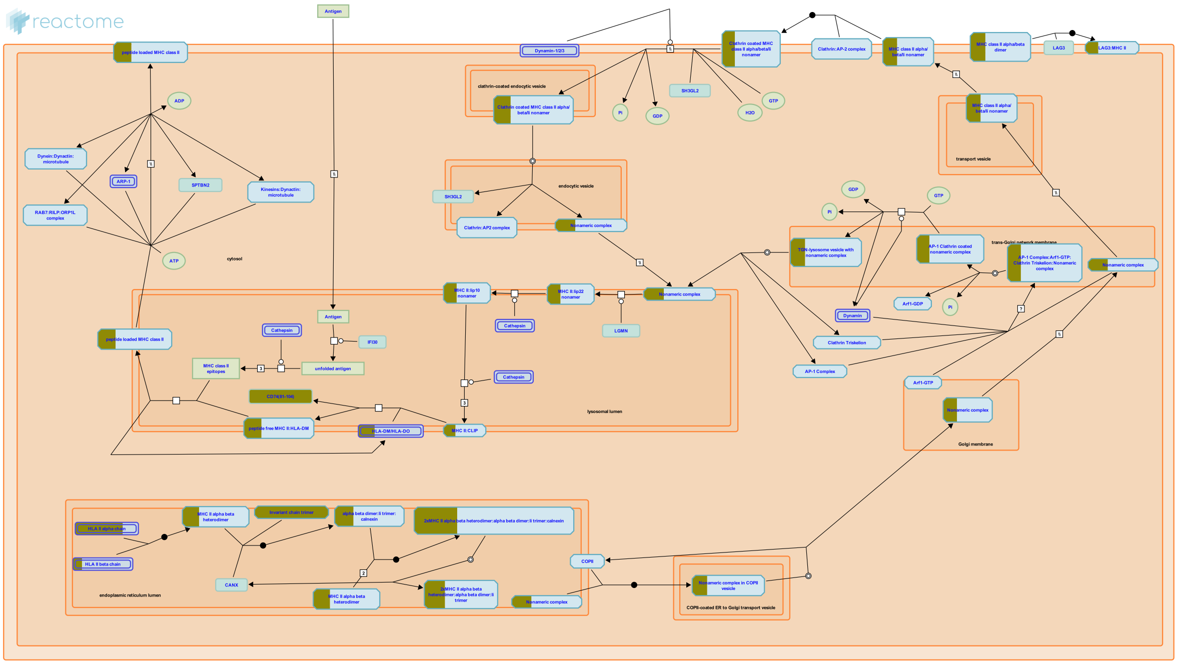

Antigen presenting cells (APCs) such as B cells, dendritic cells (DCs) and monocytes/macrophages express major histocompatibility complex class II molecules (MHC II) at their surface and present exogenous antigenic peptides to CD4+ T helper cells. CD4+ T cells play a central role in immune protection. On their activation they stimulate differentiation of B cells into antibody-producing B-cell blasts and initiate adaptive immune responses. MHC class II molecules are transmembrane glycoprotein heterodimers of alpha and beta subunits. Newly synthesized MHC II molecules present in the endoplasmic reticulum bind to a chaperone protein called invariant (Ii) chain. The binding of Ii prevents the premature binding of self antigens to the nascent MHC molecules in the ER and also guides MHC molecules to endocytic compartments. In the acidic endosomal environment, Ii is degraded in a stepwise manner, ultimately to free the class II peptide-binding groove for loading of antigenic peptides. Exogenous antigens are internalized by the APC by receptor mediated endocytosis, phagocytosis or pinocytosis into endocytic compartments of MHC class II positive cells, where engulfed antigens are degraded in a low pH environment by multiple acidic proteases, generating MHC class II epitopes. Antigenic peptides are then loaded into the class II ligand-binding groove. The resulting class II peptide complexes then move to the cell surface, where they are scanned by CD4+ T cells for specific recognition (Berger & Roche 2009, Zhou & Blum 2004, Watts 2004, Landsverk et al. 2009).

#### Edit history

| Date | Action | Author |
|---|---|---|
| 2012-02-21 | Edited | Garapati P V |
| 2012-02-21 | Authored | Garapati P V |
| 2012-02-21 | Created | Garapati P V |
| 2012-05-14 | Reviewed | Neefjes J |
| 2021-05-22 | Modified | Shorser S |

#### Entities found in this pathway (6)

| Input | UniProt Id | Input | UniProt Id | Input | UniProt Id |
|---|---|---|---|---|---|
| CD74 | P04233 | HLA-DMA | P28067 | HLA-DPA1 | P20036 |
| HLA-DPB1 | P04440 | HLA-DQB1 | P01920 | HLA-DRA | P01903, P01909 |

### 25. RIPK1-mediated regulated necrosis (R-HSA-5213460)

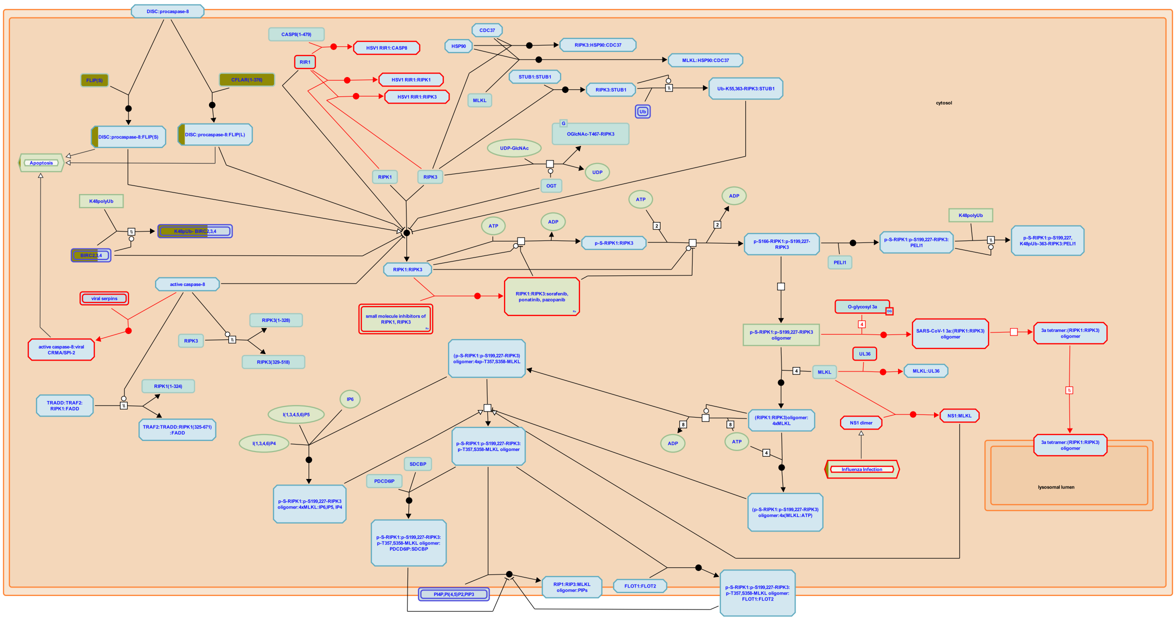

Receptor-interacting serine/threonine-kinase protein 1 (RIPK1) and RIPK3-dependent necrosis is called necroptosis or programmed necrosis. The kinase activities of RIPK1 and RIPK3 are essential for the necroptotic cell death in human, mouse cell lines and genetic mice models (Cho YS et al. 2009; He S et al. 2009, 2011; Zhang DW et al. 2009; McQuade T et al. 2013; Newton et al. 2014). The initiation of necroptosis can be stimulated by the same death ligands that activate extrinsic apoptotic signaling pathway, such as tumor necrosis factor (TNF) alpha, Fas ligand (FasL), and TRAIL (TNF-related apoptosis-inducing ligand) or toll like receptors 3 and 4 ligands (Holler N et al. 2000; He S et al. 2009; Feoktistova M et al. 2011; Voigt S et al. 2014). In contrast to apoptosis, necroptosis represents a form of cell death that is optimally induced when caspases are inhibited (Holler N et al. 2000; Hopkins-Donaldson S et al. 2000; Sawai H 2014). Specific inhibitors of caspase-independent necrosis, necrostatins, have recently been identified (Degterev A et al. 2005, 2008). Necrostatins have been shown to inhibit the kinase activity of RIPK1 (Degterev A et al. 2008). Importantly, cell death of apoptotic morphology can be shifted to a necrotic phenotype when caspase 8 activity is compromised, otherwise active caspase 8 blocks necroptosis by the proteolytic cleavage of RIPK1 and RIPK3 (Kalai M et al. 2002; Degterev A et al. 2008; Lin Y et al. 1999; Feng S et al. 2007). When caspase activity is inhibited under certain pathophysiological conditions or by pharmacological agents, deubiquitinated RIPK1 is engaged in physical and functional interactions with the cognate kinase RIPK3 leading to formation of necrosome, a necroptosis-inducing complex consisting of RIPK1 and RIPK3 (Sawai H 2013; Moquin DM et al. 2013; Kalai M et al. 2002; Cho YS et al. 2009, He S et al. 2009, Zhang DW et al. 2009). Within the necrosome RIPK1 and RIPK3 bind to each other through their RIP homotypic interaction motif (RHIM) domains. The RHIMs can facilitate RIPK1:RIPK3 oligomerization, allowing them to form amyloid-like fibrillar structures (Li J et al. 2012; Mompean M et al. 2018). RIPK3 in turn interacts with mixed lineage kinase domain-like protein (MLKL) (Sun L et al. 2012; Zhao J et al. 2012; Murphy JM et al. 2013; Chen W et al. 2013). The precise mechanism of MLKL activation by RIPK3 is incompletely understood and may vary across species (Davies KA et al. 2020). Mouse MLKL activation relies on transient engagement of RIPK3 to facilitate phosphorylation of the pseudokinase domain (Murphy JM et al. 2013; Petrie EJ et al. 2019a), while it appears that stable recruitment of human MLKL by necrosomal RIPK3 is an additional crucial step in human MLKL activation (Davies KA et al. 2018; Petrie EJ et al. 2018, 2019b). RIPK3-mediated phosphorylation is thought to initiate MLKL oligomerization, membrane translocation and membrane disruption (Sun L et al. 2012; Wang H et al. 2014; Petrie EJ et al. 2020; Samson AL et al. 2020). Studies in human cell lines suggest that upon induction of necroptosis MLKL shifts to the plasma membrane and membranous organelles such as mitochondria, lysosome, endosome and ER (Wang H et al. 2014), but it is trafficking via a Golgi-microtubule-actin-dependent mechanism that facilitates plasma membrane translocation, where membrane disruption causes death (Samson AL et al. 2020). The mechanisms of necroptosis regulation and execution downstream of MLKL remain elusive. The precise oligomeric form of MLKL that mediates plasma membrane disruption has been highly debated (Cai Z et al. 2014; Chen X et al. 2014; Dondelinger Y et al. 2014; Wang H et al. 2014; Petrie EJ et al. 2017, 2018; Samson AL et al. 2020 ). However, microscopy data revealed that MLKL assembles into higher molecular weight species upon cytoplasmic necrosomes within human cells, and upon phosphorylation by RIPK3, MLKL is trafficked to the plasma membrane (Samson AL et al. 2020). At the plasma membrane, phospho-MLKL forms heterogeneous higher order assemblies, which are thought to permeabilize cells, leading to release of DAMPs to invoke inflammatory responses. While RIPK1, RIPK3 and MLKL are the core signaling components in the necroptosis pathway, many additional molecules have been proposed to positively and negatively tune the signaling pathway. Currently, this picture is evolving rapidly as new modulators continue to be discovered.

The Reactome module describes MLKL-mediated necroptotic events on the plasma membrane.


#### Edit history

| Date | Action | Author |
|---|---|---|
| 2013-12-14 | Created | Shamovsky V |
| 2013-12-20 | Authored | Shamovsky V |
| 2014-10-31 | Reviewed | Gillespie ME |
| 2015-02-10 | Edited | Shamovsky V |
| 2015-02-15 | Reviewed | Chan FK |
| 2020-08-28 | Reviewed | Murphy JM |
| 2021-05-31 | Modified | Shorser S |

#### Entities found in this pathway (2)

| Input | UniProt Id | Input | UniProt Id |
|---|---|---|---|
| BIRC3 | Q13489, Q13490 | CFLAR | O15519-1, O15519-2 |

### 6. Identifiers found

Below is a list of the input identifiers that have been found or mapped to an equivalent element in Reactome, classified by resource.

#### Entities (102)

| Input | UniProt Id | Input | UniProt Id | Input | UniProt Id |
|---|---|---|---|---|---|
| ACSM3 | Q68CK6 | ADAM8 | P78325 | ANO9 | A1A5B4 |
| APOC1 | P02654 | AQP1 | P29972 | ARHGAP26 | Q9UNA1 |
| ARL6IP5 | O75915 | BAD | Q92934 | BIRC3 | Q13489, Q13490 |
| BTN2A2 | Q8WVV5 | CCL23 | P55773-2 | CCL25 | O15444 |
| CD74 | P04233 | CD84 | Q9UIB8 | CDC42EP4 | Q9H3Q1 |
| CFLAR | O15519-1, O15519-2 | CLDN3 | O15551 | CLDN4 | O14493 |
| COTL1 | Q14019 | CSF2RB | P24394, P32927 | CTSZ | Q9UBR2 |
| CYBA | P13498 | DAPP1 | Q9UN19 | DBI | P07108 |
| DBNL | Q9UJU6 | DSC2 | Q02487 | EIF4G2 | P78344, Q04637 |
| ENPP2 | Q13822 | FADS1 | O60427 | FADS2 | O95864 |
| FAR2 | Q96K12 | FNBP1 | Q96RU3 | FZD7 | O75084 |
| GALNT18 | Q6P9A2 | GFRA1 | P56159 | GJB3 | O75712 |
| GJB4 | Q9NTQ9 | GOSR1 | O95249 | GP2 | P55259 |
| HDGF | P51858 | HIPK2 | Q9H2X6 | HLA-DMA | P28067 |
| HLA-DPA1 | P20036 | HLA-DPB1 | P04440 | HLA-DQB1 | P01920 |
| HLA-DRA | P01903, P01909 | HS3ST1 | O14792 | HTR2B | P41595 |
| ICAM2 | P13598 | IL23A | Q9NPF7 | IL2RG | P31785 |
| ITGAM | P11215, P20702 | ITGAV | P06756 | KCNE3 | Q9Y6H6 |
| LTB | Q06643 | LYZ | P61626 | MAP1LC3A | Q9H492 |
| MYO6 | Q9UM54 | NCALD | P61601 | NCEH1 | Q6PIU2 |
| NCF1 | P14598 | NCK1 | P16333 | NECAP2 | Q9NVZ3 |
| NFKB2 | Q00653 | NIPAL3 | Q6P499 | NLGN4Y | Q8NFZ3 |
| NOXO1 | Q8NFA2 | NPB | Q8NG41 | NPPC | P23582 |
| PAK1 | Q13153 | PFKFB3 | Q16875 | PIK3AP1 | Q6ZUJ8 |
| PLAU | P00749 | POLD1 | P28340 | PON3 | Q15166 |
| PTAFR | P25105 | RB1 | P06400 | RELB | Q01201 |
| RIN3 | Q8TB24 | RNF181 | Q9P0P0 | RPL23 | P62829 |
| RPS19BP1 | Q86WX3 | RRAS | P62070 | S100A11 | P31949 |
| S100A9 | P06702 | SEMA5A | Q13591 | SERPINB1 | P30740 |
| SERPINF2 | P08697 | SLC2A6 | Q9UGQ3 | SLC9A9 | Q8IVB4 |
| SLCO2B1 | O94956 | SOCS1 | O15524 | ST5 | P78524 |
| TCF12 | Q99081 | TMSB4X | P62328 | TNFAIP3 | P21580 |
| TNFRSF11A | Q9Y6Q6 | TNFRSF11B | O00300 | TPM4 | P67936 |
| TYMP | P19971 | UPP1 | Q16831 | YWHAH | Q04917 |

| Input | Ensembl Id | Input | Ensembl Id | Input | Ensembl Id |
|---|---|---|---|---|---|
| APOC1 | ENSG00000130208 | ENPP2 | ENST00000427067 | FADS1 | ENSG00000149485 |
| HDGF | ENSG00000143321 | HLA-DPA1 | ENSG00000231389 | HLA-DPB1 | ENSG00000223865 |
| HLA-DQB1 | ENSG00000179344 | HLA-DRA | ENSG00000204287 | IL23A | ENST00000619177 |
| IL2RG | ENSG00000147168 | ITGAM | ENSG00000169896 | PTAFR | ENSG00000169403 |
| SOCS1 | ENSG00000185338 | | | | |

### 7. Identifiers not found

These 43 identifiers were not found neither mapped to any entity in Reactome.

| | | | | | | | |
|---|---|---|---|---|---|---|---|
| ABRACL | AC009041.2 | AC026471.3 | AC239800.2 | ADGRL2 | ANKRD29 | ASPHD1 | BCL3 |
| BRI3BP | BTG3 | C2CD2 | CALHM6 | CCL15 | CLPTM1 | CSRP2 | EXOC3L4 |
| FCGRT | FERMT1 | KIAA1147 | LAD1 | LINC01559 | LINC02384 | MIA | OGFRL1 |
| PARD3B | PNRC1 | PRSS36 | RASSF2 | RNF145 | SEPT11 | SH3BGRL | SNX10 |
| SOX8 | SPART | SUB1 | SULF2 | SYNPO2 | TACSTD2 | TM7SF3 | TSPAN8 |
| WEE2 | XG | ZNF593 | | | | | |
