## Supplementary material for "A proximal-to-distal survey of healthy adult human small intestine and colon epithelium by single-cell transcriptomics": File S4

reactome

### Pathway Analysis Report

This report contains the pathway analysis results for the submitted sample ''. Analysis was performed against Reactome version 77 on 24/08/2021. The web link to these results is:

https://reactome.org/PathwayBrowser/#/ANALYSIS=MjAyMTA4MjQxODI0MjBfNTY5

Please keep in mind that analysis results are temporarily stored on our server. The storage period depends on usage of the service but is at least 7 days. As a result, please note that this URL is only valid for a limited time period and it might have expired.

### Table of Contents

1. Introduction
2. Properties
3. Genome-wide overview
4. Most significant pathways
5. Pathways details
6. Identifiers found
7. Identifiers not found

### 1. Introduction

Reactome is a curated database of pathways and reactions in human biology. Reactions can be considered as pathway 'steps'. Reactome defines a 'reaction' as any event in biology that changes the state of a biological molecule. Binding, activation, translocation, degradation and classical biochemical events involving a catalyst are all reactions. Information in the database is authored by expert biologists, entered and maintained by Reactome’s team of curators and editorial staff. Reactome content frequently cross-references other resources e.g. NCBI, Ensembl, UniProt, KEGG (Gene and Compound), ChEBI, PubMed and GO. Orthologous reactions inferred from annotation for Homo sapiens are available for 17 non-human species including mouse, rat, chicken, puffer fish, worm, fly, yeast, rice, and Arabidopsis. Pathways are represented by simple diagrams following an SBGN-like format.

Reactome's annotated data describe reactions possible if all annotated proteins and small molecules were present and active simultaneously in a cell. By overlaying an experimental dataset on these annotations, a user can perform a pathway over-representation analysis. By overlaying quantitative expression data or time series, a user can visualize the extent of change in affected pathways and its progression. A binomial test is used to calculate the probability shown for each result, and the p-values are corrected for the multiple testing (Benjamini–Hochberg procedure) that arises from evaluating the submitted list of identifiers against every pathway.

To learn more about our Pathway Analysis, please have a look at our relevant publications:

Fabregat A, Sidiropoulos K, Garapati P, Gillespie M, Hausmann K, Haw R, … D’Eustachio P (2016). The reactome pathway knowledgebase. Nucleic Acids Research, 44(D1), D481–D487. https://doi.org/10.1093/nar/gkv1351.

Fabregat A, Sidiropoulos K, Viteri G, Forner O, Marin-Garcia P, Arnau V, … Hermjakob H (2017). Reactome pathway analysis: a high-performance in-memory approach. BMC Bioinformatics, 18.

#### 2. Properties

- This is an **overrepresentation** analysis: A statistical (hypergeometric distribution) test that determines whether certain Reactome pathways are over-represented (enriched) in the submitted data. It answers the question ‘Does my list contain more proteins for pathway X than would be expected by chance?’ This test produces a probability score, which is corrected for false discovery rate using the Benjamani-Hochberg method.
- 73 out of 118 identifiers in the sample were found in Reactome, where 261 pathways were hit by at least one of them.
- All non-human identifiers have been converted to their human equivalent.
- This report is filtered to show only results for species 'Homo sapiens' and resource 'all resources'.
- The unique ID for this analysis (token) is MjAyMTA4MjQxODI0MjBfNTY5. This ID is valid for at least 7 days in Reactome’s server. Use it to access Reactome services with your data.

### 3. Genome-wide overview

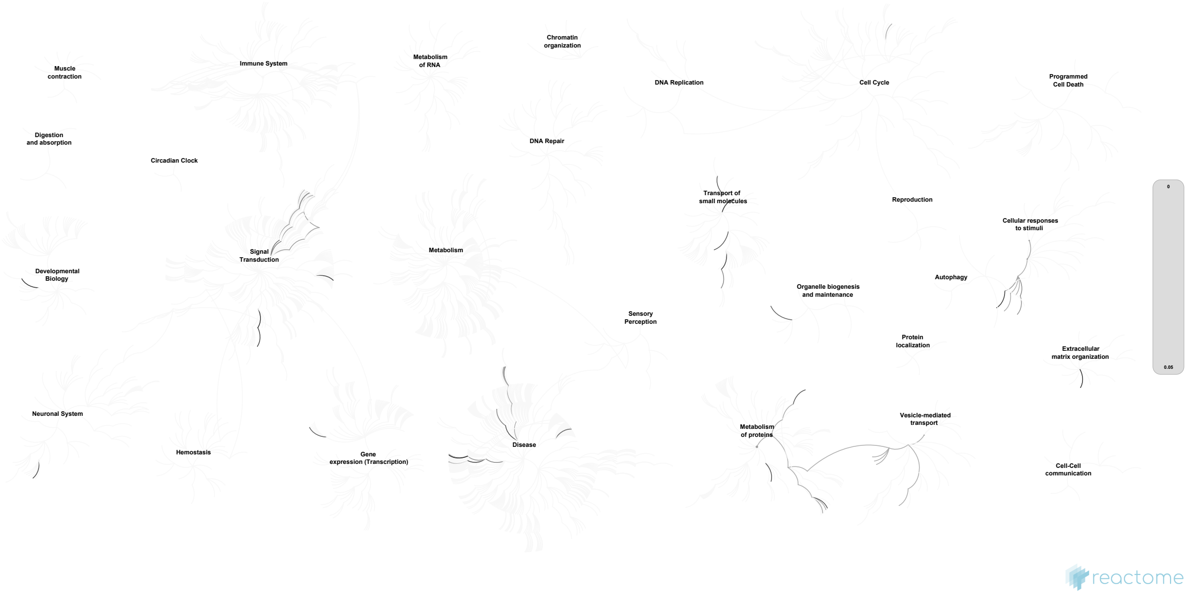


This figure shows a genome-wide overview of the results of your pathway analysis. Reactome pathways are arranged in a hierarchy. The center of each of the circular "bursts" is the root of one top-level pathway, for example "DNA Repair". Each step away from the center represents the next level lower in the pathway hierarchy. The color code denotes over-representation of that pathway in your input dataset. Light grey signifies pathways which are not significantly over-represented.

#### 4. Most significant pathways

The following table shows the 25 most relevant pathways sorted by p-value.

| Pathway name | Entities | | | | Reactions | |
|---|---|---|---|---|---|---|
| | found | ratio | p-value | FDR* | found | ratio |
| Unfolded Protein Response (UPR) | 19 / 155 | 0.011 | 2.33e-15 | 6.23e-13 | 70 / 94 | 0.007 |
| Asparagine N-linked glycosylation | 27 / 421 | 0.029 | 1.38e-14 | 1.83e-12 | 39 / 144 | 0.011 |
| ER to Golgi Anterograde Transport | 16 / 164 | 0.011 | 1.21e-11 | 7.96e-10 | 27 / 39 | 0.003 |
| Transport to the Golgi and subsequent modification | 17 / 219 | 0.015 | 8.90e-11 | 4.72e-09 | 29 / 60 | 0.004 |
| IRE1alpha activates chaperones | 12 / 101 | 0.007 | 5.60e-10 | 2.46e-08 | 52 / 53 | 0.004 |
| COPI-mediated anterograde transport | 11 / 107 | 0.007 | 1.30e-08 | 4.31e-07 | 10 / 12 | 8.88e-04 |
| XBP1(S) activates chaperone genes | 9 / 95 | 0.007 | 5.58e-07 | 1.62e-05 | 47 / 47 | 0.003 |
| Signaling by FGFR2 IIIa TM | 5 / 24 | 0.002 | 4.87e-06 | 1.16e-04 | 2 / 2 | 1.48e-04 |
| Post-translational protein modification | 35 / 1,598 | 0.11 | 4.92e-06 | 1.16e-04 | 60 / 526 | 0.039 |
| FGFR2 mutant receptor activation | 6 / 43 | 0.003 | 5.25e-06 | 1.16e-04 | 17 / 18 | 0.001 |
| Synthesis of substrates in N-glycan biosythesis | 9 / 131 | 0.009 | 7.45e-06 | 1.49e-04 | 9 / 49 | 0.004 |
| Signaling by FGFR2 in disease | 6 / 58 | 0.004 | 2.80e-05 | 5.33e-04 | 27 / 28 | 0.002 |
| Biosynthesis of the N-glycan precursor (dolichol lipid-linked oligosaccharide, LLO) and transfer to a nascent protein | 9 / 159 | 0.011 | 3.37e-05 | 5.52e-04 | 9 / 63 | 0.005 |
| Membrane Trafficking | 19 / 665 | 0.046 | 3.45e-05 | 5.52e-04 | 47 / 219 | 0.016 |
| O-linked glycosylation of mucins | 6 / 73 | 0.005 | 9.87e-05 | 0.001 | 17 / 17 | 0.001 |
| COPI-dependent Golgi-to-ER retrograde traffic | 7 / 107 | 0.007 | 1.07e-04 | 0.001 | 6 / 11 | 8.14e-04 |
| CREB3 factors activate genes | 3 / 9 | 6.19e-04 | 1.08e-04 | 0.002 | 13 / 21 | 0.002 |
| Phospholipase C-mediated cascade; FGFR2 | 4 / 25 | 0.002 | 1.24e-04 | 0.002 | 3 / 3 | 2.22e-04 |
| COPII-mediated vesicle transport | 6 / 77 | 0.005 | 1.32e-04 | 0.002 | 11 / 16 | 0.001 |
| FGFR2 ligand binding and activation | 4 / 26 | 0.002 | 1.44e-04 | 0.002 | 4 / 5 | 3.70e-04 |
| Signaling by FGFR in disease | 6 / 82 | 0.006 | 1.84e-04 | 0.002 | 27 / 106 | 0.008 |
| Synthesis of GDP-mannose | 3 / 11 | 7.56e-04 | 1.95e-04 | 0.002 | 2 / 3 | 2.22e-04 |
| PI-3K cascade:FGFR2 | 4 / 31 | 0.002 | 2.80e-04 | 0.003 | 6 / 7 | 5.18e-04 |
| Metabolism of proteins | 38 / 2,205 | 0.152 | 3.50e-04 | 0.003 | 65 / 798 | 0.059 |
| SHC-mediated cascade:FGFR2 | 4 / 33 | 0.002 | 3.54e-04 | 0.003 | 4 / 4 | 2.96e-04 |

* False Discovery Rate

### 5. Pathways details

For every pathway of the most significant pathways, we present its diagram, as well as a short summary, its bibliography and the list of inputs found in it.

#### 1. Unfolded Protein Response (UPR) (R-HSA-381119)

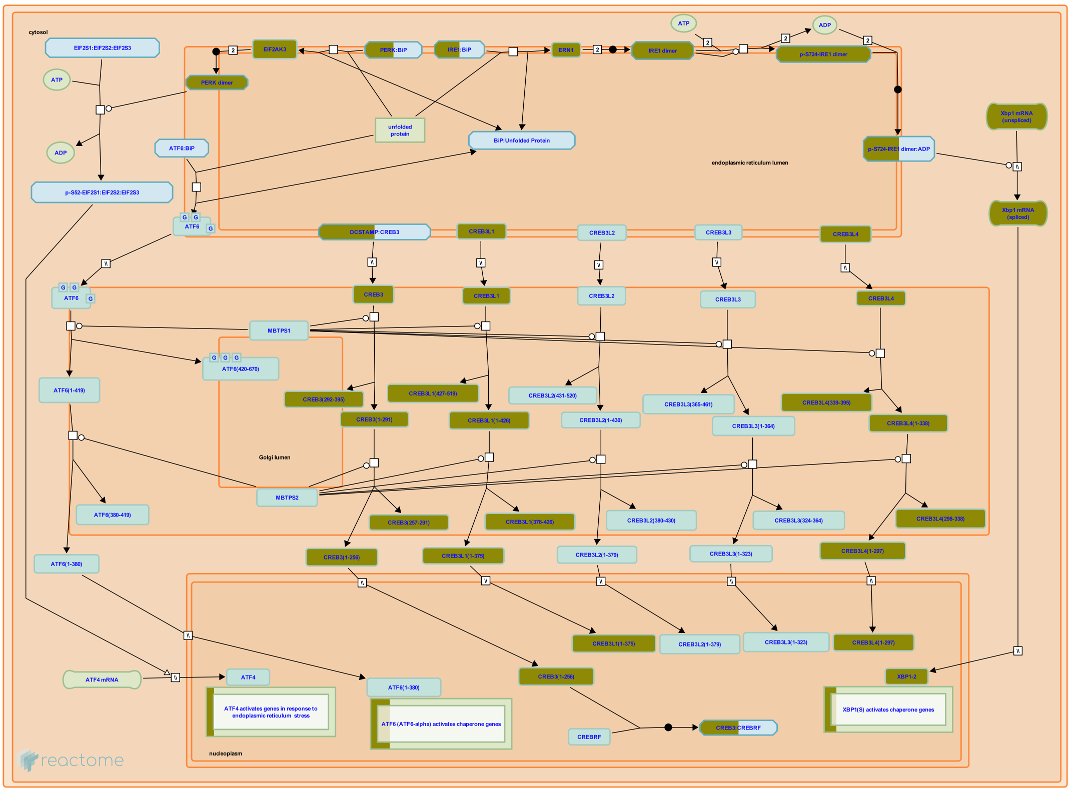


**Cellular compartments:** Golgi membrane, cytosol, endoplasmic reticulum lumen, endoplasmic reticulum membrane, nucleoplasm.

The Unfolded Protein Response (UPR) is a regulatory system that protects the Endoplasmic Reticulum (ER) from overload. The UPR is provoked by the accumulation of improperly folded protein in the ER during times of unusually high secretion activity. Analysis of mutants with altered UPR, however, shows that the UPR is also required for normal development and function of secretory cells.

One level at which the URP operates is transcriptional and translational regulation: mobilization of ATF6, ATF6B, CREB3 factors and IRE1 leads to increased transcription of genes encoding chaperones, while mobilization of PERK (pancreatic eIF2alpha kinase, EIF2AK3) leads to phosphorylation of the translation initiation factor eIF2alpha and global down-regulation of protein synthesis.

ATF6, ATF6B, and CREB3 factors (CREB3 (LUMAN), CREB3L1 (OASIS), CREB3L2 (BBF2H7, Tisp40), CREB3L3 (CREB-H), and CREB3L4 (CREB4)) are membrane-bound transcription activators that respond to ER stress by transiting from the ER membrane to the Golgi membrane where their transmembrane domains are cleaved, releasing their cytosolic domains to transit to the nucleus and activate transcription of target genes. IRE1, also a resident of the ER membrane, dimerizes and autophosphorylates in response to ER stress. The activated IRE1 then catalyzes unconventional splicing of XBP1 mRNA to yield an XBP1 isoform that is targeted to the nucleus and activates chaperone genes.


#### Edit history

| Date | Action | Author |
|---|---|---|
| 2008-11-19 | Created | May B |
| 2008-11-20 | Edited | May B, Gopinathrao G |
| 2008-12-02 | Reviewed | Matthews L, D'Eustachio P, Gillespie ME |
| 2009-06-02 | Authored | May B |
| 2010-04-30 | Reviewed | Urano F |
| 2021-05-22 | Modified | Shorser S |

#### Entities found in this pathway (10)

| Input | UniProt Id | Input | UniProt Id | Input | UniProt Id |
|---|---|---|---|---|---|
| ARFGAP1 | Q8N6T3 | CREB3L1 | Q96BA8 | CREB3L4 | O43889, Q8TEY5 |
| EIF2AK3 | Q9NZJ5 | ERN2 | O75460 | GFPT1 | Q06210 |
| HERPUD1 | Q15011 | KDELR3 | O43731 | PDIA5 | Q14554 |
| XBP1 | P17861-2 | | | | |

| Input | Ensembl Id | Input | Ensembl Id | Input | Ensembl Id |
|---|---|---|---|---|---|
| ARFGAP1 | ENSG00000101199 | GFPT1 | ENSG00000198380 | HERPUD1 | ENSG00000051108 |
| KDELR3 | ENSG00000100196 | PDIA5 | ENSG00000065485 | XBP1 | ENSG00000100219, ENST00000216037, ENST00000344347 |

#### 2. Asparagine N-linked glycosylation (R-HSA-446203)

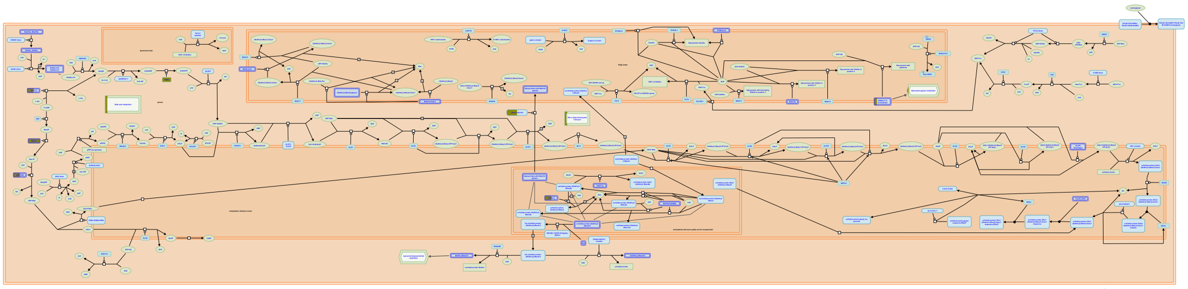

N-linked glycosylation is the most important form of post-translational modification for proteins synthesized and folded in the Endoplasmic Reticulum (Stanley et al. 2009). An early study in 1999 revealed that about 50% of the proteins in the Swiss-Prot database at the time were N-glycosylated (Apweiler et al. 1999). It is now established that the majority of the proteins in the secretory pathway require glycosylation in order to achieve proper folding.

The addition of an N-glycan to a protein can have several roles (Shental-Bechor & Levy 2009). First, glycans enhance the solubility and stability of the proteins in the ER, the golgi and on the outside of the cell membrane, where the composition of the medium is strongly hydrophilic and where proteins, that are mostly hydrophobic, have difficulty folding properly. Second, N-glycans are used as signal molecules during the folding and transport process of the protein: they have the role of labels to determine when a protein must interact with a chaperon, be transported to the golgi, or targeted for degradation in case of major folding defects. Third, and most importantly, N-glycans on completely folded proteins are involved in a wide range of processes: they help determine the specificity of membrane receptors in innate immunity or in cell-to-cell interactions, they can change the properties of hormones and secreted proteins, or of the proteins in the vesicular system inside the cell.

All N-linked glycans are derived from a common 14-sugar oligosaccharide synthesized in the ER, which is attached co-translationally to a protein while this is being translated inside the reticulum. The process of the synthesis of this glycan, known as Synthesis of the N-glycan precursor or LLO, constitutes one of the most conserved pathways in eukaryotes, and has been also observed in some eubacteria. The attachment usually happens on an asparagine residue within the consensus sequence asparagine-X-threonine by an complex called oligosaccharyl transferase (OST).

After being attached to an unfolded protein, the glycan is used as a label molecule in the folding process (also known as Calnexin/Calreticulin cycle) (Lederkremer 2009). The majority of the glycoproteins in the ER require at least one glycosylated residue in order to achieve proper folding, even if it has been shown that a smaller portion of the proteins in the ER can be folded without this modification.

Once the glycoprotein has achieved proper folding, it is transported via the cis-Golgi through all the Golgi compartments, where the glycan is further modified according to the properties of the glycoprotein. This process involves relatively few enzymes but due to its combinatorial nature, can lead to several millions of different possible modifications. The exact topography of this network of reactions has not been established yet, representing one of the major challenges after the sequencing of the human genome (Hossler et al. 2006).

Since N-glycosylation is involved in an great number of different processes, from cell-cell interaction to folding control, mutations in one of the genes involved in glycan assembly and/or modification can lead to severe development problems (often affecting the central nervous system). All the diseases in genes involved in glycosylation are collectively known as Congenital Disorders of Glycosylation (CDG) (Sparks et al. 2003), and classified as CDG type I for the genes in the LLO synthesis pathway, and CDG type II for the others.


#### Edit history

| Date | Action | Author |
|---|---|---|
| 2009-11-10 | Edited | Jassal B |
| 2009-11-10 | Authored | Dall'Olio GM |
| 2009-11-10 | Created | Jassal B |
| 2010-04-16 | Reviewed | Gagneux P |
| 2021-05-22 | Modified | Shorser S |

#### Entities found in this pathway (23)

| Input | UniProt Id | Input | UniProt Id | Input | UniProt Id |
|---|---|---|---|---|---|
| ARFGAP1 | Q8N6T3, Q9NP61 | B4GALT4 | O60513 | COG3 | Q96JB2 |
| COPB2 | P35606 | EDEM3 | Q9BZQ6 | GFPT1 | Q06210 |
| GMPPB | Q9Y5P6 | GNE | Q9Y223 | GOLGA2 | Q08379 |
| GOLGB1 | Q14789 | KDELR2 | P24390, P33947 | KDELR3 | O43731 |
| LMAN1 | P49257, Q9HAT1 | MIA3 | Q5JRA6 | NANS | Q9NR45 |
| PGM3 | O95394 | PMM2 | O15305, Q92871 | SEC16A | O15027 |
| SEC24D | O94855 | SLC35A1 | P78382 | ST6GALNAC1 | Q9NSC7 |
| TMED3 | Q9Y3Q3 | USO1 | O60763 | | |

#### 3. ER to Golgi Anterograde Transport (R-HSA-199977)

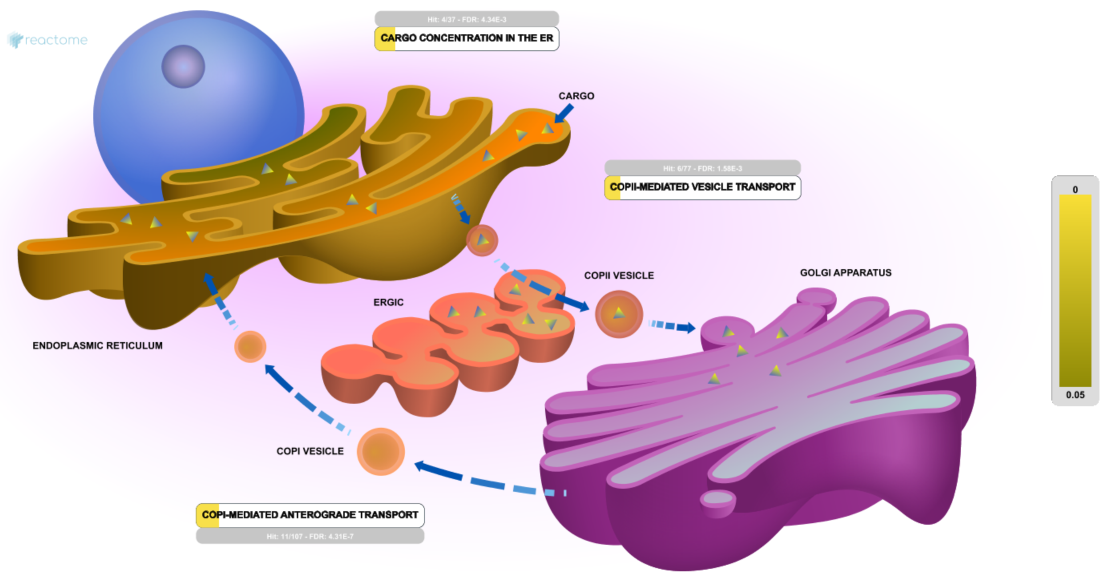


Secretory cargo destined to be secreted or to arrive at the plasma membrane (PM) leaves the ER via distinct exit sites. This cargo is destined for the Golgi apparatus for further processing.

About 25% of the proteome may be exported from the ER in human cells. This cargo is recognized and concentrated into COPII vesicles, which range in size from 60-90 nm, and move cargo from the ER to the ERGIC. Soluble cargo in the ER lumen is concentrated into COPII vesicles through interaction with a receptor with the receptor subsequently recycled to the ER in COPI vesicles through retrograde traffic.

The ERGIC (ER-to-Golgi intermediate compartment, also known as vesicular-tubular clusters, VTCs) is a stable, biochemically distinct compartment located adjacent to ER exit sites.

Retrograde traffic makes use of microtubule-directed COPI-coated vesicles, carrying ER proteins and membrane back to the ER.

##### Edit history

| Date | Action | Author |
|---|---|---|
| 2007-07-14 | Authored | Gillespie ME |
| 2007-07-14 | Created | Gillespie ME |
| 2007-07-19 | Edited | Gillespie ME |
| 2010-11-18 | Reviewed | Gagneux P |
| 2015-04-18 | Revised | Rothfels K |
| 2015-08-18 | Revised | Gillespie ME |
| 2021-05-22 | Modified | Shorser S |

#### Entities found in this pathway (13)

| Input | UniProt Id | Input | UniProt Id | Input | UniProt Id |
|---|---|---|---|---|---|
| ARFGAP1 | Q8N6T3, Q9NP61 | COG3 | Q96JB2 | COPB2 | P35606 |
| GOLGA2 | Q08379 | GOLGB1 | Q14789 | KDELR2 | P24390, P33947 |
| KDELR3 | O43731 | LMAN1 | P49257, Q9HAT1 | MIA3 | Q5JRA6 |
| SEC16A | O15027 | SEC24D | O94855 | TMED3 | Q9Y3Q3 |
| USO1 | O60763 | | | | |

#### 4. Transport to the Golgi and subsequent modification (R-HSA-948021)

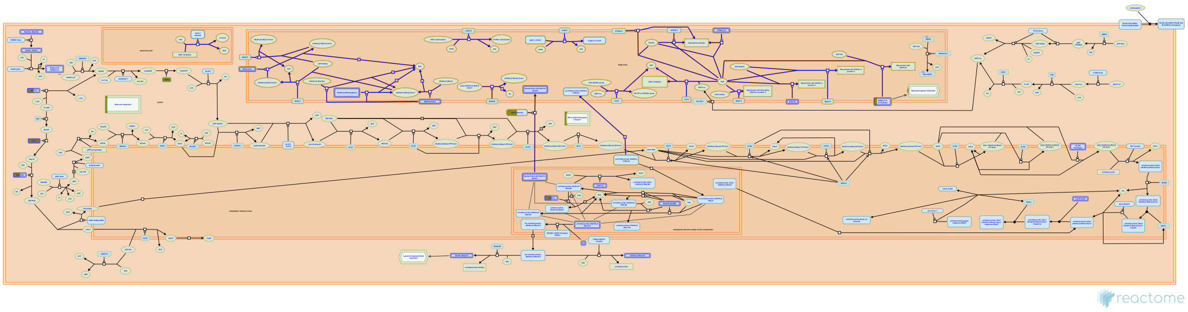

At least two mechanisms of transport of proteins from the ER to the Golgi have been described. One is a general flow requiring no export signals (Wieland et al, 1987; Martinez-Menarguez et al, 1999). The other is mediated by LMAN1/MCFD2, mannose-binding lectins that recognize a glycan signal (Zhang B et al, 2003).


##### Edit history

| Date | Action | Author |
|---|---|---|
| 2009-11-10 | Authored | Dall'Olio GM |
| 2010-09-15 | Edited | Jassal B |
| 2010-09-15 | Created | Jassal B |
| 2010-11-18 | Reviewed | Gagneux P |
| 2021-05-22 | Modified | Shorser S |

##### Entities found in this pathway (14)

| Input | UniProt Id | Input | UniProt Id | Input | UniProt Id |
|---|---|---|---|---|---|
| ARFGAP1 | Q8N6T3, Q9NP61 | B4GALT4 | O60513 | COG3 | Q96JB2 |
| COPB2 | P35606 | GOLGA2 | Q08379 | GOLGB1 | Q14789 |
| KDELR2 | P24390, P33947 | KDELR3 | O43731 | LMAN1 | P49257, Q9HAT1 |
| MIA3 | Q5JRA6 | SEC16A | O15027 | SEC24D | O94855 |
| TMED3 | Q9Y3Q3 | USO1 | O60763 | | |

#### 5. IRE1alpha activates chaperones (R-HSA-381070)

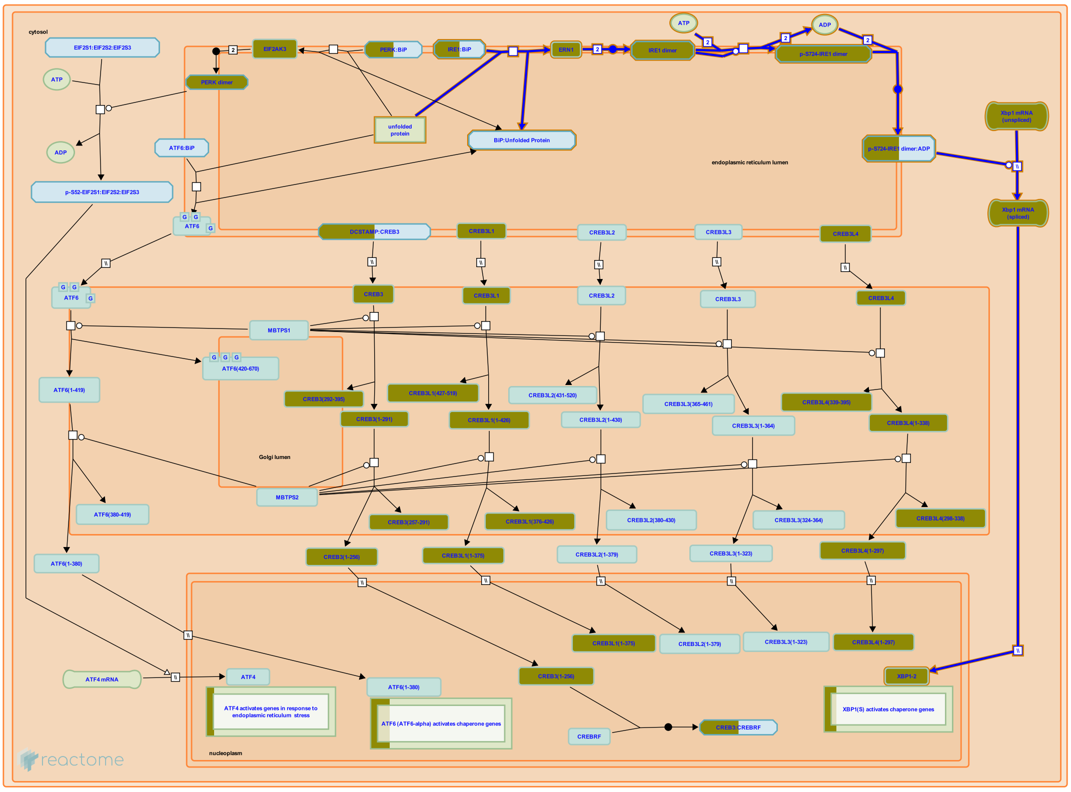

**Cellular compartments:** endoplasmic reticulum lumen, endoplasmic reticulum membrane, cytosol, nucleoplasm.

IRE1-alpha is a single-pass transmembrane protein that resides in the endoplasmic reticulum (ER) membrane. The C-terminus of IRE1-alpha is located in the cytosol; the N-terminus is located in the ER lumen. In unstressed cells IRE1-alpha exists in an inactive heterodimeric complex with BiP such that BiP in the ER lumen binds the N-terminal region of IRE1-alpha. Upon accumulation of unfolded proteins in the ER, BiP binds the unfolded protein and the IRE1-alpha:BiP complex dissociates. The dissociated IRE1-alpha then forms homodimers. Initially the luminal N-terminal regions pair. This is followed by trans-autophosphorylation of IRE1-alpha at Ser724 in the cytosolic C-terminal region. The phosphorylation causes a conformational change that allows the dimer to bind ADP, causing a further conformational change to yield back-to-back pairing of the cytosolic C-terminal regions of IRE1-alpha. The fully paired IRE1-alpha homodimer has endoribonuclease activity and cleaves the mRNA encoding Xbp-1. A 26 residue polyribonucleotide is released and the 5' and 3' fragments of the original Xbp-1 mRNA are rejoined. The spliced Xbp-1 message encodes Xbp-1 (S), a potent activator of transcription. Xbp-1 (S) together with the ubiquitous transcription factor NF-Y bind the ER Stress Responsive Element (ERSE) in a number of genes encoding chaperones. Recent data suggest that the IRE1-alpha homodimer can also cleave specific subsets of mRNAs, including the insulin (INS) mRNA in pancreatic beta cells.

#### Edit history

| Date | Action | Author |
|---|---|---|
| 2008-11-19 | Created | May B |
| 2008-11-20 | Edited | May B, Gopinathrao G |
| 2008-12-02 | Reviewed | Matthews L, D'Eustachio P, Gillespie ME |
| 2009-06-02 | Authored | May B |
| 2010-04-30 | Reviewed | Urano F |
| 2021-05-22 | Modified | Shorser S |

#### Entities found in this pathway (6)

| Input | UniProt Id | Input | UniProt Id | Input | UniProt Id |
|---|---|---|---|---|---|
| ARFGAP1 | Q8N6T3 | ERN2 | O75460 | GFPT1 | Q06210 |
| KDELR3 | O43731 | PDIA5 | Q14554 | XBP1 | P17861-2 |

| Input | Ensembl Id | Input | Ensembl Id | Input | Ensembl Id |
|---|---|---|---|---|---|
| ARFGAP1 | ENSG00000101199 | GFPT1 | ENSG00000198380 | KDELR3 | ENSG00000100196 |
| PDIA5 | ENSG00000065485 | XBP1 | ENST00000216037, ENST00000344347 | | |

#### 6. COPI-mediated anterograde transport (R-HSA-6807878)

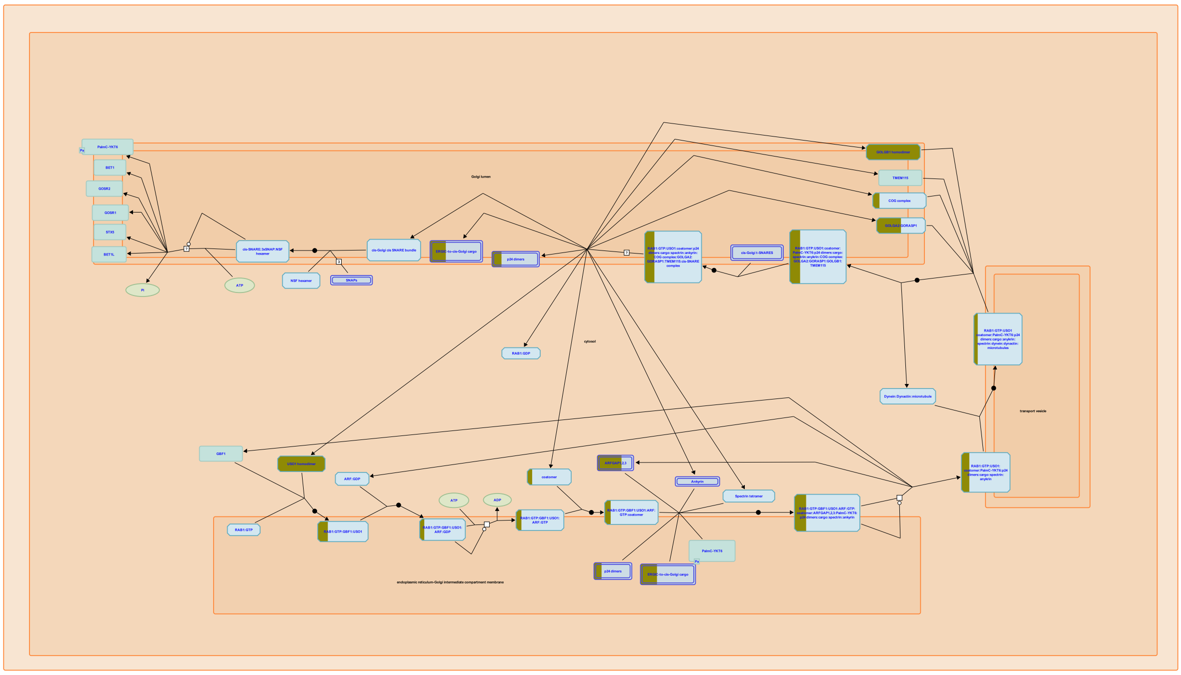

The ERGIC (ER-to-Golgi intermediate compartment, also known as vesicular-tubular clusters, VTCs) is a stable, biochemically distinct compartment located adjacent to ER exit sites (Ben-Tekaya et al, 2005; reviewed in Szul and Sztul, 2011). The ERGIC concentrates COPII-derived cargo from the ER for further anterograde transport to the cis-Golgi and also recycles resident ER proteins back to the ER through retrograde traffic. Both of these pathways appear to make use of microtubule-directed COPI-coated vesicles (Pepperkok et al, 1993; Presley et al, 1997; Scales et al, 1997; Stephens and Pepperkok, 2002; Stephens et al, 2000; reviewed in Lord et al, 2001; Spang et al, 2013).


##### Edit history

| Date | Action | Author |
|---|---|---|
| 2015-09-01 | Edited | Rothfels K |
| 2015-09-01 | Authored | Rothfels K |

| Date | Action | Author |
|---|---|---|
| 2015-09-02 | Reviewed | Gillespie ME |
| 2015-11-03 | Created | Rothfels K |
| 2021-05-22 | Modified | Shorser S |

#### Entities found in this pathway (9)

| Input | UniProt Id | Input | UniProt Id | Input | UniProt Id |
|---|---|---|---|---|---|
| ARFGAP1 | Q8N6T3, Q9NP61 | COG3 | Q96JB2 | COPB2 | P35606 |
| GOLGA2 | Q08379 | GOLGB1 | Q14789 | KDELR2 | P24390, P33947 |
| KDELR3 | O43731 | TMED3 | Q9Y3Q3 | USO1 | O60763 |

#### 7. XBP1(S) activates chaperone genes (R-HSA-381038)

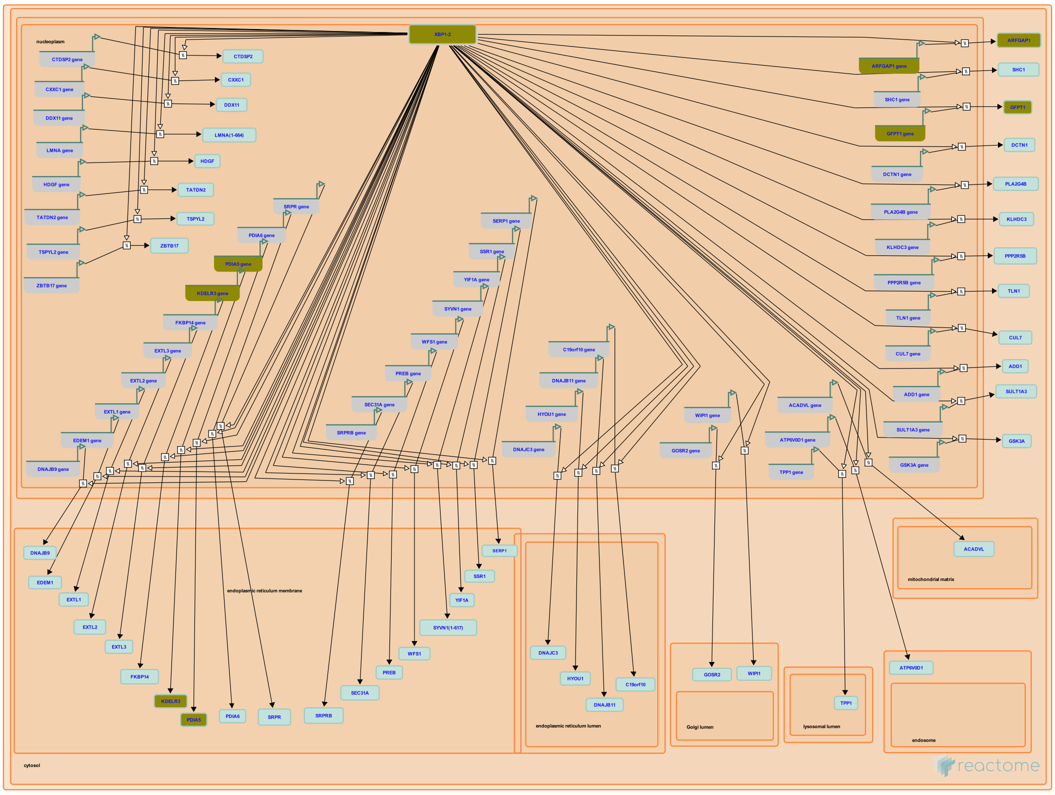

**Cellular compartments:** nucleoplasm, cytosol, endoplasmic reticulum lumen, endoplasmic reticulum membrane.

Xbp-1 (S) binds the sequence CCACG in ER Stress Responsive Elements (ERSE, consensus sequence CCAAT (N)9 CCACG) located upstream from many genes. The ubiquitous transcription factor NF-Y, a heterotrimer, binds the CCAAT portion of the ERSE and together the IRE1-alpha: NF-Y complex activates transcription of a set of chaperone genes including DNAJB9, EDEM, RAMP4, p58IPK, and others. This results in an increase in protein folding activity in the ER.


##### Edit history

| Date | Action | Author |
|---|---|---|
| 2008-11-19 | Created | May B |
| 2008-12-02 | Reviewed | Matthews L, D'Eustachio P, Gillespie ME |
| 2009-06-02 | Edited | May B |
| 2009-06-02 | Authored | May B |
| 2010-04-30 | Reviewed | Urano F |
| 2018-11-25 | Modified | May B |

#### Entities found in this pathway (5)

| Input | UniProt Id | Input | UniProt Id | Input | UniProt Id |
|---|---|---|---|---|---|
| ARFGAP1 | Q8N6T3 | GFPT1 | Q06210 | KDELR3 | O43731 |
| PDIA5 | Q14554 | XBP1 | P17861-2 | | |

| Input | Ensembl Id | Input | Ensembl Id |
|---|---|---|---|
| ARFGAP1 | ENSG00000101199 | GFPT1 | ENSG00000198380 |
| KDELR3 | ENSG00000100196 | PDIA5 | ENSG00000065485 |

#### 8. Signaling by FGFR2 IIIa TM (R-HSA-8851708)

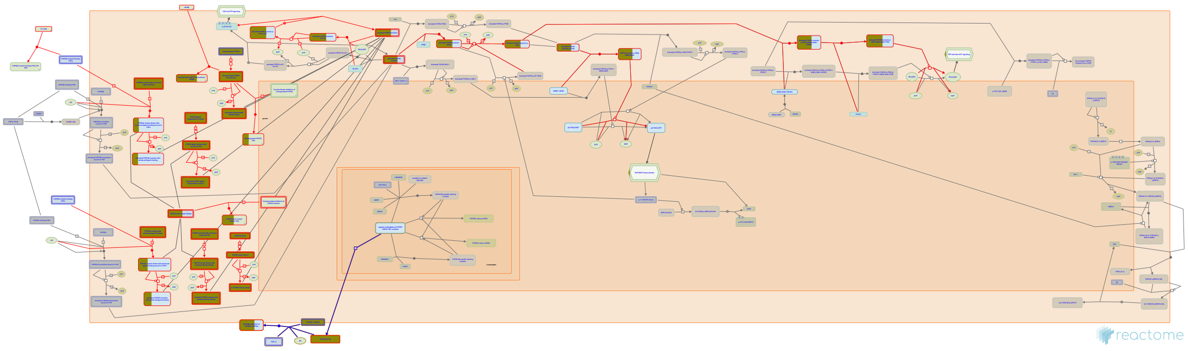

**Diseases:** acrocephalosyndactylia.

A soluble truncated form of FGFR2 is aberrantly expressed in an Apert Syndrome mouse model and inhibits FGFR signaling in vitro and in vivo. This variant, termed FGFR IIIa TM, arises from an mis-spliced transcript that fuses exon 7 to exon 10 and that escapes nonsense-mediated decay. FGFR2 IIIa TM may inhibit signaling by sequestering FGF ligand and/or by forming nonfunctional heterodimers with full-length receptors at the cell surface (Wheldon et al, 2011).


##### Edit history

| Date | Action | Author |
|---|---|---|
| 2016-01-09 | Edited | Rothfels K |
| 2016-01-09 | Authored | Rothfels K |
| 2016-01-09 | Created | Rothfels K |
| 2016-01-25 | Modified | Rothfels K |
| 2016-01-25 | Reviewed | Grose RP |

##### Entities found in this pathway (1)

| Input | UniProt Id |
|---|---|
| FGFR2 | P21802, P21802-1, P21802-18, P21802-3, P21802-5 |

#### 9. Post-translational protein modification (R-HSA-597592)

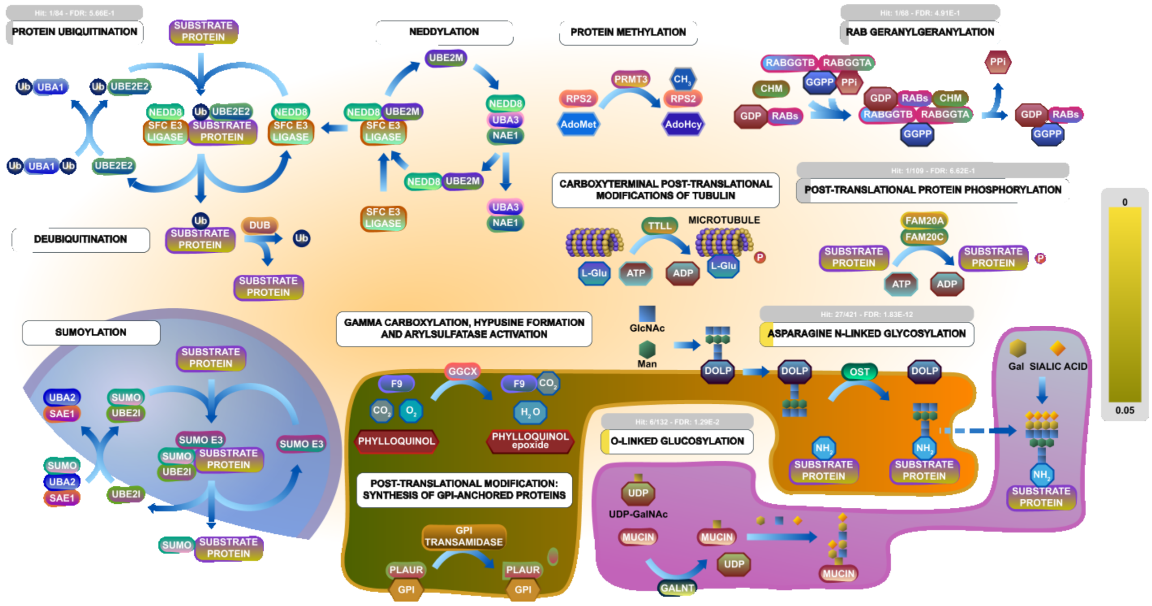


After translation, many newly formed proteins undergo further covalent modifications that alter their functional properties. Modifications associated with protein localization include the attachment of oligosaccharide moieties to membrane-bound and secreted proteins (**N-linked** and **O-linked glycosylation**), the attachment of lipid (**RAB geranylgeranylation**) or glycolipid moieties ( **GPI-anchored proteins**) that anchor proteins to cellular membranes, and the vitamin K-dependent attachment of carboxyl groups to glutamate residues. Modifications associated with functions of specific proteins include **gamma carboxylation** of clotting factors, **hypusine formation** on eukaryotic translation initiation factor 5A, conversion of a cysteine residue to formylglycine (**arylsulfatase activation**), methylation of lysine and arginine residues on non-histone proteins (**protein methylation**), **protein phosphorylation** by secretory pathway kinases, and **carboxyterminal modifications of tubulin** involving the addition of polyglutamate chains.

**Protein ubiquitination** and **deubiquitination** play a major role in regulating protein stability and, together with **SUMOylation** and **neddylation**, can modulate protein function as well.

##### References

##### Edit history

| Date | Action | Author |
|---|---|---|
| 2005-04-18 | Authored | D'Eustachio P |
| 2010-04-14 | Created | D'Eustachio P |
| 2021-05-18 | Edited | D'Eustachio P |
| 2021-05-18 | Reviewed | Stafford DW, Orlean P |
| 2021-05-22 | Modified | Shorser S |

##### Entities found in this pathway (31)

| Input | UniProt Id | Input | UniProt Id | Input | UniProt Id |
|---|---|---|---|---|---|
| ARFGAP1 | Q8N6T3, Q9NP61 | ATOH1 | P61086 | B4GALT4 | O60513 |

| Input | UniProt Id | Input | UniProt Id | Input | UniProt Id |
| --- | --- | --- | --- | --- | --- |
| COG3 | Q96JB2 | COPB2 | P35606 | EDEM3 | Q9BZQ6 |
| GALNT12 | Q8IXK2 | GALNT5 | Q7Z7M9 | GALNT7 | Q86SF2 |
| GALNT8 | Q9NY28 | GFPT1 | Q06210 | GMPPB | Q9Y5P6 |
| GNE | Q9Y223 | GOLGA2 | Q08379 | GOLGB1 | Q14789 |
| KDELR2 | P24390, P33947 | KDELR3 | O43731 | LMAN1 | P49257, Q9HAT1 |
| MIA3 | Q5JRA6 | MUC2 | Q02817 | MUC4 | Q99102 |
| NANS | Q9NR45 | PGM3 | O95394 | PMM2 | O15305, Q92871 |
| RAB27A | P51159 | SEC16A | O15027 | SEC24D | O94855 |
| SLC35A1 | P78382 | ST6GALNAC1 | Q9NSC7 | TMED3 | Q9Y3Q3 |
| USO1 | O60763 | | | | |

#### 10. FGFR2 mutant receptor activation (R-HSA-1839126)

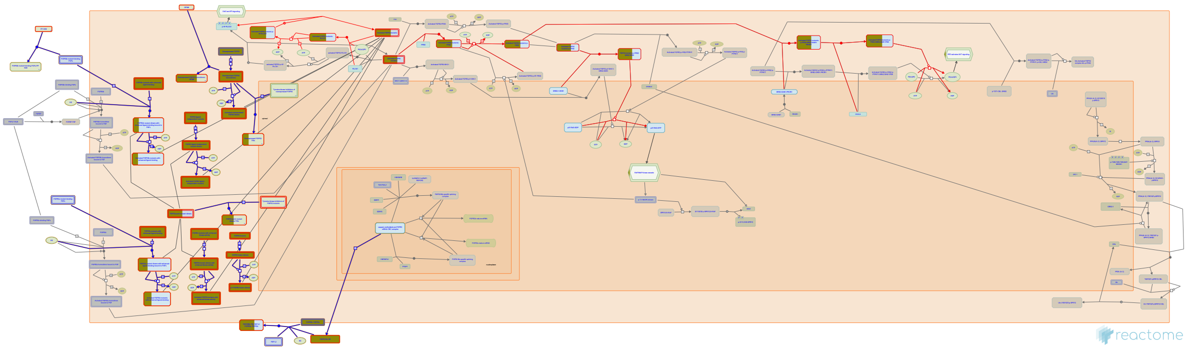

**Cellular compartments:** plasma membrane, cytosol, extracellular region.

**Diseases:** cancer, bone development disease.

Autosomal dominant mutations in FGFR2 are associated with the development of a range of skeletal disorders including Beare-Stevensen cutis gyrata syndrome, Pfeiffer syndrome, Jackson-Weiss syndrome, Crouzon syndrome and Apert Syndrome (reveiwed in Burke, 1998; Webster and Donoghue 1997; Cunningham, 2007). Activating point mutations have also been identified in FGFR2 in ~15% of endometrial cancers, as well as to a lesser extent in ovarian and gastric cancers (Dutt, 2008; Pollock, 2007; Byron, 2010; Jang, 2001). Activating mutations in FGFR2 are thought to contribute to receptor activation through diverse mechanisms, including constitutive ligand-independent dimerization (Robertson, 1998), expanded range and affinity for ligand (Ibrahimi, 2004b; Yu, 2000) and enhanced kinase activity (Byron, 2008; Chen, 2007). FGFR2 amplifications have been identified in 10% of gastric cancers, where they are associated with poor prognosis diffuse cancers (Hattori, 1996; Ueda, 1999; Shin, 2000; Kunii, 2008) , and in ~1% of breast cancers (Turner, 2010; Tannheimer, 2000). FGFR2 amplification often occur in conjunction with deletions of C-terminal exons, resulting in expression of a internalization- and degradation-resistant form of the receptor (Takeda, 2007; Cha, 2008, 2009). Signaling through overexpressed FGFR2 shows evidence of being ligand-independent and sensitive to FGFR inhibitors (Lorenzi, 1997; Takeda, 2007; Cha, 2009). More recently, FGFR2 fusion proteins have been identified in a number of cancers; these are thought to form constitutive ligand-independent dimers based on the dimerization domains of the 3' fusion partners and contribute to cellular proliferation and tumorigenesis in a kinase-inhibitor sensitive manner (Wu, 2013; Arai, 2013; Seo, 2012; reviewed in Parker, 2014).

#### Edit history

| Date | Action | Author |
|---|---|---|
| 2011-10-27 | Created | Rothfels K |
| 2012-02-10 | Authored | Rothfels K |
| 2012-05-15 | Reviewed | Ezzat S |
| 2012-05-16 | Edited | Rothfels K |
| 2016-01-09 | Revised | Rothfels K |
| 2016-01-22 | Modified | Rothfels K |

#### Entities found in this pathway (1)

| Input | UniProt Id |
|---|---|
| FGFR2 | P21802, P21802-1, P21802-17, P21802-18, P21802-3, P21802-5 |

#### 11. Synthesis of substrates in N-glycan biosynthesis (R-HSA-446219)

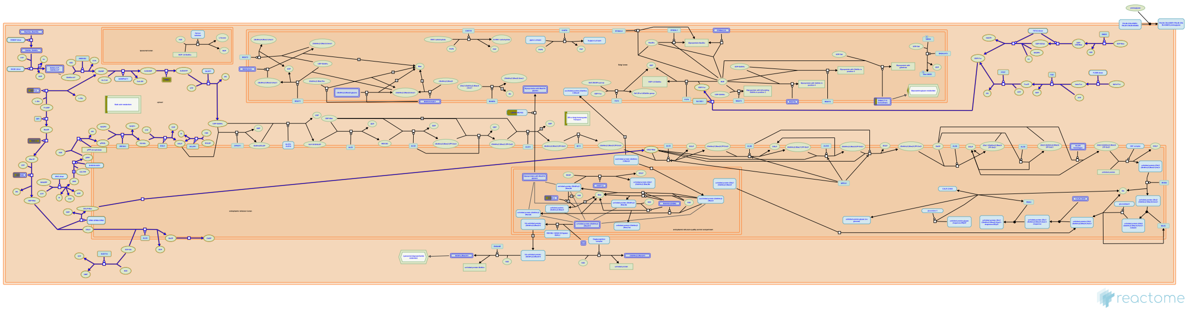

Reactions for the synthesis of the small nucleotide-linked sugar substrates that are used in the synthesis of the N-glycan precursor and in the later steps of glycosylation are annotated here.

All these nucleotide-linked sugar donors are synthesized in the cytosol; however, to participate in the later reactions of N-glycan precursor biosynthesis (when the glycan is oriented toward the lumen of the endoplasmic reticulum (ER)), these substrates must be attached to a dolichyl-phosphate molecule and then flipped toward the luminal side of the ER, through a mechanism which is still not known but which involves a different protein than the one that mediates the flipping of the LLO itself (Sanyal et al. 2008). Two of the genes encoding enzymes involved in these reactions, MPI and PMM2, are known to be associated with Congenital Disorders of Glycosylation (CDG) diseases of type I. Of these, CDG-Ia, associated with defects in PMM2, is the most frequent CDG disease reported.


##### Edit history

| Date | Action | Author |
|---|---|---|
| 2009-11-10 | Edited | Jassal B |
| 2009-11-10 | Authored | Dall'Olio GM |
| 2009-11-10 | Created | Jassal B |
| 2010-04-16 | Reviewed | Gagneux P |
| 2021-05-22 | Modified | Shorser S |

##### Entities found in this pathway (8)

| Input | UniProt Id | Input | UniProt Id | Input | UniProt Id |
|---|---|---|---|---|---|
| GFPT1 | Q06210 | GMPPB | Q9Y5P6 | GNE | Q9Y223 |
| NANS | Q9NR45 | PGM3 | O95394 | PMM2 | O15305, Q92871 |
| SLC35A1 | P78382 | ST6GALNAC1 | Q9NSC7 | | |

#### 12. Signaling by FGFR2 in disease (R-HSA-5655253)

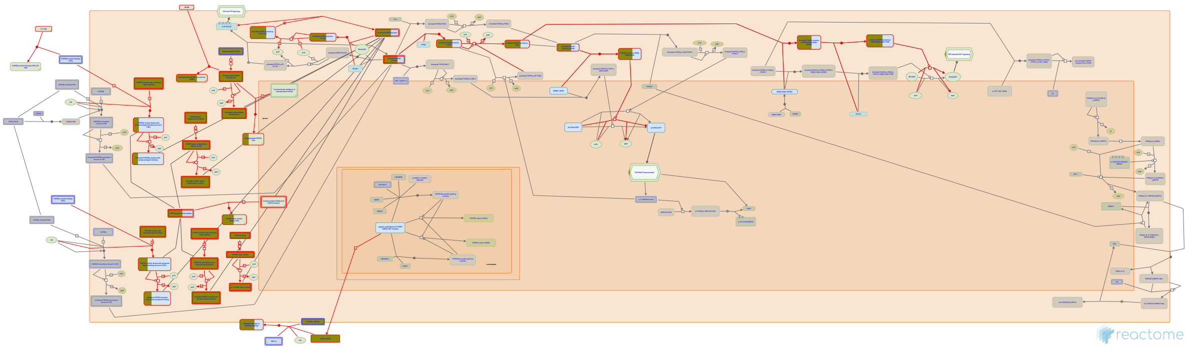


**Diseases:** cancer, bone development disease.

The FGFR2 gene has been shown to be subject to activating mutations and gene amplification leading to a variety of proliferative and developmental disorders depending on whether these events occur in the germline or arise somatically. Activating FGFR2 mutations in the germline give rise to a range of craniosynostotic conditions including Pfeiffer, Apert, Jackson-Weiss, Crouzon and Beare-Stevensen Cutis Gyrata syndromes. These autosomal dominant skeletal disorders are characterized by premature fusion of several sutures in the skull, and in some cases also involve syndactyly (abnormal bone fusions in the hands and feet) (reviewed in Webster and Donoghue, 1997; Burke, 1998; Cunningham, 2007).

Activating FGFR2 mutations arising somatically have been linked to the development of gastric and endometrial cancers (reviewed in Greulich and Pollock, 2011; Wesche, 2011). Many of these mutations are similar or identical to those that contribute to the autosomal disorders described above. Notably, loss-of-function mutations in FGFR2 have also been recently described in melanoma (Gartside, 2009). FGFR2 may also contribute to tumorigenesis through overexpression, as FGFR2 has been identified as a target of gene amplification in gastric and breast cancers (Kunii, 2008; Takeda, 2007).


##### Edit history

| Date | Action | Author |
|---|---|---|
| 2012-05-15 | Reviewed | Ezzat S |
| 2014-11-20 | Authored | Rothfels K |

| Date | Action | Author |
|---|---|---|
| 2014-12-05 | Edited | Rothfels K |
| 2014-12-05 | Created | Rothfels K |
| 2015-05-08 | Modified | Rothfels K |

#### Entities found in this pathway (1)

| Input | UniProt Id |
|---|---|
| FGFR2 | P21802, P21802-1, P21802-17, P21802-18, P21802-3, P21802-5 |

#### 13. Biosynthesis of the N-glycan precursor (dolichol lipid-linked oligosaccharide, LLO) and transfer to a nascent protein (R-HSA-446193)

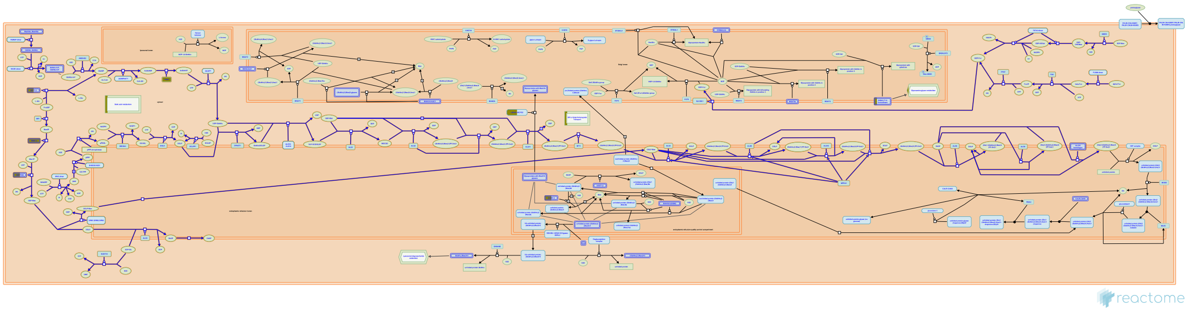

N-linked glycosylation commences with the 14-step synthesis of a dolichol lipid-linked oligosaccharide (LLO) consisting of 14 sugars (2 core GlcNAcs, 9 mannoses and 3 terminal GlcNAcs). This pathway is highly conserved in eukaryotes, and a closely related pathway is found in many eubacteria and Archaea. Mutations in the genes associated with N-glycan precursor synthesis lead to a diverse group of disorders collectively known as Congenital Disorders of Glycosylation (type I and II) (Sparks et al. 1993). The phenotypes of these disorders reflect the important role that N-glycosylation has during development, controlling the folding and the properties of proteins in the secretory pathway, and proteins that mediate cell-to-cell interactions or timing of development.


##### Edit history

| Date | Action | Author |
|---|---|---|
| 2009-11-10 | Edited | Jassal B |
| 2009-11-10 | Authored | Dall'Olio GM |
| 2009-11-10 | Created | Jassal B |
| 2010-04-16 | Reviewed | Gagneux P |
| 2021-05-22 | Modified | Shorser S |

##### Entities found in this pathway (8)

| Input | UniProt Id | Input | UniProt Id | Input | UniProt Id |
|---|---|---|---|---|---|
| GFPT1 | Q06210 | GMPPB | Q9Y5P6 | GNE | Q9Y223 |
| NANS | Q9NR45 | PGM3 | O95394 | PMM2 | O15305, Q92871 |
| SLC35A1 | P78382 | ST6GALNAC1 | Q9NSC7 | | |

#### 14. Membrane Trafficking (R-HSA-199991)

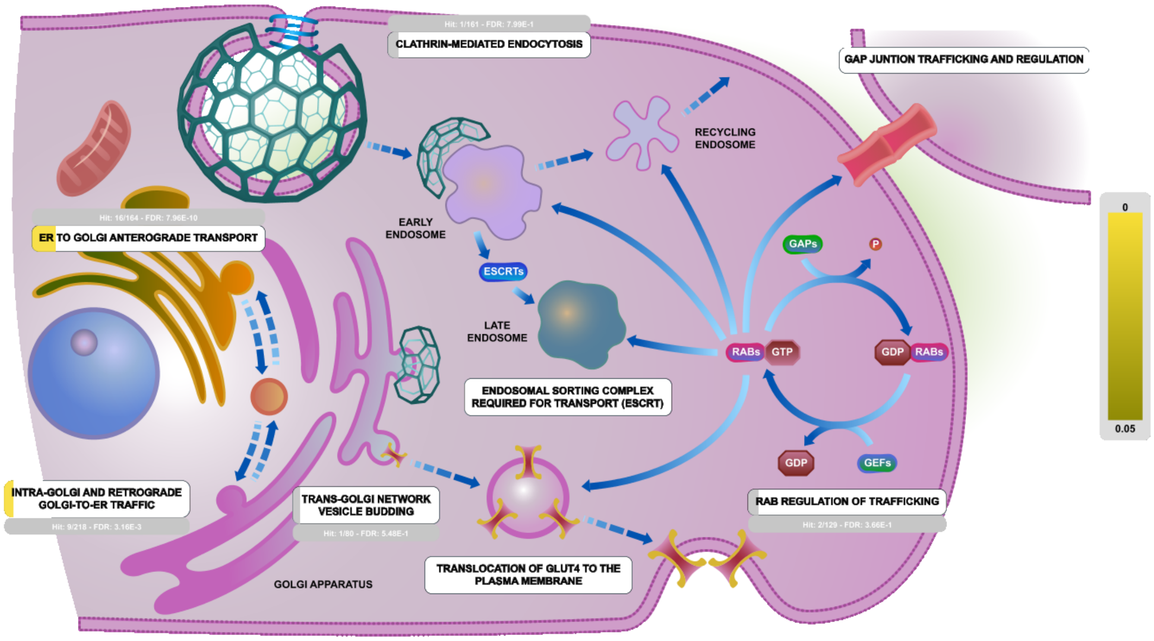


The secretory membrane system allows a cell to regulate delivery of newly synthesized proteins, carbohydrates, and lipids to the cell surface, a necessity for growth and homeostasis. The system is made up of distinct organelles, including the endoplasmic reticulum (ER), Golgi complex, plasma membrane, and tubulovesicular transport intermediates. These organelles mediate intracellular membrane transport between themselves and the cell surface. Membrane traffic within this system flows along highly organized directional routes. Secretory cargo is synthesized and assembled in the ER and then transported to the Golgi complex for further processing and maturation. Upon arrival at the trans Golgi network (TGN), the cargo is sorted and packaged into post-Golgi carriers that move through the cytoplasm to fuse with the cell surface. This directional membrane flow is balanced by retrieval pathways that bring membrane and selected proteins back to the compartment of origin.

##### Edit history

| Date | Action | Author |
|---|---|---|
| 2007-07-14 | Created | Gillespie ME |
| 2021-05-22 | Modified | Shorser S |

##### Entities found in this pathway (16)

| Input | UniProt Id | Input | UniProt Id | Input | UniProt Id |
|---|---|---|---|---|---|
| ARFGAP1 | Q8N6T3, Q9NP61 | ARL1 | P40616 | COG3 | Q96JB2 |
| COPB2 | P35606 | GOLGA2 | Q08379 | GOLGB1 | Q14789 |
| KDELR2 | P24390, P33947 | KDELR3 | O43731 | LMAN1 | P49257, Q9HAT1 |
| MIA3 | Q5JRA6 | RAB27A | P51159 | SEC16A | O15027 |

| Input | UniProt Id | Input | UniProt Id | Input | UniProt Id |
|---|---|---|---|---|---|
| SEC24D | O94855 | SYTL1 | Q8IYJ3 | TMED3 | Q9Y3Q3 |
| USO1 | O60763 | | | | |

#### 15. O-linked glycosylation of mucins (R-HSA-913709)

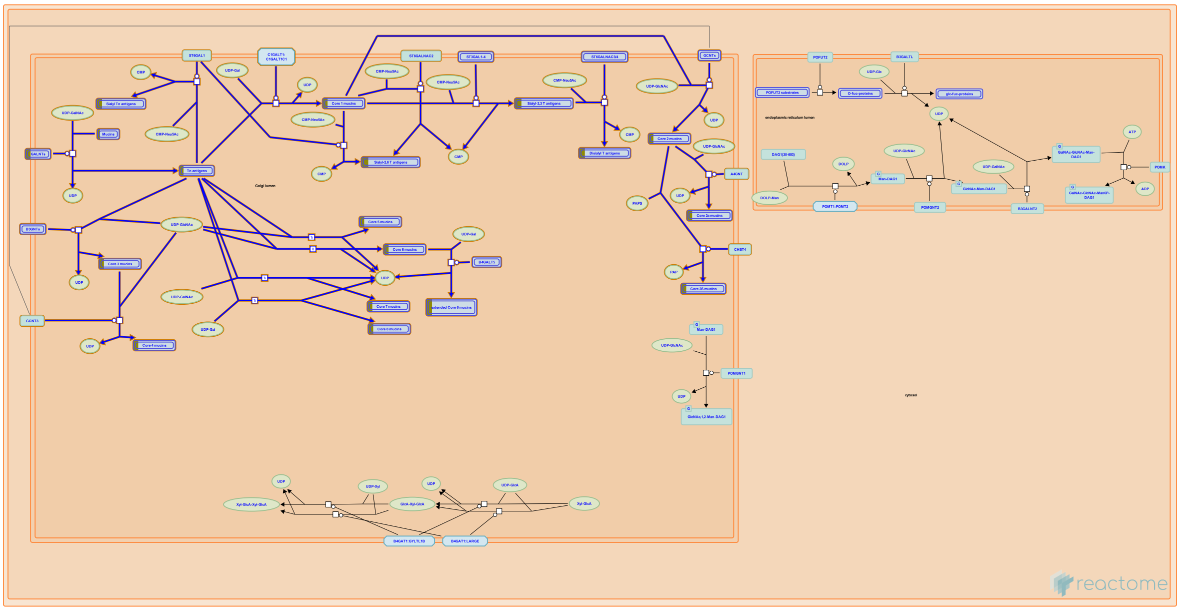

Mucins are a family of high molecular weight, heavily glycosylated proteins (glycoconjugates) produced by epithelial tissues in most metazoa. Mucins' key characteristic is their ability to form gels; therefore they are a key component in most gel-like secretions, serving functions from lubrication to cell signalling to forming chemical barriers. To date, there are approximately 20 genes that express mucins. Mature mucins are composed of two distinct regions:

(1) The amino- and carboxy-terminal regions are very lightly glycosylated, but rich in cysteines. The cysteine residues participate in establishing disulfide linkages within and among mucin monomers.

(2) A large central region rich in serine, threonine and proline residues called the variable number of tandem repeat (VNTR) region which can become heavily O-glycosylated with hundreds of O-GalNAc glycans.

N-acetyl-galactosamine (GalNAc) is the first glycan to be attached, forming the simplest mucin O-glycan. After this, several different pathways are possible generating "core" structures. Four core structures are commonly formed, several others are possible but infrequent. O-linked glycans are often capped by the addition of a sialic acid residue, terminating the addition of any more O-glycans (Brockhausen et al, 2009; Tarp and Clausen, 2008).


##### Edit history

| Date | Action | Author |
|---|---|---|
| 2010-07-19 | Edited | Jassal B |
| 2010-07-19 | Authored | Jassal B |

| Date | Action | Author |
|---|---|---|
| 2010-07-19 | Created | Jassal B |
| 2011-11-04 | Reviewed | Ferrer A |
| 2021-05-22 | Modified | Shorser S |

#### Entities found in this pathway (6)

| Input | UniProt Id | Input | UniProt Id | Input | UniProt Id |
|---|---|---|---|---|---|
| GALNT12 | Q8IXK2 | GALNT5 | Q7Z7M9 | GALNT7 | Q86SF2 |
| GALNT8 | Q9NY28 | MUC2 | Q02817 | MUC4 | Q99102 |

#### 16. COPI-dependent Golgi-to-ER retrograde traffic (R-HSA-6811434)

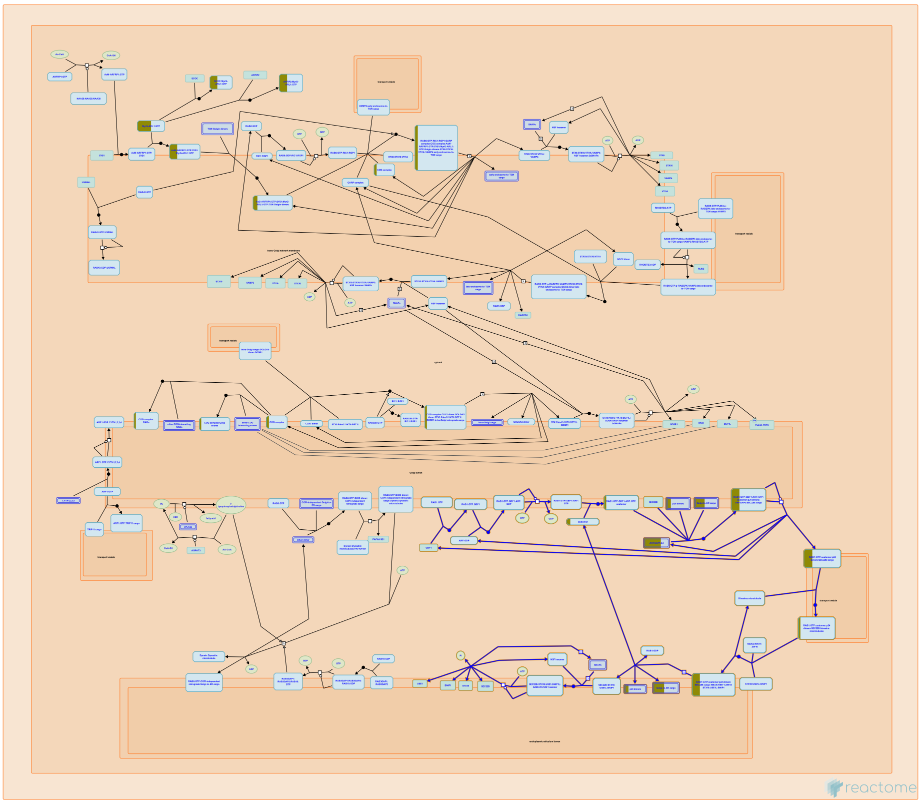

Retrograde traffic from the cis-Golgi to the ERGIC or the ER is mediated in part by microtubule-directed COPI-coated vesicles (Letourneur et al, 1994; Shima et al, 1999; Spang et al, 1998; reviewed in Lord et al, 2013; Spang et al, 2013). These assemble at the cis side of the Golgi in a GBF-dependent fashion and are tethered at the ER by the ER-specific SNAREs and by the conserved NRZ multisubunit tethering complex, known as DSL in yeast (reviewed in Tagaya et al, 2014; Hong and Lev, 2014). Typical cargo of these retrograde vesicles includes 'escaped' ER chaperone proteins, which are recycled back to the ER for reuse by virtue of their interaction with the Golgi localized KDEL receptors (reviewed in Capitani and Sallese, 2009; Cancino et al, 2013).


##### Edit history

| Date | Action | Author |
|---|---|---|
| 2015-11-09 | Edited | Rothfels K |
| 2015-11-09 | Authored | Rothfels K |
| 2015-11-19 | Created | Rothfels K |
| 2016-02-02 | Reviewed | Gillespie ME |
| 2021-05-22 | Modified | Shorser S |

#### Entities found in this pathway (5)

| Input | UniProt Id | Input | UniProt Id | Input | UniProt Id |
|---|---|---|---|---|---|
| ARFGAP1 | Q8N6T3, Q9NP61 | COPB2 | P35606 | KDELR2 | P24390, P33947 |
| KDELR3 | O43731 | TMED3 | Q9Y3Q3 | | |

#### 17. CREB3 factors activate genes (R-HSA-8874211)

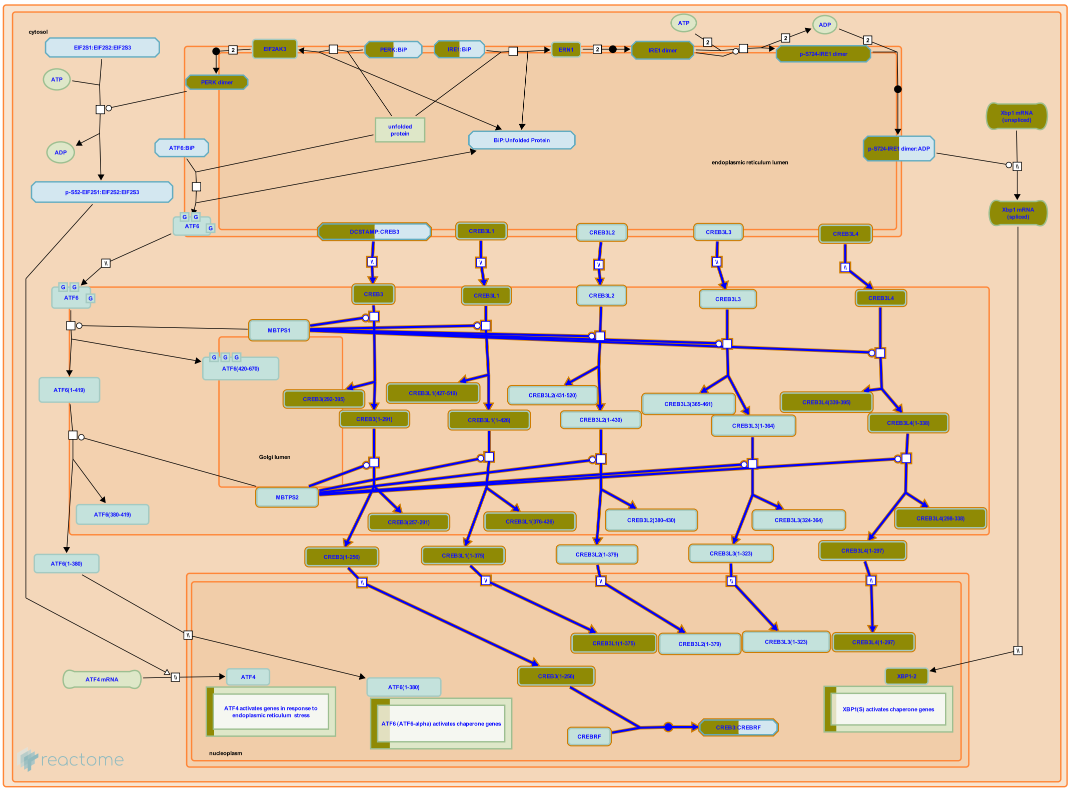

Members of the CREB3 family (also known as the OASIS family) are tissue-specific proteins that each contain a transcription activation domain, a basic leucine zipper (bZIP) domain that promotes dimerization and DNA binding, and a transmembrane domain that anchors the protein to the membrane of the endoplasmic reticulum (ER) (reviewed in Asada et al. 2011, Chan et al. 2011, Kondo et al. 2011, Fox and Andrew 2015). The family includes CREB3 (LUMAN), CREB3L1 (OASIS), CREB3L2 (BBF2H7, Tisp40), CREB3L3 (CREB-H), and CREB3L4 (CREB4). Activation of the proteins occurs when they transit from the ER to the Golgi and are cleaved sequentially by the Golgi resident proteases MBTPS1 (S1P) and MBTPS2 (S2P), a process known as regulated intramembrane proteolysis that releases the cytoplasmic region of the protein containing the transcription activation domain and the bZIP domain. This protein fragment then transits from the cytosol to the nucleus where it activates transcription of target genes. CREB3L1, CREB3L2, and CREB3L3 are activated by ER stress, although the mechanisms that cause the transit of the CREB3 proteins are not fully characterized. Unlike the ATF6 factors, CREB3 proteins do not appear to interact with HSPA5 (BiP) and therefore do not appear to sense unfolded proteins by dissociation of HSPA5 when HSPA5 binds the unfolded proteins.

#### Edit history

| Date | Action | Author |
|---|---|---|
| 2016-05-21 | Edited | May B |
| 2016-05-21 | Authored | May B |
| 2016-05-21 | Created | May B |
| 2017-02-25 | Reviewed | Schröder M |
| 2021-05-22 | Modified | Shorser S |

#### Entities found in this pathway (2)

| Input | UniProt Id | Input | UniProt Id |
|---|---|---|---|
| CREB3L1 | Q96BA8 | CREB3L4 | O43889, Q8TEY5 |

#### 18. Phospholipase C-mediated cascade; FGFR2 (R-HSA-5654221)

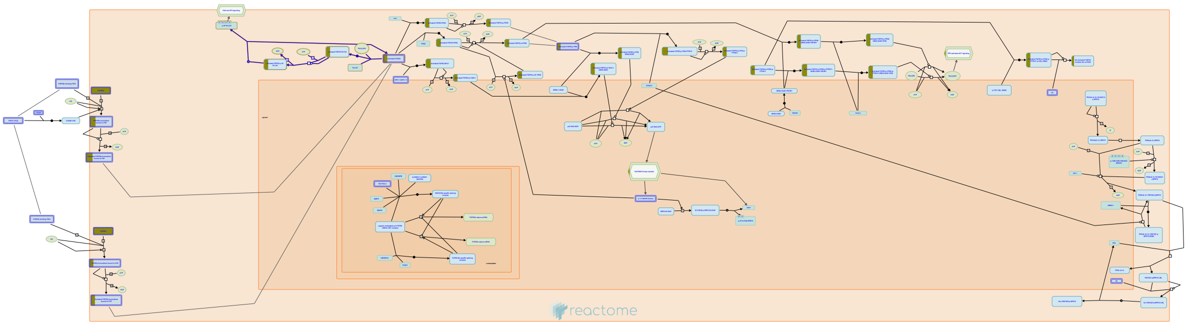

**Cellular compartments:** plasma membrane.

Phospholipase C-gamma (PLC-gamma) is a substrate of the fibroblast growth factor receptor (FGFR) and other receptors with tyrosine kinase activity. It is known that the src homology region 2 (SH2 domain) of PLC-gamma and of other signaling molecules (such as GTPase-activating protein and phosphatidylinositol 3-kinase-associated p85) direct their binding toward autophosphorylated tyrosine residues of the FGFR. Recruitment of PLC-gamma results in its phosphorylation and activation by the receptor. Activated PLC-gamma hydrolyzes phosphatidyl inositol[4,5] P2 to form the second messengers diacylglycerol (DAG) and Ins [1,4,5]P3, which stimulate calcium release and activation of calcium/calmodulin dependent kinases.


##### Edit history

| Date | Action | Author |
|---|---|---|
| 2007-01-10 | Authored | de Bono B |
| 2007-02-07 | Reviewed | Mohammadi M |
| 2010-02-03 | Edited | Jupe S |
| 2014-12-04 | Created | Rothfels K |
| 2021-05-21 | Modified | Shorser S |

##### Entities found in this pathway (1)

| Input | UniProt Id |
|---|---|
| FGFR2 | P21802-1, P21802-18, P21802-3, P21802-5 |

#### 19. COPII-mediated vesicle transport (R-HSA-204005)

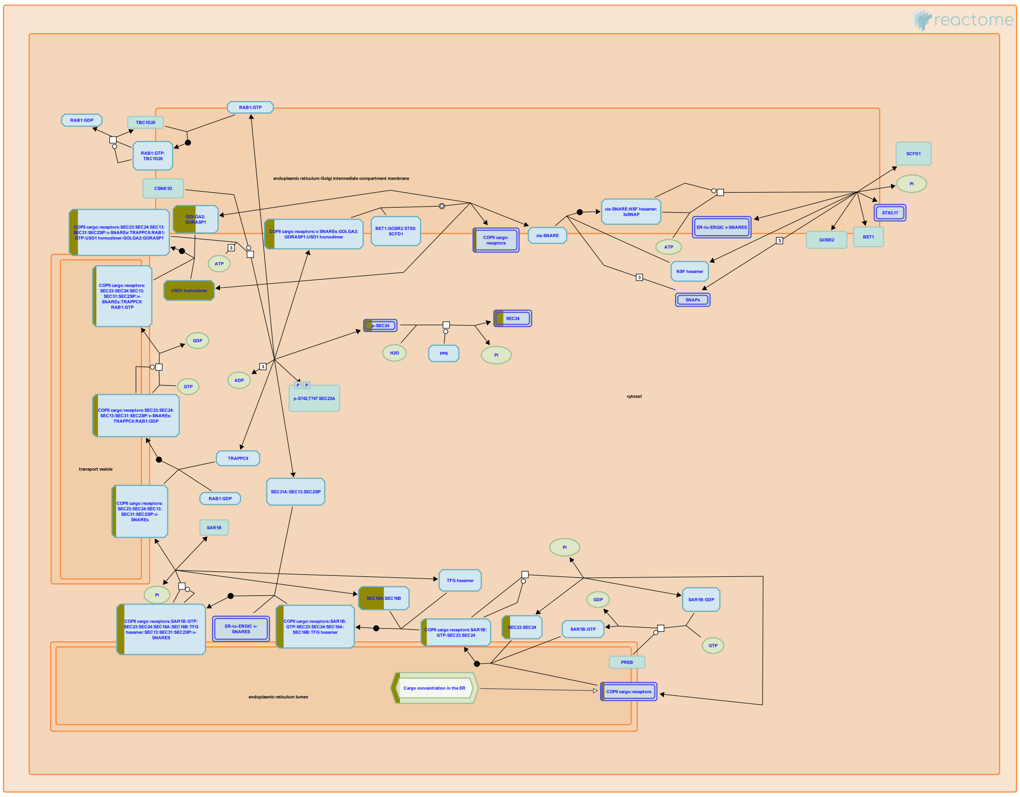

COPII components (known as Sec13p, Sec23p, Sec24p, Sec31p, and Sar1p in yeast) traffic cargo from the endoplasmic reticulum to the ER-Golgi intermediate compartment (ERGIC). COPII-coated vesicles were originally discovered in the yeast Saccharomyces cerevisiae using genetic approaches coupled with a cell-free assay. The mammalian counterpart of this pathway is represented here. Newly synthesized proteins destined for secretion are sorted into COPII-coated vesicles at specialized regions of the ER. These vesicles leave the ER, become uncoated and subsequently fuse with the ERGIC membrane.


##### Edit history

| Date | Action | Author |
|---|---|---|
| 2007-07-14 | Authored | Gillespie ME |
| 2007-11-22 | Edited | Gillespie ME |
| 2007-11-22 | Created | Gillespie ME |
| 2010-11-18 | Reviewed | Gagneux P |
| 2015-04-18 | Revised | Rothfels K |
| 2015-08-18 | Revised | Gillespie ME |
| 2021-05-31 | Modified | Shorser S |

#### Entities found in this pathway (5)

| Input | UniProt Id | Input | UniProt Id | Input | UniProt Id |
|---|---|---|---|---|---|
| GOLGA2 | Q08379 | LMAN1 | P49257, Q9HAT1 | SEC16A | O15027 |
| SEC24D | O94855 | USO1 | O60763 | | |

#### 20. FGFR2 ligand binding and activation (R-HSA-190241)

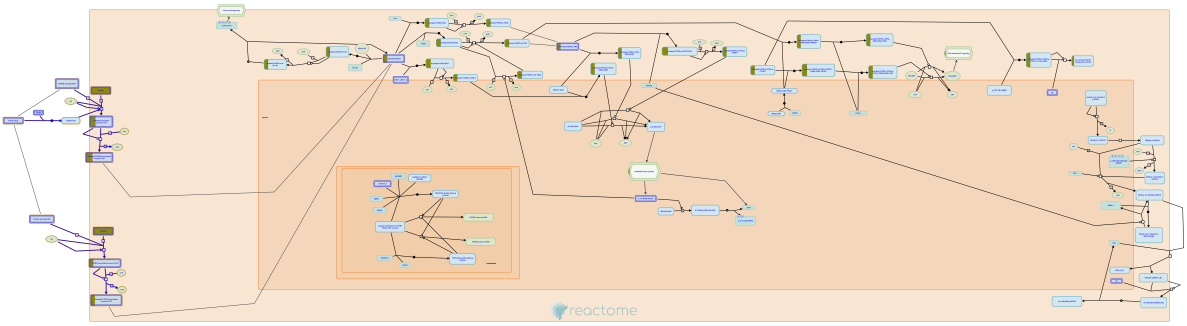

Dominant mutations in the fibroblast growth factor receptor 2 (FGFR2) gene have been identified as causes of four phenotypically distinct craniosynostosis syndromes, including Crouzon, Jackson-Weiss, Pfeiffer, and Apert syndromes. FGFR2 binds a number of different FGFs preferentially, as illustrated in this pathway.

FGFR is probably activated by NCAM very differently from the way by which it is activated by FGFs, reflecting the different conditions for NCAM-FGFR and FGF-FGFR interactions. The affinity of FGF for FGFR is approximately 10e6 times higher than that of NCAM for FGFR. Moreover, in the brain NCAM is constantly present on the cell surface at a much higher (micromolar) concentration than FGFs, which only appear transiently in the extracellular environment in the nanomolar range.


##### Edit history

| Date | Action | Author |
|---|---|---|
| 2006-12-18 | Created | de Bono B |
| 2007-01-10 | Authored | de Bono B |
| 2007-02-07 | Reviewed | Mohammadi M |
| 2007-02-11 | Edited | D'Eustachio P, de Bono B |
| 2021-05-21 | Modified | Shorser S |

##### Entities found in this pathway (1)

| Input | UniProt Id |
|---|---|
| FGFR2 | P21802-1, P21802-18, P21802-3, P21802-5 |

#### 21. Signaling by FGFR in disease (R-HSA-1226099)

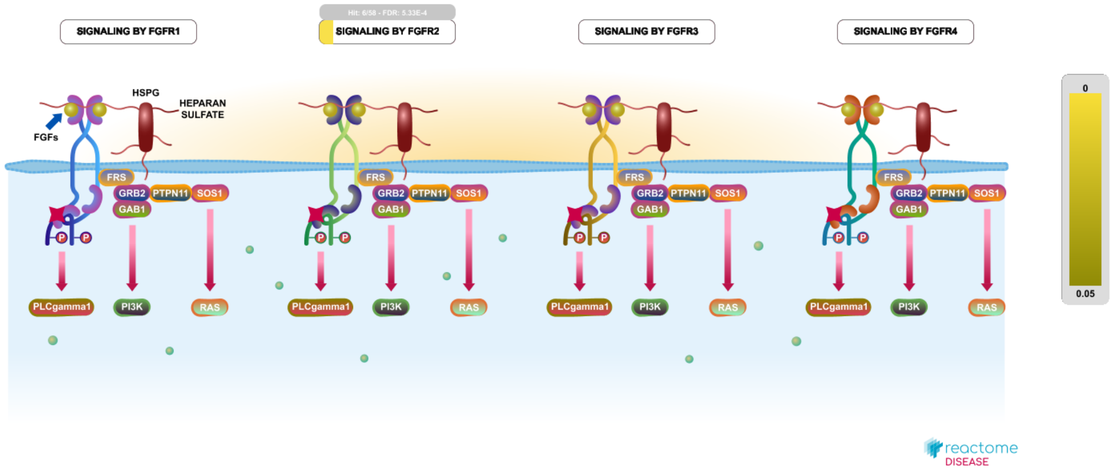


**Diseases:** bone development disease, cancer.

A number of skeletal and developmental diseases have been shown to arise as a result of mutations in the FGFR1, 2 and 3 genes. These include dwarfism syndromes (achondroplasia, hypochondroplasia and the neonatal lethal disorders thanatophoric dysplasia I and II), as well as craniosynostosis disorders such as Pfeiffer, Apert, Crouzon, Jackson-Weiss and Muenke syndromes (reviewed in Webster and Donoghue 1997; Burke, 1998, Cunningham, 2007; Harada, 2009). These mutations fall into four general regions of the receptor: a) the immunoglobulin (Ig)-like domain II-III linker region, b) the alternatively spliced second half of the Ig III domain, c) the transmembrane domain and d) the tyrosine kinase domain (reviewed in Webster and Donoghue, 1997). With the exception of mutations in class b), which affect only the relevant splice variant, these mutations may be present in either the 'b' or 'c' isoforms. These activating mutations affect FGFR function by altering or expanding the ligand-binding range of the receptors (see for instance Ibrahimi, 2004a), by promoting ligand-independent dimerization (for instance, Galvin,1996; Neilson and Friesel, 1996; d'Avis,1998) or by increasing the activity of the kinase domain (for instance, Webster, 1996; Naski, 1996; Tavormina, 1999; Bellus, 2000). Thus, a number of the point mutations found in FGFR receptors alter their activity without altering their intrinsic kinase activity. Many of the mutations that promote constitutive dimerization do so by creating or removing cysteine residues; the presence of an unpaired cysteine in the receptor is believed to promote dimerization through the formation of intramolecular disulphide bonds (Galvin, 1996; Robertson, 1998). Paralogous mutations at equivalent positions have been identified in more than one FGF receptor, sometimes giving rise to different diseases. For instance, mutation of the highly conserved FGFR2 Ser252-Pro253 dipeptide in the region between the second and third Ig domain is responsible for virtually all cases of Apert Syndrome (Wilkie, 1995), while paralogous mutations in FGFR1 (S252R) and FGFR3 (P250R) are associated with Pfeiffer and Crouzon syndromes, respectively (Bellus, 1996). FGFR4 is unique in that mutations of this gene are not known to be associated with any developmental disorders.

Recently, many of the same activating mutations in the FGFR genes that have been characterized in skeletal and developmental disorders have begun to be identified in a range of cancers (reviewed in Turner and Gross, 2010; Greulich and Pollock, 2011; Wesche, 2011). The best established link between a somatic mutation of an FGFR and the development of cancer is in the case of FGFR3, where 50% of bladder cancers have mutations in the FGFR3 coding sequence. Of these mutations, which largely match the activating mutations seen in thanatophoric dysplasias, over half occur at a single residue (S249C) (Cappellen, 1999; van Rhijn, 2002). Activating mutations have also been identified in the coding sequences of FGFR1, 2 and 4 (for review, see Wesche, 2011)

In addition to activating point mutations, the FGFR1, 2 and 3 genes are subject to misregulation in cancer through gene amplification and translocation events, which are thought to lead to overexpression and ligand-independent dimerization (Weiss, 2010; Turner, 2010; Kunii, 2008; Takeda, 2007; Chesi, 1997; Avet-Loiseau, 1998; Ronchetti, 2001). It is important to note, however, that in each of these cases, the amplification or translocation involve large genomic regions encompassing additional genes, and the definitive roles of the FGFR genes in promoting oncogenesis has not been totally established. In the case of FGFR1, translocation events also give rise to FGFR1 fusion proteins that contain the intracellular kinase domain of the receptor fused to a dimerization domain from the partner gene. These fusions, which are expressed in a pre-leukemic myeloproliferative syndrome, dimerize constitutively based on the dimerization domain provided by the fusion partner and are constitutively active (reviewed in Jackson, 2010).


#### Edit history

| Date | Action | Author |
|---|---|---|
| 2011-03-09 | Created | Rothfels K |
| 2012-02-10 | Authored | Rothfels K |
| 2012-05-15 | Edited | Rothfels K |
| 2012-05-15 | Reviewed | Ezzat S |
| 2016-01-25 | Reviewed | Grose RP |
| 2021-05-04 | Modified | Matthews L |

#### Entities found in this pathway (1)

| Input | UniProt Id |
|---|---|
| FGFR2 | P21802, P21802-1, P21802-17, P21802-18, P21802-3, P21802-5 |

#### 22. Synthesis of GDP-mannose (R-HSA-446205)

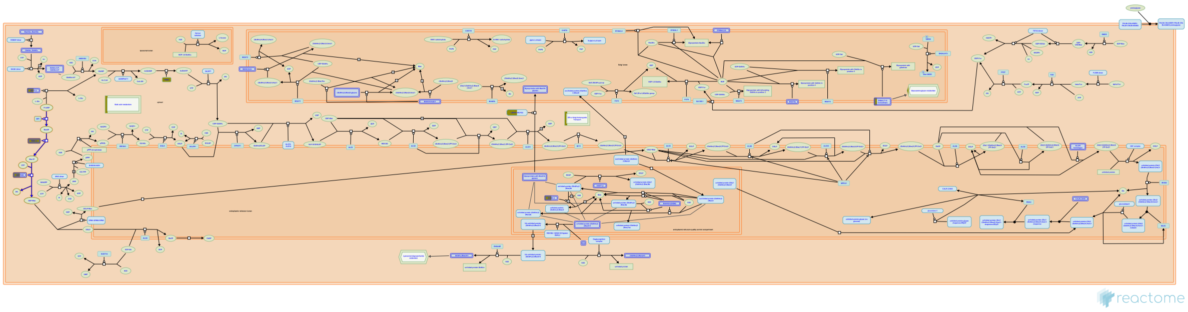

GDP-mannose is the mannose donor for the first 5 mannose addition reactions in the N-glycan precursor synthesis, and also for the synthesis of Dolichyl-phosphate-mannose involved in other mannose transfer reactions. It is synthesized from fructose 6-phosphate and GTP in three steps.

##### References

##### Edit history

| Date | Action | Author |
|---|---|---|
| 2009-11-10 | Edited | Jassal B |
| 2009-11-10 | Authored | Dall'Olio GM |
| 2009-11-10 | Created | Jassal B |
| 2010-04-16 | Reviewed | Gagneux P |
| 2021-05-22 | Modified | Shorser S |

##### Entities found in this pathway (2)

| Input | UniProt Id | Input | UniProt Id |
|---|---|---|---|
| GMPPB | Q9Y5P6 | PMM2 | O15305, Q92871 |

#### 23. PI-3K cascade:FGFR2 (R-HSA-5654695)

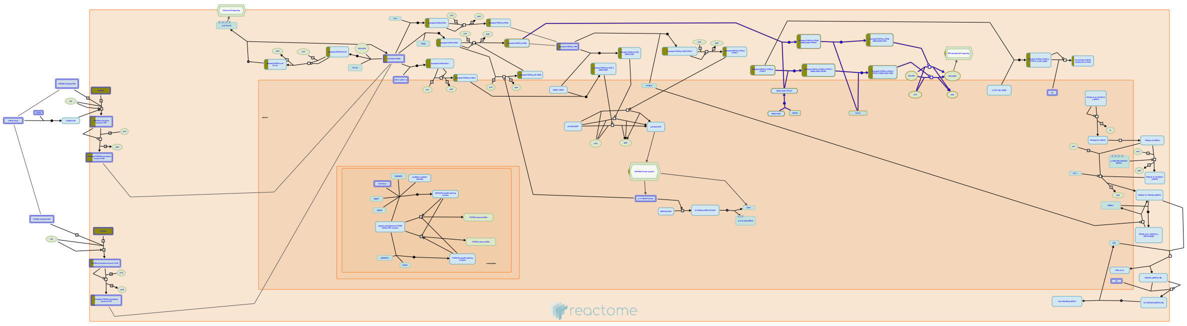

The ability of growth factors to protect from apoptosis is primarily due to the activation of the AKT survival pathway. P-I-3-kinase dependent activation of PDK leads to the activation of AKT which in turn affects the activity or expression of pro-apoptotic factors, which contribute to protection from apoptosis. AKT activation also blocks the activity of GSK-3b which could lead to additional antiapoptotic signals.


##### Edit history

| Date | Action | Author |
|---|---|---|
| 2007-01-10 | Authored | de Bono B |
| 2007-02-07 | Reviewed | Mohammadi M |
| 2010-02-03 | Edited | Jupe S |
| 2014-12-04 | Created | Rothfels K |
| 2021-05-31 | Modified | Shorser S |

##### Entities found in this pathway (1)

| Input | UniProt Id |
|---|---|
| FGFR2 | P21802-1, P21802-18, P21802-3, P21802-5 |

#### 24. Metabolism of proteins (R-HSA-392499)

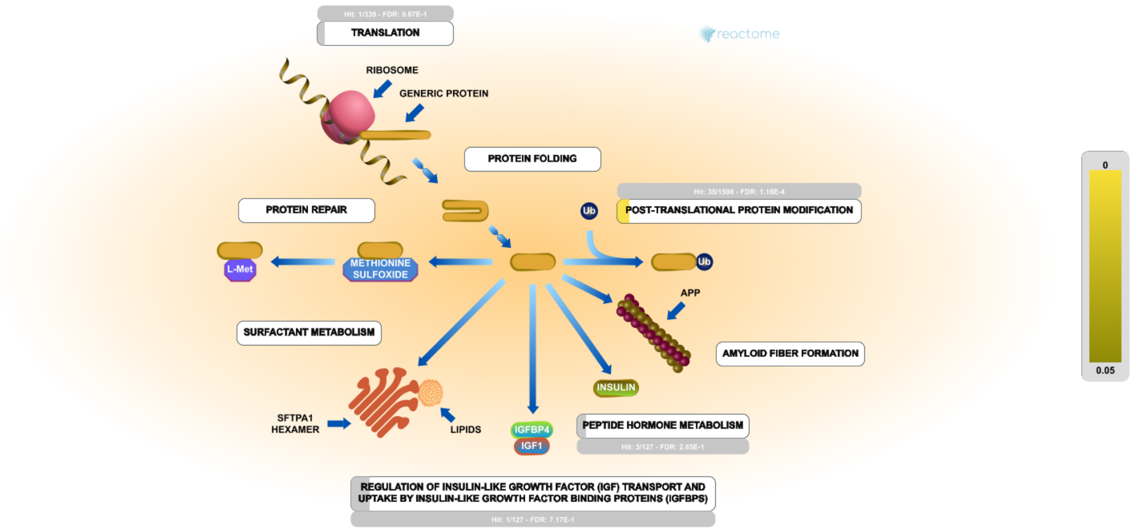


Metabolism of proteins, as annotated here, covers the full life cycle of a protein from its synthesis to its posttranslational modification and degradation, at various levels of specificity. Protein synthesis is accomplished through the process of Translation of an mRNA sequence into a polypeptide chain. Protein folding is achieved through the function of molecular chaperones which recognize and associate with proteins in their non-native state and facilitate their folding by stabilizing the conformation of productive folding intermediates (Young et al. 2004). Following translation, many newly formed proteins undergo Post-translational protein modification, essentially irreversible covalent modifications critical for their mature locations and functions (Knorre et al. 2009), including gamma carboxylation, synthesis of GPI-anchored proteins, asparagine N-linked glycosylation, O-glycosylation, SUMOylation, ubiquitination, deubiquitination, RAB geranylgeranylation, methylation, carboxyterminal post-translational modifications, neddylation, and phosphorylation. Peptide hormones are synthesized as parts of larger precursor proteins whose cleavage in the secretory system (endoplasmic reticulum, Golgi apparatus, secretory granules) is annotated in Peptide hormone metabolism. After secretion, peptide hormones are modified and degraded by extracellular proteases (Chertow, 1981 PMID:6117463). Protein repair enables the reversal of damage to some amino acid side chains caused by reactive oxygen species. Pulmonary surfactants are lipids and proteins that are secreted by the alveolar cells of the lung that decrease surface tension at the air/liquid interface within the alveoli to maintain the stability of pulmonary tissue (Agassandian and Mallampalli 2013). Nuclear regulation, transport, metabolism, reutilization, and degradation of surfactant are described in the Surfactant metabolism pathway. Amyloid fiber formation, the accumulation of mostly extracellular deposits of fibrillar proteins, is associated with tissue damage observed in numerous diseases including late phase heart failure (cardiomyopathy) and neurodegenerative diseases such as Alzheimer's, Parkinson's, and Huntington's.

##### References

##### Edit history

| Date | Action | Author |
|---|---|---|
| 2009-03-04 | Edited | Matthews L |

| Date | Action | Author |
|---|---|---|
| 2009-03-05 | Authored | Matthews L |
| 2009-03-05 | Created | Matthews L |
| 2021-05-22 | Modified | Shorser S |

#### Entities found in this pathway (34)

| Input | UniProt Id | Input | UniProt Id | Input | UniProt Id |
|---|---|---|---|---|---|
| ARFGAP1 | Q8N6T3, Q9NP61 | ATOH1 | P61086 | B4GALT4 | O60513 |
| COG3 | Q96JB2 | COPB2 | P35606 | EDEM3 | Q9BZQ6 |
| GALNT12 | Q8IXK2 | GALNT5 | Q7Z7M9 | GALNT7 | Q86SF2 |
| GALNT8 | Q9NY28 | GFPT1 | Q06210 | GMPPB | Q9Y5P6 |
| GNE | Q9Y223 | GOLGA2 | Q08379 | GOLGB1 | Q14789 |
| KDELR2 | P24390, P33947 | KDELR3 | O43731 | KLF4 | O43474 |
| LMAN1 | P49257, Q9HAT1 | MIA3 | Q5JRA6 | MUC2 | Q02817 |
| MUC4 | Q99102 | NANS | Q9NR45 | PGM3 | O95394 |
| PMM2 | O15305, Q92871 | RAB27A | P51159 | SEC16A | O15027 |
| SEC24D | O94855 | SLC30A7 | Q8NEW0 | SLC35A1 | P78382 |
| SSR3 | Q9UNL2 | ST6GALNAC1 | Q9NSC7 | TMED3 | Q9Y3Q3 |
| USO1 | O60763 | | | | |

#### 25. SHC-mediated cascade:FGFR2 (R-HSA-5654699)

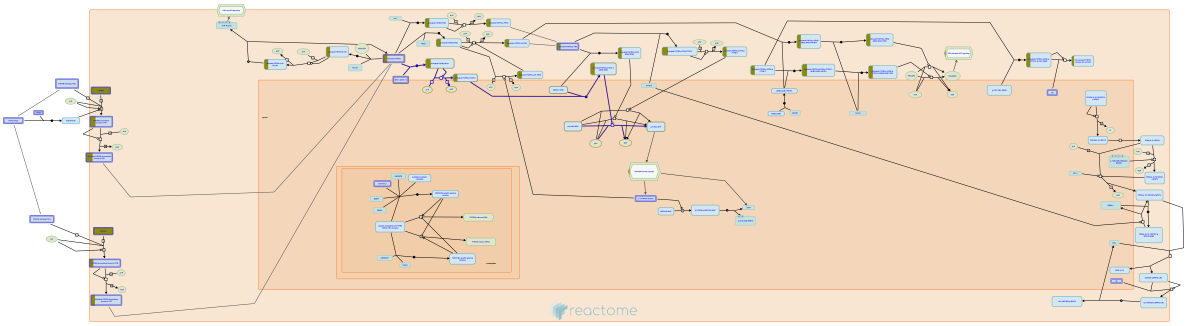

**Cellular compartments:** plasma membrane.

The exact role of SHC1 in FGFR signaling remains unclear. Numerous studies have shown that the p46 and p52 isoforms of SHC1 are phosphorylated in response to FGF stimulation, but direct interaction with the receptor has not been demonstrated. Co-precipitation of p46 and p52 with the FGFR2 IIIc receptor has been reported, but this interaction is thought to be indirect, possibly mediated by SRC. Consistent with this, co-precipitation of SHC1 and FGFR1 IIIc is seen in mammalian cells expressing v-SRC. The p66 isoform of SHC1 has also been co-precipitated with FGFR3, but this occurs independently of receptor stimulation, and the p66 isoform not been shown to undergo FGF-dependent phosphorylation. SHC1 has been shown to associate with GRB2 and SOS1 in response to FGF stimulation, suggesting that the recruitment of SHC1 may contribute to activation of the MAPK cascade downstream of FGFR.


##### Edit history

| Date | Action | Author |
|---|---|---|
| 2007-01-10 | Authored | de Bono B |
| 2007-02-07 | Reviewed | Mohammadi M |
| 2010-02-03 | Edited | Jupe S |
| 2014-12-04 | Created | Rothfels K |
| 2021-05-31 | Modified | Shorser S |

#### Entities found in this pathway (1)

| Input | UniProt Id |
|---|---|
| FGFR2 | P21802-1, P21802-18, P21802-3, P21802-5 |

### 6. Identifiers found

Below is a list of the input identifiers that have been found or mapped to an equivalent element in Reactome, classified by resource.

##### Entities (73)

| Input | UniProt Id | Input | UniProt Id | Input | UniProt Id |
|---|---|---|---|---|---|
| ANO7 | Q6IWH7 | ARFGAP1 | Q8N6T3 | ARL1 | P40616 |
| ATOH1 | P61086 | B4GALT4 | O60513 | CANT1 | Q08380 |
| CAPN8 | A6NHC0, Q6ZSI9 | CAPN9 | O14815 | CDC42EP5 | Q6NZY7 |
| CLCA1 | A8K7I4 | COG3 | Q96JB2 | COPB2 | P35606 |
| CRACR2A | Q9BSW2 | CREB3L1 | Q96BA8 | CREB3L4 | O43889, Q8TEY5 |
| EDEM3 | Q9BZQ6 | EIF2AK3 | Q9NZJ5 | ENTPD8 | Q5MY95 |
| ERLEC1 | Q96DZ1 | ERN2 | O75460 | FGFR2 | P21802, P21802-1, P21802-18, P21802-3, P21802-5 |
| FOXA3 | P55318 | GALNT12 | Q8IXK2 | GALNT5 | Q7Z7M9 |
| GALNT7 | Q86SF2 | GALNT8 | Q9NY28 | GFPT1 | Q06210 |
| GMPPB | Q9Y5P6 | GNE | Q9Y223 | GOLGA2 | Q08379 |
| GOLGB1 | Q14789 | GORASP2 | Q9H8Y8 | HDLBP | Q00341 |
| HERPUD1 | Q15011 | KCNK6 | Q9Y257 | KCNMA1 | Q12791 |
| KDELR2 | P24390, P33947 | KDELR3 | O43731 | KLF4 | O43474 |
| LMAN1 | P49257, Q9HAT1 | MB | P02144 | MCF2L | O15068 |
| MIA3 | Q5JRA6 | MLLT3 | P42568 | MUC2 | Q02817 |
| MUC4 | Q99102 | MXD4 | Q14582 | NANS | Q9NR45 |
| PDIA5 | Q14554 | PGM3 | O95394 | PLA2G10 | O15496 |
| PMM2 | O15305, Q92871 | PTGER4 | P35408 | RAB27A | P51159 |
| RAP1GAP | P47736 | SCNN1A | P37088 | SEC16A | O15027 |
| SEC24D | O94855 | SH3PXD2A | Q5TCZ1 | SLC16A7 | O60669 |
| SLC2A10 | O95528 | SLC30A7 | Q8NEW0 | SLC35A1 | P78382 |
| SLC39A7 | Q92504 | SLC50A1 | Q9BRV3 | SSR3 | Q9UNL2 |
| ST6GALNAC1 | Q9NSC7 | SYTL1 | Q8IYJ3 | TFF3 | Q07654 |
| TMED3 | Q9Y3Q3 | USO1 | O60763 | WNK4 | Q96J92 |
| XBP1 | P17861-2 | | | | |

| Input | Ensembl Id | Input | Ensembl Id | Input | Ensembl Id |
|---|---|---|---|---|---|
| ARFGAP1 | ENSG00000101199 | FOXA3 | ENSG00000170608 | GFPT1 | ENSG00000198380 |
| HERPUD1 | ENSG00000051108 | KDELR3 | ENSG00000100196 | KLF4 | ENSG00000136826 |
| MXD4 | ENSG00000123933 | PDIA5 | ENSG00000065485 | TFF3 | ENSG00000160180 |
| XBP1 | ENSG00000100219, ENST00000216037, ENST00000344347 | | | | |

### 7. Identifiers not found

These 45 identifiers were not found neither mapped to any entity in Reactome.

| | | | | | | | |
|---|---|---|---|---|---|---|---|
| AGR2 | AL390719.2 | BCAS1 | BLCAP | C3orf52 | DHX32 | DNAJC10 | ERGIC1 |
| FAM107B | FAM110C | FAM174B | FAM83E | FCGBP | FRYL | HES2 | HID1 |
| LARP1B | LRRC31 | MARVELD1 | MLPH | PLXDC2 | PODXL | PRIMPOL | REP15 |
| SEL1L3 | SGSM3 | SLC12A8 | SLC22A23 | SLC39A11 | SLC9A3R2 | SMIM14 | SPDEF |
| SPINK4 | SYTL2 | TACC1 | TAGLN | TCEA3 | TM9SF3 | TMEM184A | TMEM214 |
| TPSG1 | TSPAN13 | TTC39A | URAD | ZG16 | | | |
