## Supplementary material for "A proximal-to-distal survey of healthy adult human small intestine and colon epithelium by single-cell transcriptomics": File S5

### Supplemental Figure
### Receptor Families by Cell Lineage

### Voltage-Gated Ion Channels

#### High Expression (>0.5)

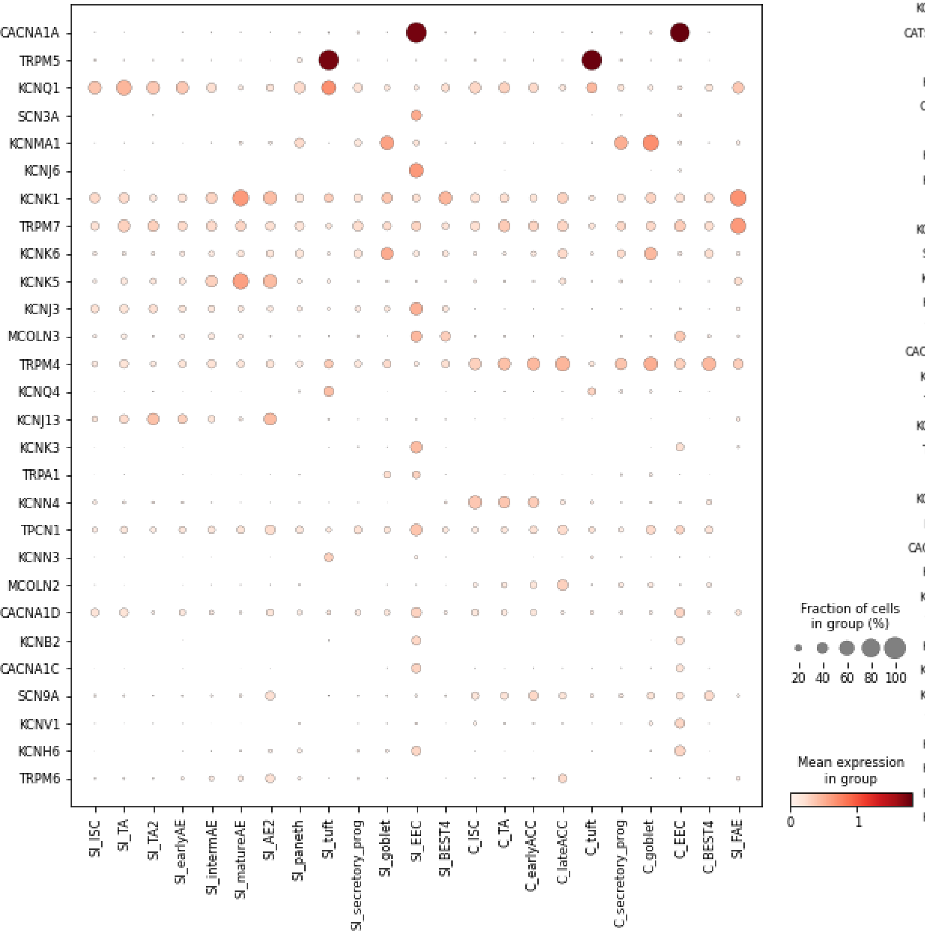


#### Mid Expression (0.05 - 0.5)

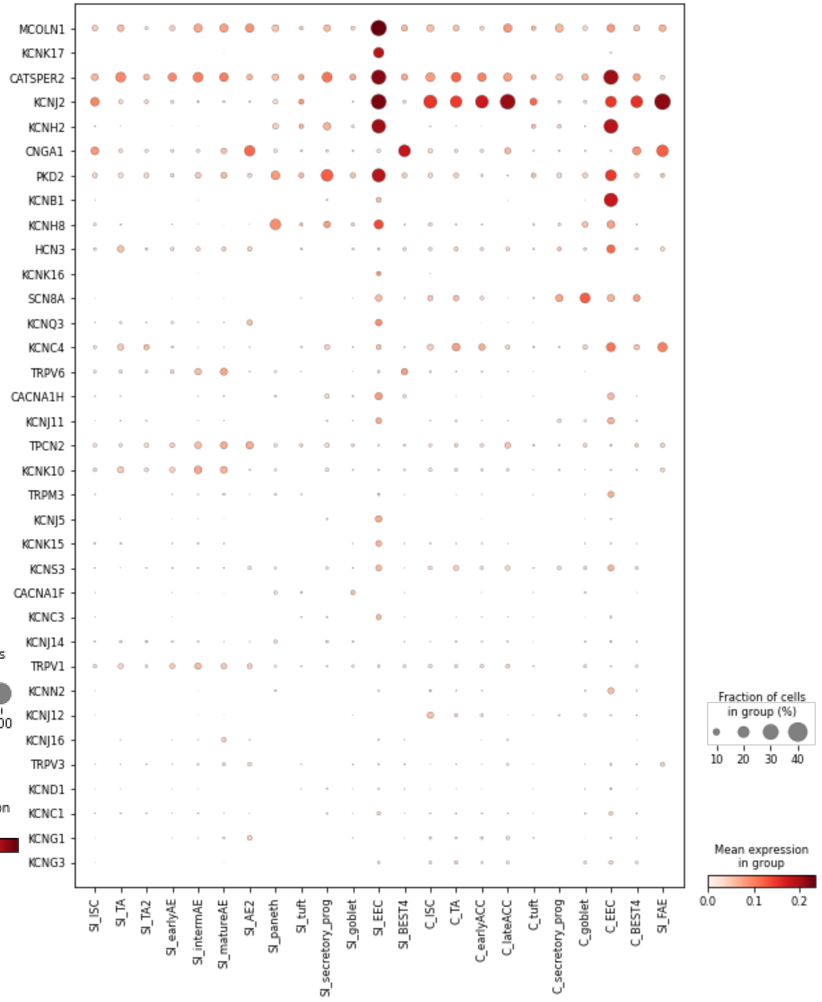


#### Low Expression (<0.05)

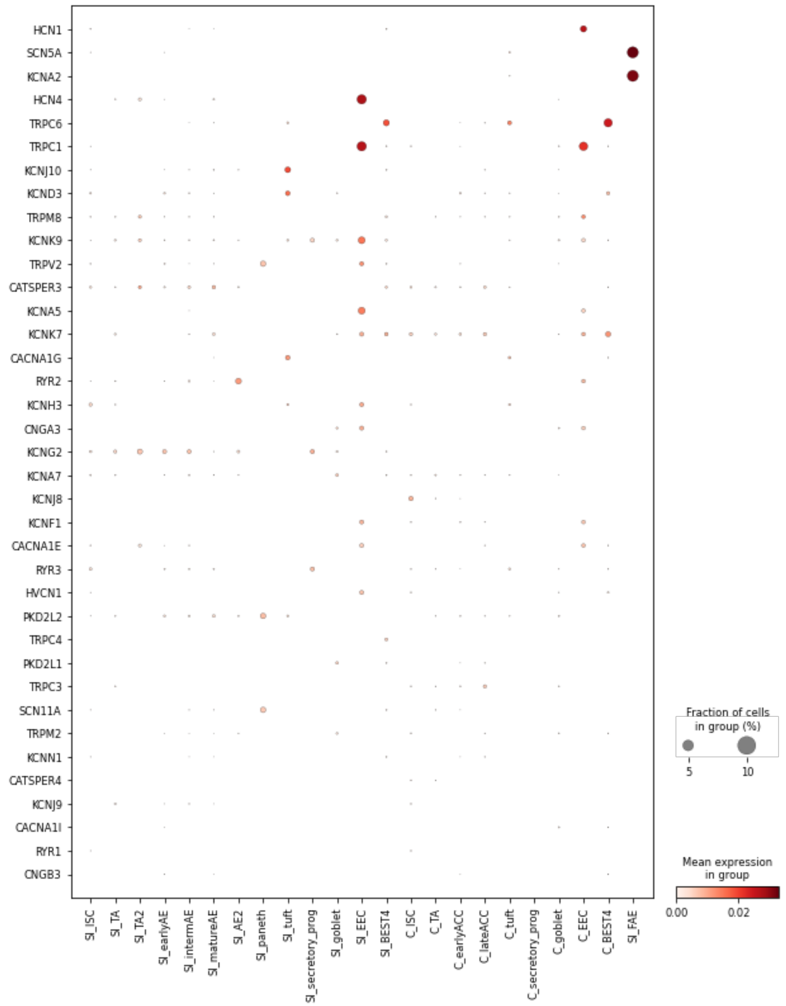

### Ligand-Gated Ion Channels

#### High Expression (>0.5)

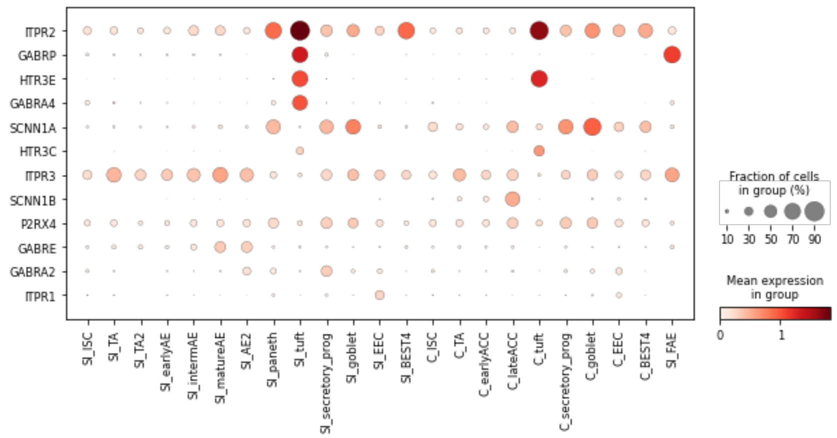


#### Mid Expression (0.05 - 0.5)

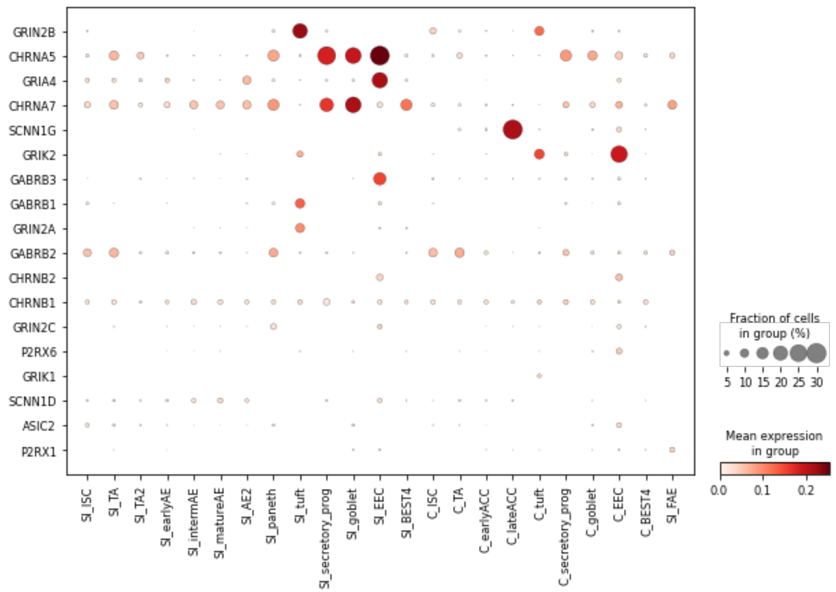


#### Low Expression (<0.05)

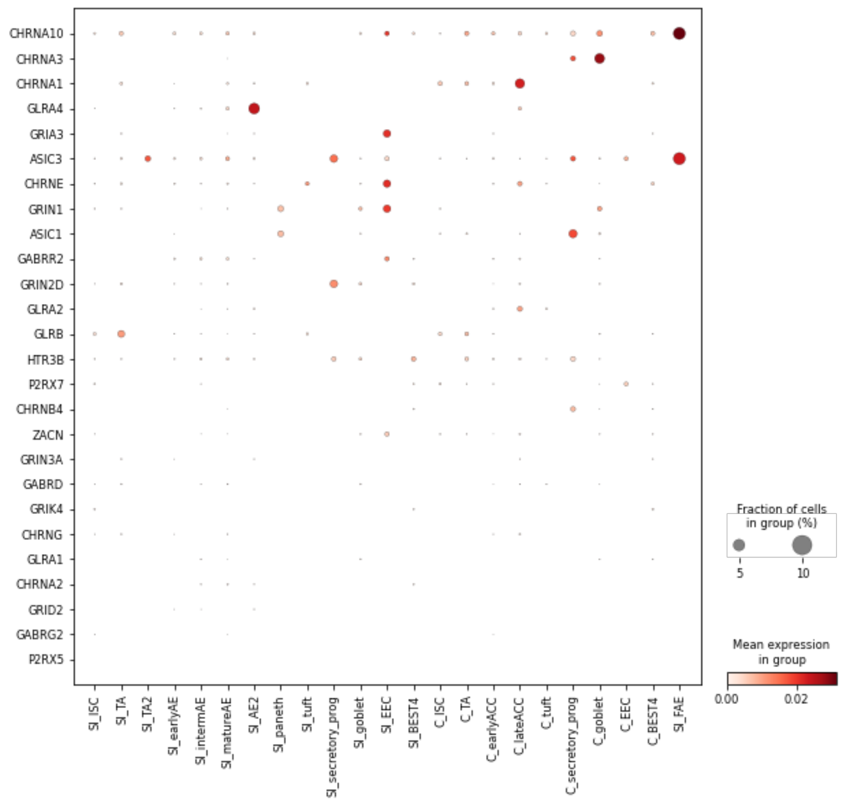

### Other Ion Channels

#### High Expression (>0.5)

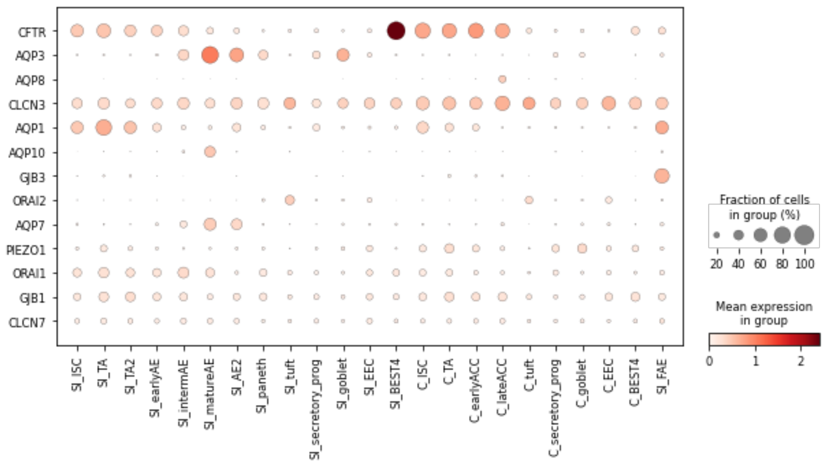


#### Mid Expression (0.05 - 0.5)

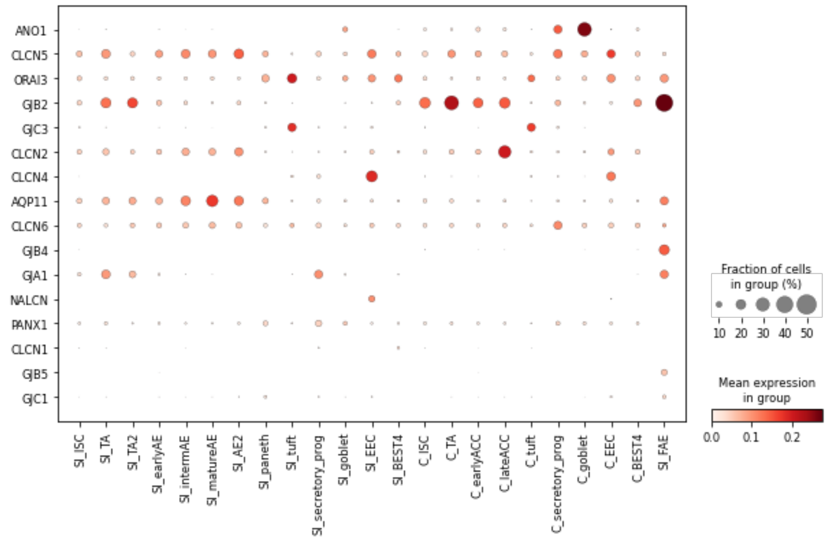


#### Low Expression (<0.05)

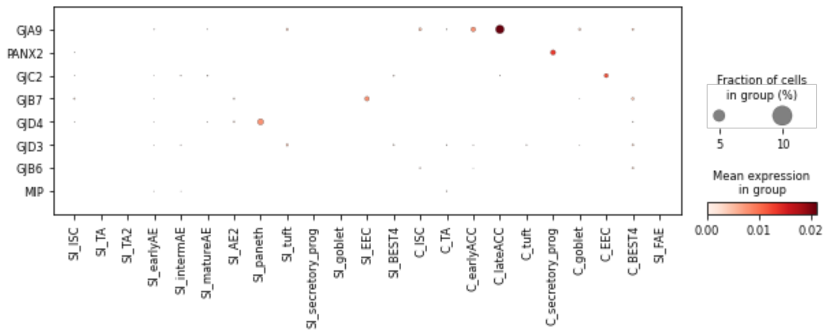

### G-Protein Coupled Receptors

#### High Expression (>0.5)

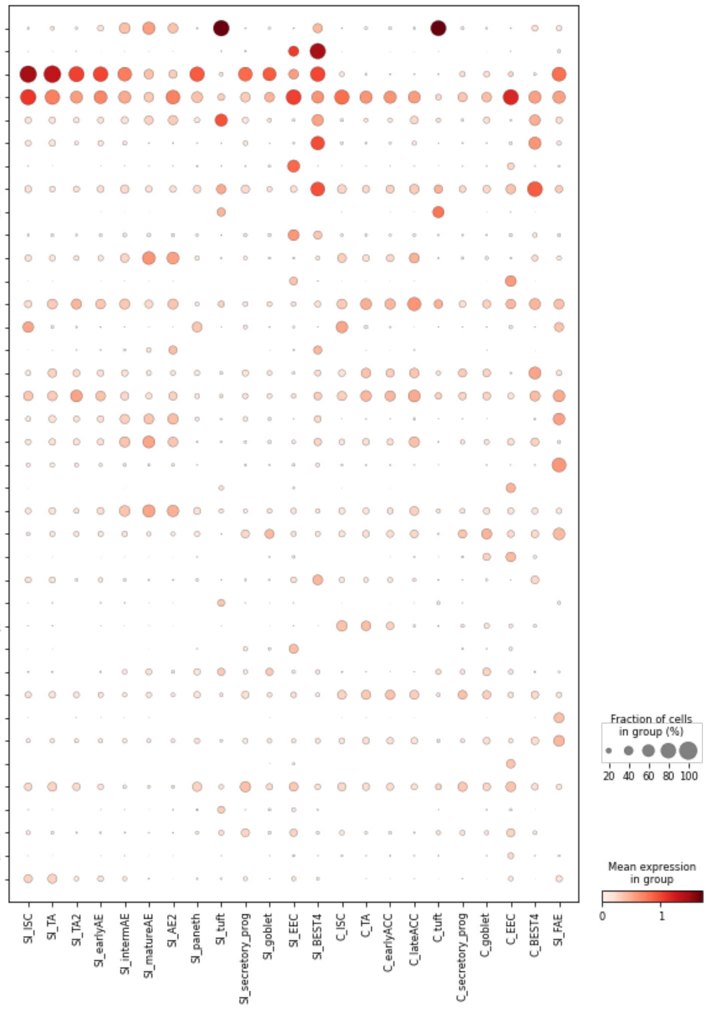


#### Mid Expression (0.05 - 0.5)

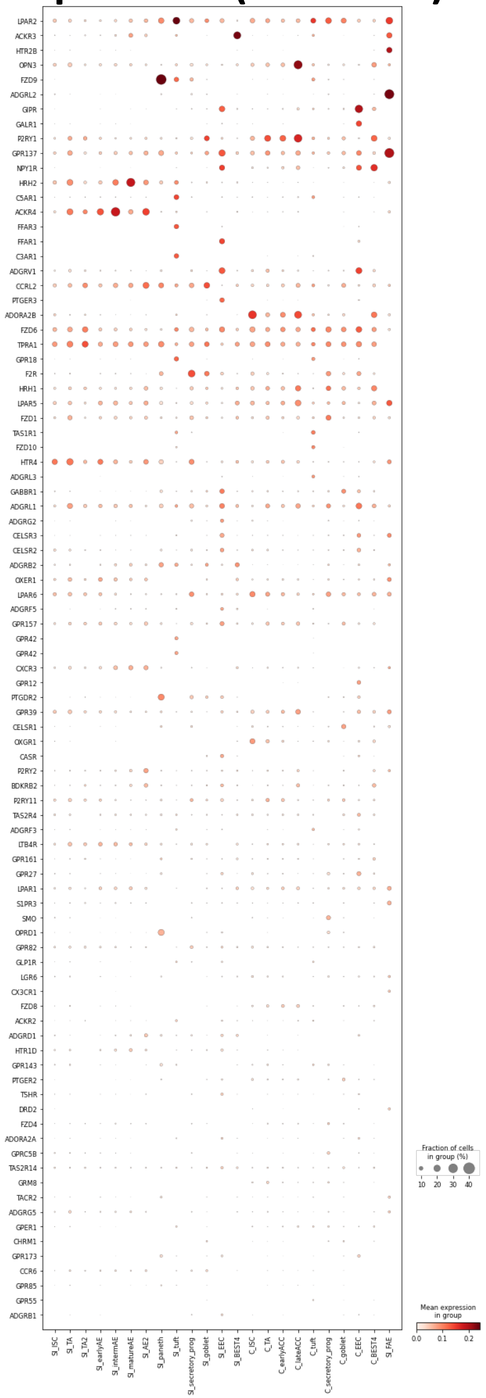

#### Low Expression (<0.05)

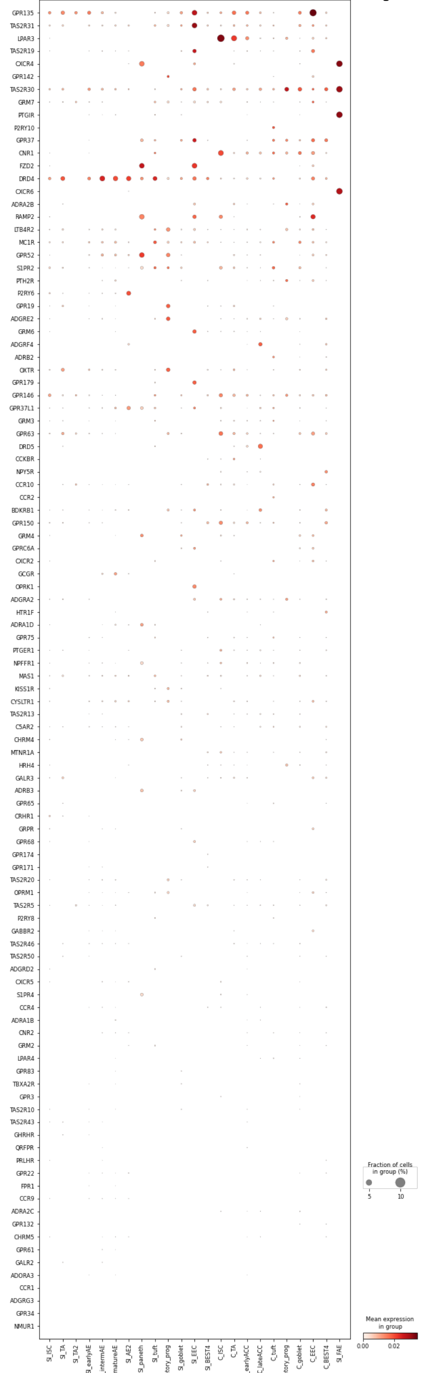

Nuclear Hormone Receptors

#### High Expression (>0.5)

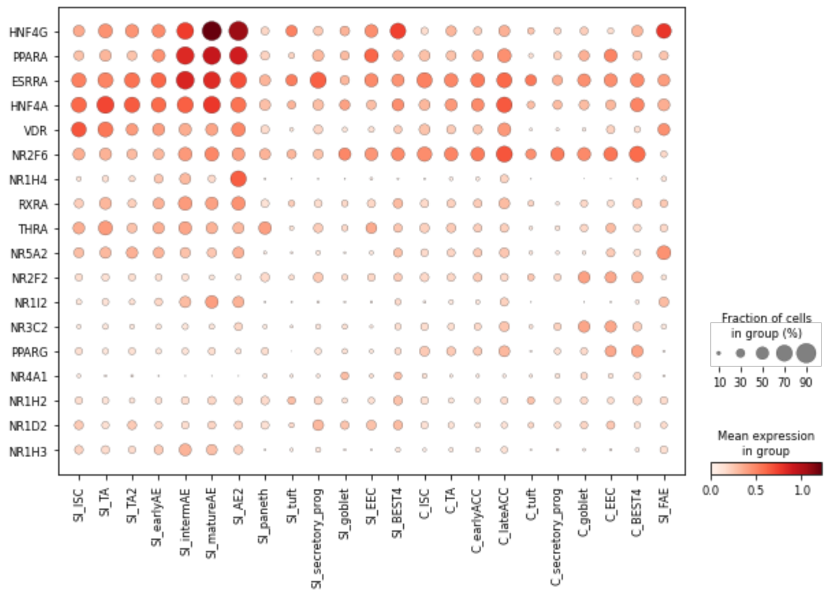


#### Mid Expression (0.05 - 0.5)

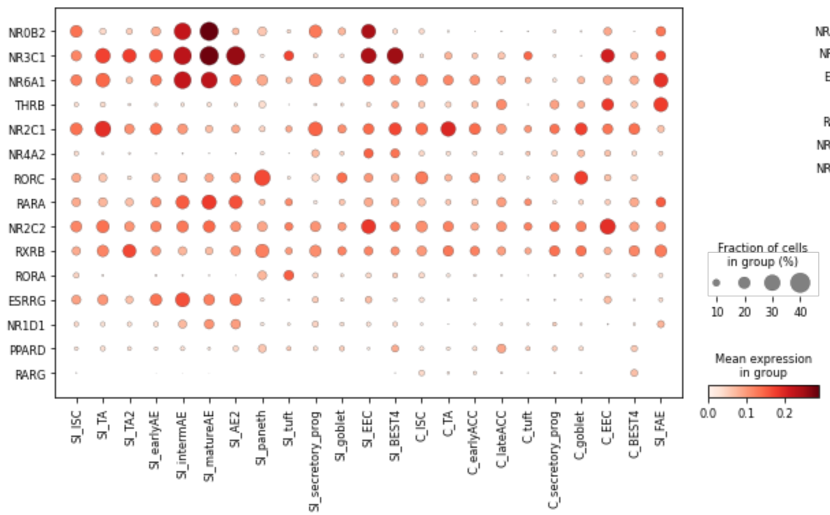


#### Low Expression (<0.05)

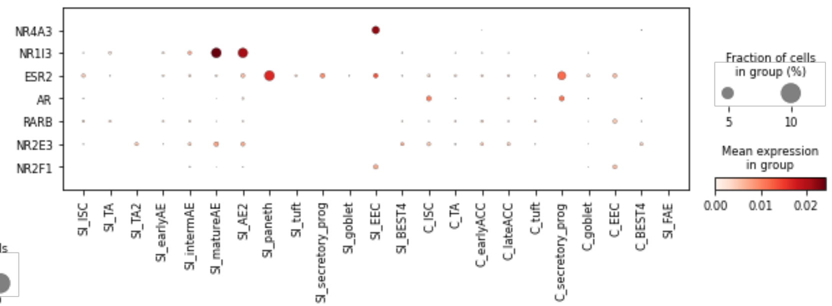

### Interleukin Receptors

#### High Expression (>0.5)

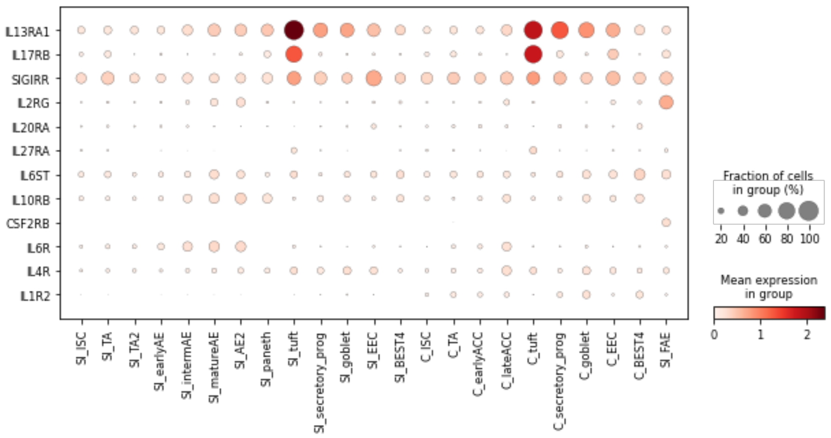


#### Mid Expression (0.05 - 0.5)

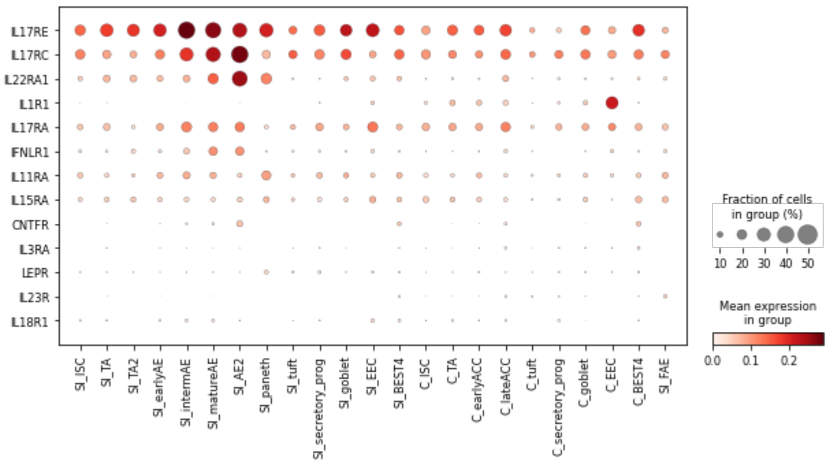


#### Low Expression (<0.05)

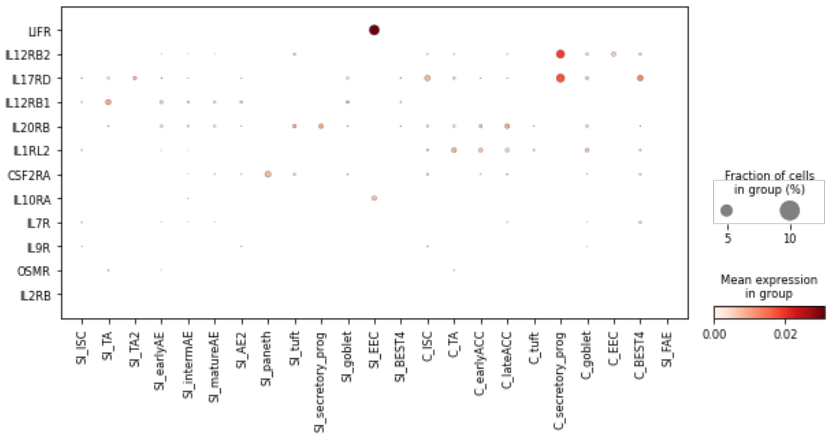

Integrins

**High Expression (>0.5)**

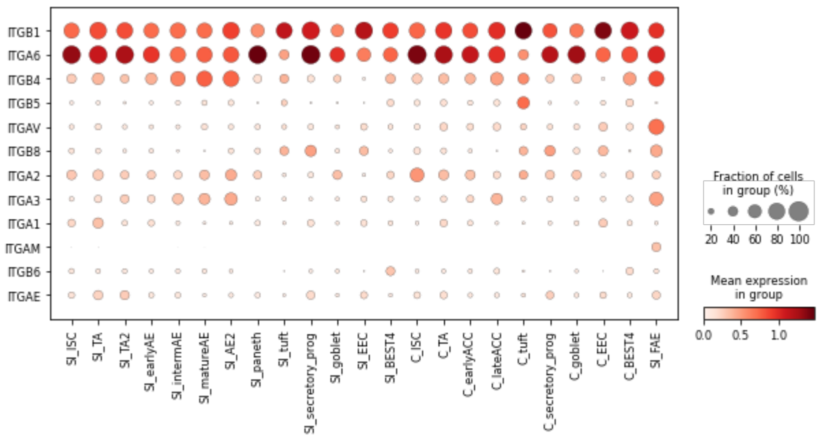


**Mid Expression (0.05 - 0.5)**

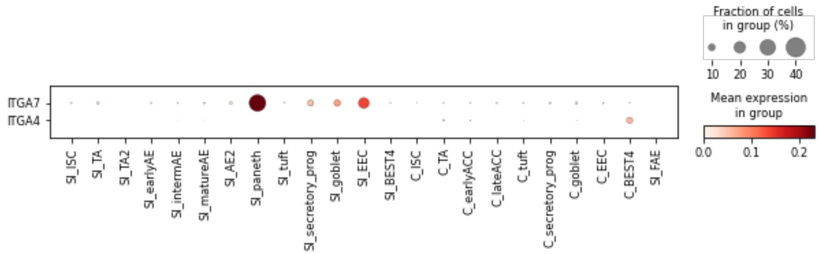


**Low Expression (<0.05)**

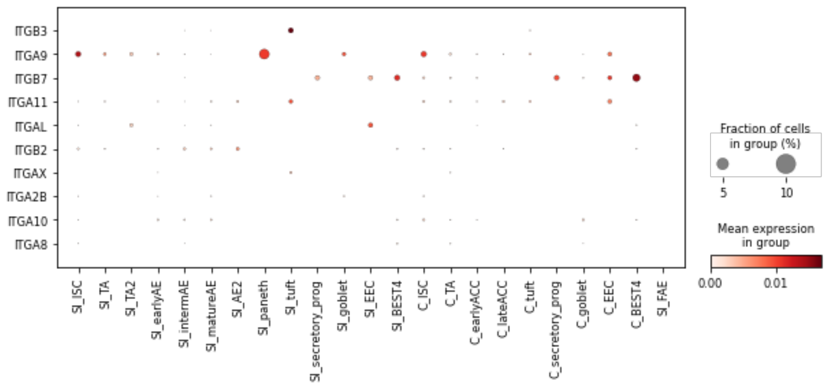

### Toll-Like Receptors

#### High Expression (>0.5)

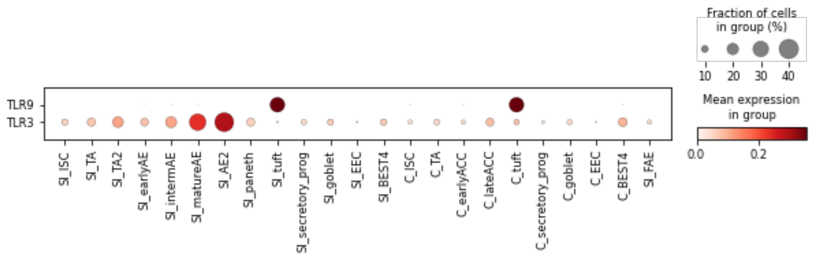


#### Mid Expression (0.05 - 0.5)

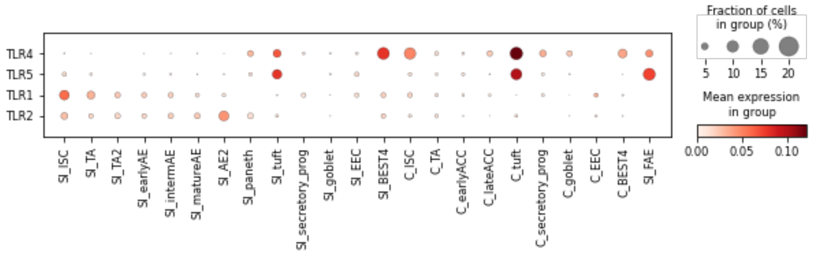


#### Low Expression (<0.05)

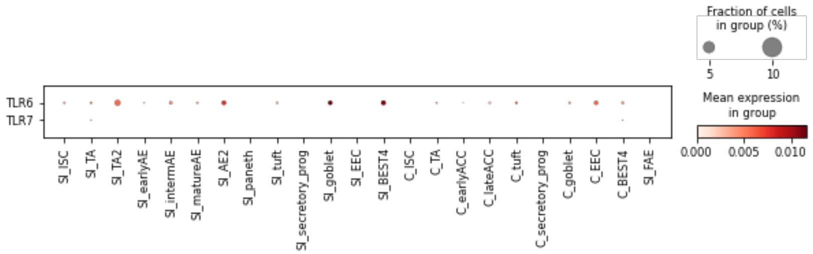

### Receptor Tyrosine Kinases

#### High Expression (>0.5)

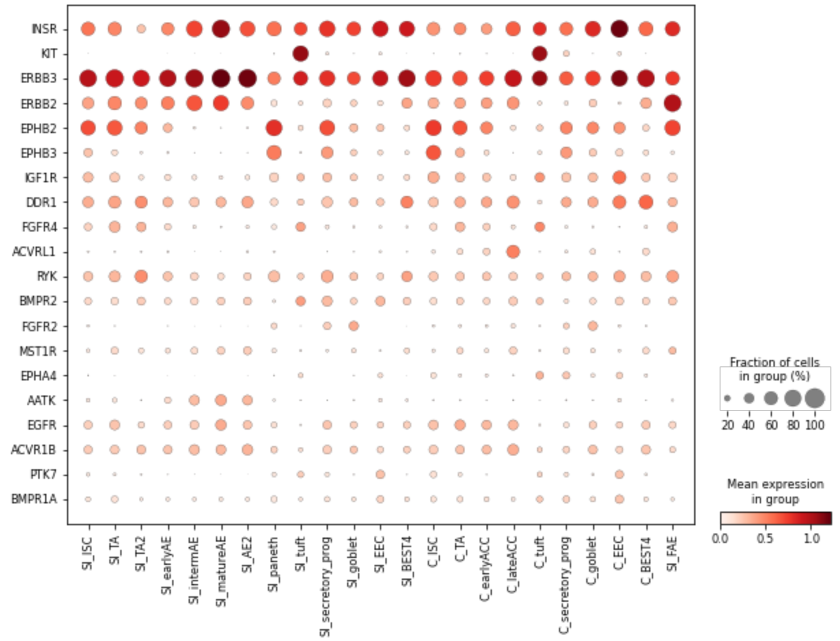


#### Mid Expression (0.05 - 0.5)

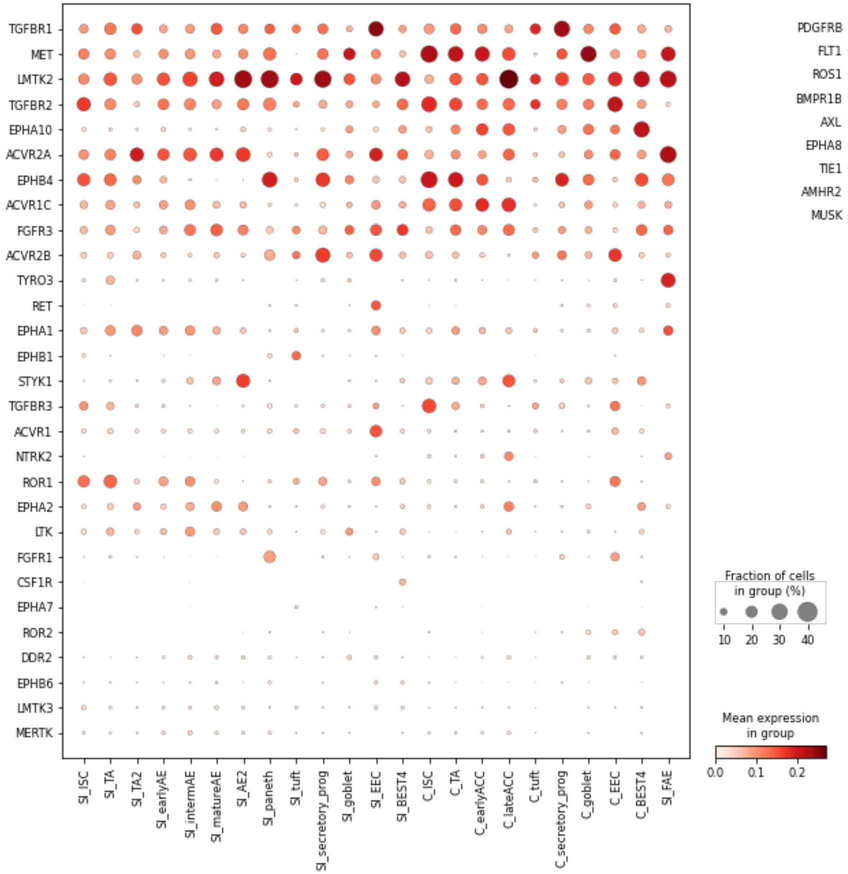


#### Low Expression (<0.05)

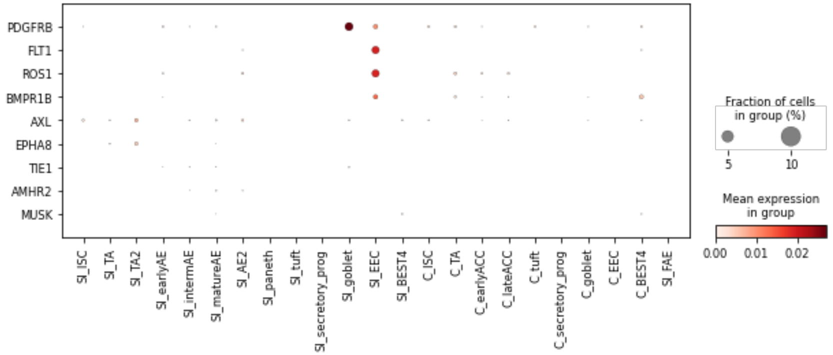

Receptor Tyrosine Phosphatases

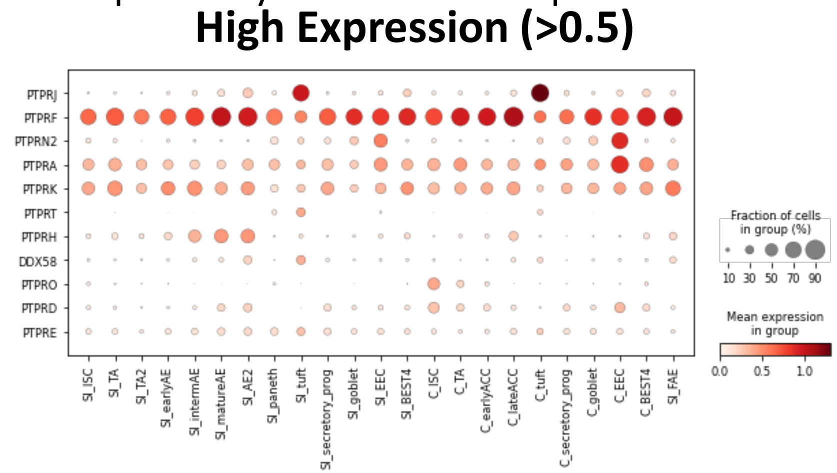


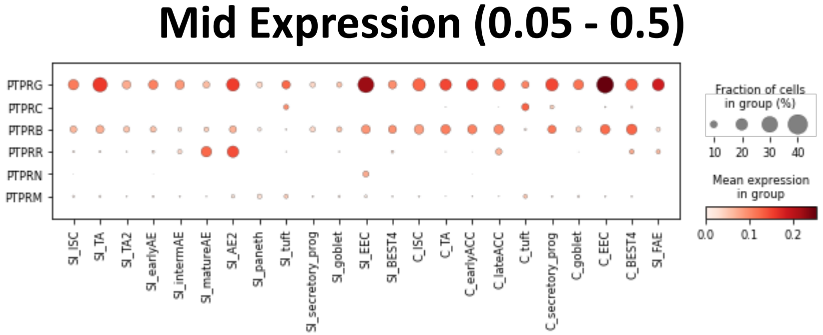


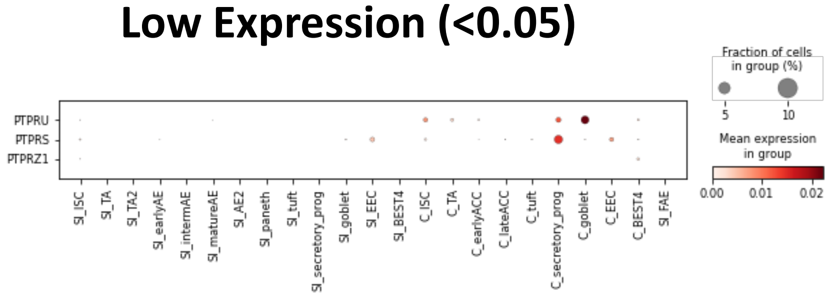

### TNF Receptors

#### High Expression (>0.5)

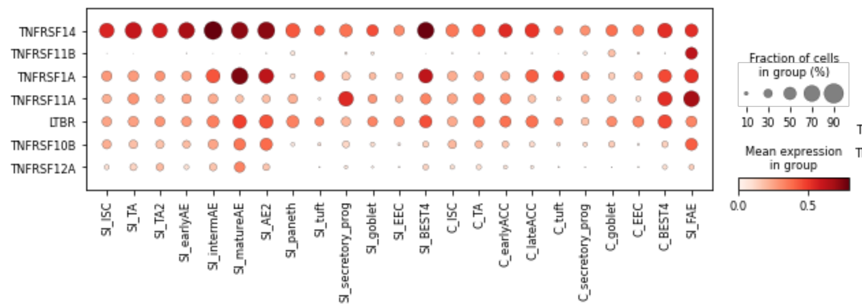


#### Mid Expression (0.05 - 0.5)

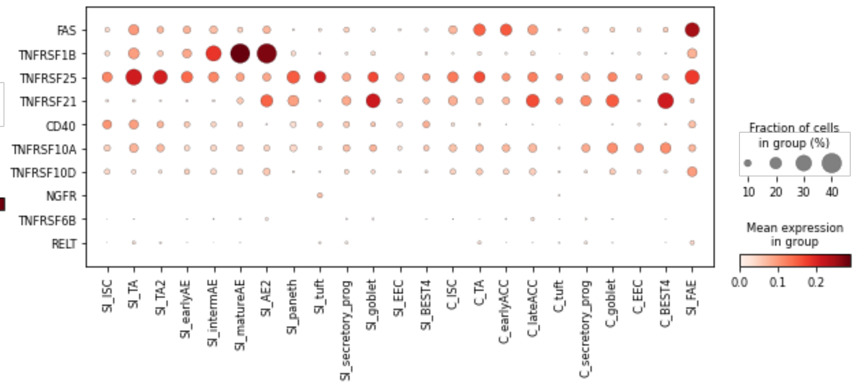


#### Low Expression (<0.05)

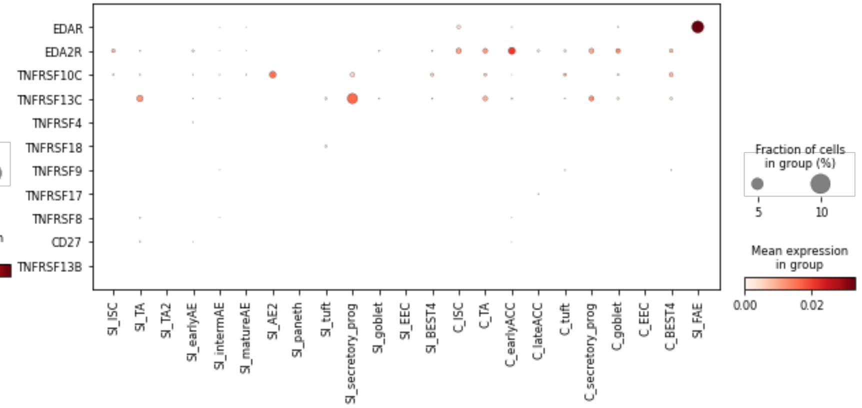

### NOD-Like Receptors

#### High Expression (>0.5)

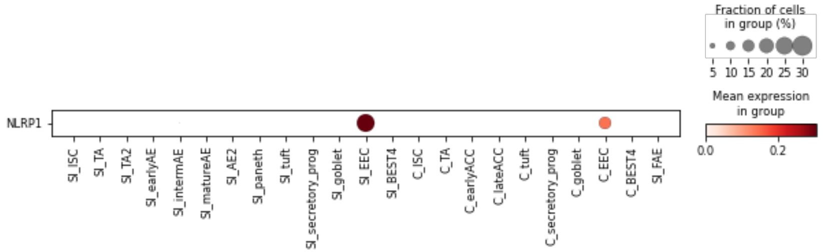


#### Mid Expression (0.05 - 0.5)

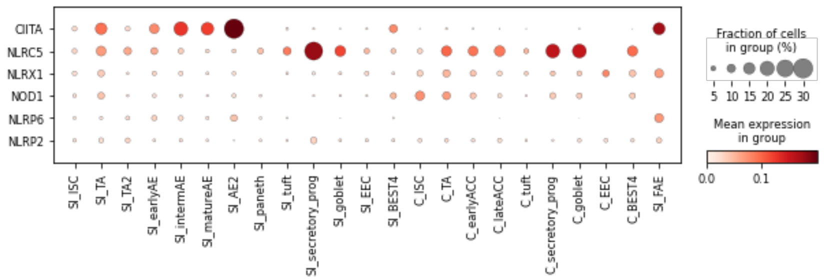


#### Low Expression (<0.05)

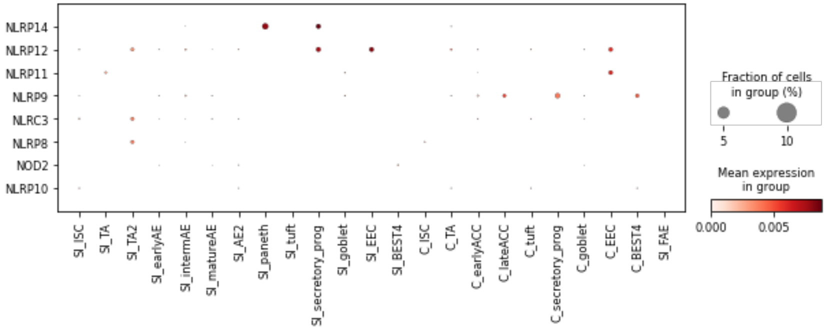

### Small Catalytic Receptor Families

#### High Expression (>0.5)

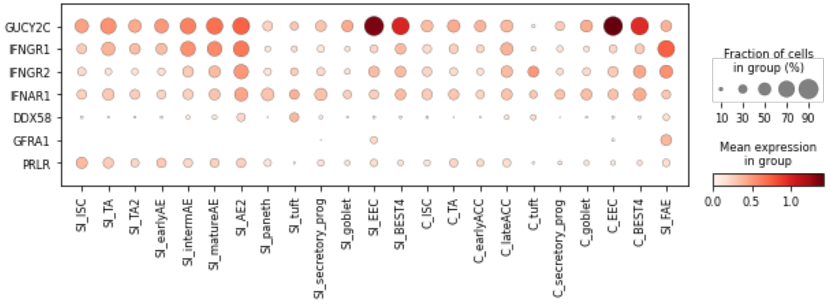


#### Mid Expression (0.05 - 0.5)

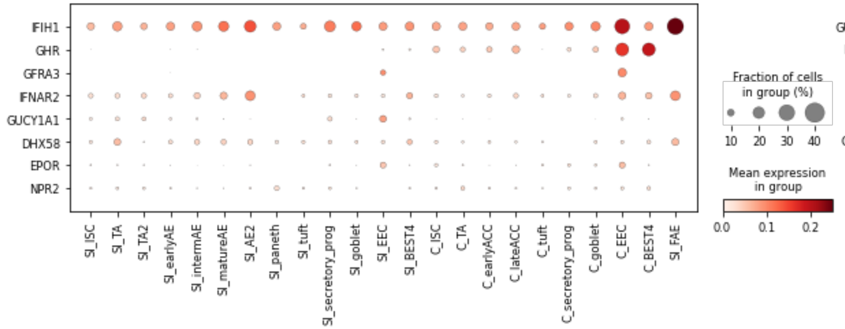


#### Low Expression (<0.05)

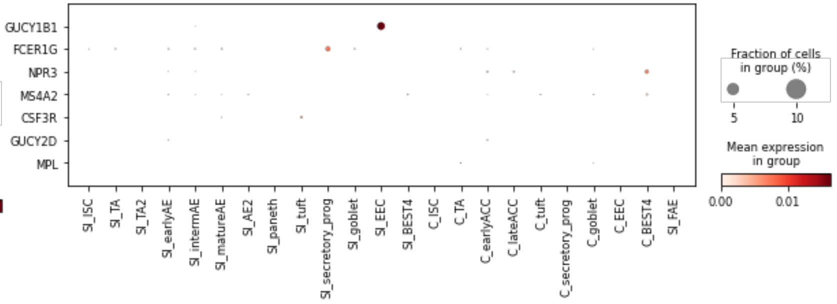
